# Mapping spatially organized molecular and genetic signatures of schizophrenia across multiple scales in human prefrontal cortex

**DOI:** 10.64898/2026.02.16.706214

**Authors:** Sang Ho Kwon, Boyi Guo, Cindy Fang, Madhavi Tippani, Svitlana V. Bach, Geo Pertea, Ryan A. Miller, Sarah E. Maguire, Artemis Iatrou, Nicholas J. Eagles, Madeline R. Valentine, Seyun Oh, Aarti Jajoo, Vijetha Balakundi, Yufeng Du, Annie B. Nguyen, Ruth Zhang, Uma M. Kaipa, Heena R. Divecha, Jashandeep S. Lobana, Joel E. Kleinman, Shizhong Han, Nikolaos P. Daskalakis, Thomas M. Hyde, Leonardo Collado-Torres, Stephanie C. Page, Kristen R. Maynard, Stephanie C. Hicks, Keri Martinowich

## Abstract

The dorsolateral prefrontal cortex (dlPFC) is central to cognitive dysfunction in schizophrenia (SCZ), yet how molecular changes are organized across cortex remains unclear. Here, we applied complementary spatial transcriptomic approaches spanning laminar domains, microenvironments, and cell types in postmortem human dlPFC. At the laminar level, SCZ-associated transcriptional changes were strongest in glia-enriched domains (layer 1/meninges and white matter), including down-regulation of microglia-associated genes, whereas genetic risk localized to neuronal-rich gray matter. We next analyzed SCZ-linked microenvironments including neuropil, neuronal, perineuronal net, and vascular compartments. Within these, neuronal and synaptic transcriptional changes were most prominent in neuropil, showing down-regulation of activity-dependent synaptic genes and inhibitory neuron markers. At the cellular level, these signals reflected intrinsic alterations localized to cell types. Across analyses, patterns converged on altered BDNF-TrkB signaling and inhibitory circuit dysfunction. Together, our findings highlight spatial scale as a key determinant in resolving neuronal and non-neuronal aspects of SCZ-associated biology.

## 1 Introduction

Schizophrenia (SCZ) is a chronic neuropsychiatric disorder affecting approximately 0.3-1% of the global population.^1–4^ SCZ is characterized by a diverse spectrum of symptoms that disrupts thoughts, emotions, and behavior. Symptoms are generally classified as positive, including hallucinations and delusions; negative, including social withdrawal and blunted affect; or cognitive, including deficits to working memory, executive function, and attention.^5,6^ The cognitive symptoms are particularly challenging because they have a substantial impact on daily functioning and are correlated with poorer quality of life, but remain largely unaddressed by existing treatments.^7,8^ These cognitive deficits have been linked to dysfunction in the dorsolateral prefrontal cortex (dlPFC).^5^ This region has undergone considerable evolutionary expansion in non-human primates and humans,^9^ and is critical for higher-order cognitive functions, including working memory, planning, and cognitive flexibility. As a neocortical structure, the dlPFC has six distinct layers, which contain differing proportions of neuronal and glial subtypes. These cell types are spatially and functionally organized into specialized microcircuits, which regulate complex cognitive processing and provide top-down control over subcortical structures.^8,10^ Neuroimaging and postmortem studies consistently suggest abnormalities in the dlPFC of individuals with SCZ,^5,11,12^ but the exact mechanisms underlying these disruptions remain poorly understood. Integration of both bulk and single-nucleus transcriptomic approaches with genome-wide association studies (GWAS) has provided some molecular-genetic evidence that SCZ vulnerability is preferentially enriched within specific cortical layers and cell types in the human prefrontal cortex, including the dlPFC.^4,5,13–17^ However, the cellular and/or spatial resolution of these approaches has limited the ability to finely localize risk to specific laminar compartments and resident cell types that comprise these microcircuits. Rapidly emerging multi-modal approaches that incorporate spatially-resolved transcriptomics (SRT) can overcome these limitations.^12,18^

We used the Visium and Xenium SRT platforms (10x Genomics) to profile gene expression in postmortem human dlPFC tissue from individuals diagnosed with SCZ compared to neurotypical controls (NTCs). We identified SCZ-associated molecular signatures linked to synaptic biology, immune signaling, and GABAergic signaling, which we mapped to discrete cortical layers and spatially-localized cell types. We extended our analyses by integrating protein detection with the SRT data to investigate how SCZ-associated gene expression varies across cellular microenvironments. Specifically, we analyzed microenvironments that have been linked with SCZ including compartments enriched for neurons, neuropil, perineuronal nets (PNNs), or vasculature. PNNs preferentially surround parvalbumin (PVALB) inhibitory neurons, which regulate synchronization of gamma oscillations that are altered in individuals with SCZ.^19,20^ PNNs maintain synapses localized to neuronal cell bodies and proximal dendrites, and regulate synaptic plasticity, which is disrupted in SCZ.^19–21^ Neurovasculature is linked to SCZ pathobiology, and directly interacts with immune signaling to regulate synaptic homeostasis.^22,23^

This integrative multi-omic strategy allowed us to contextualize how SCZ-associated signaling pathways are impacted across spatially-localized cortical domains, cellular organization, and local microenvironments. The findings underscore the value of spatially-resolved approaches to identify and localize novel molecular targets or refine our understanding of known targets. To provide broad access to the scientific community, we share both raw and processed data, along with the code used for our analyses. We also developed user-friendly web resources that allow researchers to explore and visualize the SRT data.

## 2 Results

### 2.1 Cross-diagnostic identification of data-driven spatial domains and cellular microenvironments in control and SCZ human dlPFC

Using the Visium platform, we generated SRT data from postmortem human dlPFC (Brodmann area 46) in 32 donors diagnosed with SCZ and 31 age- and sex-matched NTCs (**Figure 1A**, **Supplementary Table 1**). dlPFC tissue samples were dissected perpendicular to the pial surface and spanned cortical layers (L) 1-6 and white matter (WM) (**Figure 1B**). Instead of performing standard hematoxylin and eosin (H&E) staining, tissue sections were immunofluorescently (IF) labeled against DAPI, wisteria floribunda agglutinin (WFA), NeuN and Claudin-5, to identify cellular microenvironments associated with neuropil, PNNs, neurons, and vasculature, respectively (**Figure 1B**) for spatial proteogenomic (SPG) profiling across defined microenvironments. Following quality control (QC), we retained 279,806 high-quality spots (**Figure S1**-**Figure S3**) that were comparable across groups (NTC: 136,456; SCZ: 143,350) (**Figure S4**). Expression of *SNAP25* (neurons), *MBP* (oligodendrocytes), and *PCP4* (L5 marker) verified intact cortical structure across samples (**Figure 1B**).

**Figure 1.**
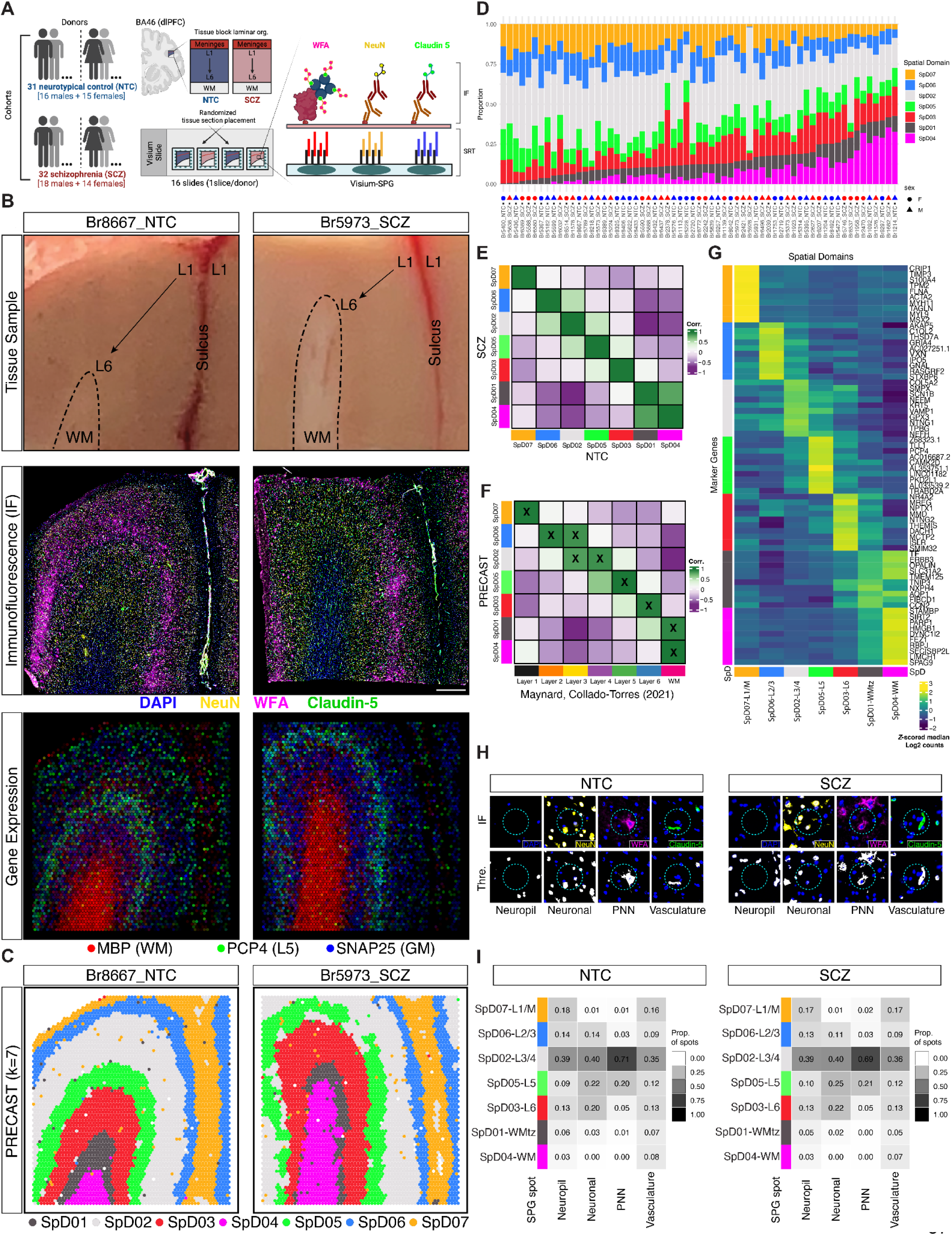
Overview of data collection and downstream processing to identify spatial domains (SpDs) corresponding to cortical layers and cellular microenvironments in the human dlPFC. (A) Schematic of the cohort and experimental workflow integrating Visium SRT and IF images to generate SPG data. Created in BioRender. Kwon, S. H. (2026) https://BioRender.com/nryjogo (B) Overview of SPG data for NTC and SCZ samples where Br8667 and Br5973 indicate donor labels, respectively. (**Top**) Tissue block images showing laminar orientation and morphology (L for cortical layers; WM for white matter). (**Middle**) IF images highlighting cellular components and microenvironments (DAPI marking nuclei in blue, NeuN marking neuronal nuclei in yellow, WFA marking PNNs in pink, and Claudin-5 marking blood vessels in green). Scale bar: 1 mm. (**Bottom**) Spot plots displaying log-transformed normalized expression (logcounts) of *SNAP25*, *MBP*, and *PCP4*, outlining gray matter, WM, and cortical layer 5, respectively. (C) Unsupervised clustering using PRECAST delineated SpDs that recapitulate the characteristic laminar organization of the human dlPFC at *k*=7. (D) Proportion of Visium spots assigned to each PRECAST SpD at *k*=7 in all donors across sex and diagnosis. (E) Correlation of enrichment *t*-statistics profiles between NTC and SCZ samples across PRECAST SpDs, demonstrating similar laminar organization. Heatmap shows correlation values (Corr.). (F) Correlation of enrichment *t*-statistics of PRECAST SpDs with manual histological layer annotations.^26,27^ Heatmap shows correlation values (Corr.), with higher-confidence annotations marked by “X”. (G) Top 10 most highly expressed marker genes for each PRECAST SpD at *k*=7. Each SpD is denoted as SpD-L where “L” represents the most confidently approximated cortical layer for a given PRECAST SpD, using the spatial registration approach shown in (**F**). The color scale of the heatmap represents the median of normalized log2 count across all samples followed with *z*-score transformation with respect to all SpDs (i.e. row-wise) (H) Classification of Visium spots (cyan circles, 55-µm diameter) into cellular microenvironments (SPG spots) based on enrichment of non-nucleated neuropil (DAPI-free, absence of blue), neuronal bodies (NeuN, yellow), PNNs (WFA, pink), and vasculature (Claudin-5, green) in NTC and SCZ samples. SPG spots are classified into four non-mutually exclusive microenvironments, allowing a single spot to belong to multiple SPG types. (**Top**) Visium spots overlaid on raw IF images. (**Bottom**) Segmentation of respective IF signal (white) was thresholded (Thre.) to classify Visium spots. (I) Heatmaps showing distribution of SPG-defined tissue microenvironments (neuropil, neuronal, PNN, and vasculature microenvironments) across PRECAST SpDs at *k*=7. Each row represents a single SpD and each column represents an SPG-defined tissue microenvironment. Values indicate the proportion of SPG spots that are localized to each SpD (column-wise summing to 1). These patterns show consistent laminar localization of the microenvironments in both NTC and SCZ samples. The higher proportions observed in L3/4 are likely due to the larger size of this SpD. A complementary within-SpD assessment is provided in **Figure S12**.

We spatially clustered all SRT samples with PRECAST^24^ using as input the top 100 marker genes for human cortical layers reported by Huuki-Myers et al..^25^ We evaluated various clustering resolutions (*k*=2-16) (**Figure S5**), and used the spatial domains (SpDs) generated with *k*=7 for downstream analysis because these seven contiguous SpDs best corresponded to the classic histological cortical layers (**Figure 1C**, **Figure S6**). Supporting our decision to cluster across all samples, the majority of variation in gene expression at *k*=7 was attributed to SpD compared to other factors (e.g., slide id, age, or diagnosis) (**Figure S7**). We annotated the SpDs by adapting a spatial registration approach,^25–27^ where gene expression profiles from our seven SpDs were compared against those derived from the manually annotated human cortical layers from Maynard, Collado-Torres et al.^27^ (**Figure 1F**). Each SpD was assigned to a canonical dlPFC layer, denoted as SpDX-L, where X represents the domain number and L represents the layer annotation: SpD07-L1/M, SpD06-L2/3, SpD02-L3/4, SpD05-L5, SpD03-L6, SpD01-WM, and SpD04-WM. For SpD07-L1/M, the M indicates inclusion of leptomeningeal structures, particularly vasculature, within this domain. The proportion of spots for each SpD was balanced across diagnosis (**Figure 1D**), and there was strong correlation in gene expression across all seven SpDs between diagnosis (**Figure 1E**). Of the two SpDs associated with WM, SpD01-WM was anatomically positioned in between the deepest gray-matter domain (SpD03-L6) and the other WM domain (SpD04-WM). Analysis of the top 10 marker genes, identified using the enrichment test (**Methods**), for SpD01-WM revealed co-enrichment of representative genes associated with neuronal functions (*CCN2*, *FIBCD1*, *NXPH4*) and oligodendrocyte functions (*TMEM125*, *SLC31A2*, *OPALIN*) (**Figure 1G**). Differential expression (DE) analysis between SpD01-WM and SpD04-WM revealed that SpD01-WM had higher expression of neuronal marker genes, including *SNAP25* and *SLC17A7*, and lower expression of oligodendrocyte genes, including *PLP1*, *MBP*, and *MOBP* (**Figure S8**, **Figure S9**, **Supplementary Table 2**). Among the neuronal genes enriched in SpD01-WM, we identified genes associated with deep layer neurons in L6, including *CCN2*, *CPLX3*, and *NR4A2*.^28^ Given its anatomical positioning at the border of the WM as well as mixed neuronal and glial transcriptomic profiles, we refined the annotation for SpD01-WM to WM transition zone (SpD01-WMtz).

To annotate Visium spots based on their inclusion within a cellular microenvironment, we segmented the high-resolution IF images of the immunostained tissue to identify regions of interest (ROI) for each IF channel (**Methods**, **Figure S10**, **Figure S11**). In each IF channel, we further classified Visium spots into one of the two categories based on spot coverage by segmented ROIs, namely SPG and non-SPG spots (**Figure 1H**, **Methods**).^25,29^ Specifically, spots were classified as ‘neuropil’/‘cell body’ per the DAPI channel, ‘neuronal’/‘non-neuronal’ per the NeuN channel, ‘PNN’/‘non-PNN’ per the PNN channel, and ‘vasculature’/‘non-vasculature’ per the Claudin-5 channel. These classifications were independent of each other such that a spot could be positive for more than one feature (e.g., both neuronal and PNN). Among all QC’ed spots (n=279,806 spots), 74,000 spots contained only DAPI signals and were classified as non-SPG across all channels (**Figure 1H**, **Figure S10**, **Figure S11**, **Methods**). Analyzing across all samples, we identified SPG spots enriched for neuropil (NTC: 71,804; SCZ: 76,161), neuronal (NTC: 28,957; SCZ: 26,060), PNN (NTC: 7,084; SCZ: 5,249), and vasculature (NTC: 10,708; SCZ: 11,198).

We examined the distribution of SPG spot types across SpDs in both NTC and SCZ samples (**Figure 1I**). For all four IF channels, the respective SPG spots were localized to SpD02-L3/4 more than any other SpDs, likely reflecting the large size of this SpD (**Figure 1D**). Neuropil spots were detected across all SpDs, with the highest enrichment in SpD02-L3/4 and SpD07-L1/M and lowest enrichment in the WM SpDs (SpD01-WMtz and SpD04-WM). Neuronal spots were primarily localized to gray matter (SpD06-L2/3 to SpD03-L6). We also observed a small proportion of neuronal spots localized to SpD01-WMtz (NTC: 756 spots, 2.6%; SCZ: 479 spots, 1.8%), potentially reflecting deep cortical neurons, which is consistent with the relatively elevated neuronal gene expression in SpD01-WMtz compared to SpD04-WM (**Figure S9**). PNN spots were enriched in SpD02-L3/4. Since PNNs preferentially surround PVALB inhibitory neurons, this data is consistent with enrichment of PVALB neurons in L4.^30^ Vasculature spots were broadly distributed across SpDs in both NTC and SCZ, with enrichment in SpD02-L3/4 (NTC: 3,744 spots, 34.9%; SCZ: 4,087 spots, 36%) and SpD07-L1/M (NTC: 1,717 spots, 16.0%; SCZ: 1,896 spots, 16.9%) (**Figure 1I**). We also quantified the proportion of SPG spot types within individual SpDs across NTC and SCZ (**Figure S12**). Among SPG spot types in SpD07-L1/M, vasculature spots showed the highest proportion after neuropil spots, potentially due to the contribution of leptomeningeal vasculature in the sulcal regions adjacent to L1 (**Figure S12**, **Figure 1B**).^31^ Neuropil spots, including the spots in WM, consistently accounted for the largest proportion in nearly all domains (**Figure S12**). This pattern may reflect the high density of neuropil, whose composition includes synaptic connections, glial processes, and extracellular matrix components, which is particularly abundant in human cortical regions that are enriched for dendritic and axonal arborizations.^32–35^

To validate our classification approach, we assessed the molecular composition of the annotated cellular microenvironments in the NTC samples by analyzing gene expression in pseudobulked SPG spots compared to non-SPG spots from the corresponding microenvironment (**Figure S13**, **Supplementary Table 3**, **Methods**). Neuropil spots, compared to non-neuropil spots, were enriched for synaptic genes including *CAMK2A*, *DLG4*, *MAP2*, and *SHANK1-3* (**Figure S13A**). Neuronal spots were enriched for neuronal markers including *SNAP25* and *RBFOX3* (**Figure S13B**). PNN spots were enriched for *PVALB* and proteoglycan-related genes (*HAPLN1* and *HAPLN4*), as well as *ERBB4*, a receptor tyrosine kinase expressed in PVALB neurons.^30,36,37^ These PNN spots were also depleted of metalloproteinase genes (*MMP16* and *MMP28*) (**Figure S13C**), which function as extracellular matrix (ECM) remodeling enzymes that degrade PNN components and regulate their structural maintenance.^20,38^ Vasculature spots were enriched for genes associated with endothelial junctions and immune response, including *CLDN5*, *PECAM1*, and *A2M* (**Figure S13D**).

We next leveraged the gene expression data to cross-validate the image-based annotation of SPG spots. Using data from existing transcriptomic studies, we compiled marker gene sets for neuropil, cortical neurons, PNN-ensheathed PVALB inhibitory neurons, and vascular cells.^25,30,39,40^ We annotated the SPG spot types using the compiled lists (referred to as gene expression-based classification, **Methods**), and observed strong concordance between image-based and gene expression-based classification, supporting the validity of image-based SPG spot annotations (**Figure S14**, **Methods**). We next compared the proportion of image-based SPG spots with marker gene expression across spots for each SpD and found that domains with higher SPG proportions generally exhibited higher expression of the marker genes of the same type, indicating concordance between morphological classification and transcriptomic profiles (**Figure S15**). We observed an increased proportion of PNN spots and elevated PNN marker gene expression in SpD02-L3/4 and SpD05-L5, supporting the biological validity of our image-based classifications (**Figure S14C**, **Figure S15C**).

### 2.2 Spatially-localized SCZ-associated differences highlight synaptic and non-neuronal transcriptional alterations

Both bulk RNA-seq and single-nucleus (snRNA-seq) studies have identified changes in gene expression associated with SCZ in the prefrontal cortex, including the dlPFC.^15–18,41^ While laminar enrichment of differentially expressed genes (DEGs) in these studies can be inferred, SRT can more directly spatially localize gene expression changes. To identify changes in gene expression associated with SCZ, we applied two different DE models to quantify the overall and within-SpD transcriptional alteration, respectively. We refer to the first model as the “layer-adjusted” DE model (**Figure S16A**). This model quantified global transcriptional alterations across tissue sections while adjusting for laminar differences using the data-driven SpDs. This analysis identified 172 DEGs (referred to as ‘layer-adjusted SCZ-DEGs’) with a false-discovery rate (FDR) < 0.10 (**Figure 2A**, **Supplementary Table 4**). The majority of these layer-adjusted SCZ-DEGs were down-regulated (58%, n=99 genes), consistent with previous bulk and snRNA-seq findings, including Ruzicka et al..^16,41^ For example, a large-scale bulk RNA-seq study (BrainSEQ Phase 2; dlPFC samples only) identified 632 DEGs at FDR < 0.10, a majority of which were also down-regulated (60%, n=379 genes).^41^ We compared our layer-adjusted SCZ-DEGs to BrainSEQ Phase 2 bulk RNA-seq DEGs and found 32 overlapping genes, including *MAPK3*, *C3*, *SERPINA3*, and *A2M* (**Figure 2B**). *MAPK3* was among the 73 up-regulated layer-adjusted SCZ-DEGs, and its dysregulation is consistent with both genetic and transcriptomic evidence linking MAPK3 signaling with SCZ.^13,42,43^

**Figure 2.**
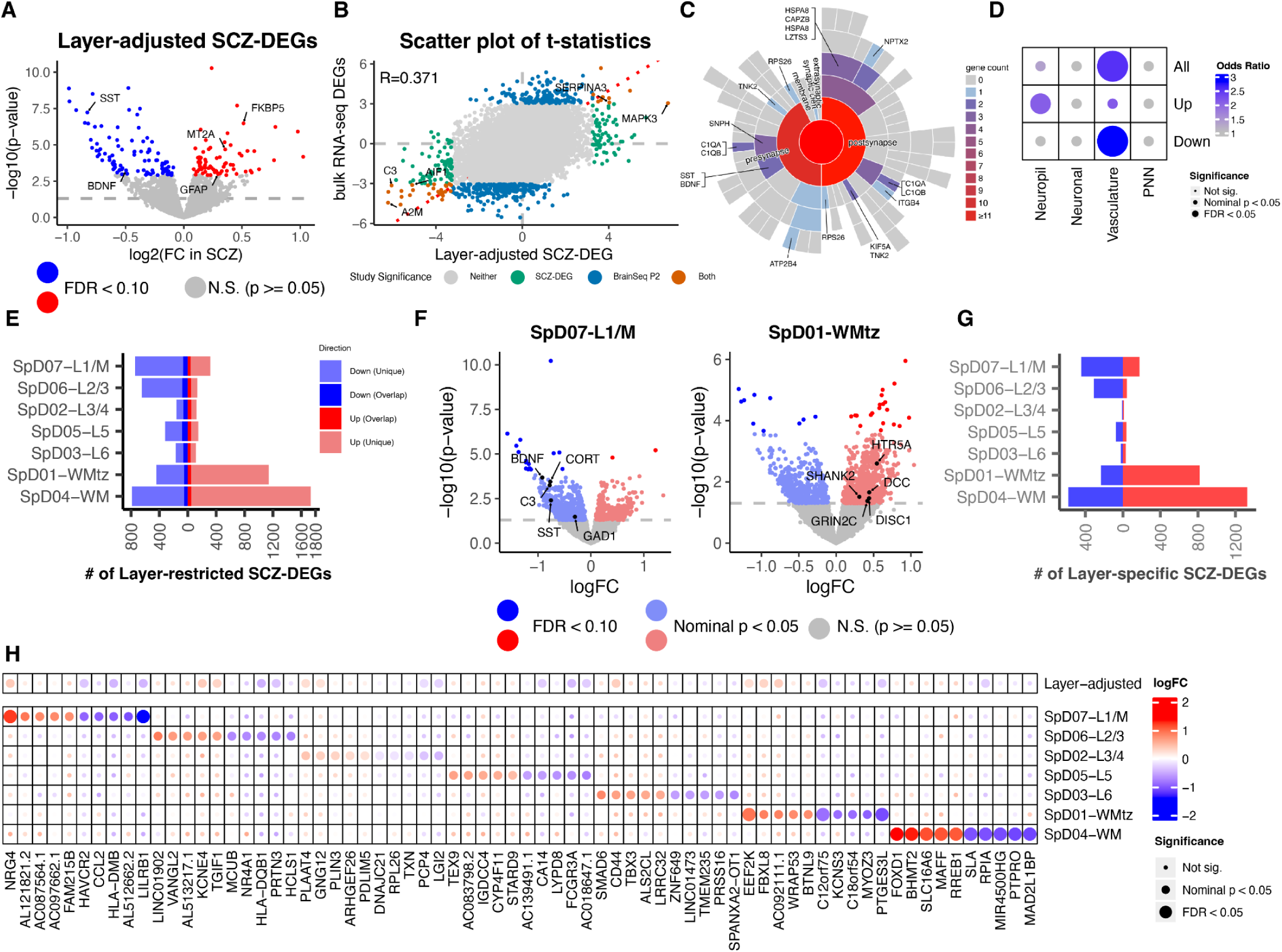
Identification and spatial characterization of SCZ-associated gene expression changes across cortical layers in the human dlPFC. (A) Volcano plot showing the log_2_ fold-change (logFC) of 172 layer-adjusted SCZ-associated DEGs (SCZ-DEGs) with FDR < 0.10. Red and blue denote up- and down-regulated SCZ-DEGs, respectively. (B) Scatter plot comparing *t*-statistics between BrainSEQ Phase 2 bulk RNA-seq DEGs previously identified in the dlPFC^41^ and layer-adjusted SCZ-DEGs identified here. (C) Layer-adjusted SCZ-DEGs mapped onto synaptic compartments using the SynGO database.^56^ Representative genes are annotated with their associated synaptic components. (D) Dot plot showing enrichment of layer-adjusted SCZ-DEGs across the four SPG-defined microenvironments involving neuropil, neuronal, vasculature, and PNN microenvironments. Enrichment significance was stratified by FDR < 0.05 and nominal *p* < 0.05. Odds ratios were indicated with the color scale. (E) Diverging bar plot showing the numbers of up- and down-regulated layer-restricted SCZ-DEGs identified at nominal *p* < 0.05 and highlighting their overlap with the layer-adjusted SCZ-DEGs identified at FDR < 0.10. Red and blue denote up- and down-regulated SCZ-DEGs, respectively. (F) Volcano plots showing layer-restricted SCZ-DEGs for SpD07-L1/M and SpD01-WMtz, respectively. Blue denotes down-regulation and red denotes up-regulation in SCZ. Points in darker shades indicate genes at FDR < 0.10, whereas lighter shades indicate genes at nominal *p* < 0.05. Red and blue denote up- and down-regulated SCZ-DEGs, respectively. (G) Diverging bar plot showing the numbers of up- and down-regulated layer-specific SCZ-DEGs at nominal *p* < 0.05. Red and blue denote up- and down-regulated SCZ-DEGs, respectively. (H) Dot plot showing logFC and corresponding statistical significance of top 5 most differentially expressed up-and down-regulated layer-specific SCZ-DEGs (ranked by logFC) within each SpD, compared with their logFCs estimated in the layer-adjusted DE model (Top row). Statistical significance was stratified by FDR < 0.05 and nominal *p* < 0.05. LogFCs were indicated with the color scale.

Layer-adjusted SCZ-DEGs were consistent with known functional alterations in SCZ. For example, neuroplasticity and glutamatergic signaling are consistently implicated in SCZ pathology,^44,45^ and our SCZ-DEGs included many genes with key roles in neuroplasticity, including *BDNF*, *FKBP5*, *MAPK3*, *NPTX2*, and *EGR1*. Consistent with previous studies in postmortem human brain,^17,46^ we found down-regulation of microglia-associated genes, including *TMEM119*, *P2RY12*, *GPR34*, *AIF1*, *TREM2*, and *C3*. GABAergic signaling dysfunction in SCZ is thought to contribute to disruptions of gamma band oscillatory synchrony and excitatory-inhibitory balance in SCZ, physiological functions that are controlled by GABA neurons.^47,48^ Consistent with this functional link to SCZ, layer-adjusted SCZ-DEGs included genes associated with GABA neuron functions, such as *CORT* and *SST*. We also identified layer-adjusted SCZ-DEGs associated with astrocyte biology, including *GFAP*, *AQP1*, and *SLC14A1*, which were up-regulated, along with *ALDH1A1*, which was down-regulated. These data support a recent study suggesting that disruptions in synaptic neuron-astrocyte programs contribute to SCZ.^15^ Finally, we identified multiple metallothionein family members (*MT1X*, *MT1M*, *MT2A*) as layer-adjusted SCZ-DEGs, which participate in heavy metal detoxification and may influence the ability of PNNs to buffer metal ions via metalloproteinase-based ECM remodeling.^49,50^ These findings may be relevant for emerging hypotheses linking developmental neurotoxicity to oxidative stress and redox dysregulation in SCZ.^51,52^ Gene set enrichment analysis (GSEA) of the layer-adjusted SCZ-DEGs further reinforced these findings, which highlighted signaling pathways involved in cellular response to metal ions, along with cellular processes, including cytokine-related signaling and myeloid cell processes. Intriguingly, the ATP metabolism and mitochondrial respiratory terms were distinctively enriched, suggesting significant metabolic alterations across cortical layers in SCZ (**Figure S17**).

Given established links between synaptic dysfunction and SCZ,^15,53–55^ we performed enrichment analysis of layer-adjusted SCZ-DEGs against SynGO annotations.^56^ SynGO identifies overrepresented synaptic terms in an experimental dataset, predicts localization of genes to pre- and post-synaptic compartments and assigns known synaptic functions (**Figure 2C**). We identified a subset of layer-adjusted SCZ-DEGs as synapse-related, including *NPTX2*, *BDNF*, *SST*, *C1QA*, and *C1QB*, spanning both the pre- and post-synaptic compartments. Corroborating the synaptic function of these genes, up-regulated layer-adjusted SCZ-DEGs were significantly enriched in SPG-defined neuropil spots (Odds ratio=2.07, nominal *p*=0.019; **Figure 2D**). Finally, several complement-related genes (e.g., *C1QA*, *C1QB*) were localized to the synapse, supporting the theory that neuroimmune regulation of synapse pruning may be disrupted in SCZ.^57,58^

To directly test whether SCZ-associated changes in gene expression were localized to specific layers, we developed a second DE model, referred to as the “layer-restricted” DE model. Compared to the previous layer-adjusted model that quantifies the overall transcriptional alteration, the “layer-restricted” model quantifies the alteration within individual SpDs using statistical interaction of SpDs and diagnosis (**Figure S16B**). This analysis identified 944 and 6,978 DEGs across all SpDs at FDR < 0.10 and nominal *p* < 0.05 respectively (**Supplementary Table 5**). To improve the power to detect small but important transcriptional signals, we decided to focus on the 6,978 nominal DEGs, and, for simplicity, refer to them as the layer-restricted SCZ-DEGs unless clarified otherwise. Layer-restricted SCZ-DEGs included 170 out of the 172 layer-adjusted SCZ-DEGs (**Figure 2E**). Consistent with snRNA-seq studies showing an up-regulation of DEGs in glial cells in SCZ,^16^ more than half of the layer-restricted SCZ-DEGs were localized to WM-associated domains (26%, n=1,581 genes for SpD01-WMtz; 36%, n=2,521 genes for SpD04-WM) and the majority of these WM-localized SCZ-DEGs show up-regulation (2,873 up-regulated vs. 1,229 down-regulated DEGs).

Up-regulated layer-restricted SCZ-DEGs in SpD04-WM reflected biological pathways related to translation/metabolic activity, cellular stress, and vascular-associated signaling. Down-regulated layer-restricted SCZ-DEGs in WM encompassed components of immune-related and cholinergic signaling (particularly, *CHRM1*, *CHRM3*, *PLCB1*, *ITPR1*, and *PRKCB*) (**Figure 2E**). These changes may indicate disrupted cholinergic activity within projection- and myelination-rich WM regions.^59–61^ In parallel, SpD01-WMtz, a region enriched in both neuronal and glial cell populations (**Figure 1G**, **Figure S9**), showed down-regulation of microglial and immune-associated genes and up-regulation of vascular stress response genes. Notably, genes involved in synaptic signaling and neurotransmission were up-regulated, including SCZ-implicated neurodevelopmental genes (*DISC1*, *DCC*), glutamatergic synaptic genes (*GRIN2C*/*D*, *SHANK1*/*2*), and neurotransmitter receptor genes such as *HTR5A* (**Figure 2F**). The detection of neuronal expression within the WM-associated regions further supports a contribution of neuronal processes to function in these domains. These findings highlight the utility of spatial transcriptomics for identifying both neuronal signatures and non-neuronal signatures within WM regions enriched for glial populations. Beyond the WM-associated domains, the majority of layer-restricted SCZ-DEGs that localized to the superficial layers, particularly SpD07-L1/M and SpD06-L2/3, were down-regulated. Microglia-associated genes (*AIF1*, *TREM2*, *C3*), GABA neuron signaling genes (*GAD1*, *SST*, *CORT*, *CRH*, *PVALB*), and genes associated with neuroplasticity (*BDNF*, *VGF*, *PTN*, *NLGN3*, *EPHB1*) were down-regulated in SpD07-L1/M, for example. Collectively, we observed layer-restricted variations in SCZ-DEGs and their expression patterns, but common biological themes emerged, including neurotransmission imbalance, microglia-associated down-regulation, vascular/stress-associated disruptions, and neurodevelopment-related alterations across cortical layers and WM regions.

Last, we identified layer-restricted SCZ-DEGs that were differentially expressed in only a single SpD. We refer to these genes as ‘layer-specific SCZ-DEGs’ (**Figure S16C**). Among the nominal layer-restricted SCZ-DEGs, 4,073 genes were layer-specific (2,416 up-regulated and 1,657 down-regulated). The number of layer-specific SCZ-DEGs was largely proportional to layer-restricted SCZ-DEGs, with the exception of SpD02-L3/4, which showed fewer layer-specific SCZ-DEGs (**Figure 2G**). Outside of the WM domains, the largest number of layer-specific SCZ-DEGs were found in SpD07-L1/M. For example, we observed up-regulation of *NRG4* in SpD07-L1/M. Neuregulin-ERBB signaling plays key roles in neural development and synaptic plasticity. While these functions are well established for several neuregulin family members, the role of *NRG4* is less studied in the brain. However, recent work suggests a role in neuronal development and morphology,^62–64^ the possibility that *NRG4* may also contribute to the regulation of neuronal structure within neuropil-rich molecular layers. Analysis of the layer-specific SCZ-DEGs with largest transcriptional alteration in each SpD (**Figure 2H**) suggests involvement of multiple signaling pathways associated with synaptic function and neuronal activity (*KCNE4*, *PDLIM5*, *EEF2K*) as well as cellular growth and differentiation (*NR4A1*, *SMAD6*, *TBX3*). We also identified genes associated with canonical immune regulation, with the majority being down-regulated (*LILRB1*, *HAVCR2*, *CCL2*, *HLA-DMB*, *HLA-DQB1*). Down-regulated genes were restricted to SpD07-L1/M and SpD06-L2/3 while *CD44*, which was up-regulated, mapped to SpD03-L6. We note that immune signaling in brain cells is relatively less understood than in peripheral tissues, and the functional significance of these gene expression changes for immune signaling or other signaling pathways they regulate is not yet clear.

### 2.3 Integrating cellular context with spatially-resolved gene expression changes reveals non-neuronal cell-type contributions in SCZ

While our laminar DE analyses identified spatially-localized SCZ-DEGs that have been attributed to synaptic function and canonical immune signaling (**Figure 2C**–**Figure 2H**), these analyses lacked cellular resolution. To provide additional biological context interpreting the spatially-localized SCZ-DEGs, we integrated our findings with an existing snRNA dlPFC atlas^25^ and a SCZ-comparative study,^16^ using gene set enrichment approaches.

We first performed cell-type enrichment analysis using layer-adjusted SCZ-DEGs and dlPFC cell-type marker genes^25^ (**Figure 3A**). Up-regulated SCZ-DEGs were enriched for marker genes of non-neuronal populations, particularly astrocytes and vascular leptomeningeal cells (VLMCs). Conversely, down-regulated SCZ-DEGs were enriched for markers of microglia and immune cells. Notably, both up- and down-regulated SCZ-DEGs were enriched in endothelial cells, pericytes (PC), and smooth muscle cells (SMC), consistent with emerging evidence of vascular signaling dysregulation in SCZ.^22,23^ Unexpectedly, we did not observe strong evidence supporting enrichment of neuronal cell-type markers among layer-adjusted SCZ-DEGs. To better understand this observation, we compared layer-adjusted SCZ-DEGs against SpD marker genes and checked their laminar localization (**Figure 3B**). SCZ-DEGs predominantly mapped to cortical domains enriched for non-neuronal cells, including SpD07-L1/M, SpD01-WMtz, and SpD04-WM, supporting an association with non-neuronal cell types. SpD07-L1/M, a domain abundant in vascular and glial cells, showed strong enrichment for both up- and down-regulated SCZ-DEGs. Together, these analyses highlight the emerging importance of non-neuronal populations in SCZ.^15,65,66^

**Figure 3.**
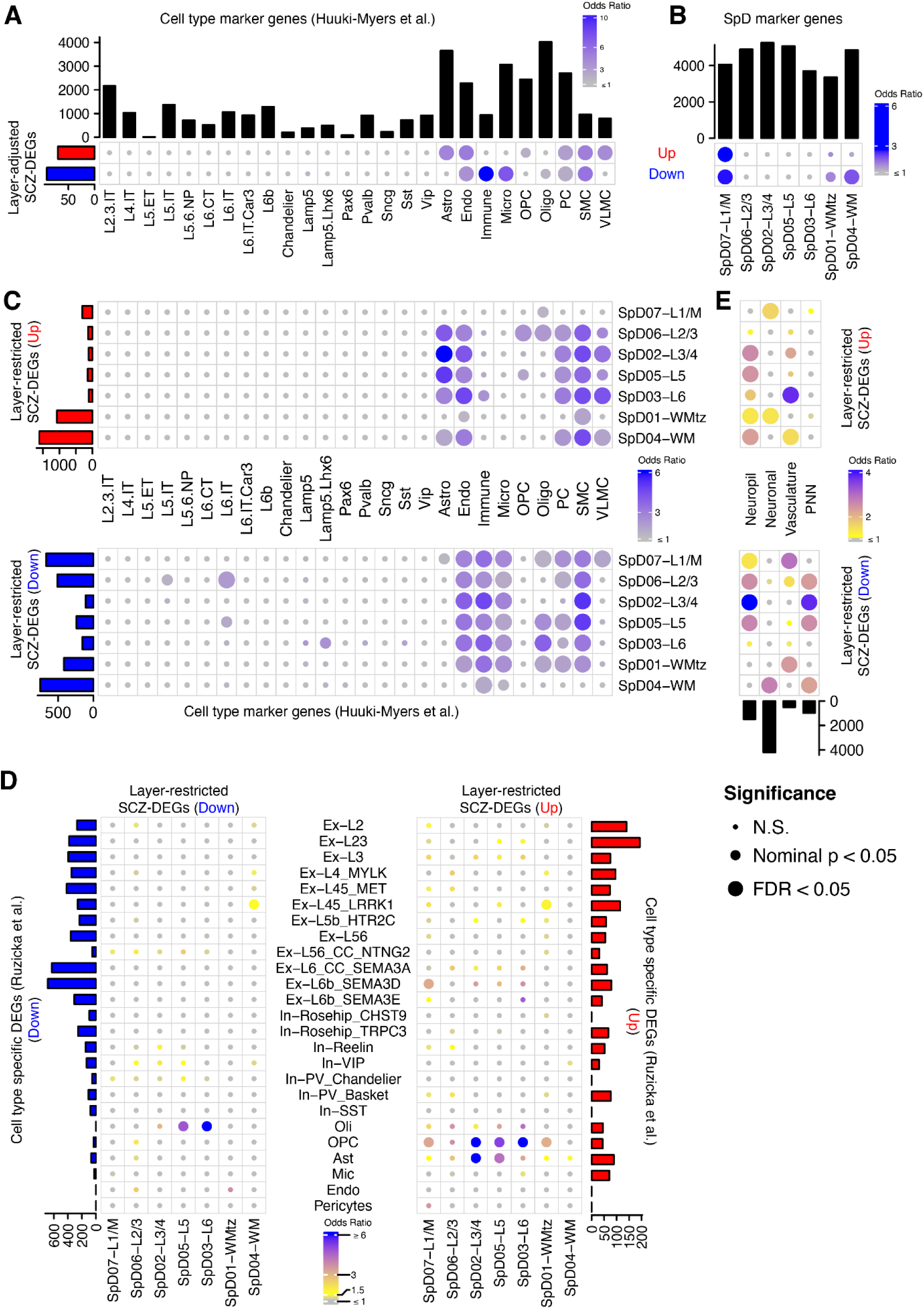
Integration of cellular context into spatially-localized SCZ-associated gene expression changes. (A) Enrichment of 172 layer-adjusted SCZ-DEGs among dlPFC cell-type marker genes from Huuki-Myers et al..^25^ Red and blue denote up- and down-regulated SCZ-DEGs, respectively. Dot size indicates the three different significance levels for gene set enrichment: non-significant (N.S.), nominal *p* < 0.05, and FDR < 0.05. Dot color indicates the odds ratio. The same color and significance schemes are applied across panels in **A**-**E**. (B) Enrichment of 172 layer-adjusted SCZ-DEGs among marker genes associated with data-driven SpDs. The color scheme for directionality of gene expression is the same as in (**A**). (C) Enrichment of nominal layer-restricted SCZ-DEGs among dlPFC cell-type marker genes from Huuki-Myers et al..^25^ (D) Enrichment analysis comparing reference cell-type-specific DEGs from Ruzicka et al.^16^ with nominal layer-restricted SCZ-DEGs. (E) Enrichment of layer-restricted SCZ-DEGs in SPG-defined microenvironment marker genes for neuropil, neuronal, vasculature, and PNN microenvironments. SpD labels and cortical layer assignments are consistent with those in (**C**).

We also performed cell-type enrichment analysis with nominal layer-restricted SCZ-DEGs (**Figure 3C**). SCZ-DEGs, particularly those that were up-regulated, were enriched for astrocyte markers in neuronal-rich domains (SpD06-L2/3, SpD02-L3/4, SpD05-L5, SpD03-L6) as well as WM (SpD04-WM), suggesting widespread astrocyte-associated transcriptional alterations.^15,65,67^ In contrast, down-regulated SCZ-DEGs were enriched predominantly for microglia and immune cell markers across all SpDs, consistent with our cell-type enrichment of layer-adjusted SCZ-DEGs (**Figure 3A**) and prior studies.^17,46^ Enrichment of oligodendrocyte markers was divergent, with down-regulated SCZ-DEGs enriched for oligodendrocyte markers in deeper layers (SpD05-L5, SpD01-WMtz), and up-regulated SCZ-DEGs enriched in superficial layers (SpD06-L2/3). While we identified relatively few instances of enrichment among neuronal marker genes within SCZ-DEGs across SpDs, there were several exceptions. For example, down-regulated SCZ-DEGs in SpD06-L2/3 were enriched for L6 intratelencephalic-projecting (IT) neuron markers. Together, the cell-type enrichment analysis of layer-restricted SCZ-DEGs delineated the spatial localization of implicated cell types and revealed contributions from both glial and neuronal populations to SCZ-associated gene expression changes across cortical layers and WM.

To determine the extent to which our spatially-localized SCZ-DEGs recapitulate cell-type-specific transcriptional alterations identified in previously published SCZ snRNA-seq studies, we compared our layer-restricted SCZ-DEGs with cell-type-specific SCZ-DEGs identified from Ruzicka et al.^16^ (referred to as “reference DEGs”) (**Figure 3D**). Reference DEGs were stratified by directionality (**Methods**), and enrichment of up- and down-regulated reference DEGs was examined within our corresponding up- and down-regulated layer-restricted SCZ-DEGs. The highest concordance was observed among glial cells, including astrocytes, oligodendrocytes, and oligodendrocyte progenitor cells (OPCs). Specifically, up-regulated astrocyte-specific reference DEGs were enriched among up-regulated SCZ-DEGs, with the strongest concordance in SpD02-L3/4. This finding provides evidence for astrocyte-associated transcriptional alterations in SCZ, where these changes may reflect circuit-level disruptions in specific cortical layers.^15,65,67^ Oligodendrocyte-specific reference DEGs exhibited strong concordance among down-regulated SCZ-DEGs in deeper cortical layers including SpD05-L5 and SpD03-L6, consistent with oligodendrocyte marker genes being enriched among layer-restricted SCZ-DEGs in these domains (**Figure 3C**). While our earlier enrichment analyses demonstrated limited enrichment of neuronal marker genes among SCZ-DEGs (**Figure 3A**, **Figure 3C**), registration of reference DEGs revealed transcriptional alterations across several excitatory and inhibitory neuronal subtypes, such as deep-layer excitatory neuron-specific DEGs (Ex-L6b_SEMA3D) overlapping in SpD07-L1/M. However, the magnitude of neuronal changes was generally less pronounced than those observed in glial populations. Indeed, our dataset was more sensitive to glial- and WM-associated alterations than the snRNA-seq dataset,^16^ highlighting the complementary strength of SRT in capturing non-neuronal transcriptional changes *in situ* (**Figure S18**). Together, these findings indicate that spatially-localized gene expression changes encompass a spectrum of SCZ-associated alterations, with glial-linked changes being more salient in our SRT dataset.

To further interrogate these glial effects and infer how our layer-restricted SCZ-DEGs related to transcriptional alterations within spatially-localized cellular compartments or synaptic microenvironments, we assessed enrichment of marker genes for the four SPG-defined microenvironments (neuropil, PNN, neuronal, vasculature) among our SCZ-DEGs (**Figure 3E**). Broadly, these microenvironment marker genes mapped onto layer-restricted SCZ-DEGs across multiple SpDs, revealing heterogenous laminar patterns. Neuropil marker genes were enriched among both up- and down-regulated SCZ-DEGs, with up-regulated SCZ-DEGs localized to mid-to-deep neuropil (SpD02-L3/4 to SpD01-WMtz) and down-regulated SCZ-DEGs to superficial-to-middle layer neuropil (SpD07-L1/M to SpD05-L5), suggesting divergent synaptic dysregulation across neuronal and glial processes. PNN marker genes were enriched among down-regulated SCZ-DEGs in middle cortical layers, particularly SpD06-L3/4, where we also observed astrocytic alterations (**Figure 3C**, **Figure 3D**). These findings may reflect SCZ-associated disruption of astrocyte-PNN interactions, which are crucial for maintaining synaptic homeostasis of PVALB neurons.^27,30,68–70^

Non-intuitively, marker genes for neuronal spots were enriched among SCZ-DEGs in glial-rich domains (SpD07-L1/M, SpD01-WMtz, SpD04-WM), possibly implicating aberrant neuron-glia interactions that may be linked to extranuclear transcripts within neuronal processes or sparse neuronal populations interspersed throughout these domains.^71–75^ Vascular marker genes were enriched among up-regulated SCZ-DEGs in SpD03-L6 and SpD04-WM and among down-regulated SCZ-DEGs in SpD07-L1/M and SpD01-WMtz, which may signify domain-specific neurovascular alterations. Overall, layer-restricted SCZ-DEGs were largely associated with neuropil and PNN microenvironments, a finding supported by enrichment of bulk RNA-seq DEGs^41^ in these domains (**Figure S19**). In summary, these data highlight the utility of SRT to capture subtle alterations in specialized cellular and extracellular compartments relevant to synaptic function and point towards potential dysregulation of neuron to non-neuronal interactions across microenvironment domains in SCZ.

### 2.4 Spatial divergence in SCZ-associated transcriptional regulation and genetic risk across neuronal and non-neuronal cortical domains

Transcription factors (TFs) are highly represented amongst genes linked to SCZ risk.^13^ To better understand biological mechanisms mediating regulation of SCZ-associated gene expression and determine whether those mechanisms are linked to SCZ genetic risk, we predicted candidate TFs that may trans-regulate the layer-restricted SCZ-DEGs utilizing the prediction algorithm ChEA3^76^ (**Figure 4A**, **Supplementary Table 6**, **Methods**). We tested enrichment of TF target gene sets within the up- and down-regulated layer-restricted SCZ-DEGs for each SpD, respectively. Candidate TFs were ranked based on their aggregated enrichment scores, where the top 10 TFs per SpD were displayed across SpDs. Many TFs recurred across multiple SpDs (e.g., NACC2, FOSB among up-regulated SCZ-DEGs; OLIG1, SOX10 among down-regulated SCZ-DEGs), suggesting cross-laminar involvement in transcriptional dysregulation. We also observed some TFs concurrently appearing within both up- and down-regulated SCZ-DEGs. These discordant patterns likely indicate a context-dependent regulation as influenced by local cell composition and microenvironment features. In addition, we identified TFs that are involved in cellular differentiation and maturation, including neuronal lineages (ATOH8, TSHZ1), synaptic plasticity and neurite remodeling (NR4A1, KLF9), and neurovascular development (SOX18). We also identified TFs exhibiting localized associations with specific SpDs. For instance, ZNF324B showed strong relevance to up-regulated SCZ-DEGs in SpD01-WMtz, while CAMTA1, PEG3, ZNF483, PURG, and CSRNP3 demonstrated relative specificity in SpD07-L1/M. Additionally, within SpD04-WM, TFs including SGSM2, FLYWCH1, TSHZ1, and TEF showed preferential associations with down-regulated SCZ-DEGs, suggesting layer-restricted regulatory involvement in downstream targets within this domain.

**Figure 4.**
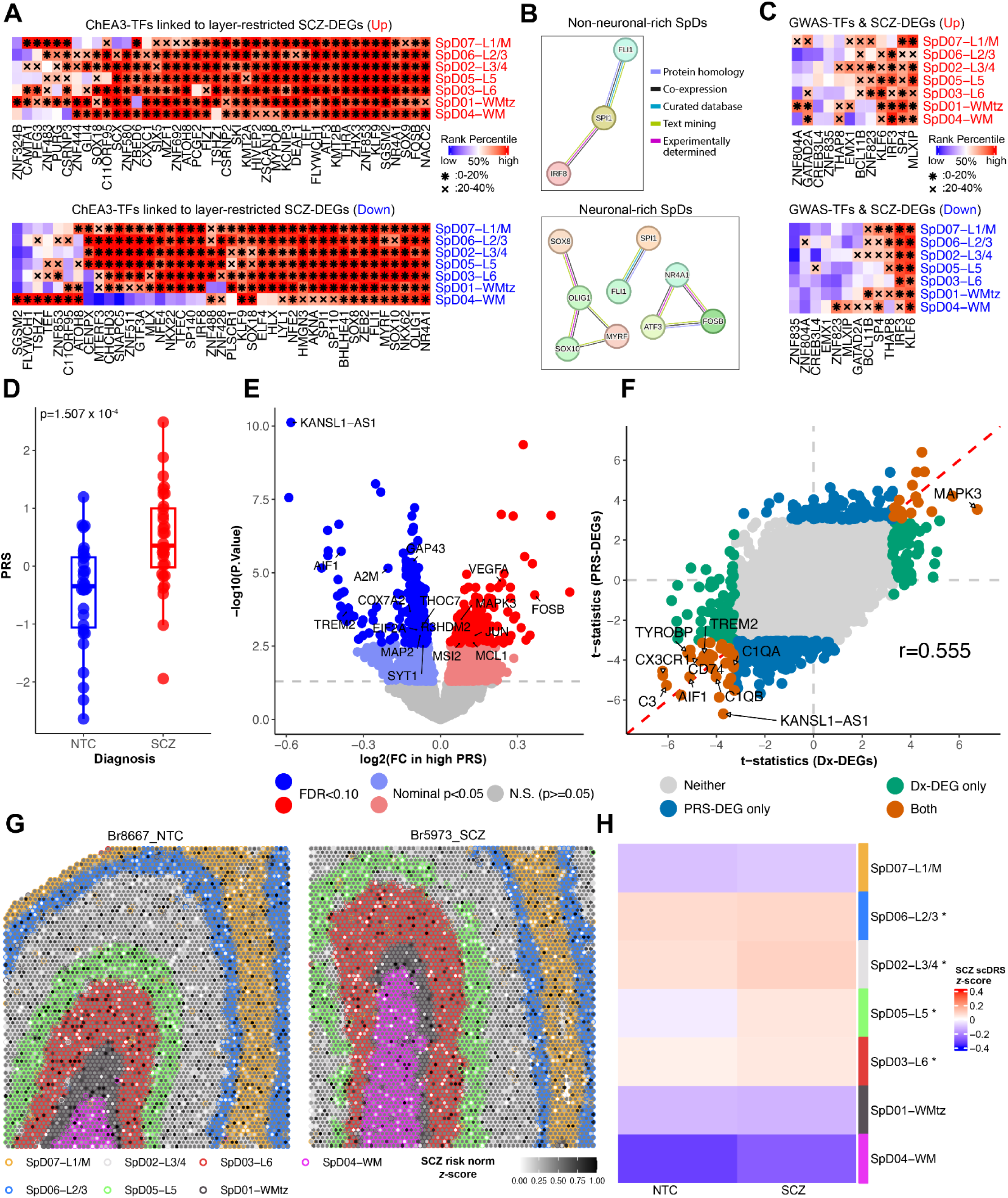
Convergence of transcriptional regulation and genetic risk on spatio-molecular differences in SCZ. (A) Heatmaps showing the top ranked 10 TFs predicted by ChEA3,^76^ per SpD, associated with layer-restricted SCZ-DEGs up-regulated (top) and down-regulated (bottom) in SCZ. Due to overlapping TFs across SpDs, the initial total of 70 TFs in each group was reduced to 42 unique TFs (up-regulated) and 41 unique TFs (down-regulated). (B) STRING TF functional networks for non-neuronal-rich (SpD07-L1/M, SpD01-WMtz, SpD04-WM) and neuronal-rich (SpD06-L2/3, SpD02-L3/4, SpD05-L5, SpD03-L6) domains. The top 10 TFs linked to up- and down-regulated layer-restricted SCZ-DEGs per SpD were selected, resulting in initial input sets of 60 TFs (non-neuronal) and 80 TFs (neuronal). After removing overlapping TFs across SpDs and unmapped entries (e.g., NKX62, NKX22), 48 TFs (non-neuronal) and 42 TFs (neuronal) remained. Networks display only connected nodes with an interaction score > 0.7. Complementary network diagrams including all TFs are shown in Figure S20 at an interaction score > 0.4 to capture broader connectivity. (C) Heatmap of ChEA3-predicted TFs associated with layer-restricted SCZ-DEGs, intersecting with prioritized SCZ GWAS risk genes from Trubetskoy et al.^13^ across PRECAST SpDs. (D) Box plot comparing predicted SCZ PRSs shows a higher mean PRS in the SCZ group (n=32 donors, red), compared to the NTC group (n=31 donors, blue); *p*=1.507 x 10^-4^. (E) Volcano plot of layer-adjusted PRS-associated DEGs (PRS-DEGs). The x-axis is the logFC and the y-axis is the -log_10_ (transformed *p*-value). Red and blue denote up- and down-regulated PRS-DEGs in high-PRS donors, respectively. Points in darker shades indicate PRS-DEGs at FDR < 0.10. Points in lighter shades indicate PRS-DEGs at nominal *p* < 0.05. (F) Scatter plot comparing *t*-statistics between PRS-DEGs and layer-adjusted SCZ-DEGs (Dx-DEGs; Dx=diagnosis) (G) Spot plots illustrating the distribution of SCZ disease relevance normalized *z*-scores in representative NTC and SCZ donors, as estimated using scDRS. Each point represents a Visium spot, shaded according to SCZ risk *z*-score (white to black). SpD boundaries are outlined and color-coded to indicate PRECAST SpDs at *k*=7. (H) Heatmap showing mean SCZ scDRS *z*-scores across SpDs in NTC and SCZ groups. The color scale reflects relative enrichment *z*-scores. Asterisks indicate SpDs with significantly increased SCZ scDRS *z*-scores relative to the NTC group (FDR < 0.05).

Given the substantial number of candidate TFs and their heterogeneous patterns across SpDs (**Figure 4A**), we used STRING database,^77^ which integrates functional and physical protein associations, to construct interaction networks of TFs and characterize their overarching biological themes. Based on the domain-specific enrichment pattern of TFs in non-neuronal-rich cortical domains (**Figure 4A**) as well as the robust mapping of SCZ-DEGs to these regions (**Figure 3A**–**Figure 3C**), we stratified ChEA3-derived TFs (top 10 ranked per SpD for each up- and down-regulated layer-restricted SCZ-DEGs, **Methods**) into two groups: non-neuronal-rich domains (SpD07-L1/M, SpD01-WMtz, SpD04-WM) and neuronal-rich domains (SpD06-L2/3, SpD02-L3/4, SpD05-L5, SpD03-L6) (**Figure 4B**, **Figure S20**). Within non-neuronal-rich domains, TF networks contained factors associated with myeloid and microglial regulatory signaling (SPI1, IRF8, FLI1), with their associated SCZ-DEGs showing coordinated down-regulation (**Supplementary Table 6**). In neuronal-rich domains, we observed a similar network associated with myeloid and stress response signaling (SPI1, FLI1, TFEC). We also identified networks implicated in oligodendrocyte differentiation and myelination (MYRF, OLIG1, SOX10), whose associated SCZ-DEGs were down-regulated, and activity-dependent stress-response programs (NR4A1, FOSB, ATF3), whose SCZ-DEGs were up-regulated (**Supplementary Table 6**). Together, these TF-associated networks suggest that layer-restricted SCZ-DEGs converge on domain-specific regulatory contexts, with coordinated down-regulation of glial and myelination-related gene programs and enrichment of activity-dependent and stress-response transcriptional programs in neuronal-rich domains.

To integrate transcriptional regulation with genetic risk, we examined whether candidate TFs intersected with known genetic factors implicated in SCZ. By cross-referencing candidate TFs from ChEA3^76^ database with prioritized SCZ-associated GWAS risk genes,^13^ we identified and visualized 12 overlapping TFs with their enrichment ranks (**Figure 4C**, **Methods**). Among these, two inflammation-related TFs, KLF6 and IRF3, were consistently ranked among the top 50% of TFs across all SpDs, frequently within the top 20-35%, across both up- and down-regulated SCZ-DEGs. Domain-specific GWAS-prioritized TFs were also noted, although their rankings were modest, including ZNF823 (linked to down-regulated SCZ-DEGs in SpD04-WM, top 17%) and CREB3L4 (linked to down-regulated SCZ-DEGs in SpD05-L5, top 36%). ZNF804A, a SCZ risk gene,^78–80^ showed target gene modules overlapping with down-regulated SCZ-DEGs in neuronal-rich SpD06-L2/3 (top 38%) and with up-regulated SCZ-DEGs within non-neuronal-rich domains (SpD07-L1/M, SpD01-WMtz; top 38% and 13%, respectively). Many SCZ-DEGs linked to ZNF804A were involved in neuronal maturation (*NR4A2*, *EGR1*, *DCC*), neurotransmission (*GRIN1*, *CACNA2D3*, *GABRB3*, *SCN2B*, *SCN8A*), and synaptic plasticity and organization (*BDNF*, *CNTN1*, *CSMD3*), supporting a convergent genetic-transcriptional disruption of neuronal circuitry and plasticity in these cortical regions (**Supplementary Table 6**, **Figure S21**).

To quantitatively assess the impact of genetic risk on gene expression, we computed polygenic risk scores (PRSs) for SCZ (**Methods**).^81^ As expected, PRS was significantly higher in the SCZ group compared to controls (*p*=1.507 x 10^-4^) (**Figure 4D**). Using the layer-adjusted model, we replaced diagnosis as the predictor with PRS. This analysis identified 354 PRS-associated DEGs (PRS-DEGs) controlling FDR < 0.10 (1,588 PRS-DEGs at nominal *p* < 0.05, **Figure 4E**, **Supplementary Table 7**, **Methods**). PRS-DEGs included genes involved in protein translation (e.g., *EIF2A*), mitochondrial respiration (*COX7A2*), apoptosis (*MCL1*), microglial function (*AIF1*, *TREM2*), vascular and immune processes (*VEGFA*, *A2M*), activity-related signaling (*FOSB*, *JUN*, *SYT1*), and neuronal structure and outgrowth (*MAP2*, *GAP43*). Several PRS-DEGs mapped to SCZ risk loci, including *MAPK3*, *MSI2*, *R3HDM2*, *THOC7* and *KANSL1-AS1*.^13^ PRS-DEGs were moderately correlated with diagnosis-associated DEGs (Pearson’s correlation=0.555, **Figure 4F**) with 55 genes shared between the two sets. Many of these shared genes were consistently down-regulated in both PRS- and diagnosis-associated analyses and were linked to microglial- and immune-related functions such as *AIF1*, *TREM2, C1QA*, *C1QB*, *C3*, *CX3CR1*, *TYROBP*, and C*D74*. GSEA on these PRS-DEGs highlighted pathways related to metabolism, gliogenesis, immune regulation, and vascular-associated processes (**Figure S22**), suggesting non-neuronal contributions arising from both PRS- and diagnosis-associated transcriptional alterations. Together, these results provide evidence that PRS-DEGs link genetic risk for SCZ to transcriptional dysregulation that extends beyond neuronal processes, potentially.^66^

To further investigate how genetic risk is spatially distributed across cortical architecture, we applied the single-cell disease relevance score (scDRS) framework^82^ to predict SCZ genetic risk at the level of individual Visium spots (**Figure 4G**, **Figure S23**, **Methods**). Comparing across SpDs, genetic risk was largely localized to neuronal-rich domains (L2-6) as compared to non-neuronal domains (**Figure 4H)**. Multi-marker analysis of genomic annotation (MAGMA)^83^ also showed enrichment of SCZ genetic risk in neuronal-rich domains (**Figure S24**). We noted a small, but significant increase of the magnitude of genetic risk in SCZ samples showing slightly higher mean scDRS *z*-scores (FDR-adjusted *p* < 0.05). Overall, these results highlight a dichotomy between neuronal localization of genetic susceptibility (**Figure 4H**) and non-neuronal localization of disease-associated changes in gene expression (**Figure 2E**).

### 2.5 SCZ-associated gene expression changes map to spatially-localized cell types

To directly measure SCZ-associated changes in spatial gene expression at the cellular level, we generated Xenium data in a subset of the donors (n=12 NTC, n=12 SCZ) (**Figure 5A**, **Supplementary Table 1**, **Methods**). Xenium provides cell-level resolution of intact tissue sections using a pre-defined set of target genes, and can accurately profile cell types that are often underrepresented in other approaches, particularly microglia.^84,85^ This is important for SCZ because transcriptional changes in these underrepresented cell types have been reported,^15,86,87^ including changes in cell-type abundance,^18,88,89^ as well as dysregulation at the level of gene expression.^15,16,90,91^ To investigate whether the transcriptional changes we observed reflect intrinsic dysregulation or differences in cell-type abundance, we designed a 300-gene panel containing cell-type-specific gene markers and the layer-adjusted SCZ-DEGs (**Figure 2A**, **Supplementary Table 8**). After QC, 1,262,965 high-quality cells were retained (NTC: 629,981; SCZ: 632,984) (**Figure S25**, **Figure S26**).

**Figure 5.**
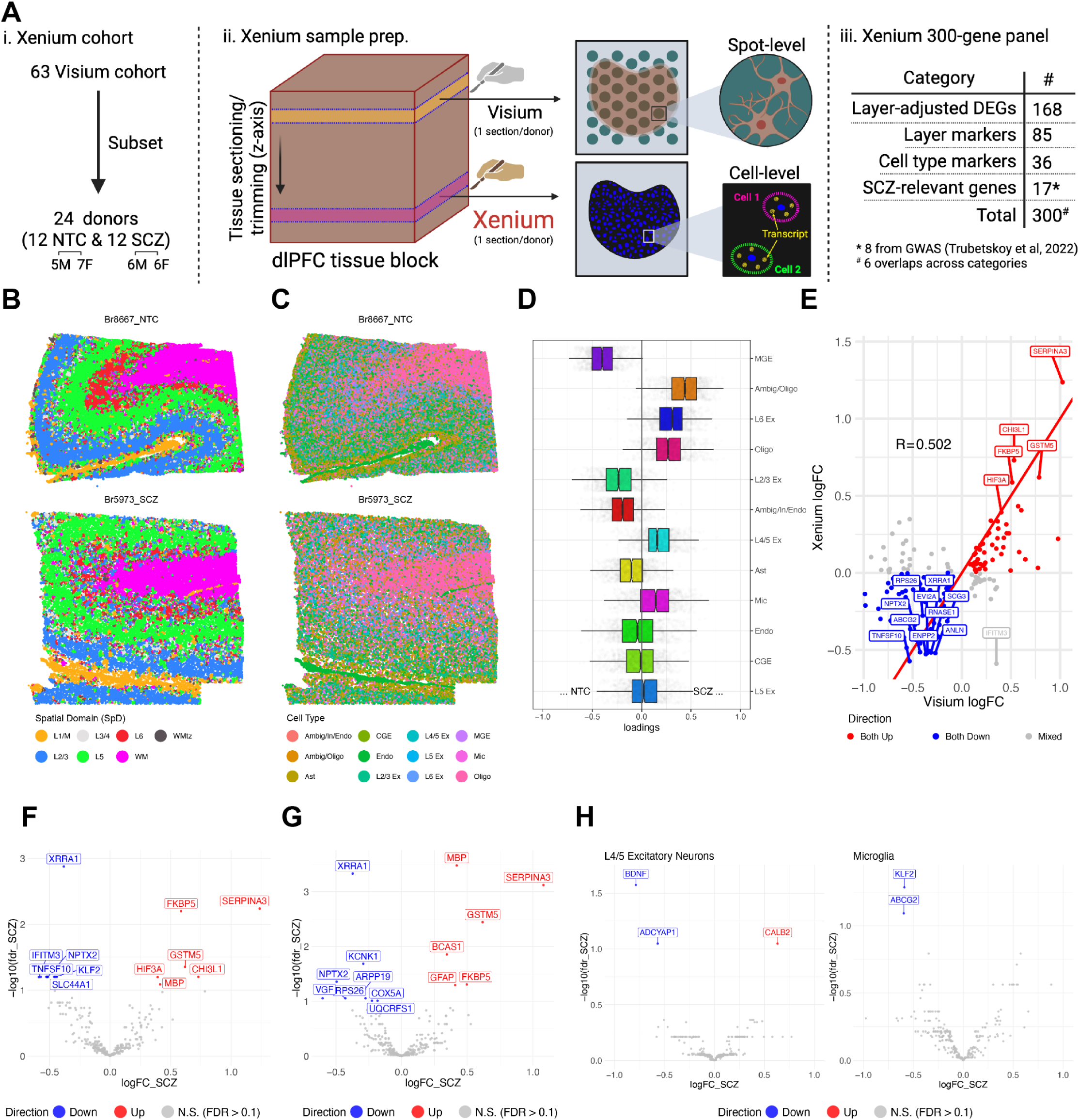
Cell-type-specific gene expression changes linked to SCZ in the human dlPFC. (A) Schematic showing the cohort selection and data generation process for the Xenium data. Notably, we highlight the single-cell resolution of the Xenium data, as opposed to the spot-level resolution of the Visium data. A table showing the 300 genes selected for the custom Xenium panel is shown, providing the number of genes in each category. Created in BioRender. Kwon, S. H. (2026) https://BioRender.com/9djvx8j (B) Cell-resolution plots of two representative tissue sections, Br8667 and Br5973 (NTC and SCZ, respectively), with cells colored by the predicted SpD label from spaTransfer after spatial smoothing. Predicted labels recapitulate the cortical layer structure of the dlPFC. (C) Same as (**B**), but with each cell colored by cell types. Cell types were annotated by performing data-driven clustering with BANKSY and then annotating clusters based on expression of the 36 cell-type markers in the gene panel. (D) Box plot of cell-type composition changes between NTC and SCZ, performed across all cortical layers. No significant changes in composition were detected between the diagnostic groups in any cell type. (E) Scatter plot showing comparison of logFC between the Visium layer-adjusted DE analysis and the Xenium layer-adjusted DE analysis, for each of the 168 Visium layer-adjusted SCZ-DEGs. (F) Volcano plot showing genes that are DEGs between NTC and SCZ, while adjusted for cortical layers. 19 genes are identified as differentially expressed at FDR < 0.10. Red and blue denote up- and down-regulated genes in SCZ, respectively. (G) Same as (**F**), but with cell-type adjustment rather than layer adjustment. 15 genes are identified as differentially expressed at FDR < 0.10. Red and blue denote up- and down-regulated genes in SCZ, respectively. (H) Volcano plots showing cell-type-specific SCZ-DEGs. Red and blue denote up- and down-regulated genes in SCZ, respectively. At FDR < 0.10, *CALB2* was up-regulated while *BDNF* and *ADCYAP1* were down-regulated in L4/5 Ex neurons, and *KLF2* and *ABCG2* were down-regulated in microglia.

We used spaTransfer,^92^ a label transfer method, to perform data-driven annotation of SpDs in the Xenium data using the Visium dataset as the reference. Specifically, each cell was annotated with a predicted cortical layer label identified using the Visium SpDs (**Figure 5B**). We found high correlation between our predicted Xenium cell-level SpDs and previous manually annotated dlPFC layers^27^ (**Figure S28**). Similar to the spot-level SpDs, the predicted cell-level SpDs accounted for the majority of variation across samples in the first two principal components (PC1 and PC2) (**Figure S27**), which suggested that the transferred annotations were not biased by diagnosis.

To further characterize cell-type composition, we used unsupervised spatial clustering with BANKSY^93^ and identified *k*=18 clusters, which we then annotated based on expression of cell-type marker genes (**Figure S29**, **Supplementary Table 8**). We grouped the 18 clusters into 12 clusters with cell-type labels; four excitatory (L2/3, L4/5, L5, and L6 ‘Ex’ neurons), two inhibitory (one ‘In’ neuron cluster consisting of a mixture of *VIP*- and *LAMP5*-expressing cells, and the other consisting of a mixture of *SST*- and *PVALB*-expressing cells), one each for astrocytes, microglia, oligodendrocytes, and endothelial cells, and two ambiguous (one expressing primarily oligodendrocyte marker genes and the other expressing inhibitory neuron and endothelial cell marker genes) (**Figure 5C**). Due to the limited number of genes on the panel, BANKSY could not separate the excitatory and inhibitory neuron subtypes, and we annotated the inhibitory neuron clusters as medial and caudal ganglionic eminence clusters, respectively (MGE for mixed *SST*/*PVALB* populations, CGE for *VIP*/*LAMP5*).^94,95^

Previous studies suggested alterations in cellular composition across cortical layers, particularly highlighting vulnerability in specific inhibitory neuronal populations in SCZ.^18,96,97^ To investigate this, we used the cacoa R package^98^ to quantify cell-type composition differences in SCZ. We used two analysis strategies: 1) globally considering all cell-level SpDs together, and 2) within each individual cell-level SpDs. We found no evidence of global cell-type composition changes (**Figure 5D**, **Figure S30**). To investigate whether specific cell types differ in abundance, and in which cell-level SpDs these differences occur, we examined the proportion of each cell type within each cell-level SpD. We found no layer-specific differences in cell-type composition, except in L2/3 where there was a significant enrichment for L6 Ex neurons in SCZ (*p*=0.002). However, because L6 Ex neurons represent a trivial proportion of cells in this domain (1.68% to 4.26%) (**Figure S31**), this finding should be interpreted with caution. Notably, we did not observe a compositional difference in microglia in any of the layers. Gene expression down-regulation has previously been observed in microglia,^17,46^ and taken together with our results, which showed down-regulation of microglial genes (**Figure 2A**, **Figure 2E**, **Figure 3A**, **Figure 3C**), this suggests microglia are likely undergoing intrinsic expression changes in SCZ.^86^ Overall, our compositional analysis did not provide strong evidence for substantial changes in cell-type abundance between NTC and SCZ, supporting the interpretation that previously observed SCZ-associated gene expression changes reflect intrinsic transcriptional changes.

To identify transcriptional changes associated with SCZ as a validation of our Visium results, we first performed layer-adjusted DE analysis on the Xenium data by quantifying expression changes associated with SCZ (**Methods**). Considering only effect sizes and not statistical significance, the majority (n=124 genes; 73.8%) of the 168 layer-adjusted SCZ-DEGs from Visium showed consistent directionality with the Xenium dataset (**Figure 5E**, Pearson’s correlation=0.502). Specifically, 72 genes were down-regulated and 52 genes were up-regulated in both analyses. Furthermore, in the Xenium analysis, we identified a total of 19 statistically significant layer-adjusted SCZ-DEGs with FDR < 0.10 (**Figure 5F**). 13 of the 19 genes were down-regulated in SCZ, again showing concordance with previous bulk and snRNA-seq studies.^16,41^ We confirmed significant differential expression of 16 of the genes identified in the Visium data. These include up-regulated *FKBP5*, *SERPINA3*, *CHI3L1*, and down-regulated *NPTX2*, *IFITM3*, and *TNFSF10*, spanning neuroplasticity and neuroinflammation, with mixed directions of changes. Of the 16 genes that were also found in the Visium layer-adjusted DE analysis, 15 genes showed the same direction of expression change. The only discrepancy in directionality was *IFITM3*, which was up-regulated in the Visium data, but down-regulated in the Xenium data. The three additional genes identified in the Xenium analysis were *KLF2*, *SLC44A1*, and *MBP*. These genes are primarily expressed in microglia (*KLF2*)^99^ and oligodendrocytes (*SLC44A1*, *MBP*),^25^ providing orthogonal evidence for cell-type-specific transcriptional alterations across non-neuronal cell types in SCZ, in concordance with our earlier findings (**Figure 3A**–**Figure 3D**). Taken together, the results from both the Visium and Xenium layer-adjusted analyses consistently identified genes associated with plasticity and cell-state dynamics, suggesting alterations in these pathways in SCZ.

Given that the layer-adjusted analysis allowed us to identify SCZ-DEGs while controlling for differences in cortical layer composition between donors, we hypothesized that controlling for cell-type composition could yield additional insight into transcriptional changes. To this end, we next performed cell-type-adjusted DE analysis (**Methods**). We identified 15 significant DEGs at FDR < 0.10 (**Figure 5G**). Up-regulation of *FKBP5*, *SERPINA3*, *IFITM3*, *GFAP*, and *GSTM5* suggest stress and glial reactivity in SCZ. In concordance with previous results, we observed down-regulation of mitochondrial-associated genes such as *UQCRFS1* and *COX5A*, suggesting reduced expression of oxidative phosphorylation-related pathways, changes that have been linked to a hypometabolic state in SCZ.^100^ We also observed down-regulation of *NPTX2*, *VGF*, and *ARPP19*, suggesting impairment in synaptic plasticity.^101–103^ Interestingly, down-regulation of the immediate early gene *NPTX2* has been implicated in SCZ.^104^ Up-regulation of the myelin-related genes *MBP* and *BCAS1* were also observed in SCZ in the cell-type-adjusted analysis (**Figure 5G**), supporting previous findings of myelin dysfunction.^97,105^ Taken together, the cell-type-adjusted DE model revealed intrinsic transcriptional changes in SCZ, independent of cell-type composition.

Next, we leveraged the single-cell resolution of the Xenium data to identify transcriptional changes occurring in discrete cell populations (**Methods**). Using this “cell-type-specific” DE model, we found significant gene expression changes in microglia and L4/5 Ex neurons at FDR < 0.10 (**Figure 5H**). In microglia, we observed significant down-regulation of *KLF2* and *ABCG2*, genes involved in cellular stress responses, consistent with the overall reduction of microglia-associated gene expression reported in previous studies.^17,46^ Interestingly, *KLF2* was also down-regulated in the Xenium layer-adjusted DE analysis, highlighting the ability of the cell-type-specific model to pinpoint the specificity of gene expression changes and map them to cellular populations. In L4/5 Ex neurons, we observed up-regulation of *CALB2* and down-regulation of *BDNF* and *ADCYAP1* in SCZ. Given that *CALB2* is associated with neuronal excitability, and *BDNF*/*ADCYAP1* have roles in synaptic strengthening and dendritic remodeling,^106^ this pattern may reflect altered synaptic plasticity in deep layer neurons in SCZ. Similar to the observations for microglia, we did not find compositional differences in L4/5 Ex neurons between NTC and SCZ. Taken together, these findings highlight transcriptional dysregulation in cellular stress and neuroplasticity within discrete cellular populations, rather than alterations in cell-type abundance.

Finally, we profiled expression patterns of some SCZ risk genes compiled from GWAS and other literature (**Supplementary Table 8**, Column G), including *GRIN2A*, *COMT*, *DISC1*, *SV2A*, and *C4A*.^13,107–110^ Although these genes did not show significant differences across the three Xenium DE models described above (**Supplementary Table 9**), several genes were expressed in specific cell types. *ZNF804A*, a TF previously linked to SCZ,^13,80^ was enriched in the MGE inhibitory neuron populations despite no significant diagnostic-group differences (**Figure S32**). Based on our earlier TF analysis (**Figure 4D**, **Figure S21**), we identified ZNF804A as a possible TF linked to the layer-restricted SCZ-DEGs in the Visium SpDs (SpD07-L1/M, SpD06-L2/3, SpD01-WMtz). Given the localization of these neurons, this suggests that the dysregulation of *ZNF804A* in MGE inhibitory neurons, which are endogenous to these layers, could drive the observed layer-restricted SCZ-DEGs, contributing to a potential cell-driven mechanism for the disruption in transcriptional changes occurring in SCZ.

### 2.6 SCZ-associated differences in gene expression map to discrete, spatially-organized cellular microenvironments

Cells are organized into distinct subcellular and microenvironment compartments that support specialized functions and shape local gene expression.^111,112^ We delineated SPG-defined cellular microenvironments enriched for non-nucleated neuropil (DAPI-free), neuronal nuclei (NeuN), PNN (WFA), and vascular compartments (Claudin-5), and showed that SCZ-DEGs, including laminar- and bulk-level transcriptional signatures, were differentially enriched across these microenvironments (**Figure 2D**, **Figure 3E**, **Figure S19**). We next directly examined diagnosis-associated gene expression changes within neuropil, PNN, and neurovascular microenvironments (**Figure 6A**, **Figure 6B**, **Supplementary Table 10**, **Methods**). We first pseudobulked neuropil, PNN, and neuronal spots within individual gray-matter SpDs (e.g., L1-L6). Given the broad distribution of blood vessels in both WM and gray matter, we pseudobulked vasculature spots across all SpDs (e.g., L1-L6 and WM). For each SPG spot classification, we evaluated SCZ-associated differences within the microenvironment while controlling for laminae. At FDR < 0.10, we identified 352, 0, 90, and 19 significant SCZ-DEGs restricted to neuropil, PNN, neuronal, and vasculature microenvironments, respectively. To capture a broader range of candidate transcriptional changes and enhance sensitivity, we examined genes at nominal *p* < 0.05. Using this threshold, we identified: neuropil (n=1,669 genes), PNN (n=559 genes), neuronal (n=1,789 genes), and vasculature (n=440 genes). The neuropil and neuronal microenvironments still exhibited the most transcriptional changes (**Figure 6A**, **Figure S36**), suggesting stronger SCZ-associated changes within local niches that are enriched for neuronal components.^13,16,53–55^ This relaxed approach identified several additional SCZ-DEGs including *NRXN2* in neuropil, and *SST* and *MAPK3* in the neuronal microenvironments (**Figure 6B**). Additional SCZ-DEGs and their microenvironment-restricted gene expression patterns included *GABRA1* for PNNs and *A2M* for vasculature (**Figure 6B**, **Supplementary Table 10**).

**Figure 6.**
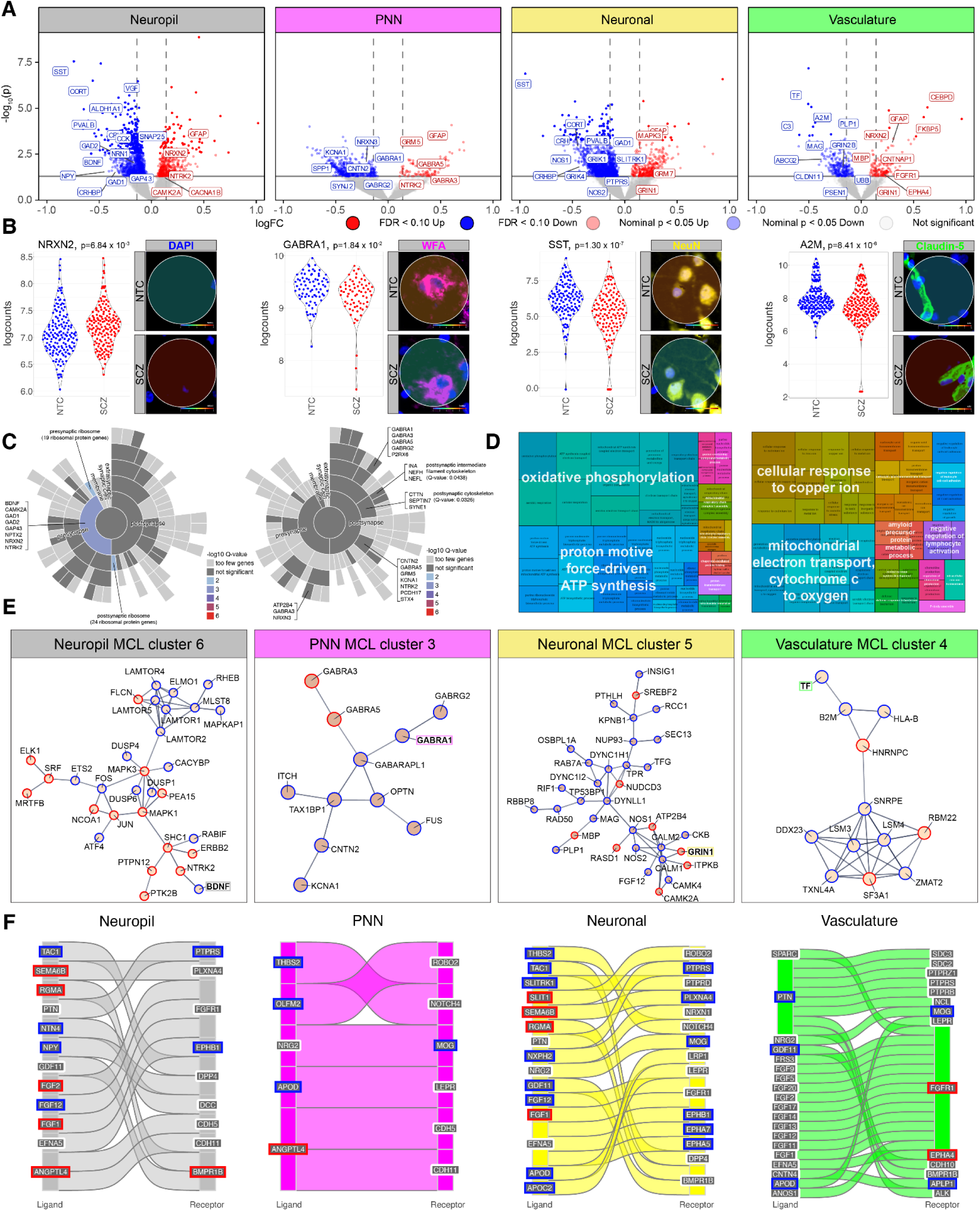
SCZ-linked differential gene expression changes across SPG-defined microenvironments in the human dlPFC. (A) Volcano plots showing microenvironment-restricted SCZ-DEGs identified in four SPG-defined microenvironments. SCZ-DEGs were identified at two significance thresholds: stringent (FDR < 0.10; red=up-regulated, blue=down-regulated; neuropil: 352, PNN: 0, neuronal: 90, vasculature: 19) and nominal (*p* < 0.05, horizontal solid line, semi-transparent red=up, semi-transparent blue=down; neuropil: 1,669, PNN: 559, neuronal: 1,789, vasculature: 440). Non-significant genes are shown in gray. Vertical dashed lines indicate reference lines at |logFC|=0.14. Representative genes are highlighted. (B) Representative microenvironment-restricted SCZ-DEGs are highlighted with their Visium spots, paired with dot plots showing their expression changes in SCZ: (i) *NRXN2* (neuropil), (ii) *GABRA1* (PNN), (iii) *SST* (neuronal), (iv) *A2M* (vasculature). Each data point represents pseudobulked gene counts from an SPG-defined microenvironment within each SpD of each donor (donor-SpD-microenvironment; blue for NTC and red for SCZ group). Spot diameter: 55 µm, Scale: logcounts. Interactive plots are available for viewing and download in both the iSEE app and Samui browser (Data Availability). (C) SynGO enrichment analyses of neuropil and PNN microenvironment-restricted SCZ-DEGs identified at nominal *p* < 0.05. Cellular Component (CC) terms significantly enriched at *q*-value < 0.01 are shown, with colors indicating enrichment significance (-log_10_ (*q*-value)) and representative genes labeled. For PNNs, selected terms significant at relaxed threshold (*q*-value < 0.05) were labeled manually to indicate significant enrichment of transcripts for postsynaptic elements. Additional supporting SynGO results for PNN using a default ‘brain-expressed’ background are provided in **Figure S33**. (D) GO-ORA of neuronal and vasculature microenvironment-restricted SCZ-DEGs identified at nominal *p* < 0.05. BP terms were tested against matched microenvironment-specific background sets, defined by all genes expressed in the respective microenvironments. Treemaps depict the top 50 significantly enriched GO-BP terms (FDR ≤ 0.05) after semantic similarity reduction (threshold=0.7). An additional neuron-focused treemap for neuronal SCZ-DEGs, generated by selectively filtering neuron-related GO-BP terms, is provided in Figure S34. (E) STRING-based protein-protein interaction (PPI) networks of microenvironment-restricted SCZ-DEGs were generated for each SPG-defined microenvironment (neuropil, PNN, neuronal, vasculature). Within each PPI network, functional clusters were identified using the built-in MCL algorithm (inflation=1.2). For each microenvironment, one representative cluster is shown here as an example of a microenvironment-restricted signaling subnetwork where a single representative gene within that subnetwork (*BDNF*, *GABRA1*, *GRIN1*, or *TF*) is highlighted with a colored box. Blue indicates down-regulation and red indicates up-regulation of gene expression in SCZ. (F) Sankey representation of candidate SCZ-linked LR interactions from Huuki-Myers et al.^25^ that match microenvironment-restricted SCZ-DEGs identified at nominal *p* < 0.05. LR pairs are shown when either the ligand or receptor intersects with one of SCZ-DEGs, with blue indicating down-regulation and red indicating up-regulation of gene expression in SCZ. The partner genes shown in gray were not significantly differentially expressed within the microenvironments in SCZ.

To infer broad functional implications of these SCZ-DEGs, we combined up- and down-regulated genes within each microenvironment and performed gene ontology (GO) enrichment analyses (**Figure 6C**, **Methods**). Given the relevance to synaptic and ECM functions, we used the SynGO web portal to interpret the neuropil and PNN microenvironment-restricted SCZ-DEGs, while enrichment analysis of general GO terms was applied to neuronal and vasculature microenvironment-restricted SCZ-DEGs. Neuropil SCZ-DEGs were significantly enriched for pre-synaptic compartments and ribosomal machinery, suggesting altered local synaptic translation and plasticity.^113–115^ PNN SCZ-DEGs (e.g., *GABRG2*, *NTRK2*) were enriched for synaptic components, with stronger enrichment in postsynaptic compartments, which may preferentially affect stabilization of synaptic inputs onto PVALB neurons surrounded by PNNs (**Figure 6C**, **Figure S33**).^116–119^ Neuronal SCZ-DEGs featured pathways associated with mitochondrial metabolism, cellular respiration, and energy production (**Figure 6C**). Enrichment of neuron-specific processes included neuronal projection, calcium ion-mediated neurotransmission, and neuronal apoptotic processes (**Figure S34**). Vasculature SCZ-DEGs showed enrichment in metabolic pathways, metal-ion homeostasis, and immune functions, with some enrichment for amyloid precursor protein metabolism (**Figure 6C**).

To complement these functional annotations and identify potential signaling pathways among SCZ-DEGs, we performed STRING network analysis restricted to physical protein-protein interactions, and clustered gene modules into subnetworks (**Methods**, **Figure 6E**, **Supplementary Table 11**).^77^ Across all microenvironments, more than 70% of genes within the top five MCL subnetworks converged on mitochondrial metabolism, RNA processing, and protein homeostasis, with SCZ-DEGs repeatedly shared across microenvironments and enriched for housekeeping pathways such as ATP metabolism and oxidative phosphorylation (**Figure S35**, **Figure S36**, **Figure S37**, **Supplementary Table 11**). Additionally, we identified subnetworks that recapitulate specialized biological functions of each microenvironment (**Figure S38**, **Supplementary Table 11**). Neuropil subnetworks highlighted genes associated with neuronal functions and signaling, including mTOR signaling pathways linked to BDNF-TrkB signaling (**Figure 6E**, MCL cluster 6), synaptic vesicle trafficking (MCL cluster 5), GABAergic signaling (MCL cluster 11), and complement-mediated neuroimmune responses (MCL cluster 8). Neuronal subnetworks also emphasized neuronal communication, particularly calcium-mediated signaling (**Figure 6E**, MCL cluster 5) and complement activation (MCL cluster 44). PNN subnetworks were marked by representation of GABAergic signaling (**Figure 6E**, MCL cluster 3) with multiple GABA receptor subunits (*GABRA1*/*3*/*5*, *GABRG2*). Vasculature subnetworks showed enrichment of SCZ-DEGs not only for vascular growth (MCL cluster 13) and NMDA receptor signaling (MCL cluster 10) but also for RNA processing linked to immune presentation and iron regulation (**Figure 6E**, MCL cluster 4). Collectively, these analyses organized SCZ-DEGs into functionally relevant subnetworks that point to shared metabolic perturbations and microenvironment-restricted signaling.

Building on our pathway-level analyses, we next examined how SCZ-associated transcriptional changes relate to cell-cell communication (CCC). We leveraged a curated set of 90 candidate ligand-receptor (LR) pairs from Huuki-Myers et al.,^25^ which integrated dlPFC snRNA-seq CCC analysis with SCZ-linked LR annotations from OpenTargets and Omnipath.^120,121^ We mapped these interactions to contextualize how transcriptional dysregulation may coordinate altered intercellular communication within microenvironments. Among the 90 LR pairs, we identified 13, 17, 6, and 26 pairs in which at least one interactor matched a microenvironment-restricted SCZ-DEG in neuropil, neuronal, PNN, and vasculature microenvironments, respectively (**Figure 6F**, **Figure S39**, **Figure S40**, **Methods**). Neuropil SCZ-DEG-matched LR pairs included those associated with axon-guidance and growth factor signaling (NTN4-DCC, EFNA5-EPHB1, SEMA6B-PLXNA4, FGF-FGFR1). Neuronal SCZ-DEG-matched LR pairs included growth- and metabolism-linked signaling (PTN-PTPR, APOD-LEPR), as well as adhesion complexes (EFNA5-EPHA5/7, NXPH2-NRXN1). PNN SCZ-DEG-matched LR pairs were characterized by ECM-, glial-, and vascular-associated interactions, such as ANGPTL4-CDH5/11 and THBS2-NOTCH4. Vasculature SCZ-DEG-matched LR pairs involved trophic signaling interactions, including FGF-FGFR1 and PTN axes, along with adhesion pairs such as EFNA5-EPHA4 and CNTN4-APLP1. Overall, we observed recurrent trophic axes (e.g., PTN-PTPR and FGF-FGFR1), which might suggest upstream signaling components shared across these specialized microenvironments.^122,123^ Taken together, integrating SCZ-DEGs with spatially-resolved LR interactions highlights candidate signaling mechanisms that may contribute to intercellular dysregulation within microenvironment domains.

### 2.7 Spatial mapping of eQTLs and colocalization with SCZ GWAS resolve context-dependent regulatory effects across dlPFC compartments

To investigate the effect of genetic variants on spatially-resolved gene expression, we integrated DNA genotype data from all 63 donors and identified expression Quantitative Trait Loci (eQTLs) using tensorQTL (**Figure 7A**, **Methods**).^124,125^ We identified nominal and independent cis-eQTLs within each SpD-defined cortical layer and SPG-defined microenvironment (**Methods**). Genes from the independent cis-eQTL variant-gene pairs (eGenes, genes containing at least one significantly associated cis single nucleotide polymorphism)^126^ were most frequently identified in neuronal-rich SpDs and microenvironments (**Figure 7B**, **Supplementary Table 12**), consistent with observations that SCZ genetic risk is enriched in these domains (**Figure 4H**). The identified nominal cis-eQTL associations included *SST* (rs17602709) in SpD05-L5, *CSMD1* (rs6559030, rs718464) in SpD01-WMtz, and *BDNF* (rs72884024, rs4923559) and *FKBP5* (rs35522880) in neuropil (**Supplementary Table 12**).

**Figure 7.**
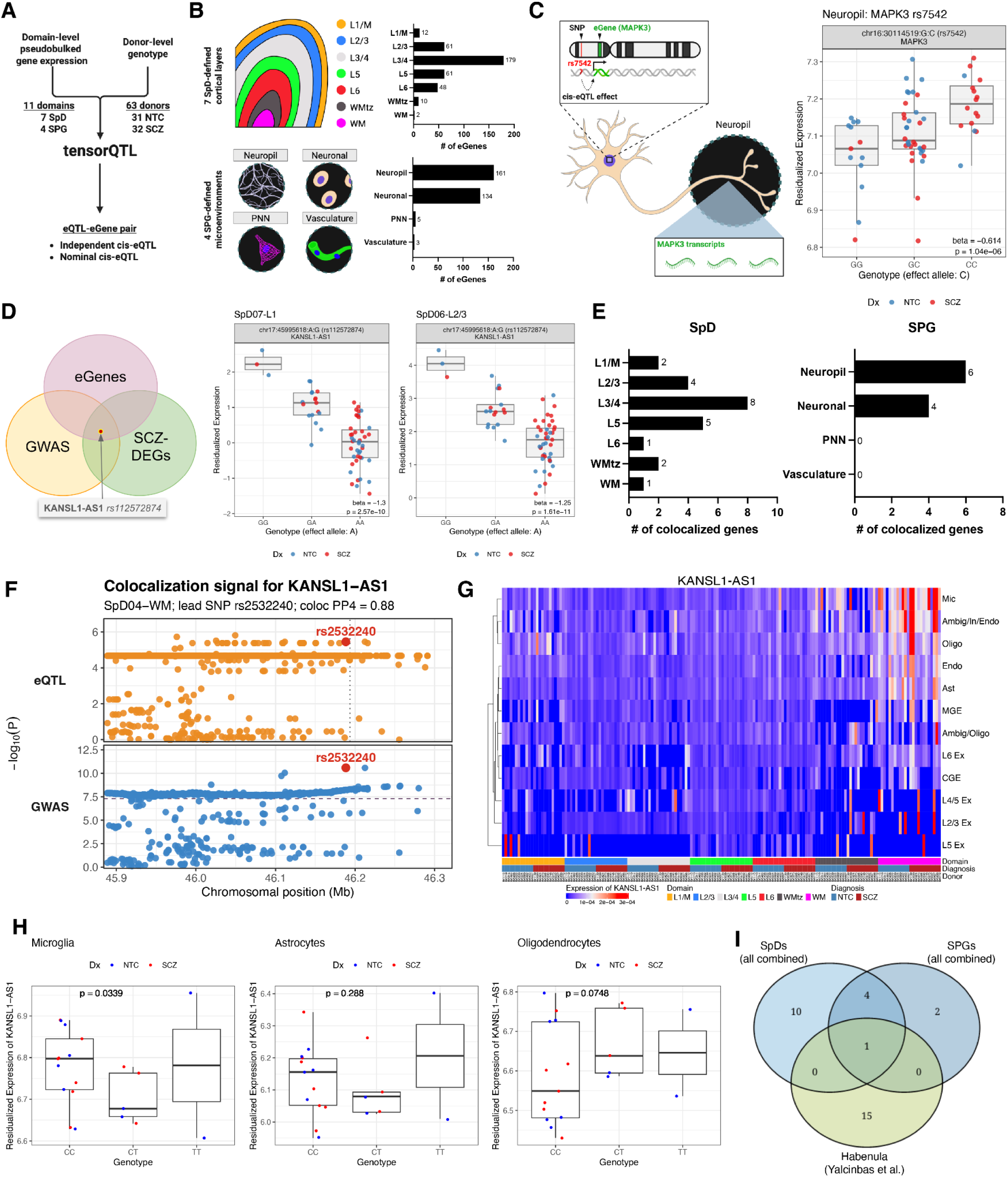
Spatial eQTL mapping and colocalization analyses across cortical layers and microenvironments in SCZ. (A) Overview of spatial eQTL mapping across the dlPFC cohort. Cis-eQTL mapping was performed using donor-level genotype data (n=63) and gene expression profiles pseudobulked for cortical layers and microenvironments. tensorQTL was used to identify cis-eQTL-eGene pairs across 7 SpDs and 4 SPG microenvironments, including independent and nominal cis-eQTLs. Created in BioRender. Kwon, S. H. (2026) https://BioRender.com/sw4sj78 (B) Bar plots summarizing the number of independent eGenes associated with spatial eQTLs identified across SpDs and SPG microenvironments, illustrating their distribution across spatial contexts. Created in BioRender. Kwon, S. H. (2026) https://BioRender.com/sw4sj78 (C) Representative example of a spatial independent eQTL signal for *MAPK3* (rs7542) in neuropil microenvironments. The schematic illustrates a cis-eQTL relationship and emphasizes that *MAPK3* transcripts were detected within neuropil regions corresponding to non-somatic neuronal compartments. Box plots show the genotype-expression association pattern across genotypes (blue for NTC and red for SCZ group). Created in BioRender. Kwon, S. H. (2026) https://BioRender.com/sw4sj78 (D) Identification of rs112572874-*KANSL1*-*AS1* as one of the two candidate independent SNP-gene pairs intersecting spatial eQTLs, layer-adjusted SCZ-DEGs, and SCZ GWAS loci. Venn diagram illustrates overlap across the three dimensions. Box plots show genotype-associated expression patterns for *KANSL1*-*AS1* (rs112572874) in SpD-defined cortical layers (L1/M and L2/3, blue for NTC and red for SCZ diagnosis group). The other candidate independent SNP-gene pair intersecting spatial eQTLs, *MAPK3* (rs7542), is visualized in **Figure S41**. (E) Summary of colocalization analyses across dlPFC layers and microenvironments. Bar plots show the number of genes with strong evidence of colocalization (PP4 ≥ 0.8) between eQTL and SCZ GWAS (European ancestry) tracks across SpD-defined cortical layers and SPG-defined microenvironments. (F) Colocalization signal for *KANSL1*-*AS1* (rs2532240) in SpD04-WM. Manhattan plots display -log_10_(*p*) values for eQTL and GWAS signals across genomic position, highlighting shared loci with strong colocalization evidence. eQTL mapping was performed within a ±1 Mb cis window, and the panel presents a zoomed-in view of the locus near the lead SNP (rs2532240, shown in red). (G) Cell-type-resolved expression of *KANSL1-AS1* from Xenium data. Heatmap shows pseudobulked expression across donor-SpD combinations (columns) and cell types (rows), based on log_2_-transformed, normalized counts. Key cell type abbreviations: MIC, microglia; Oligo, oligodendrocytes; Ast, astrocytes. Additional details on cell type labels are provided in **Figure S29** and **Supplementary Table 8**. (H) Genotype-associated expression patterns of *KANSL1*-*AS1* (rs2532240) in WM glial cell types. Box plots show expression across genotypes in microglia, astrocytes, and oligodendrocytes, where each data point represents gene expression counts aggregated at the donor level (blue for NTC and red for SCZ diagnosis group). *p*-values for differences between the C:C and C:T genotypes are shown in each plot. (I) Comparison of colocalized genes identified in this study with those detected in human habenula bulk RNA-seq data.^125^ Venn diagram depicts overlap across all 7 spatial domains (SpDs), all 4 categories of microenvironments (SPGs), and habenula-derived colocalization genes.

To assess the potential relevance of these spatially-resolved eQTLs (hereafter, spatial eQTLs) to SCZ etiology, we intersected them with SCZ GWAS risk loci^13^ and SCZ-DEGs identified in our layer-adjusted and microenvironment-level analyses (**Figure 2A**, **Figure 6A**, **Supplementary Table 12**). Cis-eQTL variant-gene pairs with overlapping signals across genetic association (GWAS), genotype-dependent regulation (eQTL), and disease-associated expression changes (SCZ-DEGs) represent candidates that may link genetic risk to downstream transcriptional alterations in SCZ within spatially defined domains. This integrative approach highlighted two intersecting genes, *MAPK3* and *KANSL1-AS1* (**Figure S41**, **Figure 7C**, **Figure 7D**). *MAPK3* showed an independent cis-eQTL signal (rs7542) within neuropil, and the same variant was also detected as a nominal cis-eQTL in neuronal and vasculature microenvironments (**Figure 7C**, **Figure S41**, **Supplementary Table 12**). The more robust signal observed in neuropil, compared to neuronal and vasculature, may suggest regulatory effects that are spatially localized to non-somatic compartments. *KANSL1-AS1* exhibited an independent cis-eQTL signal (rs112572874) in SpDs for L1/M and L2/3, supporting laminar context of genotype-dependent regulatory effects (**Figure 7D**).

To complement the spatial eQTL mapping analyses, we performed colocalization analysis (**Methods**) to evaluate whether the eQTL and SCZ GWAS signals share underlying causal variants, thereby linking genetic risk to regulatory effects. This analysis identified 15 genes across SpDs and 7 genes across microenvironments (neuropil and neuronal) with strong evidence of colocalization (PP4 ≥ 0.8) (**Figure 7E**, **Supplementary Table 13**). SCZ colocalization was mostly detected in SpD02-L3/4 (*APC2*, *ARL17B*, *DNPH1*, *LY6H*, *RNASEH2C*, *RPS17*, *SLC25A27*, and *SMG6*) and in the neuropil (*ARL17B*, *CDHR1*, *DNPH1*, *LY6H*, *RPS17*, and *SLC25A27*). *SLC25A27* showed SCZ colocalization across nearly all domains, except SpD01-WMtz. For *KANSL1-AS1*, we identified a variant (rs2532240) showing a nominal cis-eQTL association in neuropil and neuronal-rich domains in the initial tensorQTL mapping analysis (**Supplementary Table 12**), but not in WM. Intriguingly, the same variant demonstrated evidence of strong SCZ colocalization in SpD04-WM (**Figure 7F**). *KANSL1-AS1* was previously identified as both a layer-adjusted SCZ-DEG (**Figure 2A**) and a PRS-associated DEG (**Figure 4E**, **Figure 4F**). Because *KANSL1-AS1* was included in our Xenium panel (**Supplementary Table 8**), we examined its expression pattern at cellular resolution, which revealed higher expression in glial cell types within WM compared to other cell types (**Figure 7G**). We then specifically evaluated genotype-associated expression patterns of this variant in cell types with higher expression of *KANSL1-AS1* (microglia, astrocytes, and oligodendrocytes in the WM). Despite limited genotype representation in the Xenium cohort, we observed potential genotype-associated differences, particularly between the C:C and C:T genotypes (**Figure 7H**), within microglia (p=0.0339) and more limited yet similar trend in astrocytes (p=0.288). In contrast, oligodendrocytes showed changes in the opposite direction (p=0.0748), suggesting that these regulatory effects may act in a cell-type-specific context.

Because genetic regulation of transcription can differ across brain regions,^41^ we asked whether our spatially-resolved colocalization signals are specific to the dlPFC or shared with other brain regions. We leveraged a bulk RNA-seq dataset that shared some overlapping donors with our study (n=10), which compared control and SCZ in the human habenula, a region with significantly different cell-type composition compared to the dlPFC.^125^ Using these data, we performed cross-region comparisons of genes with colocalization signals (**Supplementary Table 13)**. We identified several SCZ colocalization genes that were not detected in the habenula dataset, including *ARL17B*, *DNPH1*, *LY6H*, and *RPS17* (**Figure 7I**, **Supplementary Table 13**). *LY6H* encodes a neuronal membrane protein involved in nicotinic acetylcholine receptor signaling, a pathway implicated in cognition and SCZ,^127,128^ and is associated with SCZ as a protein QTL (pQTL).^129^ We identified strong SCZ colocalization signals (PP4 ≥ 0.8) for *LY6H* across multiple neuronal-rich dlPFC layers (SpD02-L3/4, SpD05-L5) and microenvironments (neuropil and neuronal) (**Figure S42**). *LY6H* showed nominal genotype-associated expression changes (SpD02-L3/4, nominal *p* < 0.0001; neuronal and neuropil microenvironments, nominal *p* < 0.0001; **Supplementary Table 12**) and nominal SCZ-associated expression changes in neuronal microenvironment (nominal *p=*0.025; **Supplementary Table 10**).

Finally, comparison with prior SCZ-related gene-prioritization resources, including an adult multi-region brain eQTL/GWAS candidate-causal gene set,^130^ a developing-brain xQTL colocalization atlas,^131^ and the PsychENCODE SCZ risk-gene resource,^132^ revealed that 11 of our colocalization genes are unique to the spatial dlPFC analysis in this study (**Figure S43**, **Supplementary Table 13**). *LY6H* and *KANSL1-AS*1 were among these genes, along with *APC2*, *CDHR1*, *EFEMP1*, and *SETDB2,* spanning distinct cortical compartments across layers and microenvironments. Together, these results suggest that spatially resolved analysis can highlight regulatory signals not captured by bulk or non-spatial reference resources, indicating that genetic effects may be preferentially localized across cortical compartments and contribute to disease-associated changes within spatial-anatomical contexts.

## 3 Discussion

Here we applied complementary spatial transcriptomic approaches, integrating Visium SRT, spatial proteogenomic (SPG) profiling and Xenium *in situ* analysis to interrogate SCZ-associated biology across multiple spatial resolutions in the human dlPFC. Data-driven SpDs derived from the Visium data recapitulated classic cortical cytoarchitecture while also capturing finer molecular and anatomical features, including the superficial L1/M domain and the gray-white matter transition zone (WMtz).^25,133,134^ DE analyses across these SpDs revealed spatially localized disease-associated transcriptional changes that were most prominent in glia-and vascular-enriched domains (L1/M, WM, and WMtz). These findings highlight the ability of spatial domain-based analyses to resolve disease-associated signals in anatomical features that may be underrepresented or obscured in dissociative approaches, including L1/M and transitional white matter.^135–137^

Given the importance of cellular compartments in shaping local gene expression,^111,112^ we leveraged protein-guided image segmentation to resolve transcriptional patterns within defined cellular microenvironments, including neuropil, neuronal, PNN, and vascular compartments. Within these microdomains, SCZ-associated changes were enriched for neuronal and synaptic biology programs, particularly within neuropil- and neuronal-associated microdomains, indicating that spatial resolution and compartmental context are critical for detecting disease-associated signals. We provided orthogonal validation and cellular context using targeted Xenium profiling, which showed that cell-type abundances were largely similar across diagnosis, supporting the interpretation that laminar- or bulk-level gene expression changes primarily reflect intrinsic cellular alterations rather than shifts in cell-type composition. Cell-resolved analyses further localized laminar-level signals to specific populations, for example, mapping *BDNF*-associated transcriptional changes to L4/5 excitatory neurons. Together, these analyses demonstrate that integrating multiple scales links laminar- and microenvironment-level changes to cell-type-specific perturbations to reveal complementary aspects of SCZ-associated biology.

Within this framework, we observed transcriptional alterations in neuronal-rich gray-matter SpDs, particularly in L2/3, consistent with prior studies implicating neuronal pathology in SCZ. We additionally detected deep-layer (L6) excitatory neuronal signatures within superficial layers (L1-3) in SCZ, suggesting the presence of molecular signatures that deviate from canonical laminar organization. This pattern may reflect alterations in developmental neuronal migration^138–140^ or contributions from transcripts localized to distal neuronal processes.^74,141,142^ Consistent with this interpretation, we observed more robust neuronal gene expression changes within neuropil microenvironments, with additional changes detected in neuronal, PNN, and vascular microenvironments. To our knowledge, this study is the first to analyze transcription associated with neuropil, PNNs, and vasculature in the human cortex, establishing a framework for molecular profiling within these specialized synaptic, extracellular, and neurovascular niches. Ligand-receptor analyses further contextualized these findings by nominating candidate trophic and adhesion signaling interactions within these compartments, including FGF-FGFR1 and PTN-PTPR interactions,^122,123^ which represent potential targets for future mechanistic investigation.

In interpreting these findings, it is important to emphasize that gene expression changes identified through case-control comparisons cannot readily distinguish transcriptional alterations related to disease pathogenesis from those that are illness-associated factors, and may primarily reflect illness state. These include general health-related effects of chronic illness, medication exposure, substance use, and technical factors related to postmortem RNA quality. As such, we cannot determine the extent to which these factors contribute to the observed transcriptional differences.^41,143,144^ To partially mitigate these limitations, we integrated analyses using genetic information from study donors, as genetic risk is not influenced by illness-associated or postmortem confounds. Polygenic risk was predominantly localized to neuronal-rich SpDs, consistent with evidence that neurons constitute the primary substrate of genetic vulnerability, whereas transcriptional alterations were more prominently localized to glial- and vascular-rich SpDs. These observations indicate that genetic risk and transcriptional alterations localize to distinct spatial contexts, but their relationship to one another is unresolved. One possibility is that neuronal genetic susceptibility influences non-neuronal transcriptional programs through neuron-glia signaling.^145–147^ Alternatively, these non-neuronal transcriptional patterns may reflect downstream or compensatory processes associated with illness state. Resolving these possibilities will require future studies using experimental models.

A key aspect of our study is the integration of laminar, microenvironment, and cellular context for genes associated with SCZ risk. These include *ZNF804A*, *MAPK3*, and *KANSL1-AS1*, which map to established SCZ risk loci.^13^ At the laminar-level, TF analyses predicted ZNF804A-associated regulons spanning multiple SpDs, including L1/M, L2/3, and WMtz with SCZ-DEGs target genes, including *BDNF* and *DCC*. Xenium data further localized *ZNF804A* to MGE-derived inhibitory neurons, providing cellular context for its potential contribution to GABAergic dysfunction in SCZ. Similarly, *MAPK3* and *KANSL1-AS1* were identified as both layer-adjusted SCZ- and PRS-DEGs. Integrating spatial and microenvironment information with eQTL and colocalization analyses further refined the context of genotype-associated regulatory effects. For example, *MAPK3* eQTL signals localized to neuropil whereas *KANSL1-AS1* localized to WM glial populations. In addition, *LY6H*, an established SCZ-associated pQTL,^129^ showed dlPFC-specific cis-eQTL effects that localized to L3/4 SpD and neuronal microenvironments. Together, these findings demonstrate how integrating spatial and anatomical context can provide biological insight into potential mechanisms for how SCZ risk genes mediate disease.

Our transcriptional signatures also provide insight into biological pathways associated with circuit-level alterations in SCZ. Down-regulation of activity-dependent synaptic genes (*BDNF*, *VGF*, *NPTX2*) and inhibitory neuron markers (*SST*, *CORT*, *PVALB*) is consistent with disruption of trophic and inhibitory signaling in SCZ. In this context, dysregulation of *NTRK2,* which encodes the BDNF receptor TrkB, was observed in PNN microenvironments. BDNF-TrkB signaling is known to regulate the function of PVALB neurons, including their ability to generate gamma oscillatory activity, providing a potential link between these transcriptional changes and circuit-level dysfunction in SCZ.^118,119,148,149^ *FKBP5*, a key mediator of glucocorticoid-mediated stress signaling, was up-regulated in SCZ,^150–152^ and pathways related to mitochondrial metabolism and RNA processing were consistently dysregulated across analyses.^18,90,153–155^ We also observed spatially divergent glial-vascular changes, including up-regulation of astrocytic programs (*GFAP*, *SERPINA3*), attenuation of microglial signatures (*AIF1*, *TREM2*, *C3*), and heterogeneous oligodendrocyte- and vascular-associated signals in SCZ. The down-regulation of microglia-associated genes was notable, warranting further investigation on the potential for reconfigured cellular states, given their role in synapse pruning and neuroinflammation.^46^ Together, these findings indicate coordinated alterations across synaptic, stress-related, metabolic, and non-neuronal gene programs that may contribute to cortical vulnerability in SCZ.

In summary, we provide a resource-scale atlas that maps the spatio-molecular neuroanatomy of the dlPFC in SCZ. By integrating analyses across laminar domains, microenvironments, and cell types, we show how disease-associated transcriptional programs are organized within spatial-anatomical context. These findings suggest that SCZ-associated biology spans neuronal and non-neuronal contexts, with genetic risk and transcriptional alterations localizing to distinct but complementary spatial domains. We also provide publicly available web browsers (Samui Browser, iSEE) to facilitate exploration of these data and support hypothesis generation and data-driven discovery.

### 3.1 Limitations

The biological heterogeneity of SCZ and cohort size may limit statistical sensitivity for detecting spatially localized and genetically-mediated effects, particularly within the neuronal-rich cortical domains and PNN microenvironments, as well as for analyses of genetic loci with lower minor allele frequencies. Across analyses, we observed neuroimmune- and microglia-associated transcriptional changes, particularly within glial and non-neuronal domains. Interpretation of neuroimmune-associated transcriptional changes in postmortem human brain studies is inherently challenging, and RNA-based measurements do not directly reflect functional immune activity or cellular state dynamics. However, the observed trend toward down-regulation of these genes is not consistent with increased immune activity or inflammation.^46,146,156^ Our data do support the existence of altered neuroimmune-associated transcriptional programs and potential changes in neuron-glial interactions, but do not provide direct support for models of immune activation in SCZ. Postmortem research studies of neuropsychiatric disorders have inherent potential confounds including biased exposure to medications, increased incidence of substance use, and disease-associated comorbidities, which complicate assigning results to cause versus consequence.^143^ Inclusion of analyses that incorporated genetic risk for SCZ helped to mitigate this issue, but these approaches still do not establish causality. Analyses of cell-cell communication and ligand-receptor interactions rely on co-expression and curated interaction databases, and therefore do not establish direct binding, signaling directionality, or functional interaction. Some LR pairs, including NTN4-DCC and THBS2-NOTCH4 have limited experimental evidence for direct binding and would require future experiments to validate. Technical aspects of the platforms also contribute to limitations,^157,158^ including relatively low gene coverage and gene-level resolution, which hinder comprehensive eQTL or transcriptome-wide association analyses. Given that SCZ eQTLs showed transcript specificity in bulk RNA-seq data,^144^ spatial long-read technology will be important for future studies. Here we leveraged single-cell eQTL approaches that pseudobulk gene expression across a specified domain, and thus some spatial eQTL signals are likely conservative and will require replication in larger cohorts. In addition, emerging computational methods should allow future studies to use geospatial statistical models to directly leverage the 2D spatial information in eQTL analyses.

## 4 Methods

### 4.1 Postmortem human tissue samples

Postmortem human brain tissue from the Lieber Institute for Brain Development (LIBD) Human Brain and Tissue Repository was collected through the following sites and protocols at the time of autopsy with informed consent from the legal next of kin: the Office of the Chief Medical Examiner of the State of Maryland, under the Maryland Department of Health IRB protocol #12–24, the Departments of Pathology at Western Michigan University Homer Stryker MD School of Medicine and at the University of North Dakota School of Medicine and Health Sciences under WCG IRB protocol #20111080. Additional samples were consented through the National Institute of Mental Health Intramural Research Program under NIH protocol #90-M-0142, and were acquired by LIBD via material transfer agreement. Details of tissue acquisition, handling, processing, dissection, clinical characterization, diagnoses, neuropathological examinations, and QC measures have been described previously.^159^ Fresh frozen coronal brain slabs from 64 donors were selected for dlPFC sub-dissections. The cohort was divided into two groups of 32 donors, one composed of individuals diagnosed with SCZ and the other composed of NTC. One NTC sample failed anatomical validation and was removed from the study, resulting in a total of 63 dlPFC tissue blocks for the study (one per donor). Cohorts were balanced for age (NTC: mean 46.80 years; SCZ: mean 47.55 years) and sex (NTC: 16 males, 15 females; SCZ: 18 males, 14 females). Fresh frozen coronal slabs at the level of the anterior striatum were rapidly sub-dissected with a hand-held dental drill, perpendicular to the pial surface, targeting Brodmann Area (BA) 46. Cortical brain blocks were approximately 8 X 8 X 5 mm in dimensions, were collected across a sulcus to preserve the integrity of L1, and encompassed the complete laminar structure, spanning cortical L1-L6 and WM. Following dissection, tissue blocks were stored at -80°C in sealed cryogenic bags until cryosectioning. Demographics for the 63 donors are listed in **Supplementary Table 1**. Experimenters were blind to clinical diagnosis of the donors.

### 4.2 Tissue processing and anatomical validation

Tissue blocks were mounted onto round chucks with OCT (Tissue-Tek Sakura), acclimated to the cryostat (Leica CM3050s) to -14°C, ∼50 µm of tissue was trimmed from the block to achieve a flat surface, and several 10-µm sections were collected for anatomical validation (AV). Three AV measures were implemented: 1) visual inspection of the blocks to assess inclusion of the sulcus as well as WM and gray matter; 2) H&E staining to assess cellular integrity of all cortical laminae and the WM; and 3) multiplex RNAscope single molecule fluorescence *in situ* hybridization (smFISH) to ensure presence of molecular markers for all 6 layers and the WM. H&E staining was performed as previously described and images were acquired using an Aperio CS2 slide scanner (Leica) equipped with a 20x/0.75NA objective and a 2x doubler. RNAscope smFISH was performed as previously described,^160^ and images were acquired using the Nikon AX-R confocal microscope equipped with a 2x/0.1 NA objective or a Vectra Polaris slide scanner (Akoya Biosciences) at 20x magnification (0.75 NA). Probes for established marker genes (Advanced Cell Diagnostics, ACD) were used to identify cortical layers in the human cortex,^27^ including *SLC17A7* (Cat No. 415611) indicative of L1-6; *RELN* (Cat No. 413051) and *AQP4* (Cat No. 482441) indicative of L1; *HPCAL1* (Cat No. 846051) indicative of L2; *FREM3* (Cat No. 829021) indicative of L3; *RORB* (Cat No. 446061) indicative of L4; *TRABD2A* (Cat No. 532881) indicative of L5; *NR4A2* (Cat No. 582621) indicative of L6; and *MBP* (Cat No. 411051) indicative of WM. After AV, blocks were again acclimated to the cryostat (Leica CM3050s), mounted onto chucks, scored with a razor into 6.5 X 6.5 mm squares, and 10-µm sections were mounted onto the Visium Spatial Gene Expression Slide (Cat No. 2000233, 10x Genomics). Each Visium slide included four sections from four individual donors, which were also balanced across slides for diagnosis and sex. Adjacent 10-µm tissue sections were mounted onto glass slides for further validation studies (e.g., RNAscope).

### 4.3 Visium-SPG data generation and image preprocessing

#### Visium-SPG staining

IF staining was performed according to the manufacturer’s instruction (10x Genomics, CG000312 Rev D) with slight modifications to avoid staining artifacts introduced by ribonucleoside vanadyl complex (RVC) after methanol fixation. In brief, following cryosectioning, tissue was fixed in pre-chilled methanol, treated with BSA-containing blocking buffer without RVC, and incubated for 30 minutes at room temperature with staining reagents using biotinylated WFA (Sigma Aldrich, Cat No. L1526, 1:200 and 1:600) for perineuronal nets and anti-NeuN antibody conjugated to Alexa Fluor 555 (Sigma Aldrich, Cat No. MAB377A5, 1:50) for neurons. Two different lots of WFA were optimized prior to Visium-SPG staining to account for their lot-to-lot variations. Following a total of 5 washes with RVC-containing buffer, streptavidin conjugated to Alexa Fluor 647 (Thermofisher, Cat No. S32357, 1:400) and anti-Claudin-5 antibody conjugated to Alexa Fluor 488 (Thermofisher, Cat No. 352588, 1:50) was applied for 30 minutes at room temperature to fluorescently label WFA-reactive perineuronal nets and endothelial cells for vascular structures. DAPI (Thermofisher, Cat No. D1306, 1:3000, final 1.67 μg/ml) was used for nuclear counterstaining. The slide was coverslipped with 85% glycerol supplemented with RiboLock RNase inhibitor (Thermofisher, Cat No. EO0384, final 2 U/μl) and scanned on a Vectra Polaris slide scanner (Akoya Biosciences) at 20x magnification (0.75 NA). The exposure time per channel was determined manually, slide by slide, for six different band filters: DAPI (mean 3.2, range 2.3-3.9 msec), Opal 520 (mean 237.8, range 200-290 msec), Opal 570 (mean 240.3, range 200-290 msec), Opal 620 (mean 6.4, range 2-7 msec), Opal 690 (mean 397.2, range 120-550 msec), and autofluorescence (100 msec, fixed) (**Supplementary Table 14**).

#### Visium-SPG cDNA synthesis and library preparation

Immediately after slide imaging, each tissue section was permeabilized for 12 minutes and processed for cDNA synthesis, amplification, and library construction according to the Visium Spatial Gene Expression User Guide (10x Genomics, CG000239 Rev F). The resulting whole transcriptome Visium libraries were sequenced on Illumina NovaSeq 6000 with S4 200 cycle kit using a loading concentration of 300 pM (read 1: 28 cycles, i7 index: 10 cycles, i5 index: 10, read 2: 90 cycles).

#### Spectral unmixing

For image data preprocessing, we applied spectral unmixing methods following the slide scanner manufacturer’s instructions to reduce noise from multi-plexed spectral profiles of Alexa fluorophores and mitigate autofluorescence from postmortem human tissue, such as lipofuscin.^161^ Image preprocessing and spectral unmixing was performed using `Phenochart` Whole Slide Viewer and `inForm` Automated Image Analysis Software (both from Akoya Biosciences). After acquiring image data using a Vectra Polaris slide scanner, the files were subsequently processed in `Phenochart`. Annotations were drawn to outline the entire tissue section for each QPTIFF file, ensuring no overlap between stamped tiles when processed in batch mode using `inForm`. The annotated QPTIFF images were then processed for spectral unmixing in `inForm`. Spectral fingerprints were previously generated from single-positive control samples to establish reference spectral profiles for DAPI and individual Alexa fluorophores and were stored in the `inForm` spectral library.^25,29^ We selected spectral fingerprints for DAPI, Alexa 488, Alexa 555, and Alexa 647 to isolate fluorescence signals corresponding to nuclei, Claudin-5, NeuN, and WFA, respectively. The same unmixing algorithm was applied uniformly to all samples to ensure consistency in image preprocessing and outputs. This process decomposes multi-spectral profiles into spectrally unmixed, multi-channel TIFF tiles using `inForm`. Each multi-component TIFF tile represented a stamped tissue area, with fluorescence signals assigned to separate channels for DAPI, Alexa 488, Alexa 555, and Alexa 647. Autofluorescence, including lipofuscin, was segregated into a dedicated “AF” channel.

#### Image stitching

The spectrally unmixed multichannel tiles of the entire slide (∼600 TIFF files) were stitched using the `VistoSeg::InformStitch`^162^ function to recreate the multichannel whole slide TIFF image. This function utilizes the X and Y coordinates embedded in each tile’s filename to accurately position the tiles. Additionally, an offset parameter was used to exclude any extraneous dead space beyond the fiducial frames of all capture areas.

#### Image splitting

Next, this stitched image was split along the y-axis into four individual capture area images in multichannel TIFF format using the VistoSeg::splitSlide_IF^162^ function.

### 4.4 Visium gene expression data processing and analysis

#### Visium raw gene expression data processing and QC

The FASTQ files of all 63 Visium SPG samples were preprocessed and aligned with their corresponding immunofluorescence images following 10x Genomics `SpaceRanger` pipeline (v2.1.0),^163^ and later contained in a `SpatialExperiment` (v1.12.0) object.^164^ To identify spots that were compromised and hence deemed low quality, we used `SpotSweeper` (v0.99.2)^165^ to remove spots with extremely low library complexity (sum_umi < 100, sum_gene < 200) as well as local spot-level outliers and compromised regions (**Figure S2**, **Figure S3**). Out of 63 Visium SPG samples, there were two regional artifacts found and annotated using `SpotSweeper` (**Figure S3**). Low-quality spots and regional artifacts were later discarded for downstream analysis. Overall, we discarded 3,001 (1.06 %) spots, with 4 to 1,077 spots per sample with a median of 14 (**Figure S4**). Empirical cumulative distributions functions (CDFs) of three QC metrics, including total number of unique molecular identifiers (UMIs), total number of unique genes detected, and proportion of mitochondria genes, were compared between two diagnostic groups (**Figure S4**). No visual difference between the CDFs of the two diagnostic groups suggests that there was no evidence of bias with respect to the diagnosis. QC metrics, spot-level descriptive statistics are visualized using `scater` (v1.34.0)^166^; spot plots are made with `escheR` (v1.3.2).^167^

#### Data-driven SpD identification

To identify SpDs in a data-driven and scalable fashion, we applied the spatially-aware clustering algorithm `PRECAST` (v1.6.4)^24^ to all 63 samples. `PRECAST` provides native data integration to account for complex batch effects, and hence is effective to predict SpDs across experimental batches and diagnostic groups. We chose 723 genes for feature selection as input to `PRECAST`, which consisted of the top 100 laminar marker genes of Sp09D in Huuki-Myers et al.,^25^ and examined multiple possible numbers of clusters (*k*=2-16) (**Figure S5**). After examining the predicted SpDs of *k*=2-16, we found that `PRECAST` SpDs of *k*=7 matched most closely with laminar organization of the classic histological layers in a spatially congruent fashion, and was used in the following downstream analysis.

#### Marker gene identification for each PRECAST SpD

To characterize the transcriptomic profiles of PRECAST SpDs, we created pseudobulked profiles based on combinations of donor and individual domains from the data-driven SpDs at *k*=7 using `spatialLIBD::registration_pseudobulk`^26^ (we used `spatialLIBD` v1.18.0 in all analyses unless noted otherwise). We first calculated the proportion of variance in gene expression explained by key variables, including PRECAST SpDs, Visium slide id, donor diagnosis status, sex, implemented using `scater::plotExplanatoryVariables`,^166^ assuming that data points in the pseudobulked dataset were independent (**Figure S7**). Then, the pseudobulk profiles at the donor and SpD levels was then used to identify maker genes across the PRECAST SpDs using two models, an enrichment test and a pairwise test, as previously described in Maynard, Collado-Torres et al.,^27^ while adjusting for donor age, sex and Visium slide ids. The enrichment test examines differences in expression between one SpD and all other SpDs, implemented in `spatialLIBD::registration_stats_enrichment`. The pairwise test examines the differences in expression between every pair of SpDs, implemented in `spatialLIBD::registration_stats_pairwise` (**Figure S8**, **Figure S9**).

#### PRECAST SpD annotation via spatial registration

To annotate the data-driven SpDs, we used `spatialLIBD::layer_stat_cor` to spatially register seven PRECAST SpDs to the previously established manual annotation of cortical structures.^27^ Specifically, the function computes the Pearson correlation of enrichment *t*-statistics from each data-driven domain of seven PRECAST SpDs and each manual annotated layer, where the top 100 genes of each manually annotated layer are considered to calculate the Pearson correlation. We then used `spatialLIBD::annotate_registered_clusters` to further assess the confidence of the annotation for each pair of PRECAST SpDs and manually annotated cortical layers. For PRECAST SpDs that were registered to multiple manually annotated cortical layers, we found that they formed hybrid-layer identities. The calculation of the confidence score is detailed in Huuki-Myers et al.^25^ and an “X” is placed on the heatmap for the SpDs we were most confident in. The spatial registration procedure was also performed to compare PRECAST SpDs between the two diagnostic groups, further validating that the PRECAST SpDs captured the same cortical structures across diagnostic groups.

### 4.5 Visium-SPG image processing

#### Segmentation for each IF channel

We utilized Visium-SPG and stained tissue sections for DAPI (nuclei), NeuN (neurons), WFA (PNN), and Claudin-5 (blood vessels) (**Methods 4.3**, **Figure S10**, **Figure 1A**). Following staining, we performed imaging and preprocessing to obtain raw multichannel images per tissue section. We employed a custom image processing pipeline to segment and identify contours in the DAPI, NeuN and Claudin-5 channels (**Figure S10**).

Using Python libraries such as `SciPy`,^168^ `NumPy`,^169^ `OpenCV`,^170^ and `scikit-image(v0.20)`,^171^ we processed images by converting them to grayscale and applying binary thresholding with Otsu’s method (implemented via `scikit-image::threshold_otsu`) to isolate the signal. Contours were detected using `OpenCV::findContours` function, filtered based on area to remove noise. Contours with an area < 50 pixels for DAPI, area < 100 pixels for NeuN and Claudin-5 regions smaller than 5,000 pixels were removed to exclude artifacts. Binarized images were saved for further analysis. We extracted intensity histograms (x-axis - pixel intensity, y-axis - density) (**Figure S11B**) for all samples and noticed that the WFA signal was predominantly located in the tail of the intensity histogram. Consequently, we applied the T-point thresholding method for isolating PNN microenvironments (**Figure S11C**).^172,173^ We developed a custom function (https://github.com/LieberInstitute/spatialDLPFC_SCZ/blob/main/code/image_processing/01_triangle_threshold_right_tail.m) in MATLAB to implement this method, specifically designed to exclude the initial peak at 0 in the intensity histogram corresponding to the dark background.^174^ The WFA signal, situated in the right tail of the histogram, was isolated by drawing a straight line from the histogram’s maximum to its minimum on the right side (red line, **Figure S11B**). The threshold for extracting PNN microenvironments was determined at the x-axis value where the distance between this line (connecting the maximum and minimum of the histogram) and the histogram was maximal (green line in **Figure S11B**). Subsequently, to separate the WFA signal from the background tissue in the extracted PNN microenvironments (**Figure S11C**), adaptive thresholding was applied. Further refinement involved masking the signal with autofluorescence segmentations (obtained using simple Ostu’s thresholding) to eliminate regions corresponding to meninges and blood vessels (**Figure S11D**). Finally, a size threshold of 30,000 pixels or greater was imposed on the segmentations to remove spots associated with fluorophore aggregates, thus refining the identification of genuine WFA signals (**Figure S11E**).

#### Quantification and spot classification

Segmentations from all channels were combined into a single .mat file per sample and, along with the raw image (**Figure S10A**), were provided to `VistoSeg::CountNuclei`^162^ function to extract spot-level fluorescent metrics across all channels. Metrics for each channel included number of segmented regions per SPG spot (if any pixel of the segmented region lies inside the SPG spot), number of segmented regions per SPG spot (if the centroid of the segmented region lies inside the SPG spot), proportion of spot pixels covered by segmented signal, and mean intensity of all segmented pixels per spot. We used the proportion spot covered with segmented pixels in each channel to classify spots into SPG vs. non-SPG, using the thresholds PDAPI < 5%, 30% > PNeuN > 5%, PWFA > 5%, 20% > PClaudin > 5%, (**Figure S10B**). Given the multimodal distribution, all spots with intensities above the background level of 0 were included for all IF channels. Spots covered with less than 5% DAPI signal were classified as neuropil, spots covered with 5-30% NeuN signal were classified as neuronal, spots covered with more than 5% WFA signal were classified as PNN and spots covered with 5-20% Claudin-5 signal were classified as vasculature (**Figure S10**). Notably, when visualizing the proportion of spots from each SpD across microenvironments, we observed similar distributions in both SCZ and NTC samples, as expected, supporting the robustness of our classification scheme (**Figure S12**). We validated spot classification in the NTC samples to avoid potential confounding effects of diagnosis using a two-pronged approach.

#### Validation of spot classification using marker genes respective to SPG-defined microenvironments

We performed pseudobulk DE analysis comparing SPG and non-SPG spots for each IF channel within NTC samples (n=31 donors) (**Supplementary Table 3**). Pseudobulking using `spatialLIBD::registration_pseudobulk` (v1.16.2) was conducted by aggregating counts per sample and spot classification, resulting in 62 data points per channel. As expected, DEGs included *CAMK2A* in neuropil spots (**Figure S13A**), *SNAP25* in neuronal spots (**Figure S13B**), *PVALB* enriched in PNN spots (**Figure S13C**), and *CLDN5* in vasculature spots (**Figure S13D**). These findings corroborate the accuracy of our classification methodology.

#### Validation of spot classification by comparison with gene expression of reference marker gene list

To further validate our image-based spot classification, we compared our results with gene expression patterns of marker genes corresponding to neuropil, neuronal, PNNs and vasculature (**Figure S14**), as defined in publicly available reference datasets.^25,30,39,40^ For each channel, spots were classified as SPG or non-SPG based on the summed expression of marker genes, using empirically defined thresholds: > 1,000 counts to determine neuropil spots (**Figure S14A**), > 250 counts to determine neuronal spots (**Figure S14B**), > 5,000 counts to determine PNN spots (**Figure S14C**), and > 2,500 counts to determine vasculature spots (**Figure S14D**). These gene expression-based classifications were visually compared to those derived from image-based metrics. We observed strong concordance between the image- and gene-based classifications across all IF channels, with image-defined SPG spots exhibiting higher expression of their corresponding marker gene sets.

Furthermore, when analyzing mean summed marker gene expression per SpD, we found that domains with a higher proportion of image-based SPG spots also exhibited higher marker gene expression (**Figure S15**). We observed enrichment of neuropil spots in SpD06-L2/3 through SpD01-WMtz (**Figure S15A**), neuronal spots in SpD06-L2/3 through SpD01-WMtz (**Figure S15B**), PNN spots in SpD02-L3/4 and SpD05-L5 (**Figure S15C**), and vasculature, showed substantial proportions in SpD02-L3/4 through SpD04-WM (**Figure S15D**). However, some domains, such as SpD07-L1/M for neuropil and vasculature, showed meaningful proportions of SPG spots but lower marker gene expression (**Figure S15A**, **Figure S15D**). Collectively, these data support the validity of our approach for annotating SPG spot types based on their cellular characteristics while demonstrating minimal differences in density across SpDs between NTC and SCZ samples. Together, these results establish our ability to perform differential gene expression testing within these cellular environments across diagnosis.

### 4.6 DE analysis between SCZ and NTC donors

#### Identification of layer-adjusted, layer-restricted, and layer-specific SCZ-DEGs across diagnosis

With the donor-SpD pseudobulked data described above, we investigated SCZ-associated changes in gene expression using `limma` (v3.62.2).^175^ To identify SCZ-DEGs, we employed two linear regression models, the *layer-adjusted* model and the *layer-restricted* model, to quantify the overall and laminar-specific dysregulation, respectively (**Figure S16**). In the layer-adjusted model, we included diagnosis status as the variable of interest and PRECAST SpDs as one of the covariates, accounting for the excess variation attributed to differences across layers. Other confounders such as age, sex, and slide id were adjusted in the model. The layer-adjusted analysis was implemented using `spatialLIBD::registration_stats_enrichment`.^26^ In comparison, the *layer-restricted* model included an additional statistical interaction term between diagnosis status and PRECAST SpDs, capturing the laminar-specific effect. For each PRECAST SpD, a contrast matrix was constructed and tested to quantify the DE, implemented using `limma::topTable`. As a result, we detected 944 SCZ-DEGs (927 unique) when controlling FDR < 0.10, or 6,978 SCZ-DEGs (5,183 unique) at nominal *p* < 0.05 (**Supplementary Table 5**). By excluding layer-restricted SCZ-DEGs detected in more than one SpD, we identified a total of 4,073 genes that are uniquely significant for each of the PRECAST SpDs, namely *layer-specific* SCZ-DEGs (**Figure S16**).

#### Identifying microenvironment-restricted SCZ-DEGs across SPG-defined microenvironments

To identify SCZ-associated transcriptional alterations within our SPG-defined microenvironments for neuropil, PNNs, neuronal, and vasculature (**Section 2.1**, **Methods 4.5**), we conducted DE tests within each of these four microenvironments following the layer-adjusted DE model (**Methods 4.6**). For each of the SPG-defined microenvironments, we constructed a new donor-SpD pseudobulked dataset by subsetting to include SPG spots only. Non-SPG spots were excluded prior to the donor-SpD pseudobulking procedure. To assure statistical validity, we excluded donor-SpD pseudobulked data points, i.e., SpDs that were specific to each donor or that contained less than 10 SPG spots. For neuropil, PNN, and neuronal microenvironments, we restricted the analysis to gray-matter-associated SpDs, including SpD07-L1/M, SpD06-L2/3, SpD02-L3/4, SpD05-L5 and SpD03-L6 in the layer-adjusted DE model. For the vasculature microenvironments, all seven SpDs were included in the pseudobulked data. In the layer-adjusted DE model, we used donor diagnosis as the variable of interest while adjusting sex, age, and SpD as covariates. As SPG spot classifications were channel-independent, each comparison was made separately.

### 4.7 Enrichment analysis of SCZ-DEGs

#### Gene ontology analyses of layer-adjusted and microenvironment-restricted SCZ-DEGs

GSEA was performed using `clusterProfiler::gseGO` (v4.14.6)^176^ to evaluate enrichment of GO terms for biological processes (BPs) among layer-adjusted SCZ-DEGs. Genes were ranked based on their *t*-statistics in descending order, and enrichment scores were computed against the GO-BP gene sets obtained from `org.Hs.eg.db` (v3.20.0).^177^ Significantly enriched GO-BP terms were selected using an adjusted *p*-value threshold of 0.05. Treemaps clustered similar terms based on semantic similarity and were used to visualize the GO terms using `rrvgo::treemapPlot` (v2.4-4) (**Figure S17**).^178^

GO over-representation analysis (ORA) was performed for microenvironment-restricted SCZ-DEGs identified in the neuronal and vasculature microenvironments. ORA was conducted to identify enrichment of GO-BP terms, using a background gene set specific to each microenvironment. Microenvironment-restricted SCZ-DEGs were defined at nominal *p* < 0.05, and enriched GO-BP terms were selected at FDR ≤ 0.05. Treemaps displayed the top 50 terms after semantic-similarity reduction (threshold=0.7), and an additional neuron-focused treemap was generated by keyword filtering neuron-related terms (**Figure 6C**, **Figure S34**, Figure S37).

`SynGO` (version 1.2; 20231201; https://www.syngoportal.org) was used to annotate synaptic location (Cellular Component, CC) and function (BP) for both the layer-adjusted SCZ-DEGs (FDR < 0.10; **Figure 2C**) and the microenvironment-restricted SCZ-DEGs for neuropil and PNNs (nominal *p* < 0.05; **Figure 6C**). Unless noted otherwise, analysis used matched, analysis-specific background gene sets. Annotated genes and enriched terms (FDR < 0.01; ≥ 3 matching input genes) were reported with representative gene counts or enrichment statistics, as appropriate. Evidence filters were set to the inclusive default (“no filters; include all annotations”).

#### Enrichment of SCZ-DEGs among cell type, SpD, and SPG spot type marker genes

To localize the cell-type and SpD contributions among the layer-adjusted and layer-restricted SCZ-DEGs, we conducted a one-sided Fisher’s exact test, implemented with `spatialLIBD::gene_set_enrichment` (v1.18.0),^26^ to examine enrichment of gene sets. The marker genes were compiled respectively from (i) cell-type markers previously reported in Huuki-Myers et al.,^25^ (ii) PRECAST SpD markers detected using the layer DE model for marker gene identification (**Methods 4.4**; Marker gene identification for each PRECAST SpD), and (iii) SPG markers identified via pseudobulk DE analysis comparing SPG and non-SPG spots (**Methods 4.5**; Validation of spot classification using marker genes respective to SPG-defined microenvironments). Only those genes with log_2_ fold-change (logFC) > 0, and FDR-adjusted *p* < 0.05 were included in the marker gene sets. The selected marker gene sets were compared against the up- and down-regulated gene sets of the layer-adjusted and layer-restricted SCZ-DEGs respectively.

#### Enrichment of layer-restricted SCZ-DEGs among reference cell-type-specific SCZ-DEGs

To cross-reference and spatially localize the cell-type-specific SCZ-DEGs, we used `spatialLIBD::gene_set_enrichment` to examine the enrichment of layer-restricted SCZ-DEGs among the reference cell-type-specific SCZ-DEGs reported in Ruzicka et al..^16^ In line with the previous reporting, we included only SCZ-DEGs whose absolute log fold-change (|logFC|) was > 0.1 and adjusted *p* < 0.05, and separated the SCZ-DEGs by up- and down-regulation. Enrichment tests were performed only between gene sets of the same directionality (i.e., up-regulated with up-regulated, down-regulated with down-regulated)

#### TF enrichment analysis

To identify candidate TFs potentially regulating the observed spatially-localized gene expression changes, particularly our layer-restricted SCZ-DEGs in SCZ, we performed TF enrichment analysis using ChIP-X Enrichment Analysis version 3 (`ChEA3`).^76^ For each SpD, we ran `ChEA3` on the sets of up- and down-regulated layer-restricted DEGs, separately. For each TF, enrichment was quantified and reported using the mean-rank approach suggested by `ChEA3`. To facilitate cross-domain comparisons, enrichment ranks were normalized by the total number of TFs tested (n=1,632 TFs) to compute a rank percentile (rank / n * 100), where lower percentile values indicate stronger enrichment. We then focused on TFs overlapping with SCZ GWAS-prioritized genes curated in Trubetskoy et al. (Supplementary Table 12, “Prioritised” sheet, n=120 genes).^13^ Normalized values were visualized as heatmaps.

#### STRING network analyses

We performed network-based analyses using the `STRING` database (version 12.0, https://cn.string-db.org/)^77^ to characterize functional and/or physical interactions separately for (i) `ChEA3`-derived candidate TFs and (ii) microenvironment-restricted SCZ-DEGs (**Figure 4B**, **Figure 6E**). For both analyses, we selected the organism as Homo Sapiens for human proteins and used active interaction sources including text mining, experimental evidence, curated databases, co-expression, genomic neighborhood, gene fusion, and co-occurrence.

For the TF analysis, we selected and combined the top 10 highly ranked TFs associated with up- and down-regulated layer-restricted SCZ-DEGs, yielding a total of 20 TFs per SpD. These were then pooled into two composite groups according to domain characteristics: non-neuronal-rich domains (SpD07-L1/M, SpD01-WMtz, SpD04-WM; total 60 TFs, with overlap across SpDs) and neuronal-rich domains (SpD06-L2/3, SpD02-L3/4, SpD05-L5, SpD03-L6; total 80 TFs, with overlap across SpDs). TF networks were analyzed using the ‘full STRING network’ option integrating both functional and physical interactions, with interaction confidence thresholds set at high (> 0.7) and additionally at medium (> 0.4) to encompass broader transcriptional dysregulation patterns. The default STRING background gene set was used for these TF analyses.

For microenvironment-restricted SCZ-DEGs, the SCZ-DEGs identified within neuropil, neuronal, PNN, and vasculature microenvironments at nominal *p* < 0.05 were analyzed as ‘physical subnetworks’ to specifically capture direct protein-protein interactions and associated signaling pathways. Matched background gene sets were applied, and interactions were filtered at a high-confidence threshold (> 0.7). Functional modules were defined using the built-in Markov Cluster (MCL) algorithm with an inflation parameter set at 1.2 and further characterized using built-in gene ontology tools available within the `STRING` web portal (**Figure S38**).

### 4.8 SCZ polygenic risk score calculation

#### QC of genotype data

The 63 donors included in this study were originally genotyped across multiple heterogeneous batches using various Illumina SNP BeadChip microarrays. To ensure consistency, genotype data processing was performed in two stages: batch-level imputation and study-level merging. First, quality control (QC) was performed separately on each original genotyping batch using `PLINK` (v1.90).^179^ Pre-imputation QC excluded variants with minor allele frequency (MAF) < 0.5% (`--maf 0.005`), variant missingness > 5% (`--geno 0.05`), and Hardy-Weinberg equilibrium (HWE) *P* < 1x10^-5^ (`--hwe 1e-5`). Each batch was then phased and imputed separately using the NHLBI TOPMed imputation service and the hg38 reference panel.^180^ Post-imputation, we only retained variants within each batch if they met an imputation quality metric R^2^ > 0.9 and a batch-level MAF > 0.05. Subsequently, the 63 specific donors were extracted from their respective post-QC imputed batches and merged into a single dataset. To generate the final analytical set, we performed another round of QC on the merged dataset using `PLINK` to remove variants with missing call rates > 5% (`--geno 0.05`), HWE *P* < 1x10^-6^ (`--hwe 1e-6`) and MAF < 5% (`--maf 0.05`). To examine the genetic ancestry of the donors, we used the `global ancestry pipeline` (https://github.com/nievergeltlab/global_ancestry) to calculate the European ancestry proportion. Among all 63 donors, 60 donors have a predicted European ancestry proportion greater than 0.9; three donors, including Br2378, Br2039, and Br5622, had a European ancestry proportion between 0.6 and 0.9. When calculating European ancestry proportion, the variant exclusion criteria included missing call rates exceeding 1% (`--geno 0.01`), HWE below 1x10^-6^ (`--hwe 1e-6`), SNP missingness exceeding 1% (`--mind 0.01`), heterozygosity, and MAF above 1% (`--maf 0.01`).

#### PRS calculation

We computed polygenic risk scores (PRS) using the standard LD-pruning and *p*-value thresholding approach implemented in PLINK2.^181^ SNPs were pruned using *p*-value-informed clumping with an LD threshold of r^2^=0.1 within a 1,000-kb window. Eighteen PRS were generated from the pruned SNP sets using the following *p*-value thresholds: 5e-08, 1e-06, 1e-07, 1e-04, 0.001, 0.01, 0.05, 0.1, 0.2, 0.3, 0.4, 0.5, 0.6, 0.7, 0.8, 0.9 and 1. Weights for PRS calculation were defined as the natural logarithm of the odds ratios estimated from the meta-analysis of the PGC3 SCZ GWAS of European ancestry samples.^13^ The PRS used for DE analyses were selected at the *p*-value threshold (0.001) that yielded the smallest association *p*-value between PRS and SCZ case-control status in our dataset.

#### PRS-associated DE test

To determine PRS-DEGs, we adapted the layer-adjusted DE model described in **Methods 4.6**. In the layer-adjusted DE model, normalized PRSs replaced donor diagnosis as the variable of interest, while the first three genetic PCs were included as covariates in addition to SpD, age, sex, and slide id. FDR-adjusted *p*-values were calculated to control the FDR. GSEA was performed on the PRS-DEGs using their *t*-statistics and visualized as a treemap (**Figure S22**).

### 4.9 Spot-level polygenic risk enrichment via scDRS

Single-cell disease relevance score (`scDRS`, v1.0.4) framework^82^ was used to quantify SCZ polygenic risk enrichment across SpDs. First, MAGMA v1.10^83^ gene-based analysis was performed on the SCZ GWAS summary statistics^13^ using 1KGP3 as the LD reference panel, generating per-gene association statistics. This resulted in a gene list with MAGMA *p*-values and was further used as input to the `scdrs munge-gs` command to construct the SCZ-associated gene set filtered at FDR < 0.05. For the expression data, the `SpatialExperiment` object was manually converted to an `AnnData` .h5ad file containing log-normalized counts. A covariate file was generated including sequencing depth, detected genes, mitochondrial content, sex, age, and RIN, to control for technical and biological confounders. The `scdrs compute-score` command was run with the argument `--n-ctrl 1000`, generating 1,000 sets of matched control genes via Monte Carlo sampling (with mean/variance matching) to compute normalized scDRS scores for each spot. Finally, downstream analysis with the `scdrs perform-downstream` command was performed, specifying SpD labels (spd_label) as the grouping variable and enabling gene-level analyses.

### 4.10 Xenium data generation

#### Xenium custom panel design

A custom panel targeting 300 genes was designed to characterize spatial gene expression patterns at enhanced cellular resolution using the 10x Genomics Xenium platform (**Supplementary Table 8**). The 300-gene panel consisted of four gene categories (**Figure 5Aiii**): (1) 168 layer-adjusted DEGs, (2) 85 cortical layer marker genes based on findings from Huuki-Myers et al.,^25^ (3) 18 SCZ-associated risk genes reported from prior literature and GWAS (references included in **Supplementary Table 8**), and (4) 35 broad brain cell-type gene markers, including excitatory neurons, inhibitory neurons, astrocytes, oligodendrocytes, microglia, endothelial cells, and immune cells. Of the 172 SCZ-DEGs, only 168 were included as 4 genes were recommended for exclusion by the manufacturer, including 3 long non-coding RNAs (*NIPBL-DT*, *LINC01736*, *LINC01535*) and *SNHG25.* With the exclusion of 6 overlapping genes across multiple categories, the final panel consisted of 300 unique genes, which was assessed and reviewed via the Xenium Panel Designer (10x Genomics).

#### Xenium slide preparation

Xenium tissue slides were prepared and run on the Xenium analyzer (10x Genomics, Cat No. PN-1000481) in groups of two using the 10x Genomics (Pleasanton CA) v1 workflow. Slides were processed in pairs (6 sections per experimental run), with each pair consisting of one slide containing 2 NTC and 1 SCZ tissue sections, and the other slide containing 2 SCZ and 1 NTC tissue sections, balancing the number of NTC and SCZ samples (3 each per run) and minimizing batch effects across the 4 experimental runs. Procedures related to cryosectioning and slide storage were followed according to the demonstrated protocol (10x Genomics, CG000579 rev C). Briefly, tissue blocks were acclimated to -16°C on a CM3050S Cryostat (Leica Biosystems) and Xenium slides were placed in the cryostat. Sections were mounted within the sample positioning area (10.45 mm x 22.45 mm) and slides were stored in a slide mailer at -80°C for 3-4 days before downstream fixation and permeabilization steps (10x Genomics, CG000581 Rev C). Briefly, slides were allowed to incubate at 37°C for 1 minute and then were fixed for 30 minutes in 40 mL of 3.7% formaldehyde (ThermoFisher Scientific), washed in 40 mL 1xPBS (ThermoFisher Scientific) and then permeabilized in 1% SDS solution (Millipore Sigma) diluted in 0.2-mm filtered nuclease-free water (Integrated DNA Technologies) for 2 minutes. Following removal of the SDS solution, slides were washed 3x in 1xPBS and then immersed in 40 mL chilled 70% methanol (Millipore Sigma, Cat No. 34860) diluted in 0.2-mm filtered nuclease-free water. After 60 minutes, slides were removed and rinsed 2x in 1xPBS (40 mL/wash) and placed in the Xenium Cassettes. We completed probe hybridization, ligation, and amplification steps according to manufacturer’s instructions outlined in CG000582 Rev E. In brief, our Xenium Standalone Custom 300-member gene panel (10x Genomics, Cat No. PN-1000563) was resuspended in 700 µL TE Buffer (ThermoFisher Scientific), vortexed, and maintained at room temperature. Probes were diluted in Xenium Probe Hybridization Buffer and applied to Xenium slides for overnight (16-24 hours) at 50°C. The next day, hybridization reactions were washed 2x in 0.05% 1xPBS-Tween-20 (ThermoFisher Scientific, Cat No. 28320) and incubated in 500 mL Xenium Post Hybridization Wash Buffer at 37°C for 30 minutes. After 3 washes of 0.05% 1xPBS-Tween-20, probes were ligated at 37°C for 2 hours in a reaction consisting of 87.5% Xenium Ligation Buffer, 2.5% Xenium Ligation Enzyme A and 10% Xenium Ligation Enzyme B. After 3 washes of 0.05% 1xPBS-Tween-20, rolling circle products were amplified in a reaction consisting of 90% Xenium Amplification Mix and 10% Xenium Amplification Enzyme for 2 hours at 30°C. After amplification was completed, slides were washed 2x in TE buffer, 3x in 1xPBS (1 mL/wash) and quenched for autofluorescence with 1% Reducing Agent B (10x Genomics) diluted in 1xPBS for 10 minutes at room temperature. Following 3 rinses of 100% Ethanol (Millipore Sigma), slides were dehydrated for 5 minutes at 37°C and then rehydrated in 1xPBS followed by a 2-minute incubation of 0.05% 1xPBS-Tween-20. Nuclei were then stained in Nuclei Staining Buffer (10x Genomics)and washed 3x in 0.05% 1xPBS-Tween-20. Slides were then stored overnight in the dark at 4°C and imaged the following day.

#### Xenium analyzer

Imaging of all slides took place on the Xenium Analyzer (10x Genomics, Cat No. PN-1000569) located at the Johns Hopkins Sidney Kimmel Comprehensive Center (SKCCC). Reagents and procedures needed to run the Xenium v1 workflow were followed according to the manufacturer’s instructions (10x Genomics, CG000584 Rev F). The most up-to-date software version (v3.1.0.0) and onboard analysis versions (v.3.1.0.4) were selected for all instrument runs. All runs passed a ‘readiness test’ confirming that the system was working optimally and that the instrument was ready for use. Reagents bottles necessary for imaging were prepared accordingly: Bottle 1 – Milli-Q ultrapure water (ThermoFisher Scientific, Cat No. 50131948), Bottle 2 – 0.05% 1xPBS Tween-20, Bottle 3 – Milli-Q ultrapure water and Bottle 4-0.1% Tween-20, 50mM KCL (Teknova, Cat No. P0330), 50% DMSO (Millipore Sigma, Cat No. 41639) diluted in nuclease-free water (ThermoFisher Scientific, Cat No. AM9932). Xenium reagent bottles, Xenium cassettes, an Object Wetting Consumable, pipette tip racks (10x Genomics, Cat No. PN-1000461) and decoding reagent plates (10x Genomics, Cat No. PN-1000461) were loaded according to details provided by the manufacturer. Following successful loading of the instrument, panel ROI selection was performed and the run was initiated. Runs took on average 48 hours to complete.

### 4.11 Xenium data analysis

#### Data QC and normalization

Data QC was performed on the Xenium data to remove low quality cells from downstream analyses. Within each sample, low quality cells were identified based on percentage negative control features, number of detected genes, and total counts observed in each cell. We used the negative control probes, negative control codewords, and unassigned codewords from the Xenium panel. We removed all cells with expression of any of the above three features in the 99th percentile. Cells that were outliers in terms of number of genes detected (5 median absolute deviations, MADs, below the median) or in terms of total counts (5 MADs above or below the median) were identified using `scater::isOutlier` and removed. The raw counts from the retained cells were then library size-normalized and log_2_-transformed (log-normalized) using `scuttle::logNormCounts` (v1.12.0).^166^

#### SpD clustering and annotation

To identify SpDs in the Xenium data, label transfer was performed using the `spaTransfer::transfer_labels` (v0.0.0.9) function.^92^ The Visium dataset described above was used as the reference dataset and the annotated PRECAST SpD7 labels (at *k*=7) were transferred into the Xenium dataset. Briefly, `spaTransfer` computes non-negative matrix factorization (NMF) on the log-normalized counts matrix of the Visium reference dataset. This results in a loadings matrix and a factors matrix, similar to PCA. spaTransfer then uses the loadings matrix to project the log-normalized counts matrix of the target Xenium dataset and learn the Xenium factors matrix through non-negative least squares. The reference factors are used to fit a multinomial model with the formula PRECAST SpD7 labels ∼ NMF factors. The trained multinomial model is then given the Xenium factors and asked to predict SpD labels for each cell in the Xenim dataset. Finally, spatially-aware neighbourhood smoothing similar to `Banksy::smoothLabels` (v0.99.0)^93^ is used to increase spatial contiguity of predicted domains. For our purposes, we performed NMF with 50 factors and performed spatial smoothing with each cell’s 10 nearest neighbours based on physical distance. We then validated the transferred SpDs using spatial registration against manually annotated human cortical layers from Maynard, Collado-Torres et al.,^27^ similar to the approach for the Visium dataset described above.

#### Cell-type clustering and annotation

Clustering was performed with the `Banksy` package^93^ in order to identify spatially-aware cell types in the Xenium data. All 24 samples were clustered jointly following the BANKSY multi-sample tutorial. To create the BANKSY neighbourhood gene expression matrix, we set the `k_geom` parameter to 10 and `compute_agf` to be `FALSE`. The `lambda` parameter which controls the weighting on the spatial information versus the gene expression information was set to 0.1, and the clustering resolution was set to 0.7. This yielded 18 total clusters which we then annotated with cell-type labels.

The annotations were based on expression of marker genes which were included in the Xenium panel. The marker genes included in the Xenium panel spanned 7 major cell types: astrocytes, endothelial cells, inhibitory neurons, excitatory neurons, microglia, microglia/immune cells, and oligodendrocytes. The inhibitory and excitatory marker genes also allowed us to separate these cells into subclasses: SST and PVALB inhibitory neurons, VIP and LAMP5 inhibitory neurons, L2/3 excitatory neurons, L4/5 excitatory neurons, L5 excitatory neurons, and L6 excitatory neurons. Annotation of the 18 clusters resulted in a total of 12 cell types, with two of the clusters being ambiguous, likely due to the gene panel not having the resolution to confirm the identity of these subpopulations. Investigation of QC metrics within these groups gave no indication that these clusters are driven by low quality cells, so they were retained in downstream analyses. To validate the cell-type annotation, we examined the relationship between the SpDs and cell types by computing the proportion of each cell type within the SpDs (**Figure S30**). For each cell type, proportions were similar between NTC and SCZ. Cell types were also enriched in the expected cell-level SpDs; for example, oligodendrocytes were predominantly enriched in the WM domain, and L2/3 Ex neurons were primarily enriched in the L2/3 domain.

#### Identifying case-control DEGs & functional annotation

##### DE across cortical layers

Similar to the Visium DE analysis, the counts matrix was first pseudobulked to the donor-domain level, using the predicted SpD annotations from `spaTransfer`. Any pseudobulk samples with fewer than 10 cells were dropped. We first performed layer-adjusted DE analysis using `limma` through `spatialLIBD::registration_stats_enrichment`. Since the pseudobulking procedure introduced “replicates” for each donor, we used `limma::duplicateCorrelation` to account for possible correlation between these replicates. We adjusted for SpDs, age, sex, and the xenium slide for each sample.

##### DE across cell types

The cell-type-adjusted DE analysis was performed similarly to the layer-adjusted DE analysis. The counts matrix was pseudobulked to the cell type-donor level, using the cell-type annotations described above. We used `limma` through the `spatialLIBD::registration_stats_enrichment` function, again accounting for correlation between replicates with `limma::duplicateCorrelation.` We adjusted for cell type, age, sex, and the Xenium slide for each sample. In order to identify expression changes occurring within specific populations, we leveraged the cell-type information to perform cell-type-stratified DE analysis. For each of the 12 cell types we annotated, we first subsetted the counts matrix to only cells from one type. We then pseudobulked these cells to the donor level, dropping samples with fewer than 10 cells. Similar to as described above, `spatialLIBD::registration_stats_enrichment` was run, this time adjusting for age, sex, and the xenium slide for each donor.

#### Differential cell-type abundance and density analysis

To assess whether the abundance of cell types differed between NTC and SCZ, we employed compositional data analysis techniques using the `cacoa` R package (v0.5.0).^98^ Briefly, the `cacoa::estimateCellLoadings` function first transforms the abundances of the 12 cell types into a 11-dimensional simplex space using the isometric log-ratio transformation. A surface that best separates the NTC and SCZ samples in this 11-dimensional simplex space is then estimated, and the contribution of each cell type to the surface’s orientation is computed. To test for the statistical significance of each cell type, an empirical background distribution is constructed by resampling cells and samples. We first ran this analysis without stratifying the data to SpDs to identify global changes in cell-type abundance. We then ran a layer-stratified version of this analysis by subsetting to cells within a SpD and testing for differences in cell-type abundance within that domain.

### 4.12 Cell-cell communication within SPG-defined microenvironments

To evaluate how SCZ-associated transcriptional changes intersect with LR signaling within SPG-defined microenvironments, we compared our microenvironment-restricted SCZ-DEGs to a curated set of candidate LR pairs in Supplementary Table 5 from Huuki-Myers et al..^25^ Specifically, we considered LR pairs that were annotated for SCZ genetic risk and further subsetted to those that were identified as mediating cell-cell communication in a dlPFC snRNA-seq dataset. We then subsetted our microenvironment-restricted SCZ-DEGs to those with nominal *p* < 0.05. For each of the four microenvironments, we assessed whether any of these genes were present as either a ligand or a receptor within the curated list of LR pairs. When a gene matched one of the LR interactors, we plotted the corresponding LR pair in a Sankey plot (**Figure 6E**), along with its interaction partner, independent of whether the partner gene was significantly differentially expressed in SCZ.dependent of whether the partner gene was significantly differentially expressed.

### 4.13 Spatial eQTL analysis and colocalization analysis

#### Characterizing spatial eQTLs with Visium data

To identify genetic variants associated with domain- and microenvironment-specific gene expression, we performed cis-eQTL mapping utilizing pseudobulked gene expression profiles. Expression data from the `SpatialExperiment` objects were aggregated at the donor level, stratifying by seven SpDs and four SPG microenvironments (neuropil, neuronal, PNN, vasculature), amounting to a total of eleven contexts in which eQTL and colocalization analyses were performed independently.

To address the inherent sparsity and mean-variance relationships in pseudobulked spatial transcriptomics, we implemented a targeted data transformation pipeline prior to latent variable extraction.^182^ For mapping, pseudobulked expression matrices underwent trimmed mean of M-values (TMM) log_2_ counts per million [log2(CPM+1)] normalization (`edgeR` v4.8.2),^183^ followed by a rank-based inverse normal transformation across donors, adapting the standard GTEx Consortium eQTL discovery workflow.^184^ To calculate latent expression principal components (`exprPCs`), TMM log_2_(CPM) values were converted to *z*-scores, and PCA was restricted to the top 5,000 highly variable genes (HVGs), ranked by their variance-to-mean ratio. The optimal number of hidden expression components was estimated using `sva::num.sv` (v3.58.0),^185^ yielding 6 to 9 `exprPCs` per context.

We conducted eQTL mapping using `tensorQTL` v1.0.10^124^ to test for associations between genetic variants and gene expression within a 1 Mb cis-window around the gene Transcription Start Site (TSS): TSS - 1 Mb, TSS + 1 Mb. Our baseline linear model adjusted for diagnosis, sex, age, slide id, the first five genotyping principal components (`snpPCs`), and the context-specific `exprPCs`. Cis-eQTL gene-level significance (FDR < 0.05) was determined empirically using `tensorQTL`’s permutation-based `map_cis` with FDR estimated by the *q*-value procedure. Conditionally independent eQTL signals within each context were subsequently identified using the `map_independent` module, restricted to genes significant at FDR < 0.05 in `map_cis`. Nominal eQTL signals were identified using `map_nominal`, which returns nominal *p*-values for variant-gene pairs. These associations were then filtered at FDR < 0.05 after Benjamini-Hochberg adjustment.

To determine if the identified eQTLs are colocalized with SCZ GWAS risk variants, we performed bayesian colocalization analysis integrating our cis-eQTL summary statistics with the latest SCZ GWAS summary statistics from the Psychiatric Genomics Consortium (PGC3, European ancestry).^13^ We used `coloc` v5.2.3,^186^ running the `coloc.abf` function on variants shared between the eQTL and GWAS datasets. Evidence for colocalization was evaluated based on the posterior probabilities of independent causal variants (PP3) versus a shared causal variant (PP4). We defined "strong colocalization" loci as those meeting the probability threshold of PP4 ≥ 0.8. We assessed robustness with a prior-sensitivity analysis with `coloc::sensitivity`, and found no differences in the strong colocalization results.^187^

#### Characterizing spatial eQTLs with Xenium data

Using the 24 Xenium donors, we next investigated the cell-type specificity of *KANSL1-AS1* expression, which was identified as an eGene gene with a strong colocalization signal (rs2532240). To do so, we pseudobulked the counts from each donor-domain-cell type combination and examined the library size-normalized and log_2_-transformed counts in each pseudobulk sample. Based on a heatmap of this data, we identified *KANSL1-AS1* to be relatively enriched in microglia, astrocytes, and oligodendrocytes located in WM.

To further investigate the cell-type specificity of the genotype-expression relationship, we compared the gene expression to the genotypes of all 24 Xenium donors at (rs2532240). Since this colocalization result was found in SpD04-WM, we subsetted the Xenium data to the cells in the WM domain and pseudobulked to the donor-cell type level. The pseudobulked data then underwent TMM and log2(CPM+1) normalization using `edgeR`. We calculated top latent `exprsPCs` (similar to above), but using all features in the Xenium data. We then fit a linear model as described for the Visium data above using the function `jaffelab::cleaningY`.^188^ Specifically, we regressed out diagnosis, sex, age, Xenium slide id, the same first five `snpPCs` as above, and Xenium `exprPCs` to get the residualized expression of *KANSL1-AS1* to compare to the genotypes of the Xenium donors. The number of `exprPCs` to include as covariates for the model were identified using `sva::num.sv`. We then compared the expression level of *KANSL1-AS1* across genotypes within each cell type. One-sided *t*-tests were performed between the C:C and C:T genotypes to identify significant changes in expression level.

### 4.14 Interactive data visualization and exploration

To assist interactive exploration of the Visium SRT data, along with immunostaining data, we provided the web-based data visualization tools, including `Samui Browser`^189^ for visualizing fluorescent images, nuclei segmentation, as well as gene expression data, and `iSEE` apps.^190^ These tools can be used to explore donor-SpD pseudobulked gene expression data generated from all spots or subsets of spots classified into four SPG-defined microenvironments (neuropil, neuronal, PNN, vasculature). Further details on accessing the Samui browser and iSEE apps can be found via https://research.libd.org/spatialDLPFC_SCZ/#interactive-websites.

## 4.15 Data and code availability

Raw and processed data are available through Gene Expression Omnibus (GEO) under accession GSE307403 (Visium) and GSE307404 (Xenium). Project data was also uploaded to the National Institute of Mental Health Data Archive (NDA) and can be found with study ID 3054 and its associated DOI 10.15154/k6s8-2839. We provide web-based applications (iSEE, Samui Browser) for exploring the current Visium SRT dlPFC datasets, as listed at https://research.libd.org/spatialDLPFC_SCZ. The code for this project is publicly available through GitHub at https://github.com/LieberInstitute/spatialDLPFC_SCZ and https://github.com/LieberInstitute/spatialDLPFC_SCZ_XENIUM. Zenodo archives have been made for both repositories.^174,191^

## 4.16 Acknowledgments, Funding, Authorship Contributions

### Acknowledgements

Portions of some figures were created with BioRender.com. We thank the LIBD neuropathology team, particularly James Tooke and Amy Deep-Soboslay, for curation of the brain samples and assistance with tissue dissections. We thank the staff and physicians at the brain donation sites, and the generosity of the brain donors and their families, without whom this work would not be possible. We thank members of the Martinowich, Maynard, Page, and Hicks labs as well as Dr. Daniel R. Weinberger for critical reading of the manuscript and feedback. We thank the Joint High Performance Computing Exchange (JHPCE) for providing computing resources for these analyses. We thank members of the Johns Hopkins Integrated Genomics Center for assistance generating the 10x Genomics Xenium data. Finally, we thank the families of Connie and Steve Lieber and Milton and Tamar Maltz for their generous support.

### Funding

This project was supported by R01MH126393 (KM), R01MH123183 (KRM/SCH), R35GM150671 (SCH), and the Lieber Institute for Brain Development.

### Author Contributions

Conceived and Designed the Study: KRM, SCH, KM

Performed Experiments and Collected Data: SHK, SVB, SEM, MRV, SO, YD, ABN, RZ, HRD, JSL, TMH

Software: BG, SCP, RAM

Formal Analysis: BG, CF, MT, AJ, AI, VB, UMK, GP, NJE, LCT, SH

Data Curation: BG, XZ, RAM

Tissue Resources: SVB, JEK, TMH

Writing – original draft: BG, SHK, CF, MT, KRM, KM

Writing – review & editing: SHK, BG, CF, MT, SVB, TMH, SCP, KRM, SCH, KM

Visualization: BG, SHK, CF, MT

Supervision: ND, SCP, KRM, SCH, KM

Funding acquisition: KRM, SCH, KM

Project Administration: KRM, SCH, KM

## Supplemental Figures

**Figure S1.**
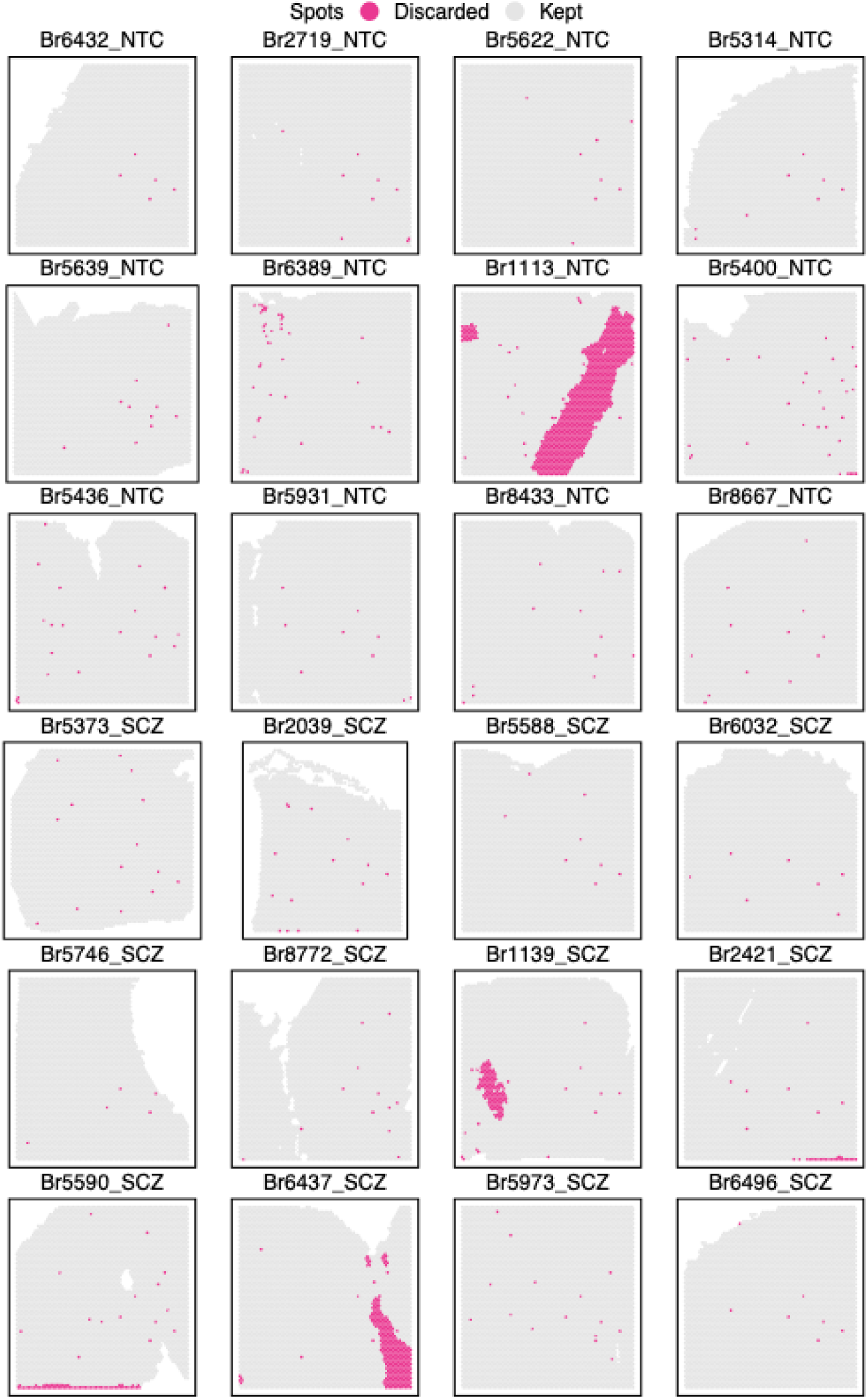
Representative overview of excluded spots. Spots with library size less than 100 and fewer than 200 unique genes detected, and tissue artifacts (**Figure S3**) were flagged for removal. Representative spot plots showing spots flagged for removal from a subset of the cohort (24/63 samples).

**Figure S2.**
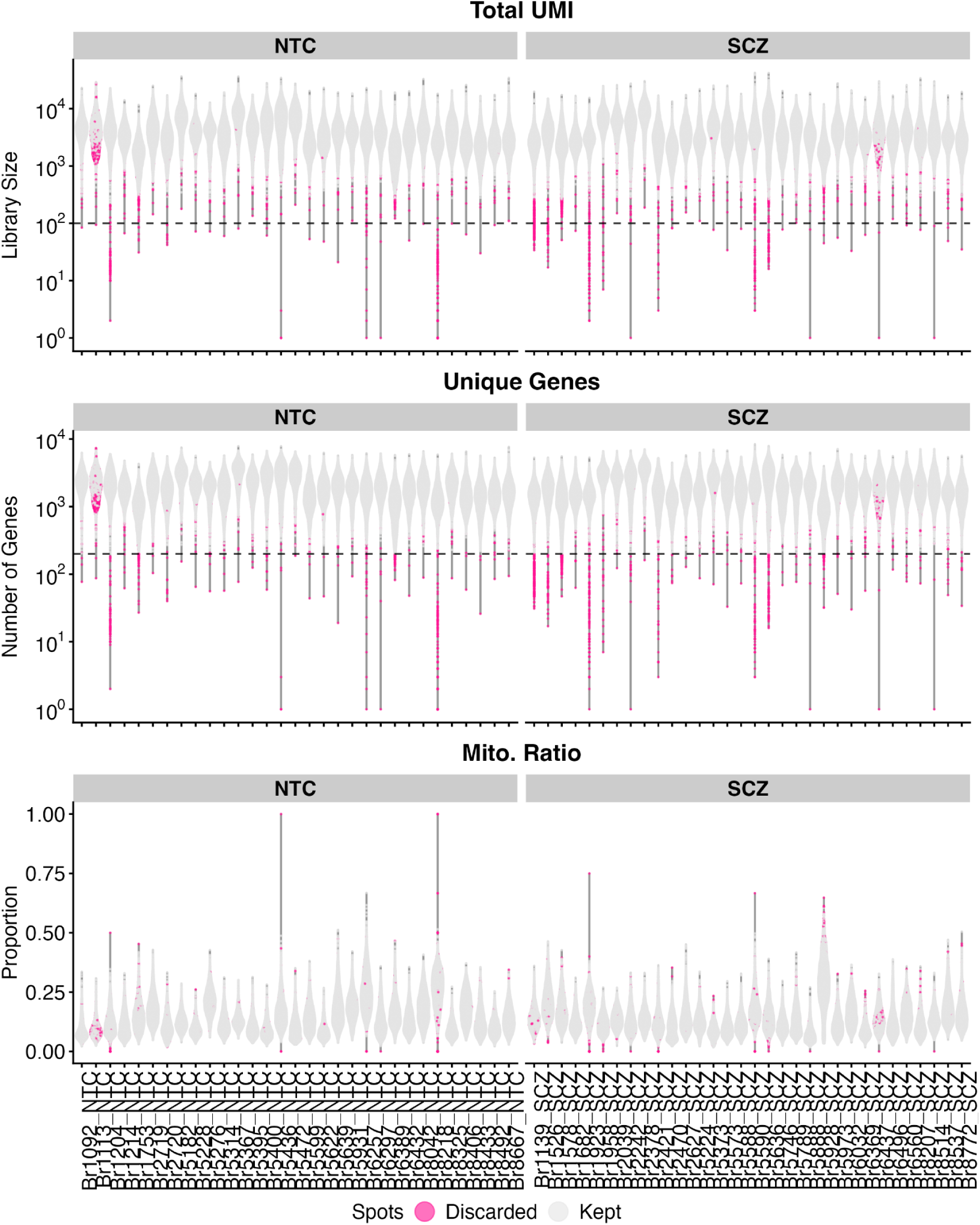
Distributions of QC metrics stratified by diagnosis. Violin plots showing spot-level distributions of three QC metrics, stratified by diagnosis (NTC and SCZ): total unique molecular identifiers (Total UMIs) also known as library size (**top**), number of detected genes (Unique Genes) (**middle**), and proportion of mitochondrial genes (Mito. Ratio) (**bottom**). Dotted lines represent thresholds used to identify low-quality spots (UMIs < 100 and genes < 200).

**Figure S3.**
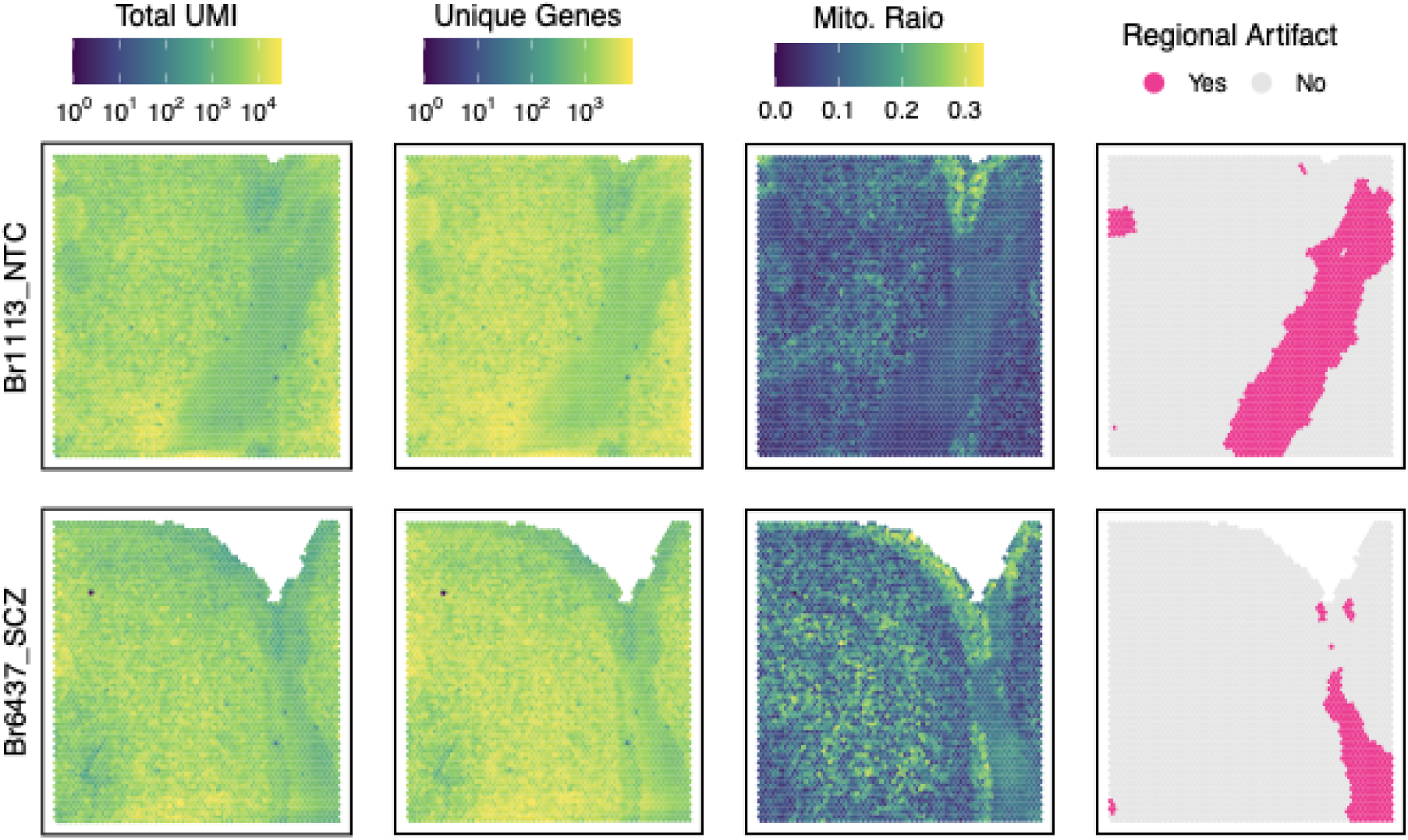
Regional artifacts identified by SpotSweeper. Examples from two samples (Br1113_NTC and Br6437_SCZ) illustrate technical artifacts introduced during experimental procedure, leading to diagonal scratches traversing across tissue and low-quality spots underlying the affected regions. SpotSweeper^165^ identified these regional artifacts and flagged them for removal.

**Figure S4.**
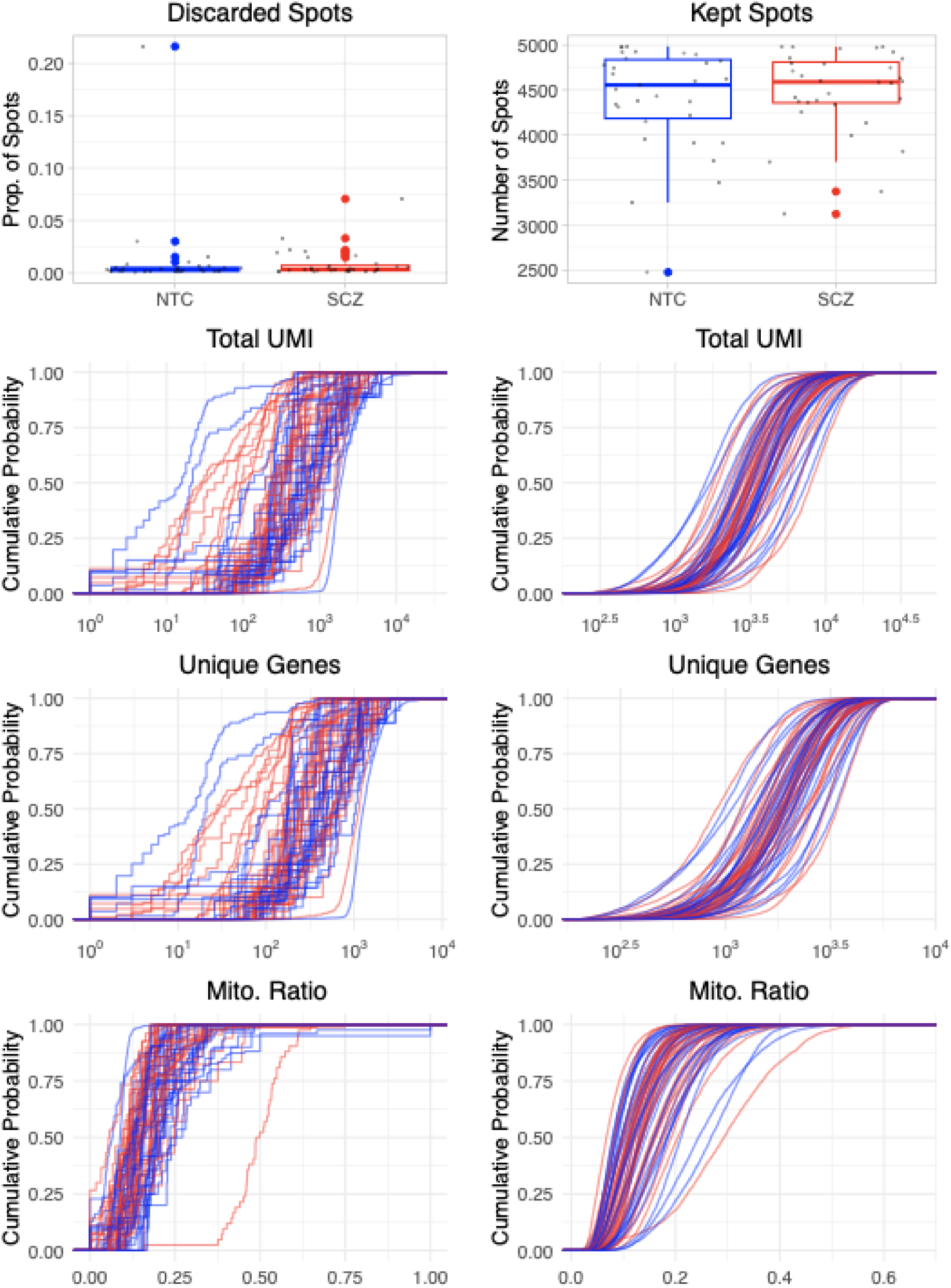
Post-QC characterization of discarded and retained spots indicates no evidence of diagnosis-related bias. (**Top left**) Proportion of spots discarded in NTC samples. (**Top right**) Number of spots retained after QC in NTC and SCZ samples. (**Bottom**) Empirical cumulative distribution functions (CDFs) of three QC metrics (total UMIs, number of unique genes, and proportion of mitochondrial genes) plotted for discarded (**left**) and retained (**right**) spots by diagnosis.

**Figure S5.**
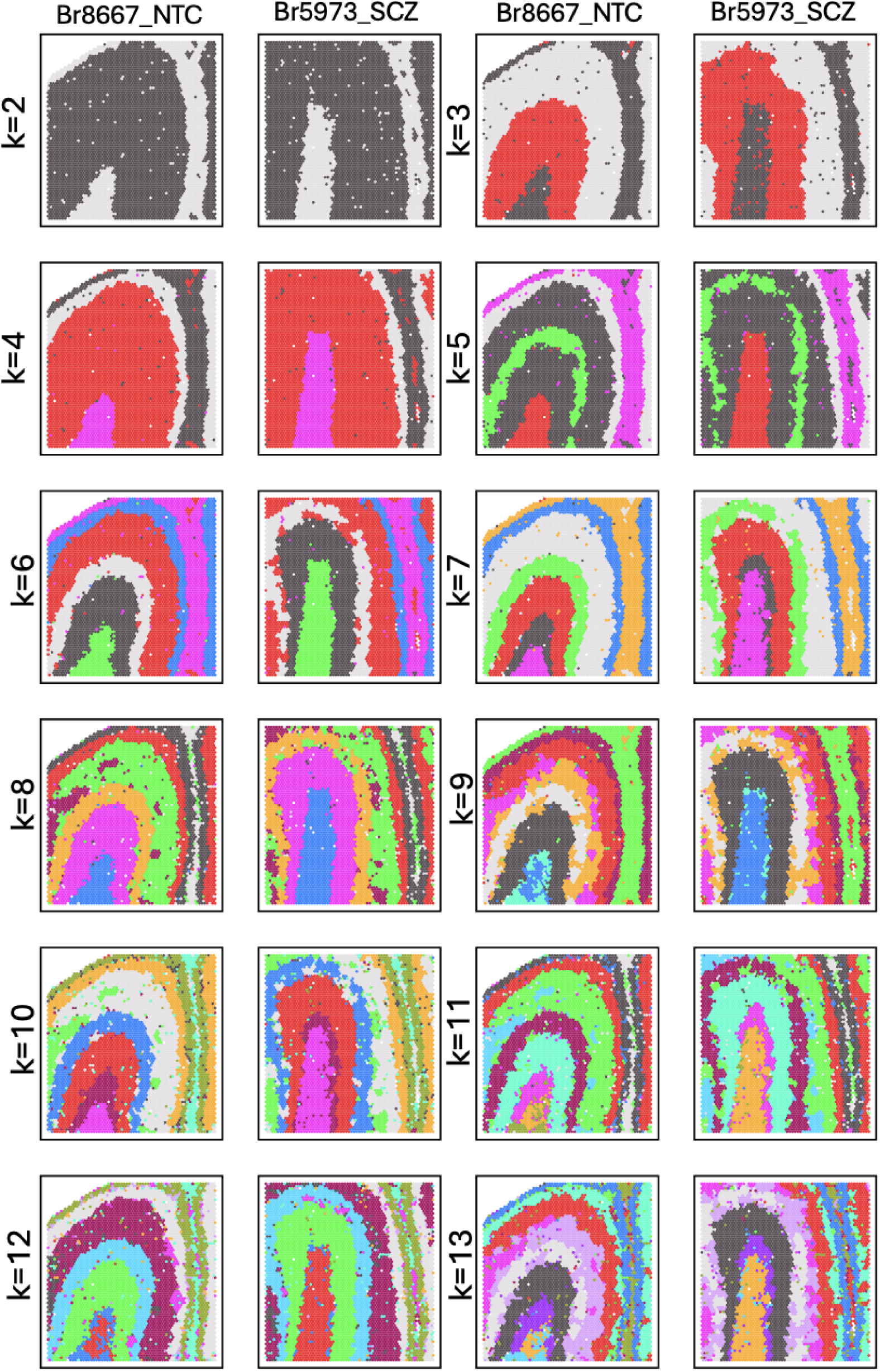
Spot plots of data-driven SpDs at different clustering resolutions. Unsupervised spatial clustering using PRECAST was performed at multiple resolutions (*k*=2-13), as shown in one representative sample from each diagnosis (Br8667 and Br5973, respectively). *k*=7 was chosen for downstream analysis because the patterning best correlated with established histological layers.

**Figure S6.**
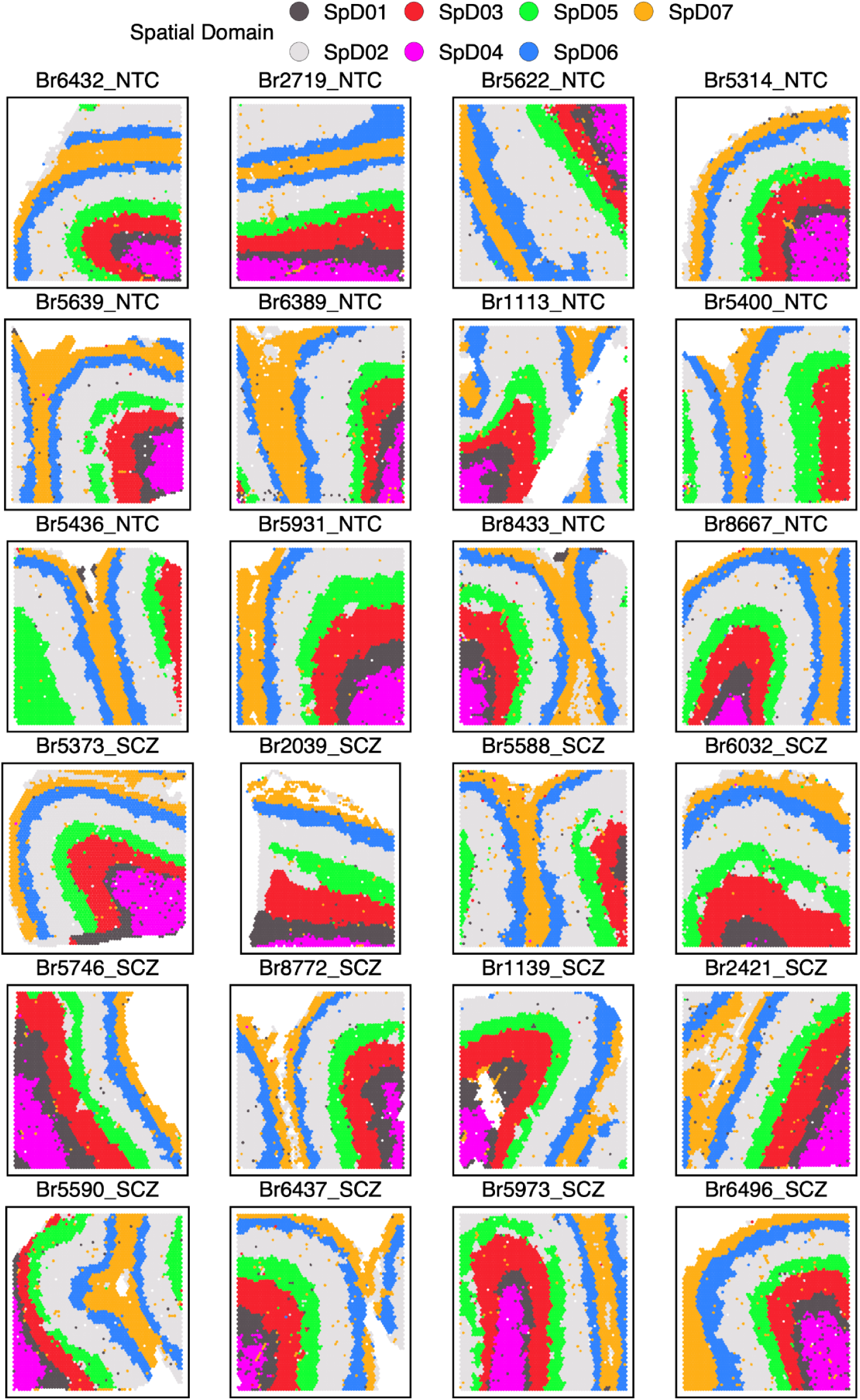
Spot plots of the *k*=7 PRECAST SpDs across a subset of the cohort. Spatial clustering patterns identified using PRECAST are shown for a subset of the cohort (24/63; 12 NTC and 12 SCZ). Colors represent the SpDs.

**Figure S7.**
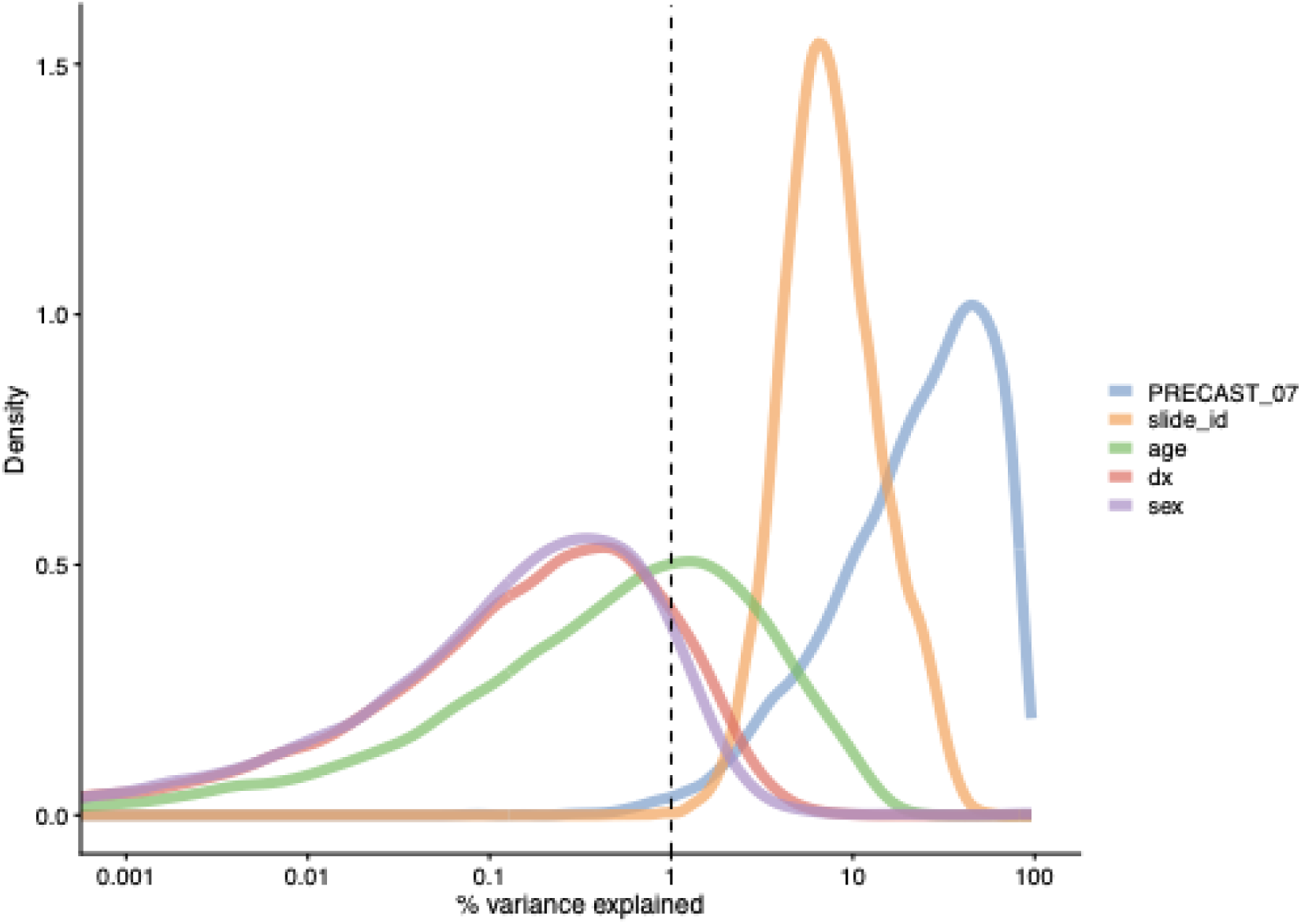
Density curves showing the gene-level variance explained by different variables in the *k*=7 SpDs. Spatial domain (SpD; PRECAST_07) accounts for the highest percentage of variance explained, whereas diagnosis contributes to a modest amount of variance, supporting the computational strategy to jointly clustering across the diagnostic groups.

**Figure S8.**
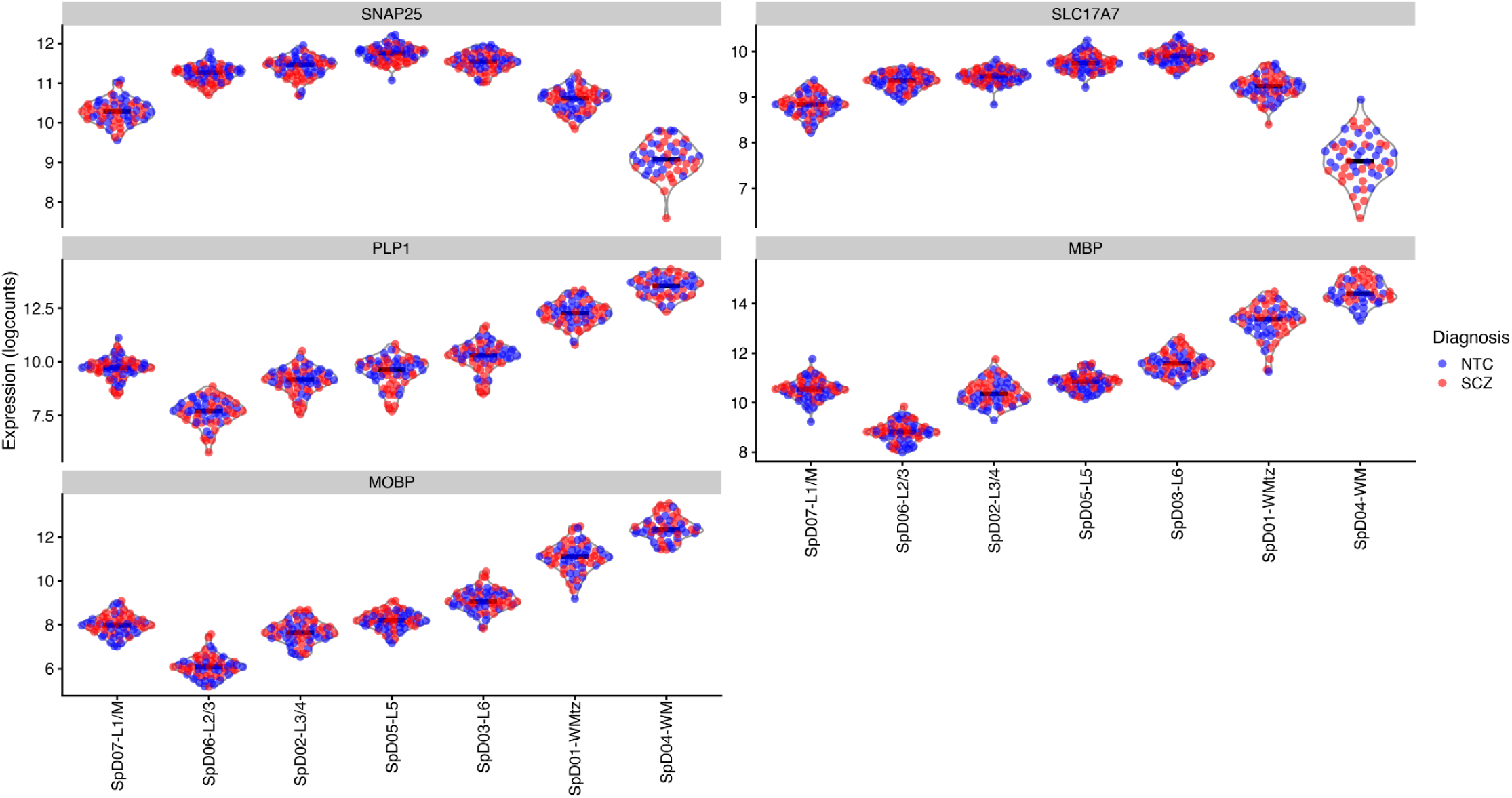
Violin plots of representative neuronal and oligodendrocyte marker gene expression across SpDs. Gene expression levels were pseudobulked within each individual brain donor for each SpD. Compared to SpD04-WM, SpD01-WMtz exhibited higher expression of neuronal marker genes, including *SNAP25* and *SLC17A7*, while showing relatively lower expression of oligodendrocyte genes, such as *PLP1*, *MBP*, and *MOBP*.

**Figure S9.**
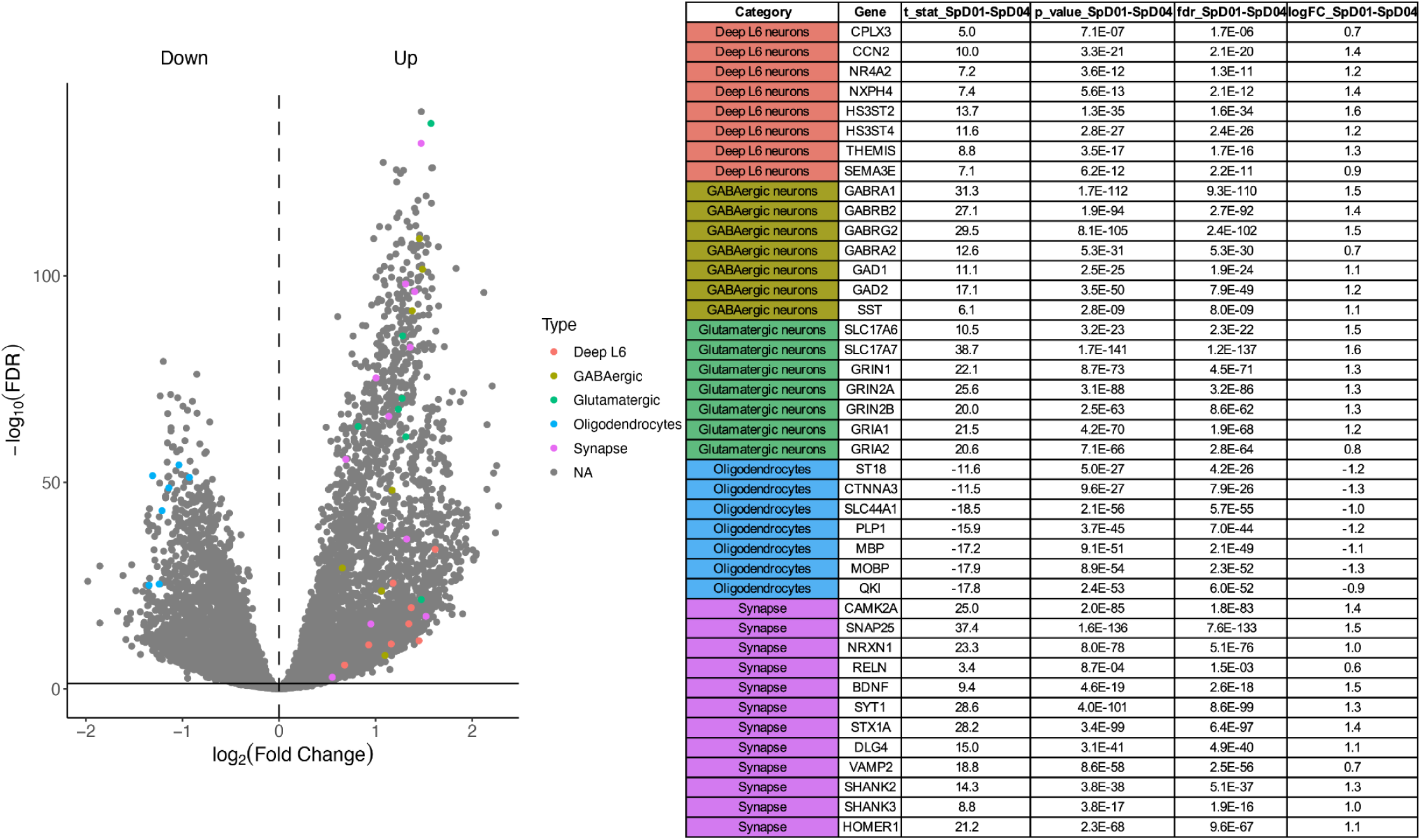
Volcano plot of DEGs between SpD01-WMtz and SpD04-WM. Representative cell-type-associated genes are visualized in a volcano plot (left) to illustrate relative enrichment of neuronal and synapse-related markers and depletion of oligodendrocyte markers in SpD01-WMtz compared to SpD04-WM. Genes without annotation are labeled as “NA” for “Not Annotated.” The horizontal line denotes the significance threshold of FDR=0.05. The corresponding genes highlighted in the volcano plot are listed in the table (right).

**Figure S10.**
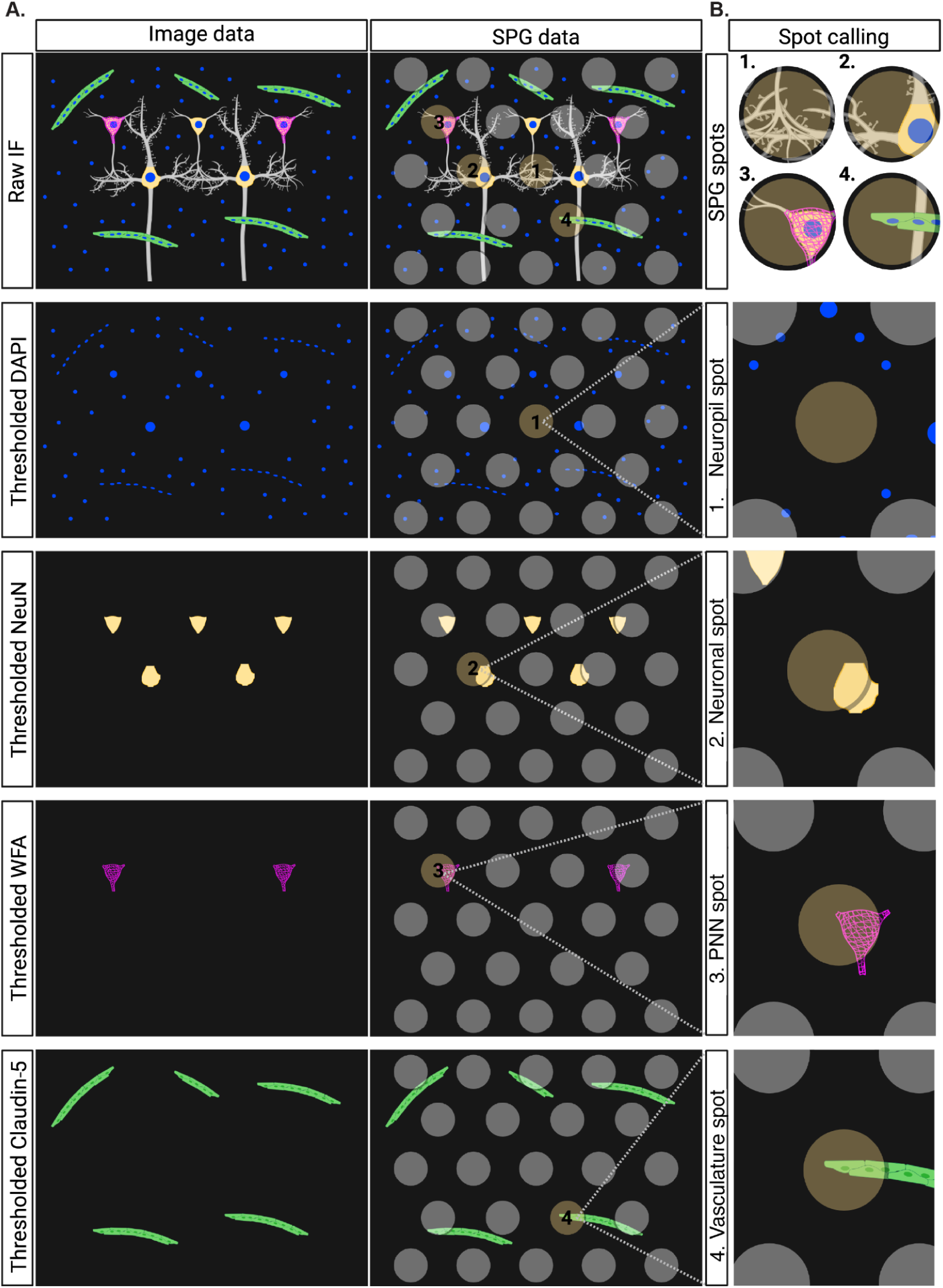
Schematic of the IF image-based strategy for classifying SPG spots into cellular microenvironments. (**A**) Image data (Column 1) displays representative raw IF images and their corresponding thresholded binary image for each channel. SPG data (Column 2) shows these raw and thresholded IF image data overlaid with the Visium spot grid. (**B**) Spot calling (Column 3) depicts how SPG spots (highlighted in brown) were classified into SPG-defined cellular microenvironments based on enrichment of IF signals: 1. Neuropil spots lacking DAPI IF signal, 2. Neuronal spots with positive NeuN IF signal, 3. PNN spot with WFA IF signal, and 4. Vasculature spots with Claudin-5 IF signal. Created in BioRender. Kwon, S. H. (2026) https://BioRender.com/e4mjp52

**Figure S11.**
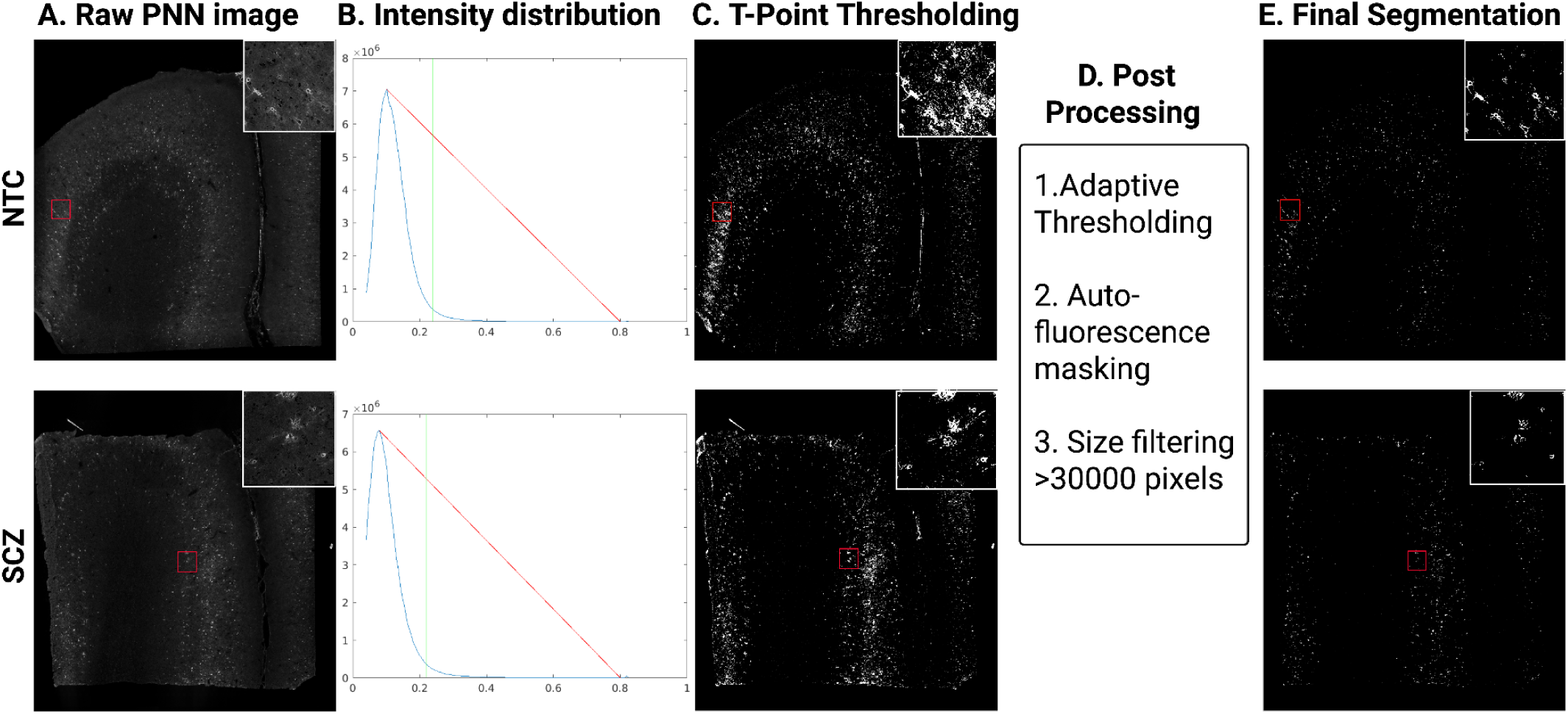
WFA segmentation workflow to identify PNN regions of interest. (**A**) Raw PNN channel images display the raw WFA IF images for representative NTC and SCZ samples (Br8667 and Br5973, respectively). (**B**) Intensity distribution shows the intensity histograms for the images presented in (**A**). The method for extracting the T-point threshold is illustrated,^172,173^ where the red line connects the maximum and minimum intensities on the right side of the histogram. A green threshold is selected at the point where the distance between the histogram and the red line is maximized. (**C**) T-point thresholding depicts the binary images resulting from T-point thresholding applied to the raw images shown in (**A**). (**D**) Post-processing describes the additional image processing applied, including adaptive thresholding (to remove tissue background noise), autofluorescence masking, and size filtering to eliminate meninges and other noises from the T-point thresholded image. (**E**) Final segmentation shows the final segmented images containing PNN ROIs after all image processing steps, which were used for PNN spot classification.

**Figure S12.**
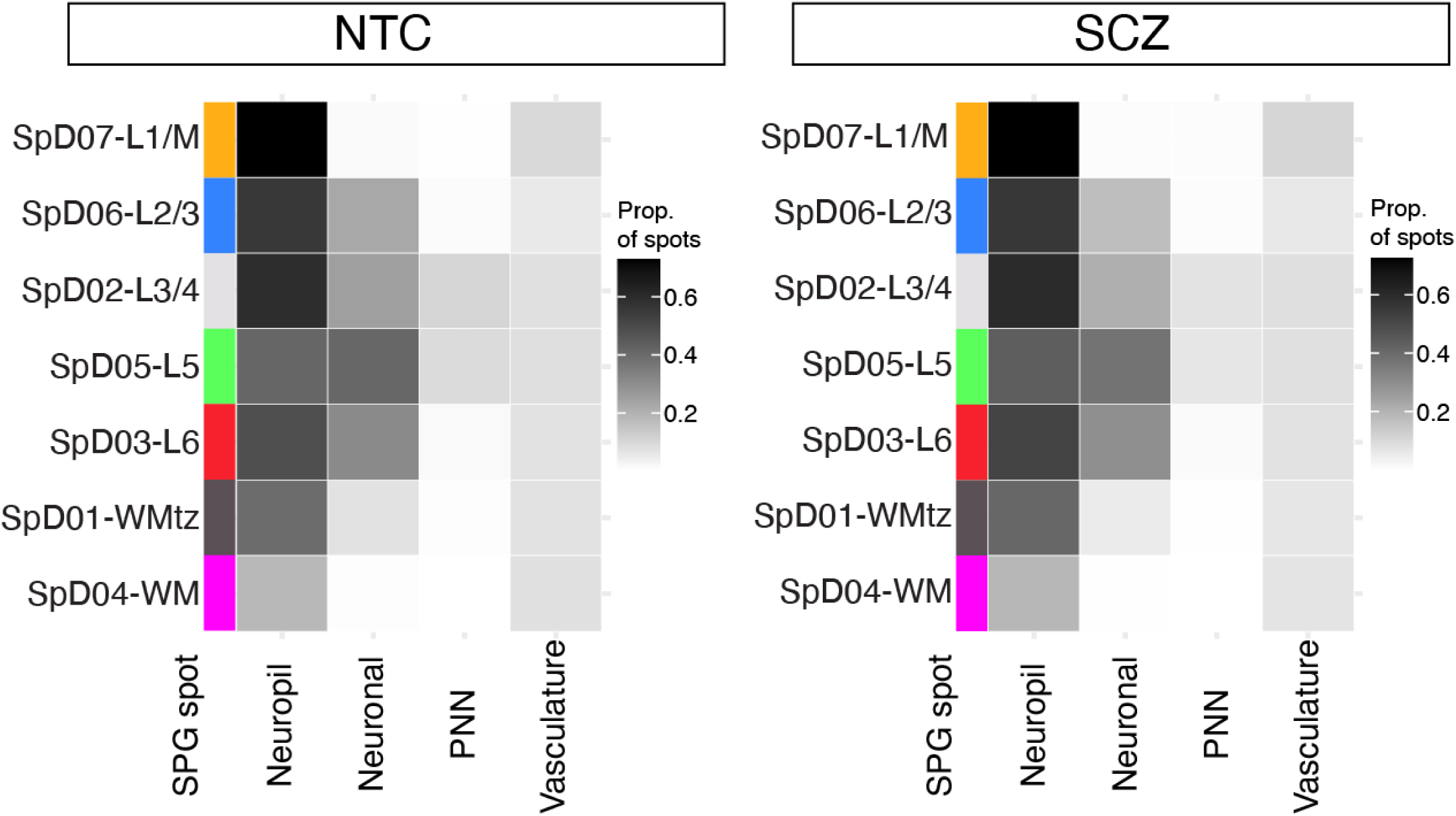
Enrichment of SPG spots per SpD. Heatmaps show, for each SpD, the proportion of SPG spots that are positive for each microenvironment category (neuropil, neuronal, PNN, and vasculature) calculated independently per category. Because assignments are non-mutually exclusive, spots can contribute to multiple categories, and proportions across rows do not sum to 1. Results are shown separately for NTC (left) and SCZ (right) samples. Overall, the distributions indicate highly similar SPG-defined microenvironment compositions across cortical domains between diagnostic groups.

**Figure S13.**
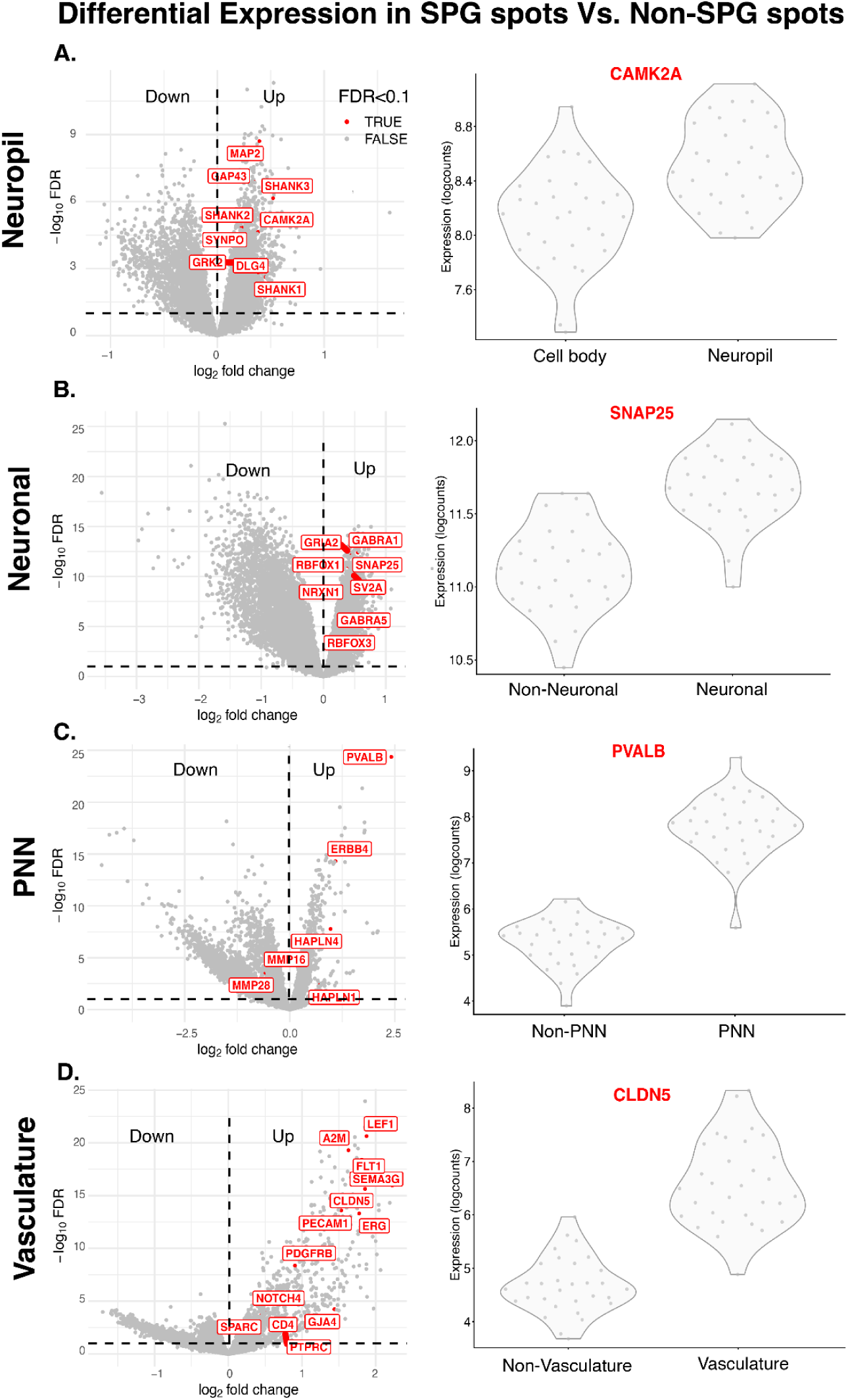
DE analysis across SPG-defined microenvironments within NTC samples. (**A**) Top DEGs between neuropil spots and cell body-enriched non-neuropil spots (cell body) highlight higher gene expression of synaptic genes such as *CAMK2A* in neuropil spots. Detailed DE results are provided in **Supplementary Table 3**, Sheet 1. (**B**) Top DEGs in neuronal spots demonstrate higher gene expression of neuron-associated genes such as *SNAP25* compared to non-neuronal spots. Detailed DE results are provided in **Supplementary Table 3**, Sheet 2. (**C**) Top DEGs in PNN spots highlight higher gene expression of PNN-associated genes such as *PVALB* compared to non-PNN spots. Detailed DE results are provided in **Supplementary Table 3**, Sheet 3. (**D**) Top DEGs in vasculature spots highlight higher gene expression of blood vessel-associated genes such as *CLDN5* compared to non-vasculature spots. Detailed DE results are provided in **Supplementary Table 3**, Sheet 4.

**Figure S14.**
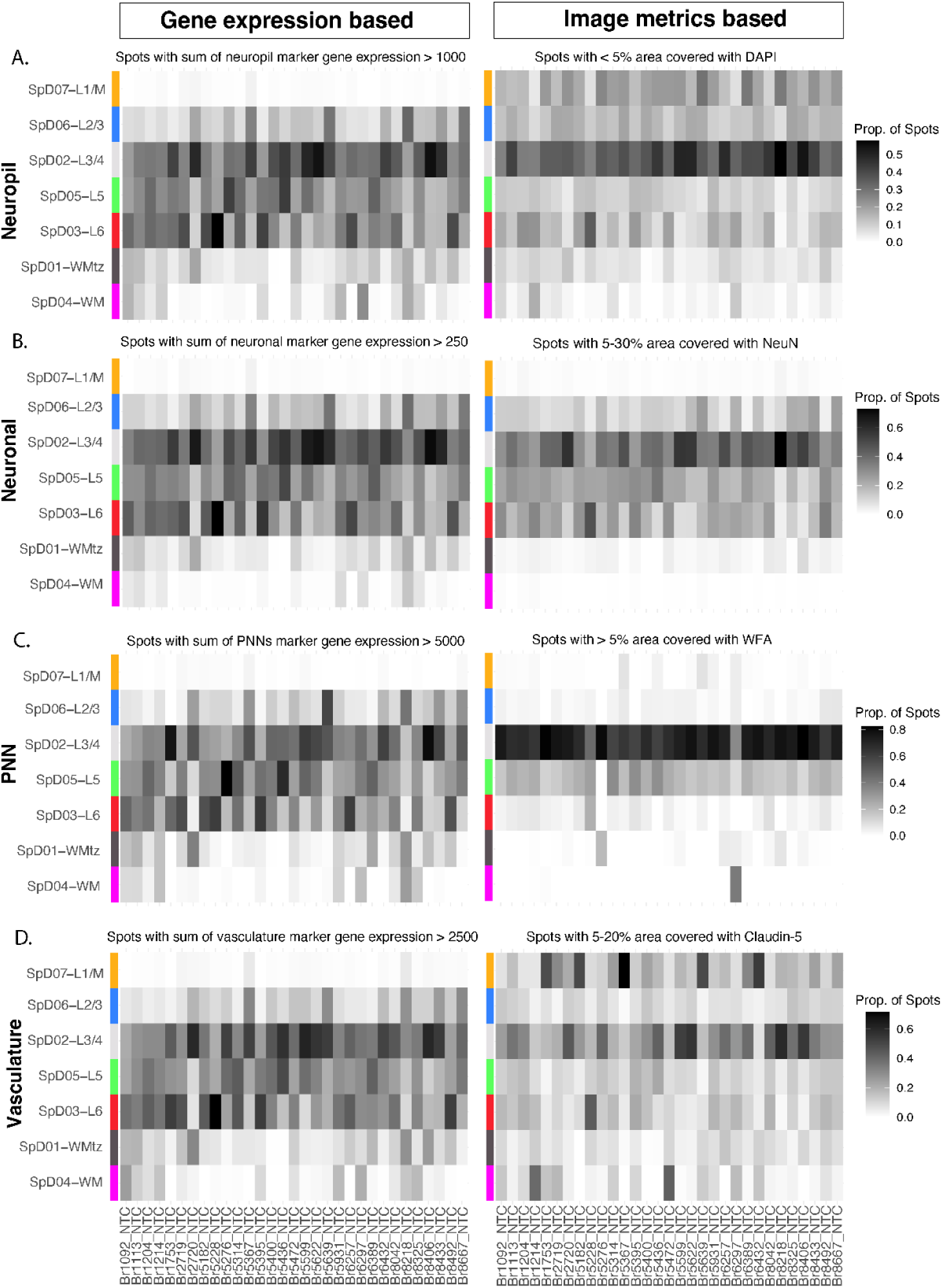
Enrichment of SPG spots per NTC donor across SpDs. (**A**) Proportion of neuropil spots identified: by summed marker gene expression on the left (spots with > 1,000 counts, marker genes from Niu et al.^39^), and by image-based metrics on the right (spots with < 5% DAPI signal coverage). (**B**) Left heatmap displays the proportion of neuronal spots identified by summed marker gene expression (spots with > 250 counts, marker genes from Huuki-Myers et al.^25^), compared with the right heatmap showing the proportion of neuronal spots identified using image-based metrics (spots with 5-30% NeuN signal coverage). (**C**) Left heatmap displays the proportion of PNN spots identified by summed marker gene expression (spots with > 5,000 counts, marker genes from Lupori et al.^30^), compared with the right heatmap showing the proportion of PNN spots identified using image-based metrics (spots with > 5% WFA signal coverage). (**D**) Left heatmap displays the proportion of vasculature spots identified by summed marker gene expression (spots with > 2,500 counts, marker genes from Garcia et al.^40^), compared with the right heatmap showing the proportion of vasculature spots identified using image-based metrics (spots with 5-20% Claudin-5 signal coverage).

**Figure S15.**
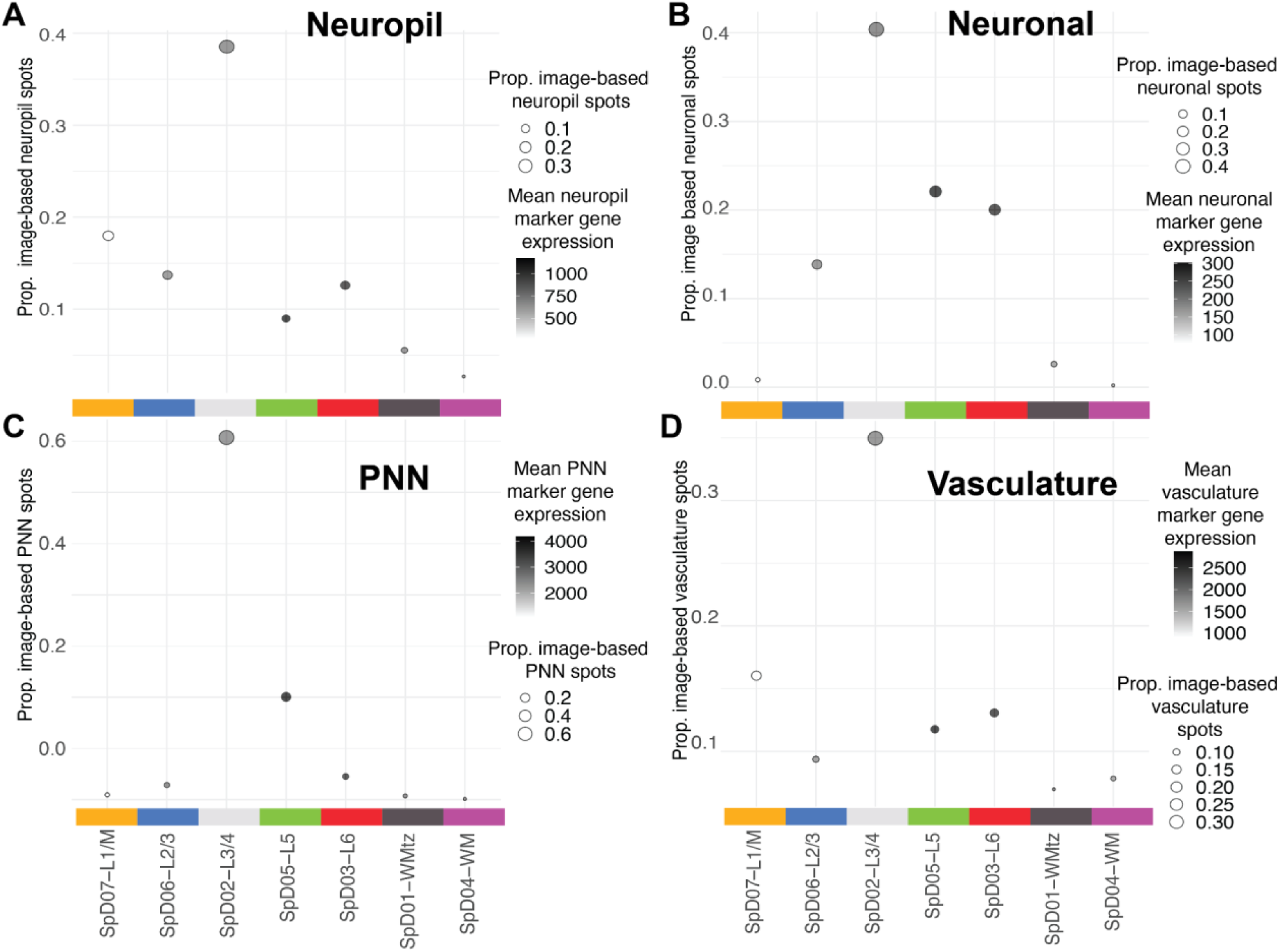
Validation of SPG spots using reference datasets. (**A**) Dot plots show the proportion of image-based neuropil spots (identified as spots lacking DAPI IF signal) per SpD (dot size) and the mean of summed marker gene expression from Visium spots within the SpD (dot color). Image-based SPG spot classification validated with gene expression of marker gene list derived from Niu et al.^39^ (Supple_Table_10_Neuropil, sheet=10_human_synase_markers). (**B**) Neuronal spots are identified as spots containing 5-30% coverage of segmented NeuN signal. Dot plots show the proportion of image-classified neuronal spots per SpD (dot size) and the mean of summed marker gene expression from Visium spots within the same domain (dot color). Marker gene list derived from Huuki-Myers et al.^25^ (TableS13_marker_stats_supp_table.xlsx, sheet=marker_stats_supp_table). (**C**) PNN spots are identified as spots containing > 5% coverage segmented WFA signal. Dot plots show the proportion of image-classified PNN spots per SpD (dot size) and the mean of summed marker gene expression from Visium spots within the same domain (dot color). Marker gene list derived from Lupori et al.^30^ (Supple_Table_2_Vas.xlsx, sheet=Post Mortem Vascular Subcluster). (**D**) Vasculature spots are identified as spots containing 5-20% coverage of segmented Claudin-5 signal. Dot plots show the proportion of image-classified vasculature spots per SpD (dot size) and the mean of summed marker gene expression from Visium spots within the same domain (dot color). Marker gene list derived from Garcia et al.^40^ (Supple_Table_DataS4_PNN.xlsx, sheet=PNN Energy).

**Figure S16.**
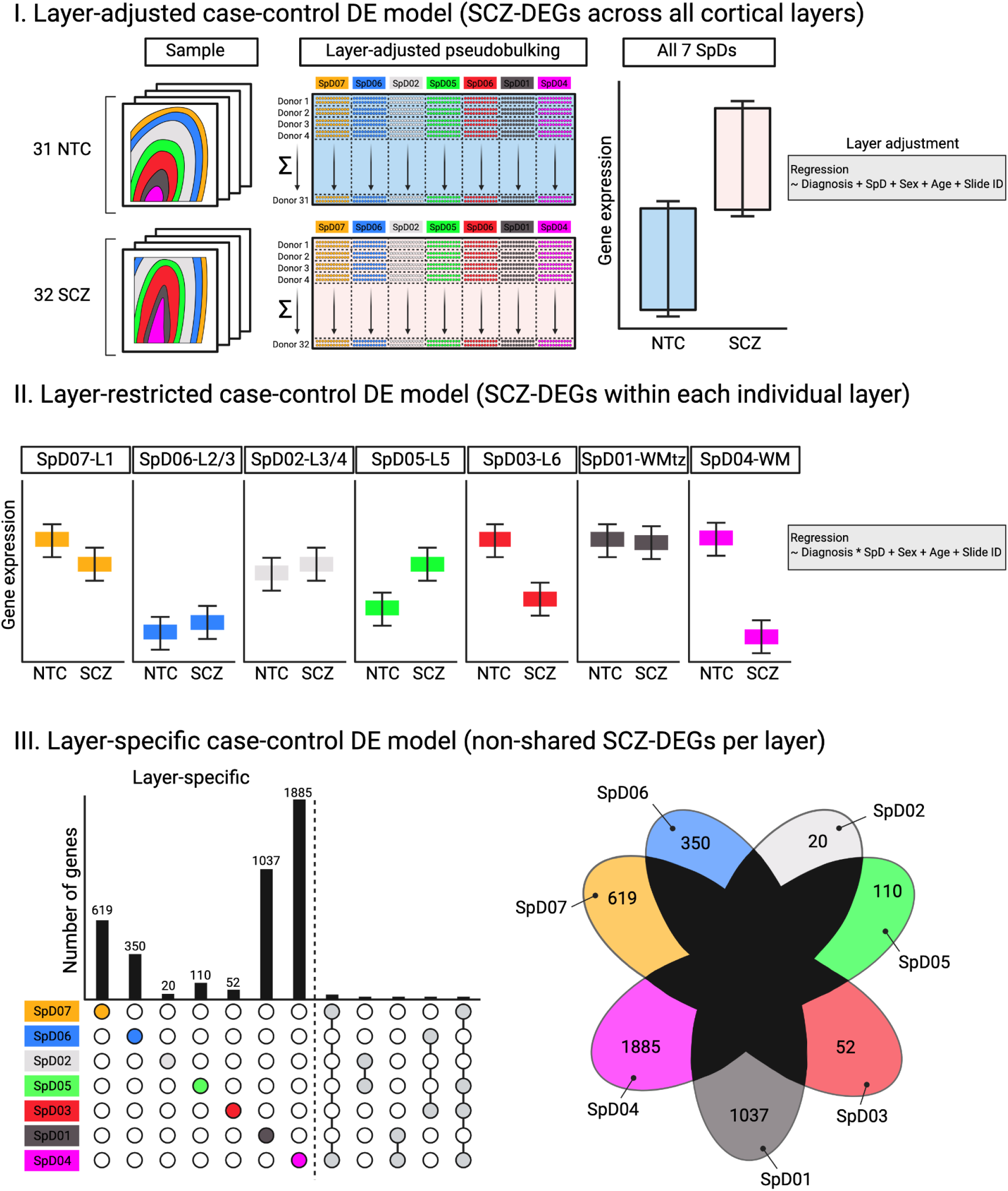
Schematic of statistical models for identifying SCZ-associated DEGs (SCZ-DEGs). Three statistical models were used (I) ‘layer-adjusted’ to identify SCZ-DEGs across all cortical layers, (II) ‘layer-restricted’ to identify SCZ-DEGs within each layer, and (III) ‘layer-specific’ to identify SCZ-DEGs that are unique to each layer. Created in BioRender. Kwon, S. H. (2026) https://BioRender.com/krzesaz

**Figure S17.**
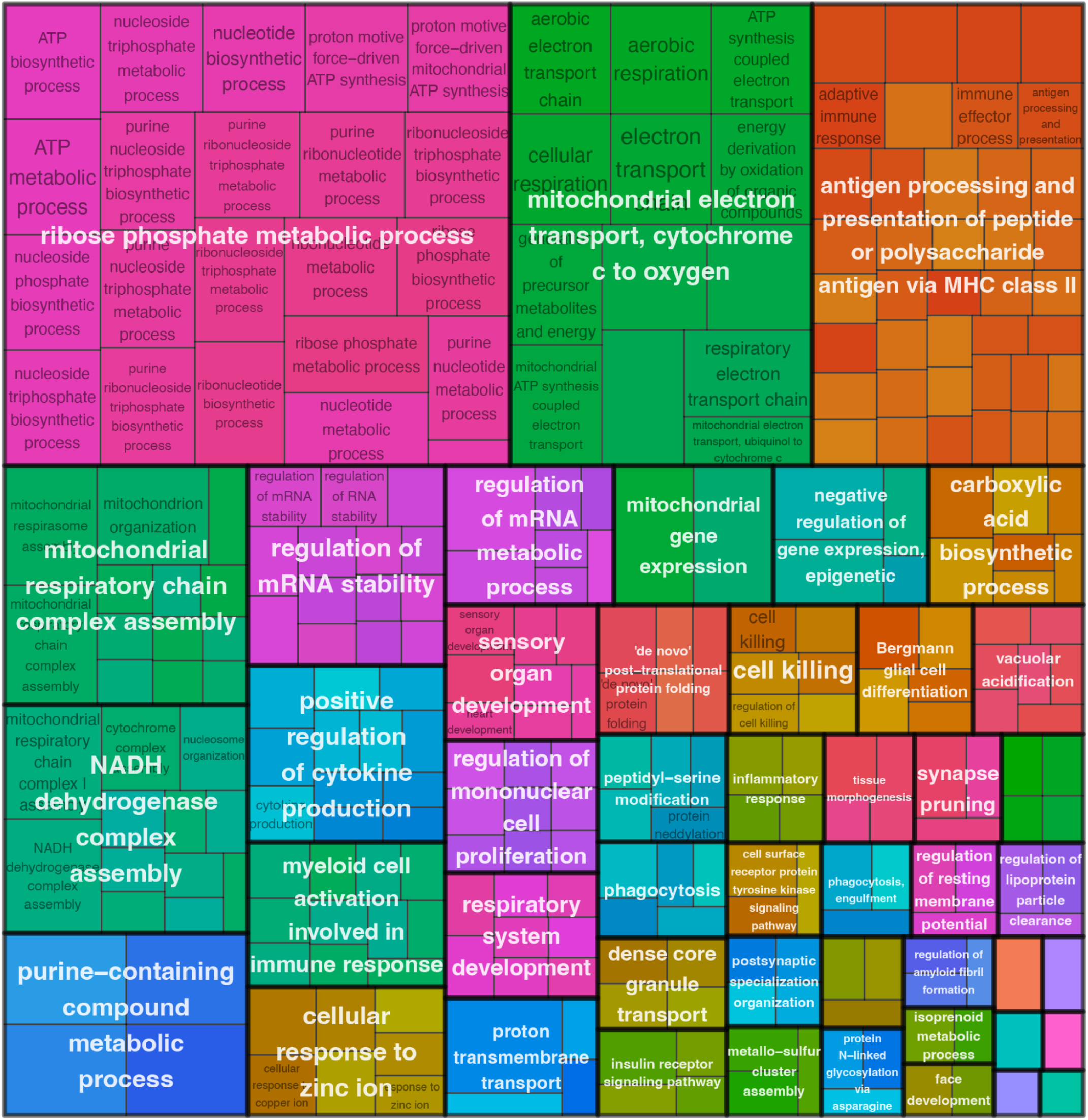
Treemap representation of GO-BP terms enriched by GSEA based on the layer-adjusted DE model. GSEA of all genes ranked by *t*-statistics from the layer-adjusted DE model identified significant overrepresentation of GO-BP terms (adjusted *p* < 0.05). Enrichment converging on immune-inflammatory responses and cellular responses to metal ions was observed, along with distinct enrichment of ATP metabolism and mitochondrial respiration, suggesting layer-wide metabolic alterations across cortical layers.

**Figure S18.**
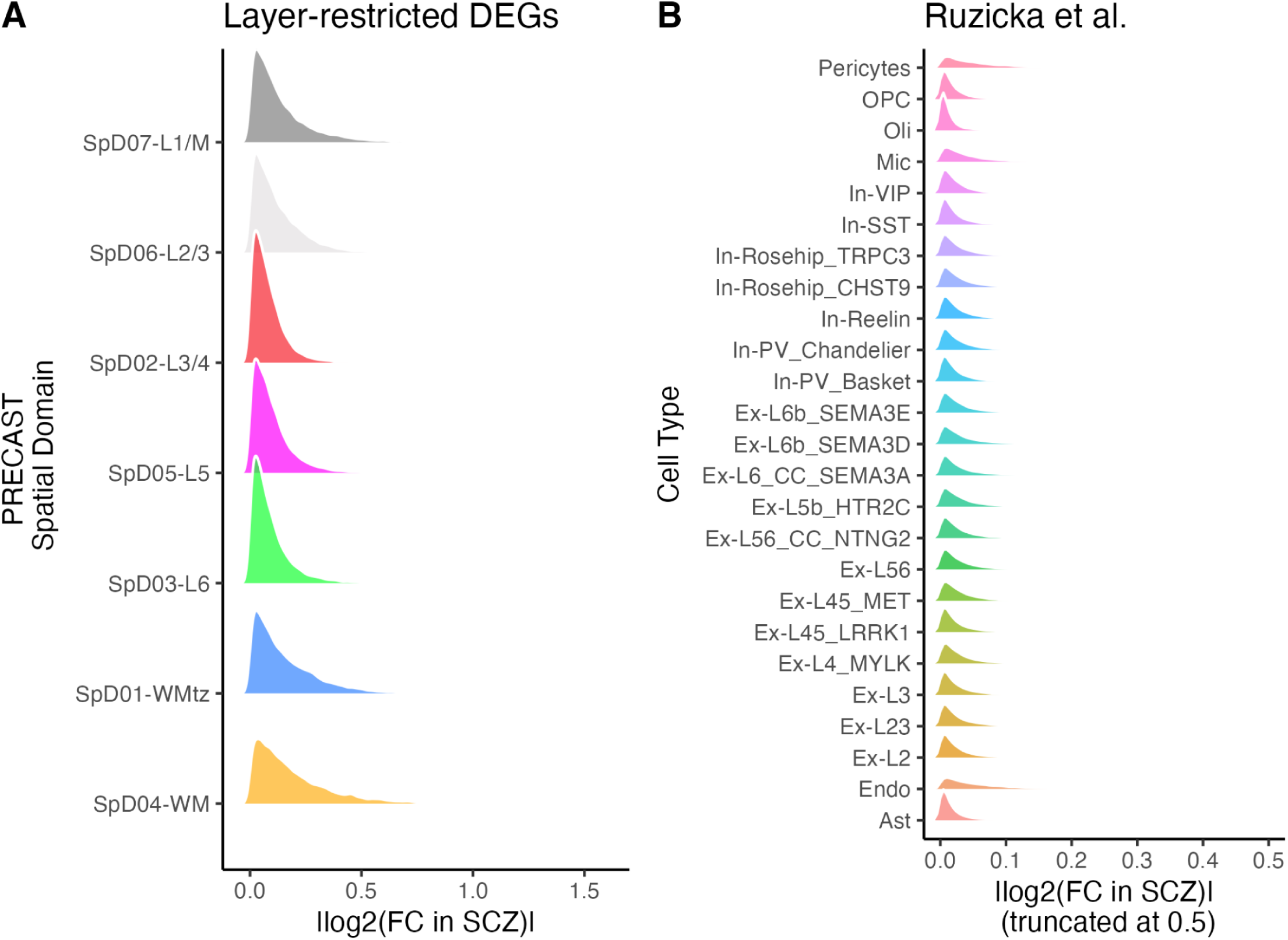
Comparing SCZ-associated gene expression changes detected in the current Visium SRT study and in a snRNA-seq study. (**A**) Distributions of absolute value of logFC (|log2FC|) in SCZ detected in the layer-restricted DE analysis, stratified by PRECAST SpDs. (**B**) Distributions of |log2FC| detected in cell-type-specific DE in Ruzicka et al.,^16^ stratified by cell types. Due to a small amount of extreme outliers, n=148 genes (0.04%) were removed from the ridge plot due to plot truncation at 0.5. Overall, there were more glial- and WM-related signals detected than neuronal signals in the current Visium SRT dataset. This observation is consistent with cell-type-specific gene expression changes reported in Ruzicka et al..^16^ Future SRT studies may require a larger sample size to detect nuanced neuronal signals in the neuronal-rich cortical domains.

**Figure S19.**
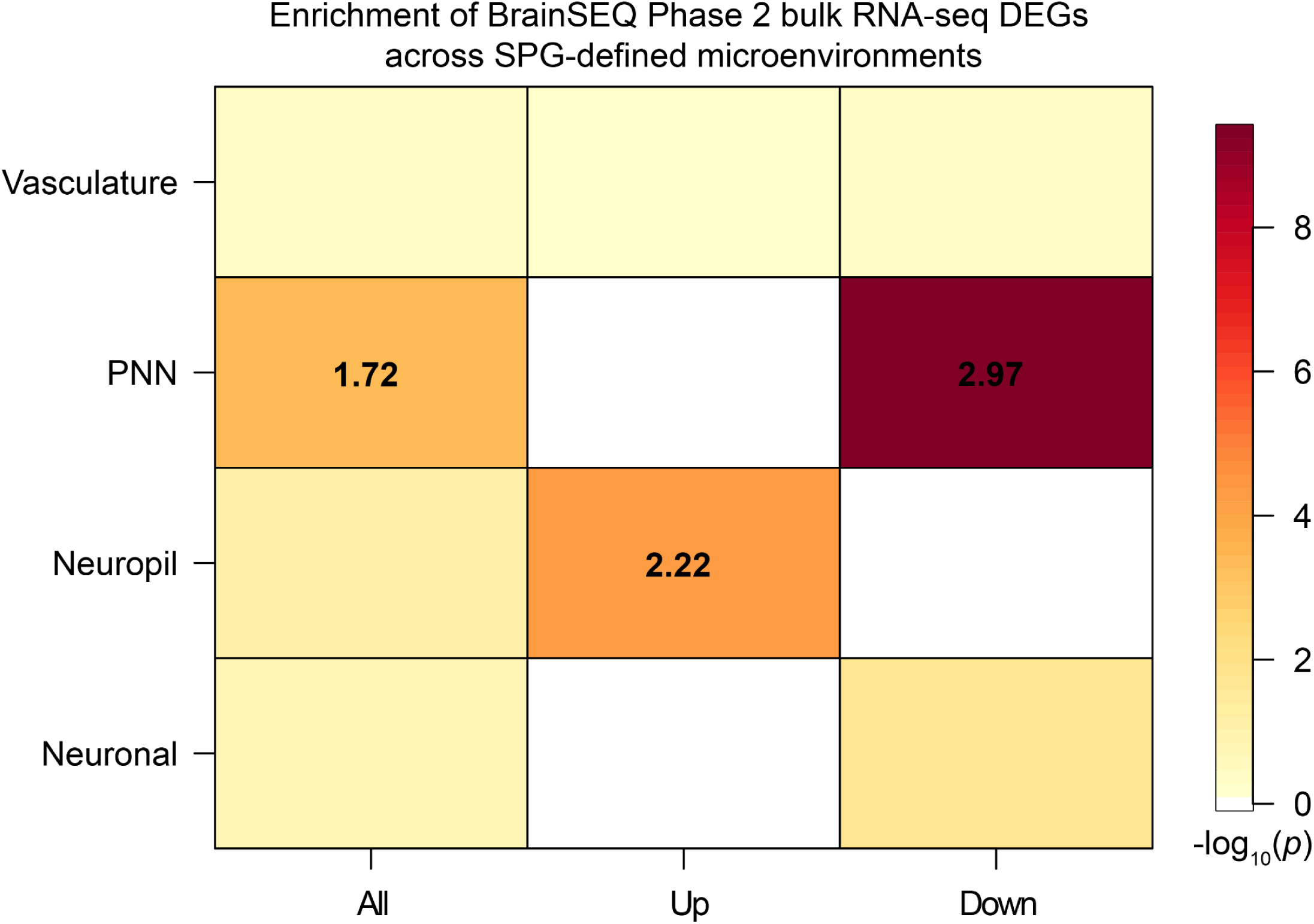
Enrichment of BrainSEQ Phase 2 bulk RNA-seq DEGs in SPG spots across four SPG-defined microenvironments. DEGs previously identified in the BrainSEQ Phase 2 bulk RNA-seq study^41^ were tested for enrichment in SPG spots representing distinct cellular and extracellular microenvironments (All: both up- and down-regulated bulk RNA-seq DEGs; Up: up-regulated bulk RNA-seq DEGs only; Down: down-regulated bulk RNA-seq DEGs only). Robust enrichment was observed in PNN spots, particularly for down-regulated DEGs, whereas up-regulated DEGs were predominantly enriched in neuropil spots. These findings emphasize the significance of synaptic and extracellular components in SCZ and demonstrate the utility of our SRT dataset in contextualizing prior findings within anatomical tissue compartments. Color scales indicate -log_10_ (*p*-value), denoting the significance of the enrichments, and the numbers within the cells indicate the odds ratios for the significant enrichments.

**Figure S20.**
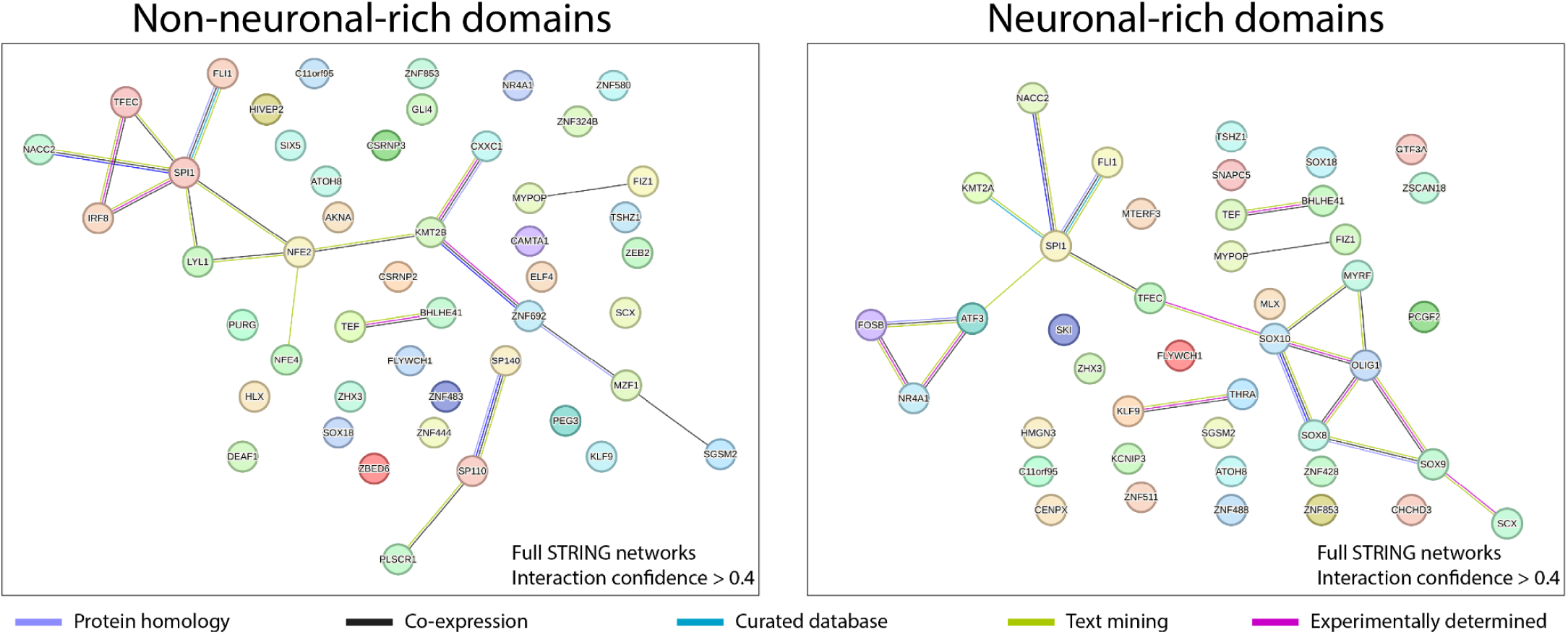
STRING network analysis of ChEA3-derived TFs linked to layer-restricted SCZ-DEGs across non-neuronal-rich and neuronal-rich cortical domains. STRING TF interaction networks for non-neuronal-rich (SpD07-L1/M, SpD01-WMtz, SpD04-WM) and neuronal-rich (SpD06-L2/3, SpD02-L3/4, SpD05-L5, SpD03-L6) domains highlighting functional and physical TF modules suggestive of transcriptional dysregulation hubs in SCZ. For each SpD, the top 10 TFs linked to both up- and down-regulated layer-restricted SCZ-DEGs per SpD were selected, yielding initial input sets of 60 TFs (non-neuronal) and 80 TFs (neuronal). After removing overlapping TFs and unmapped TFs (e.g., NKX62, NKX22), 48 TFs (non-neuronal) and 42 TFs (neuronal) remained for network construction. Networks are shown with all nodes including those without connections at an explorative interaction confidence (score > 0.4) compared to Figure 4B, revealing more extensive network connectivity.

**Figure S21.**
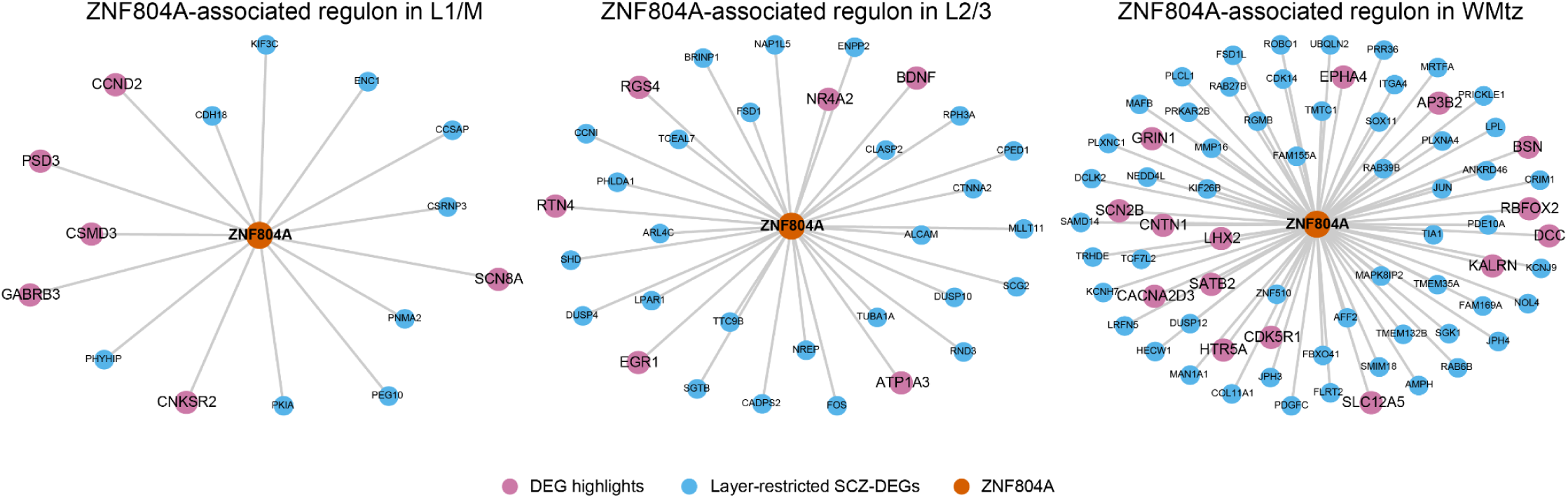
Network diagram illustrating the ChEA3-predicted ZNF804A regulon with its associated layer-restricted SCZ-DEGs across selected SpDs (SpD07-L1/M, SpD06-L2/3, and SpD01-WMtz). ZNF804A-associated SCZ-DEGs were identified using ChEA3 performed on input sets of both up- and down-regulated layer-restricted SCZ-DEGs. The networks shown here represent the ZNF804A-related subset of the full ChEA3 results (provided in **Supplementary Table 6**), highlighting the predicted regulons formed by ZNF804A and its associated SCZ-DEGs in the selected SpDs referenced in Figure 4C. Each panel shows the predicted associations between ZNF804A (orange) and its associated input SCZ-DEGs (blue). Edges indicate predicted TF-target associations only and do not represent interaction strength or biological significance. Neuron- and synapse-associated genes among the associated SCZ-DEGs are highlighted in pink.

**Figure S22.**
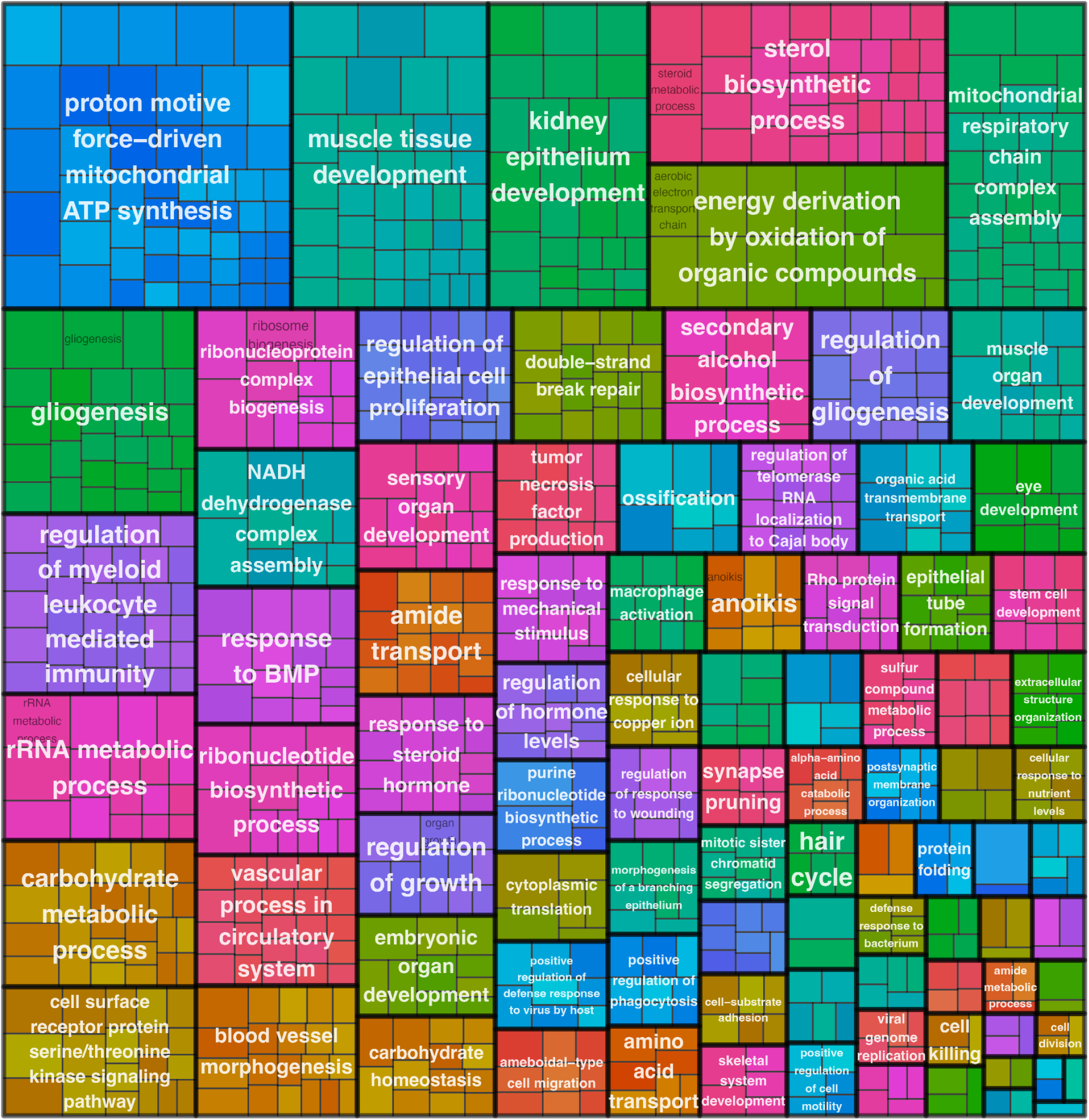
Treemap representation of GO-BP terms enriched by GSEA from the PRS-based DE model. GSEA of all genes ranked by *t*-statistics from the PRS-based DE model illustrates biological pathways associated with genetic risk for SCZ, identifying significant GO-BP terms (adjusted *p* < 0.05). Enrichment converged on metabolism, gliogenesis, immune regulation, and vascular-associated processes, some of which relate to non-neuronal cellular processes.

**Figure S23.**
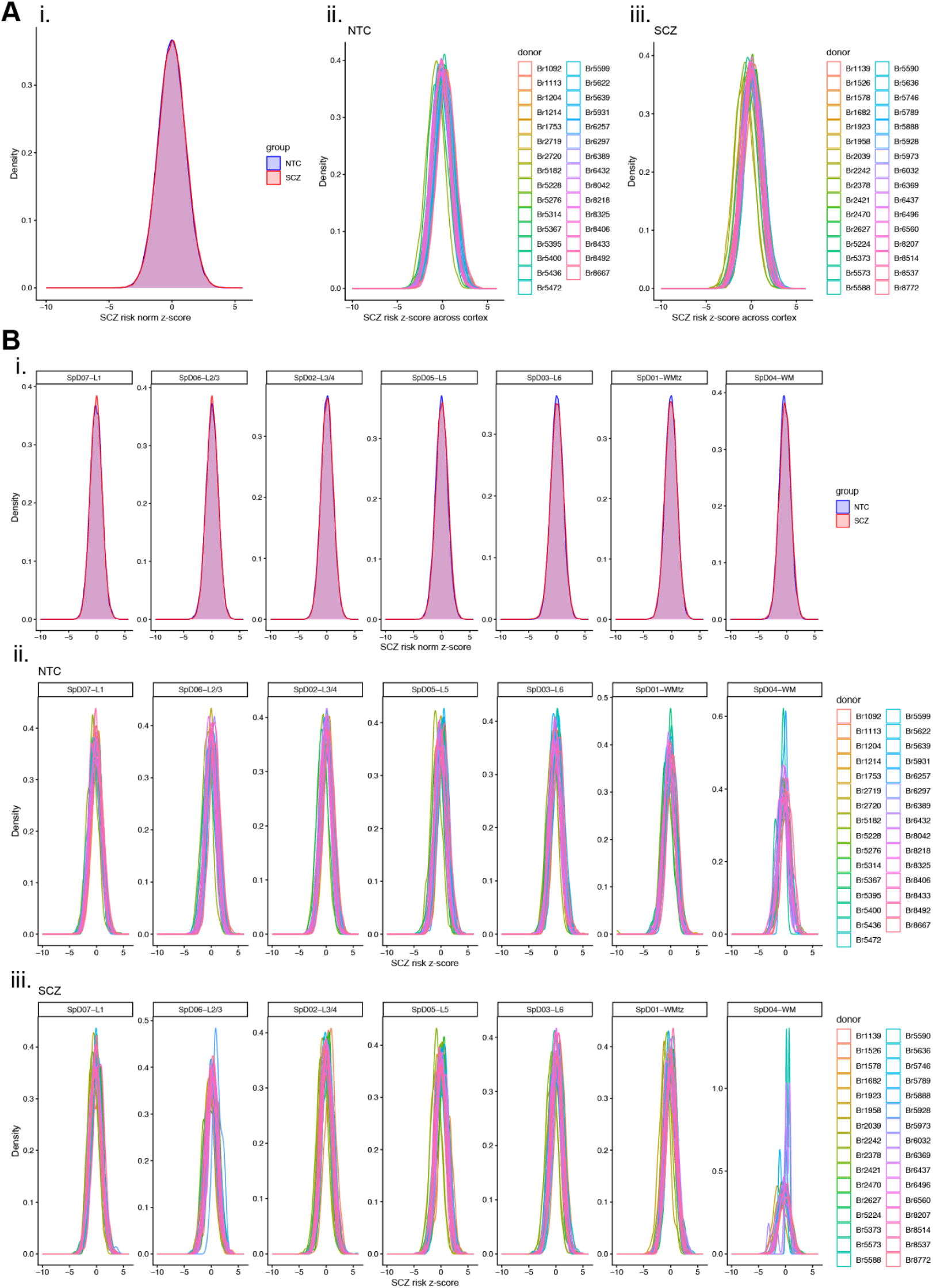
Density plots showing spatial distribution of SCZ scDRS *z*-scores across PRECAST SpDs. (**A**) Density plots of SCZ scDRS *z*-scores across PRECAST SpDs in all NTC and SCZ donors, colored by diagnostic group (i. normalized), and shown separately for (ii.) NTC and (iii.) SCZ donors. (**B**) Density plots of SCZ scDRS *z*-scores within each SpD across all NTC and SCZ donors, colored by diagnostic group (i. normalized), and shown separately for (ii.) NTC and (iii.) SCZ donors.

**Figure S24.**
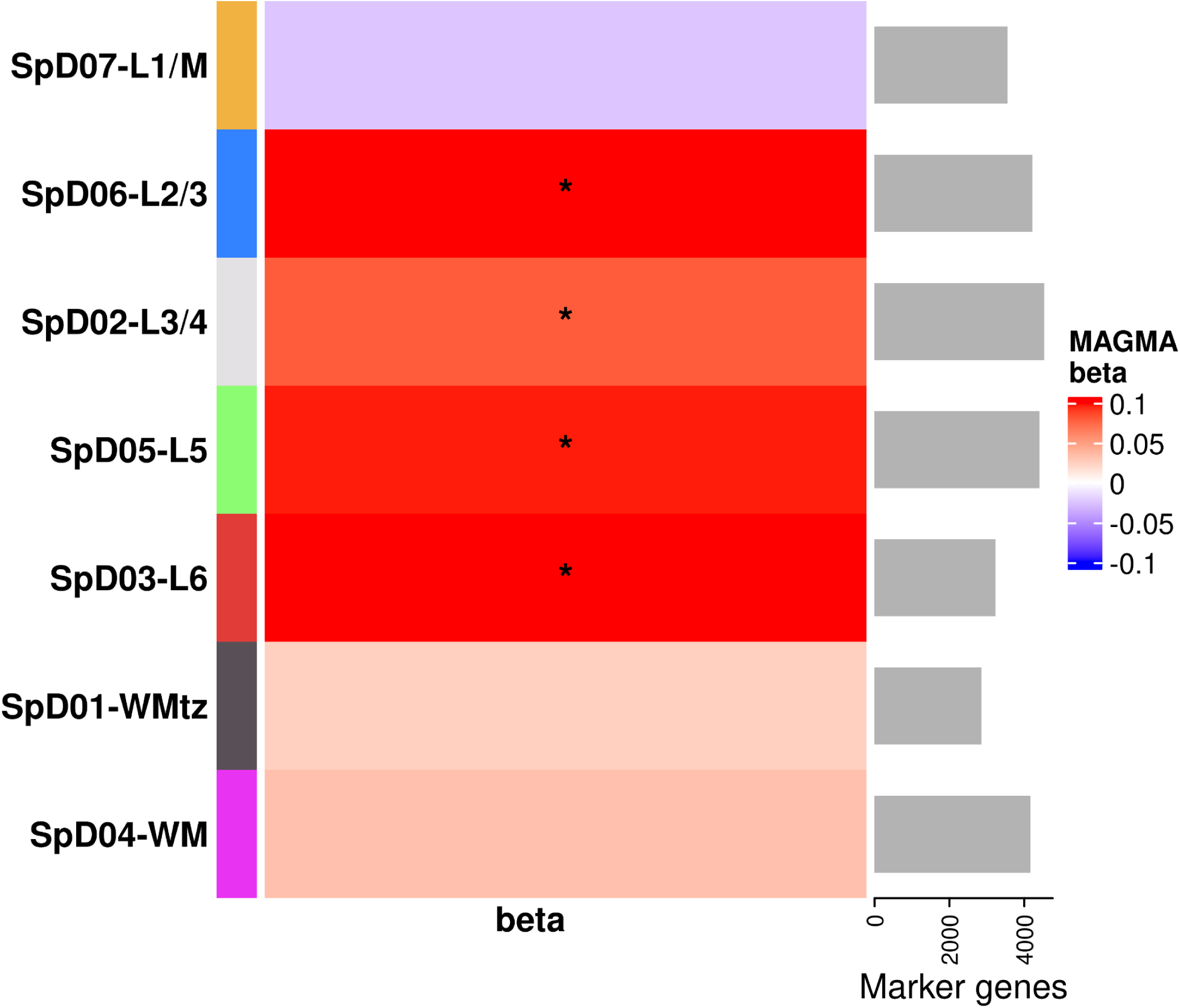
MAGMA gene set analysis showing predominant localization of SCZ risk enrichment patterns in neuronal-rich cortical domains (L2-6). The MAGMA^83^ heatmap shows the regression coefficient (beta) in each cortical layer with the asterisk denoting significant enrichment (FDR-adjusted *p* < 0.05) of each layer’s marker genes (# in bar plot) in SCZ GWAS. Relatively high SCZ risk enrichment patterns across neuronal-rich domains spanning L2-6 were observed, consistent with the scDRS *z*-score analysis in **Figure 4H**.

**Figure S25.**
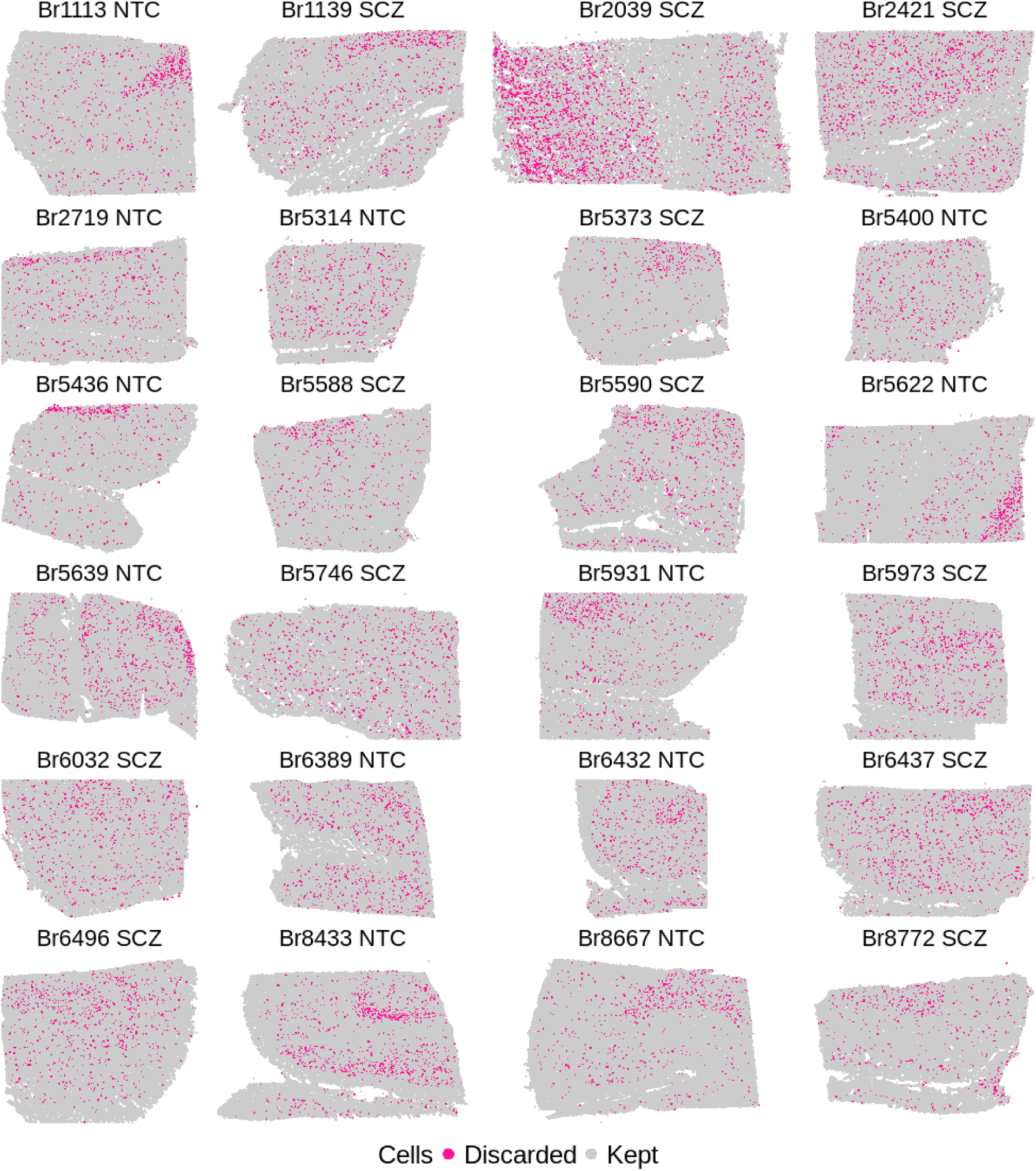
Cell-level plots of excluded cells in the Xenium dataset. Each point represents a single cell for the n=12 NTC and n=12 SCZ tissue samples. Discarded cells are pink while retained cells are gray. Cells with expression of negative control probes, negative control codewords, or unassigned codewords above the 99th percentile of all cells within a tissue section were discarded. Cells that were outliers in terms of number of genes detected (5 MADs below the median) or in terms of total counts (5 MADs above or below the median) were also discarded.

**Figure S26.**
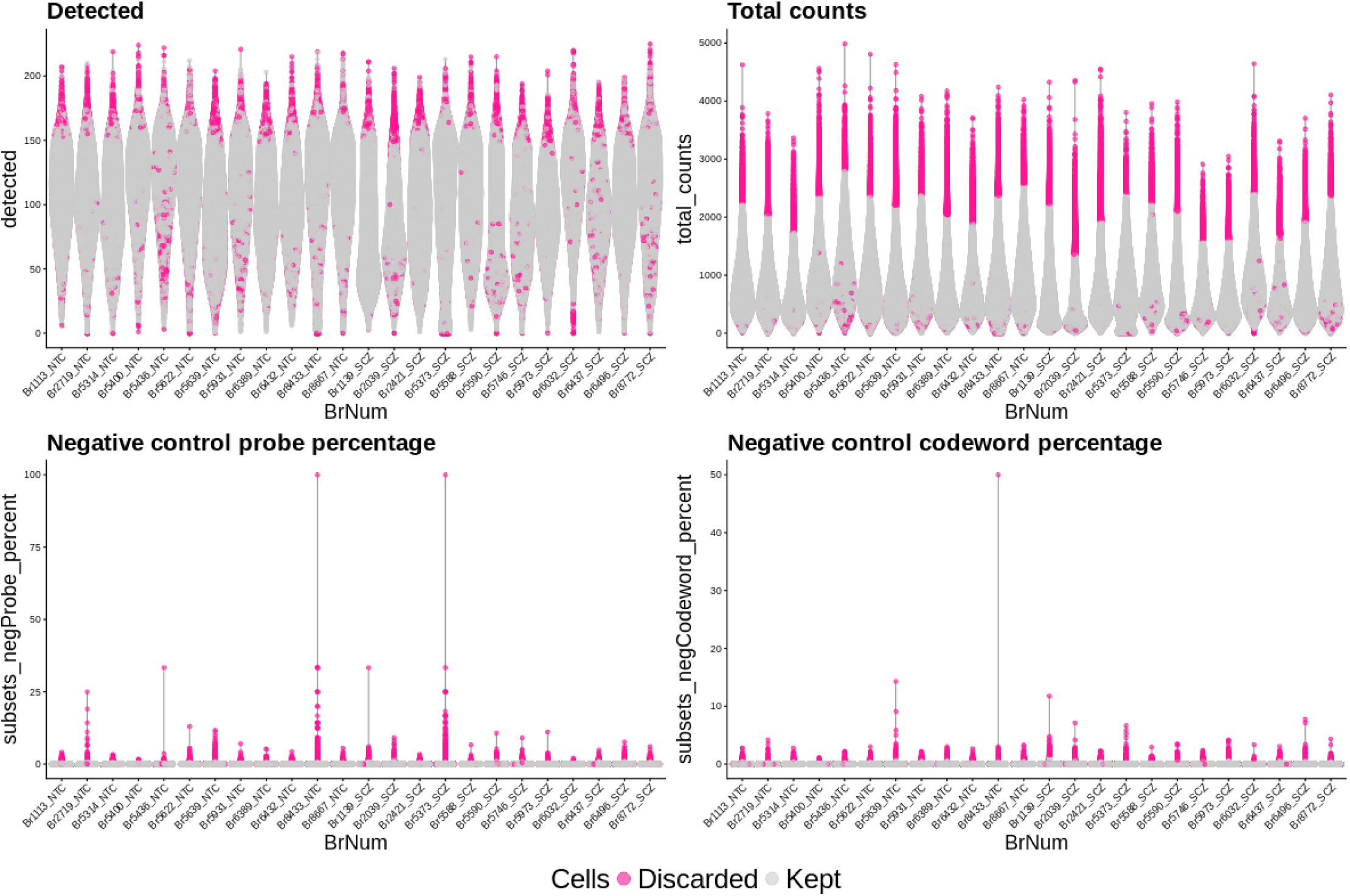
Violin plots showing distribution of QC metrics within each sample. The per-cell QC metrics used were the number of detected genes, the total counts across all genes, the percentage of counts from negative control probes, and the percentage of counts from negative control codewords. Each point represents a cell, and cells are colored by whether they were discarded (pink) or kept (gray). Cells that were outliers in terms of number of genes detected (5 MADs below the median) or in terms of total counts (5 MADs above or below the median) were also discarded. Cells with expression of negative control probes, negative control codewords, or unassigned codewords above the 99th percentile of all cells within a tissue section were discarded.

**Figure S27.**
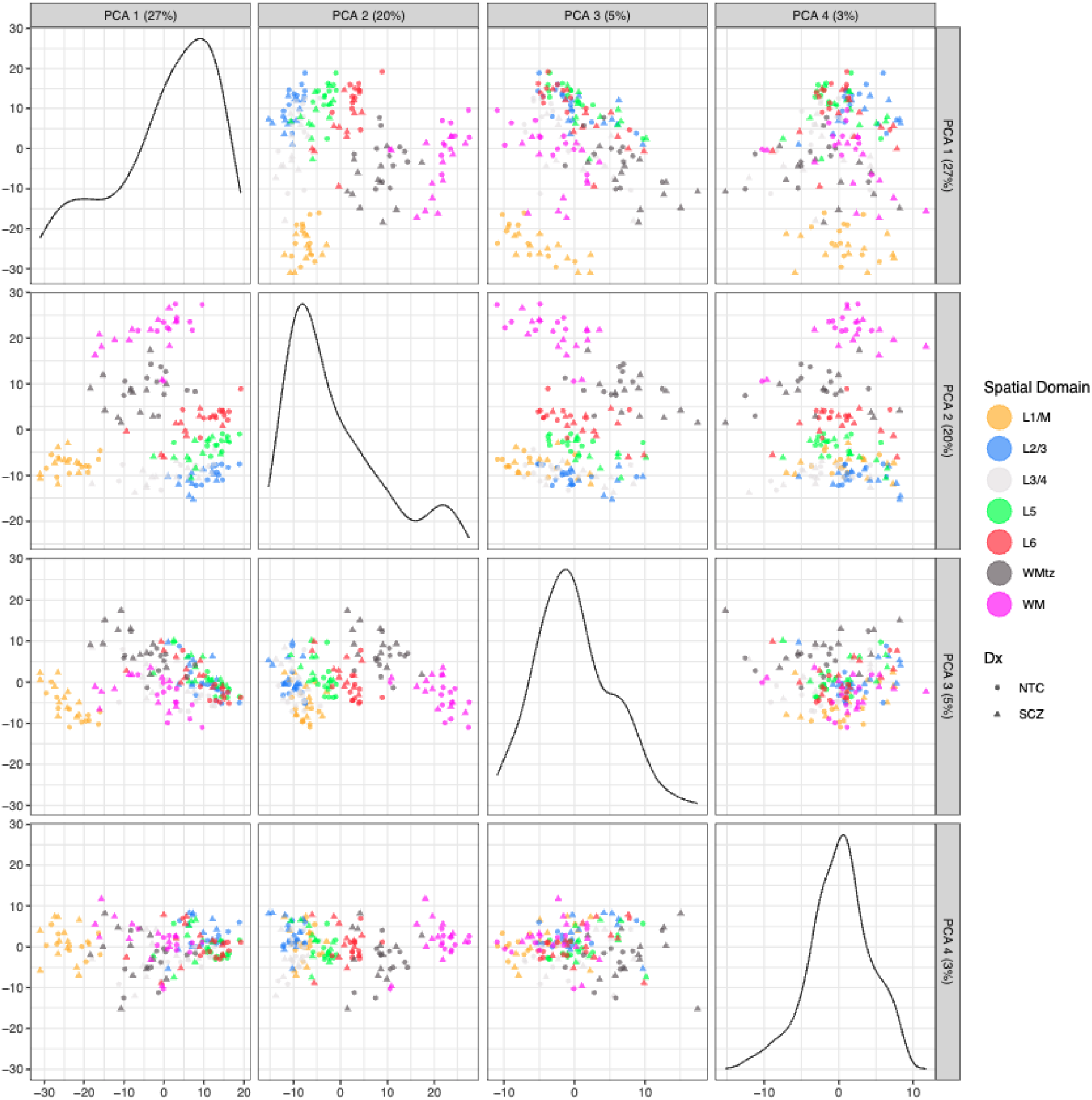
Scatter plots of the top 4 principal components (PC) from principal component analysis (PCA) performed on the pseudobulked Xenium data. Each point represents a unique donor-SpD combination. Points are shaped by the diagnosis of the donor and colored by the SpD. PC1 and PC2 both appear to be driven by the difference between the SpDs, with PC1 specifically being driven by SpD07-L1/M. For brevity, ‘SpD0X’ labels are not shown in the plot.

**Figure S28.**
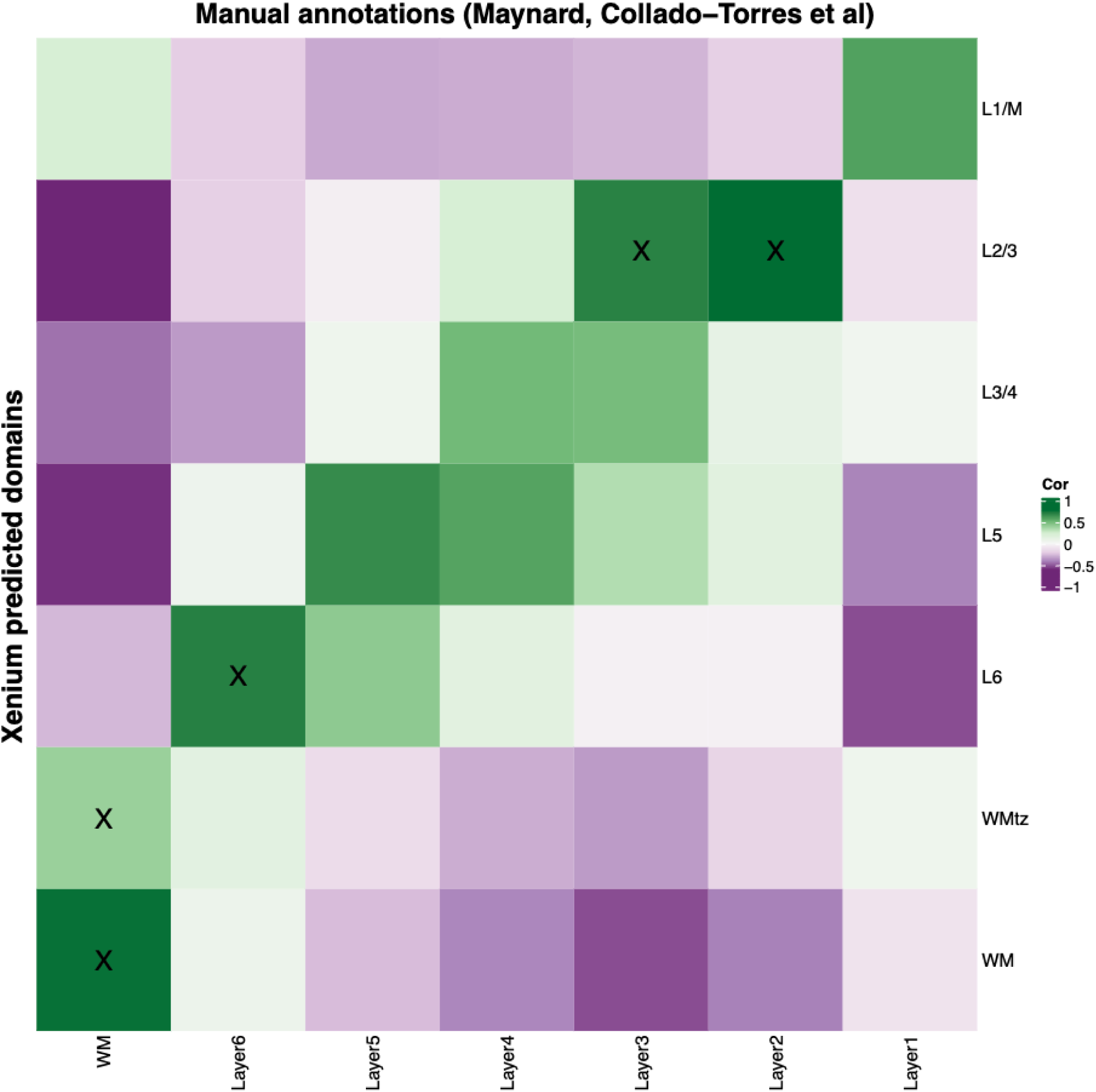
Spatial registration correlation heatmap of Xenium predicted cell-level domains against manual annotations. Spatial registration of spaTransfer predicted domains on the Xenium data (y-axis) against manual annotations of dlPFC layers from Maynard, Collado-Torres et al..^27^ Higher values (green) indicate higher Pearson correlation in enrichment *t*-statistics between predicted cell-level domains and manual annotations. To validate our predictions, predicted cell-level domains were then annotated with their best matching manual annotation based on this correlation. Annotations with high confidence are denoted with an ‘X’. For brevity, ‘SpD0X’ labels are not shown in the heatmap.

**Figure S29.**
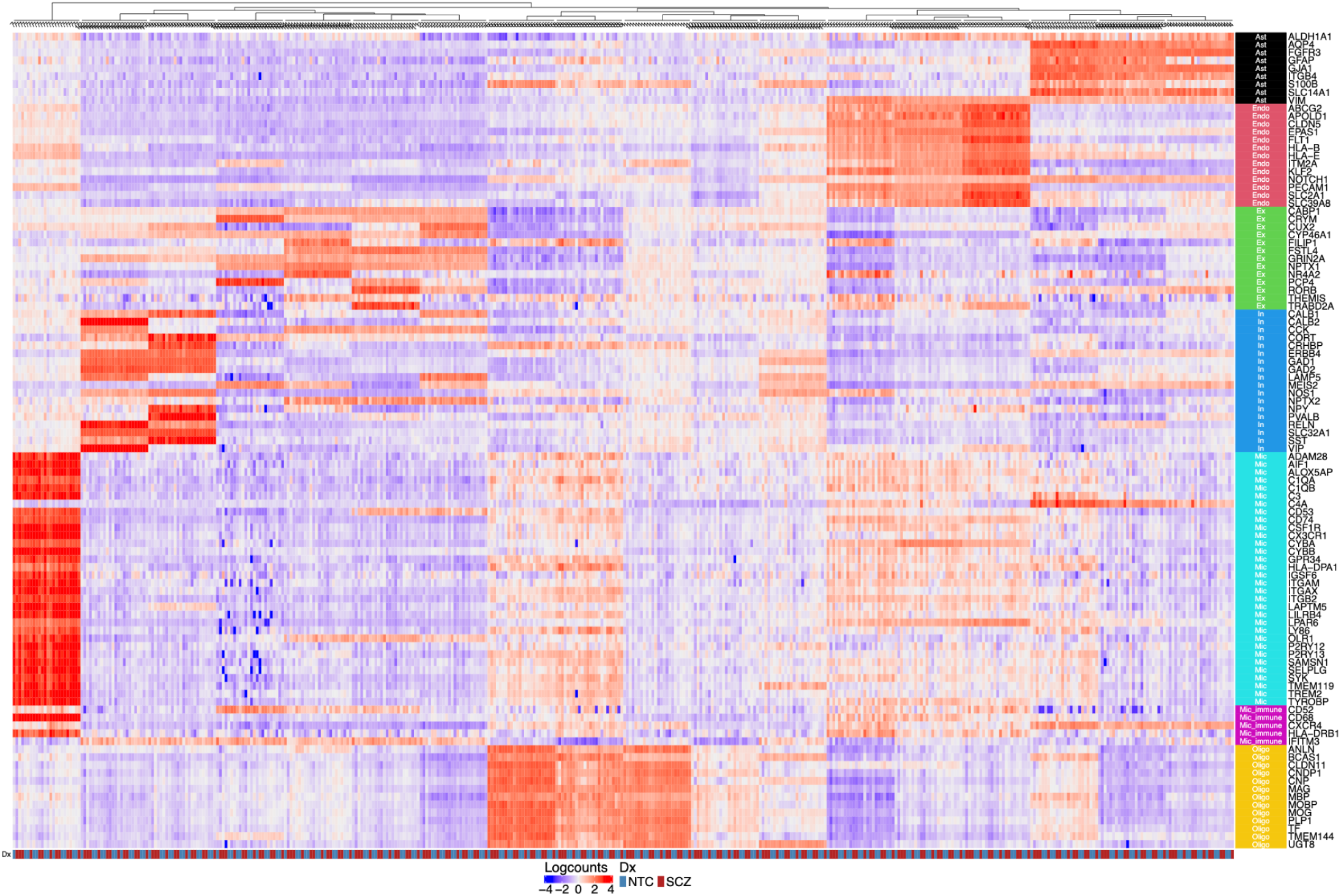
Heatmap of the marker gene expression for each of the 18 BANKSY clusters. Each column is a pseudobulked sample at the donor-cluster level, and each row is a cell-type marker gene included on the Xenium panel. Expression values are log-transformed library-size-normalized counts. Rows were standardized to have a mean of 0 and a variance of 1. Marker genes are grouped by their representative cell type. Cell type abbreviations: Ast, astrocytes; Endo, endothelial cells; Ex, excitatory neurons; In, inhibitory neurons; MIC, microglia; MIC_immune, microglia- or immune-associated cells; Oligo, oligodendrocytes.

**Figure S30.**
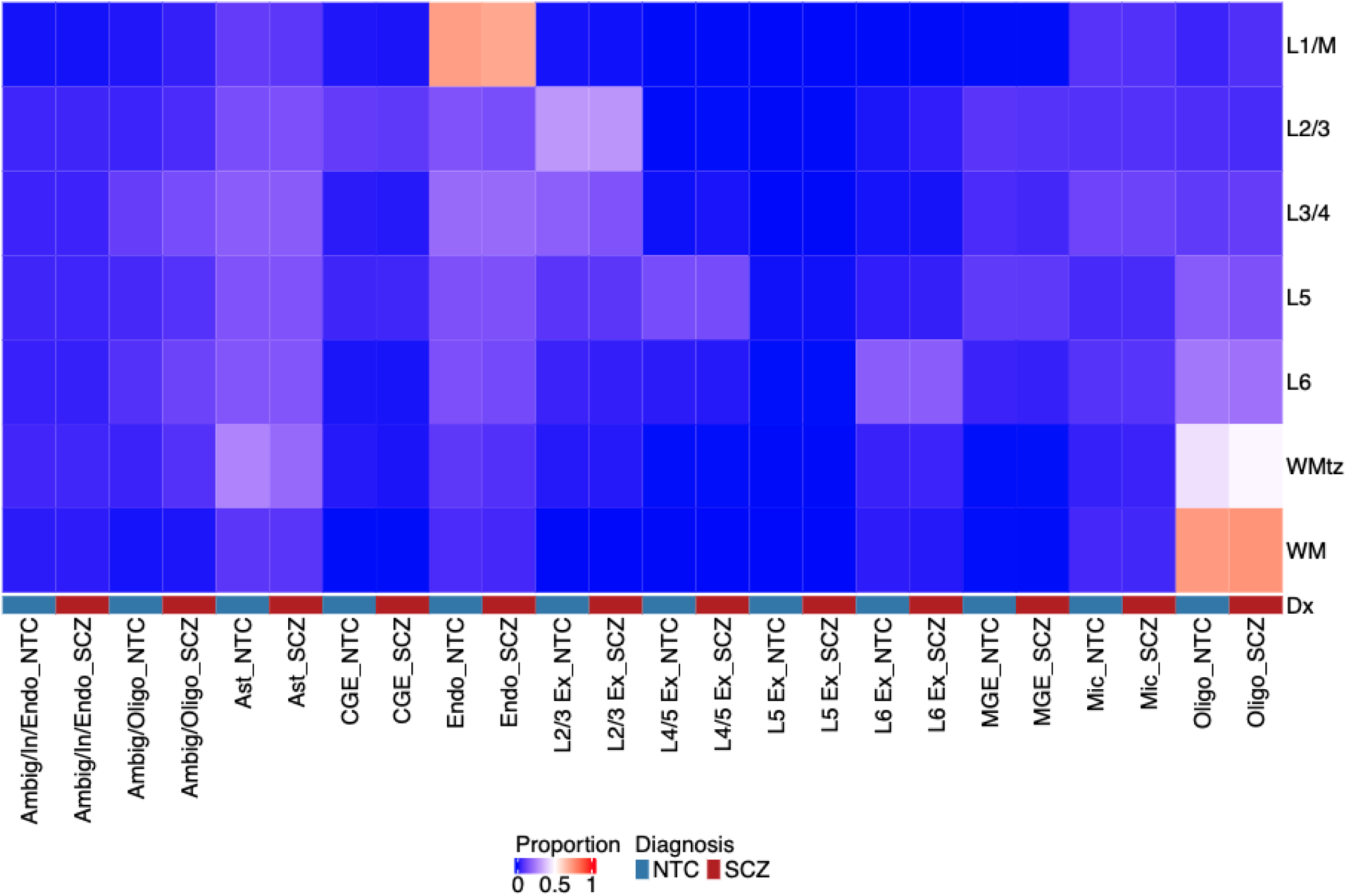
Heatmap of cell-type proportion within each SpD. Each column represents a unique cell type-diagnosis combination, and each row represents a SpD. The heatmap is colored by the proportion of a cell type within a SpD, computed across all donors within a diagnostic group. Across diagnosis, astrocytes and oligodendrocytes are enriched in the SpD04-WM and SpD01-WMtz, and endothelial cells are enriched in SpD07-L1/M. The neuronal cell types show enrichment in SpD06-L2/3 through SpD03-L6. For brevity, ‘SpD0X’ labels are not shown in the heatmap.

**Figure S31.**
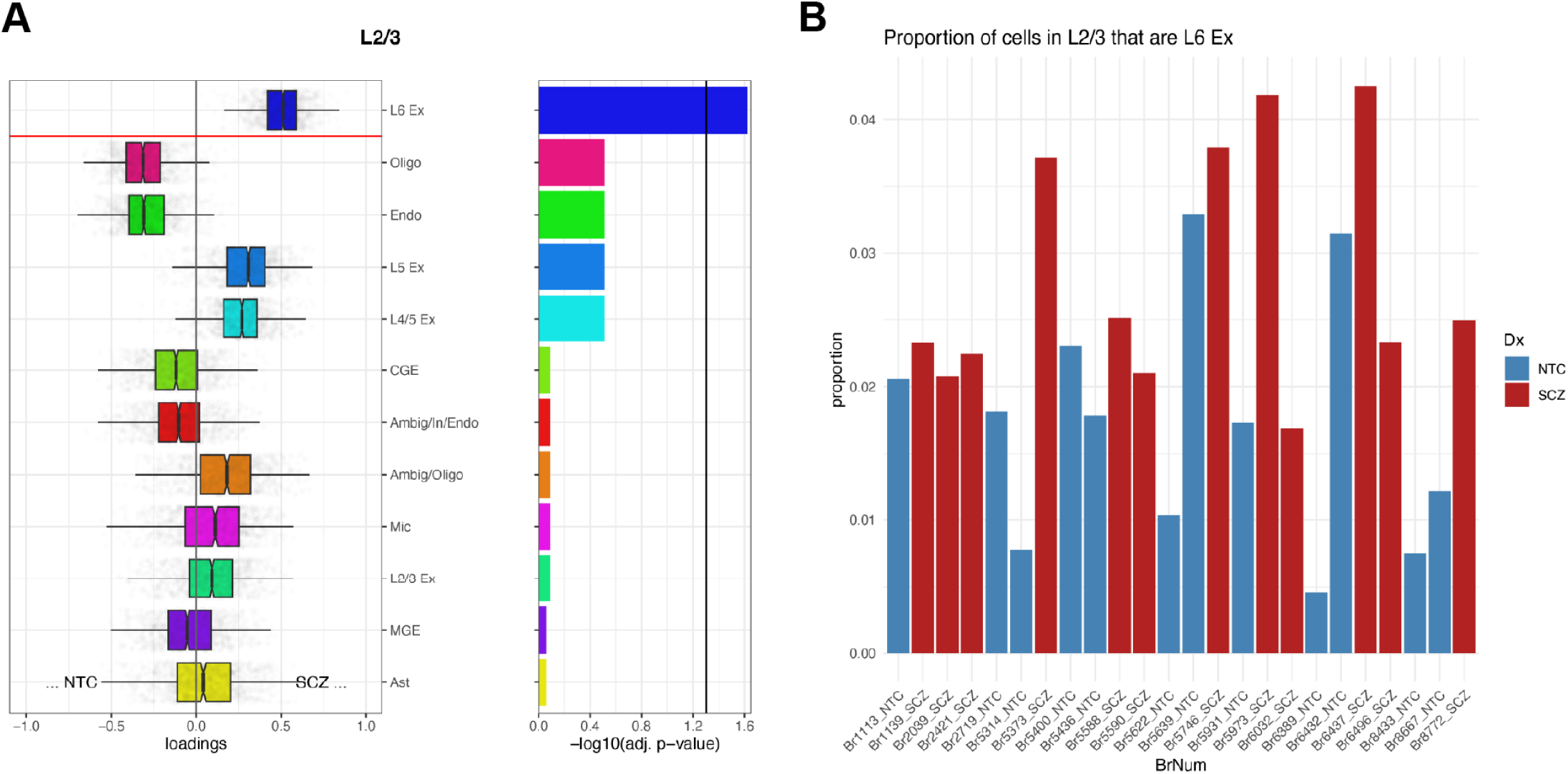
Cell-type composition analysis reveals enrichment of L6 Ex neurons in L2/3 in SCZ. (**A**) Compositional data analysis results for SpD06-L2/3 from the cacoa analysis.^98^ The x-axis of the box plot represents the loadings from linear discriminant analysis. Loadings greater than zero indicate enrichment in SCZ, and loadings less than zero indicate enrichment in NTC. Cell types in the box plot are ordered vertically from lowest to highest *p*-value. Cell types above the red line have statistically significant enrichment. The corresponding bar chart depicts -log_10_ (adjusted *p*-value). The black line indicates the *p*-value cutoff for significance. (**B**) Bar chart showing proportions of cells in SpD06-L2/3 that are L6 Ex neurons. Despite significant enrichment of L6 Ex neurons seen in (**A**), the proportion of L6 Ex neurons in L2/3 remains low for both NTC and SCZ. Notably, the percentage of L6 Ex neurons in SpD06-L2/3 ranged from 1.68% to 4.26% for SCZ donors.

**Figure S32.**
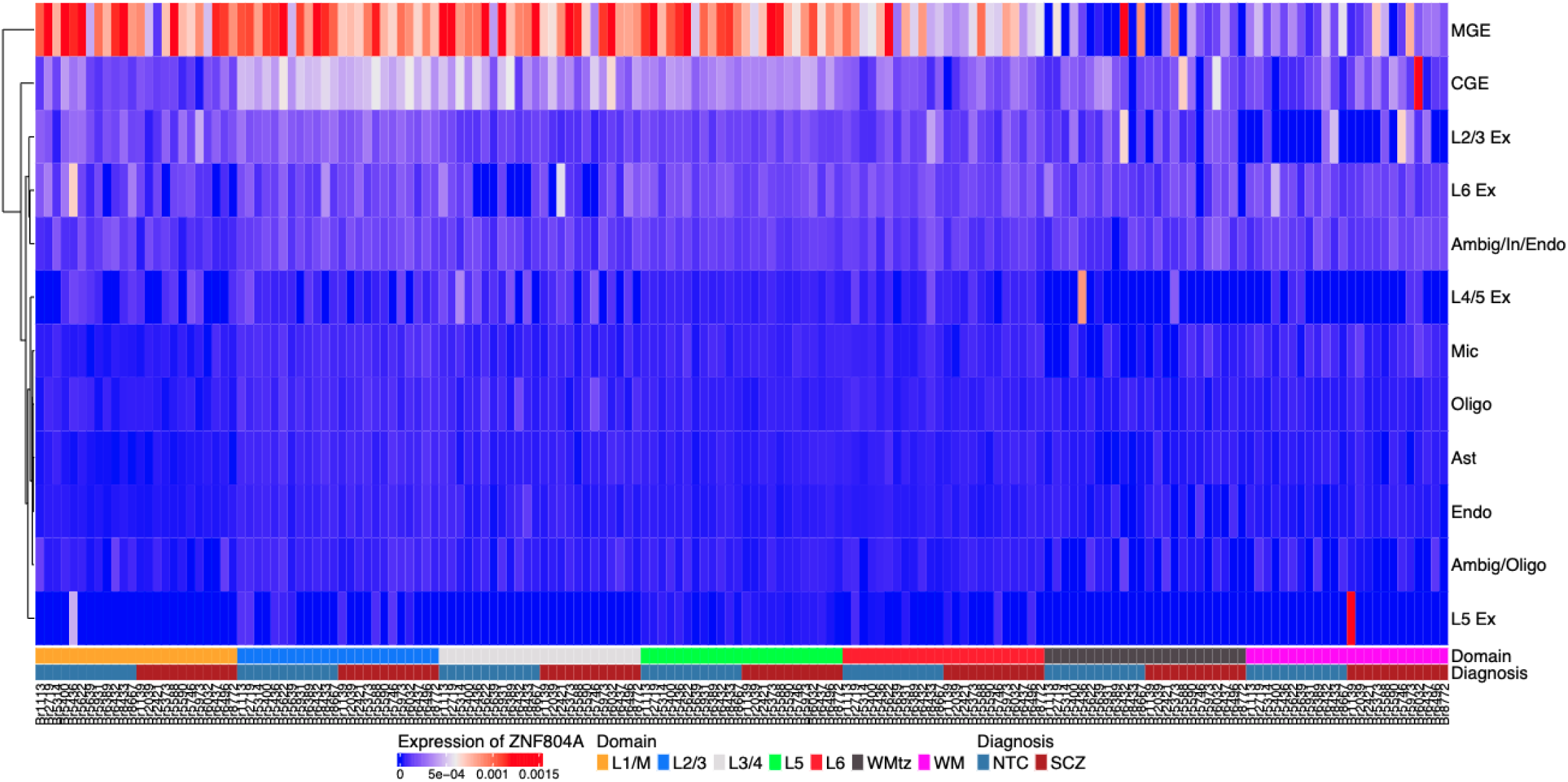
Heatmap showing pseudobulked expression of transcription factor ZNF804A in the Xenium data. Each column represents a unique pseudobulked donor-SpD combination and each row represents a cell type. Expression values are log-transformed library-size-normalized counts of *ZNF804A*. *ZNF804A* appears to be enriched in the MGE cell type across diagnosis.

**Figure S33.**
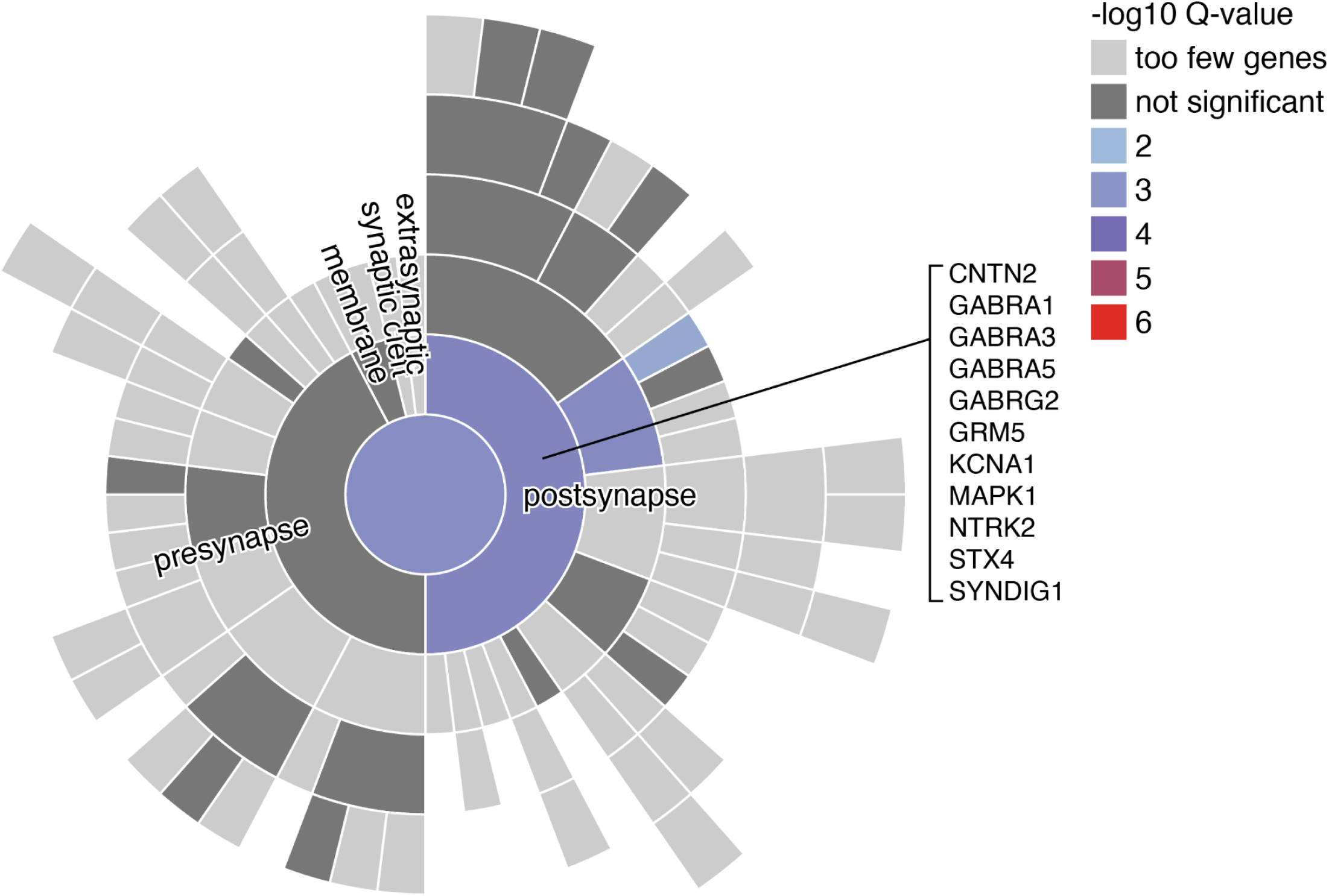
Complementary SynGO enrichment analysis of PNN microenvironment-restricted SCZ-DEGs. SynGO enrichment analysis (CC) was performed for microenvironment-restricted SCZ-DEGs identified at nominal *p* < 0.05 within PNN microenvironments using the default ‘brain-expressed’ background gene set, supporting significant enrichment of genes associated with postsynaptic compartments at a stringent threshold (FDR < 0.01). Tile color indicates enrichment significance (-log_10_ (*q*-value)), with representative synaptic genes labeled.

**Figure S34.**
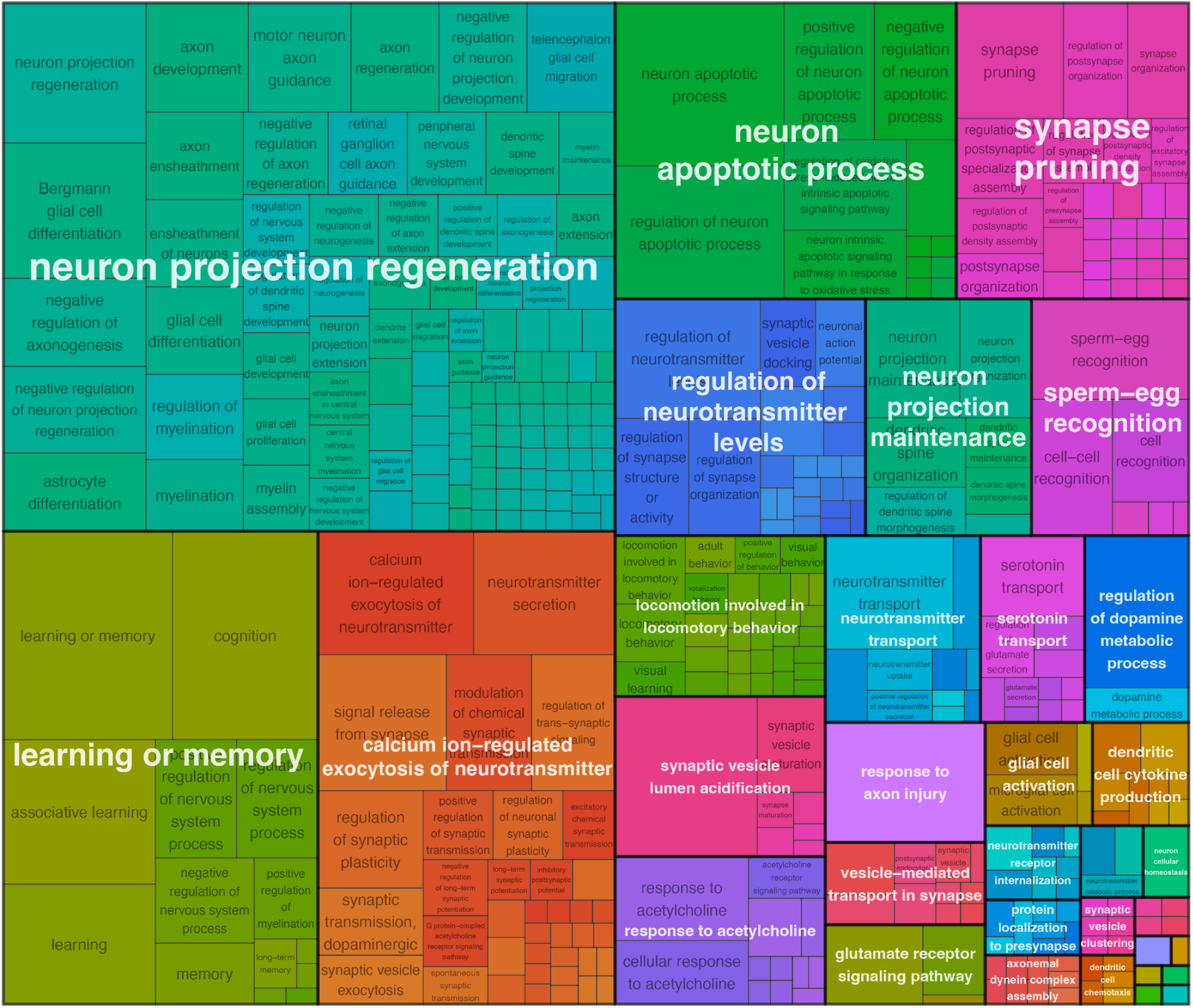
GO-BP enrichment treemap highlighting neuronal-related biological processes in the SPG defined neuronal microenvironments. GO-ORA analysis (GO-BP, FDR ≤ 0.05) was performed on neuronal microenvironment-restricted SCZ-DEGs identified at nominal *p* < 0.05. The top 50 neuron-associated enriched GO-BP terms (similarity threshold=0.7) were summarized and visualized as a treemap, supporting enrichment of neuronal signatures within these microenvironment-restricted SCZ-DEGs. Tile size reflects enrichment significance (-log_10_ (adjusted *p*-value)).

**Figure S35.**
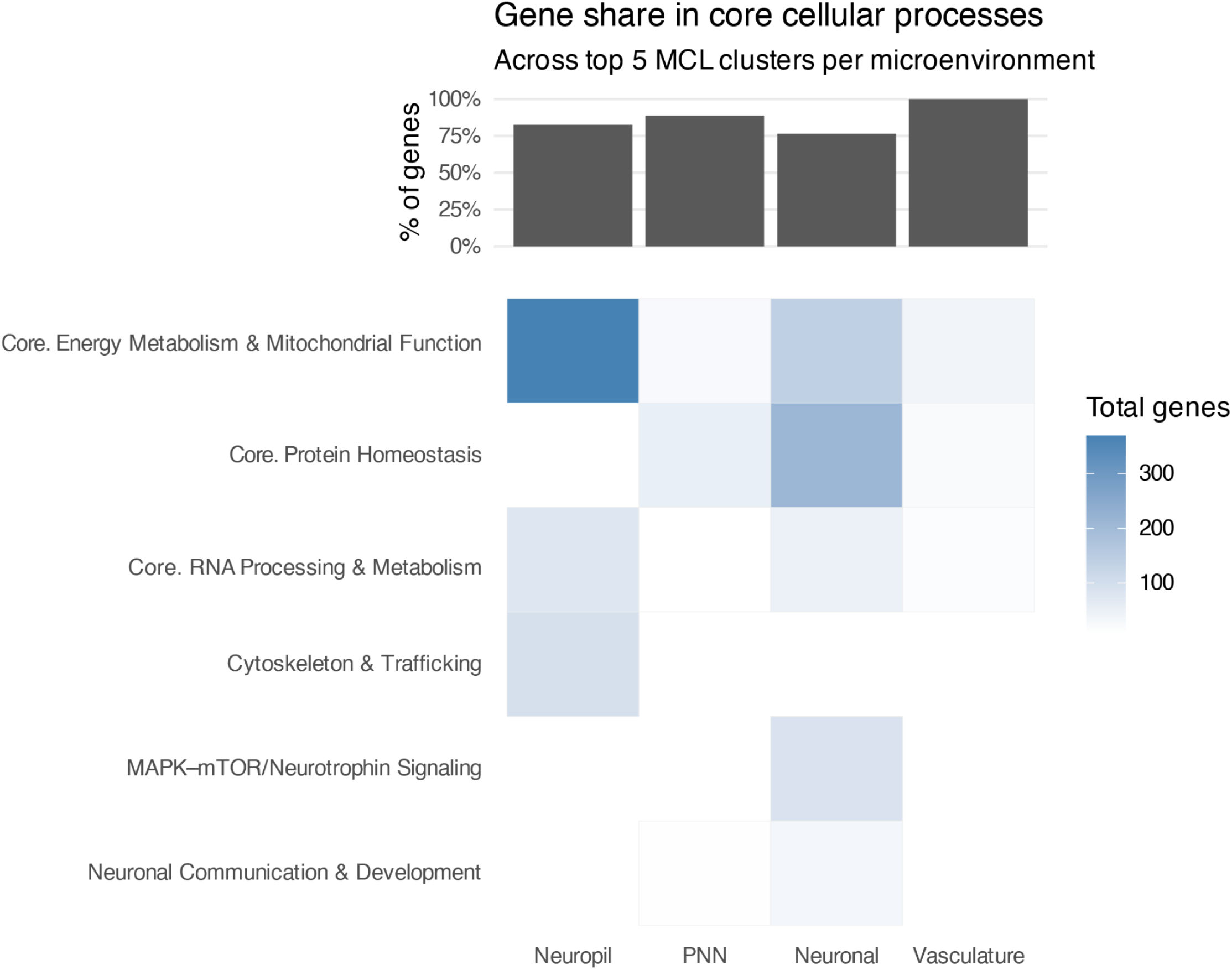
Heatmap and bar plot summarizing the representation of key biological processes across microenvironments, focusing on the top 5 clusters (Clusters 1-5) identified from STRING-based protein-protein interaction (PPI) networks using Markov clustering (MCL). The bar plot (top) shows the gene share (%) of microenvironment-restricted SCZ-DEGs (nominal *p* < 0.05) contributing to three core cellular processes, 1) energy metabolism & mitochondrial function, 2) protein homeostasis, and 3) RNA processing & metabolism, calculated relative to the total number of genes aggregated across the top five clusters within each microenvironment. The heatmap (bottom) displays the total number of microenvironment-restricted SCZ-DEGs assigned to each biological category, with darker blue indicating higher gene counts. All analyses were performed separately for each microenvironment (neuropil, PNN, neuronal, vasculature).

**Figure S36.**
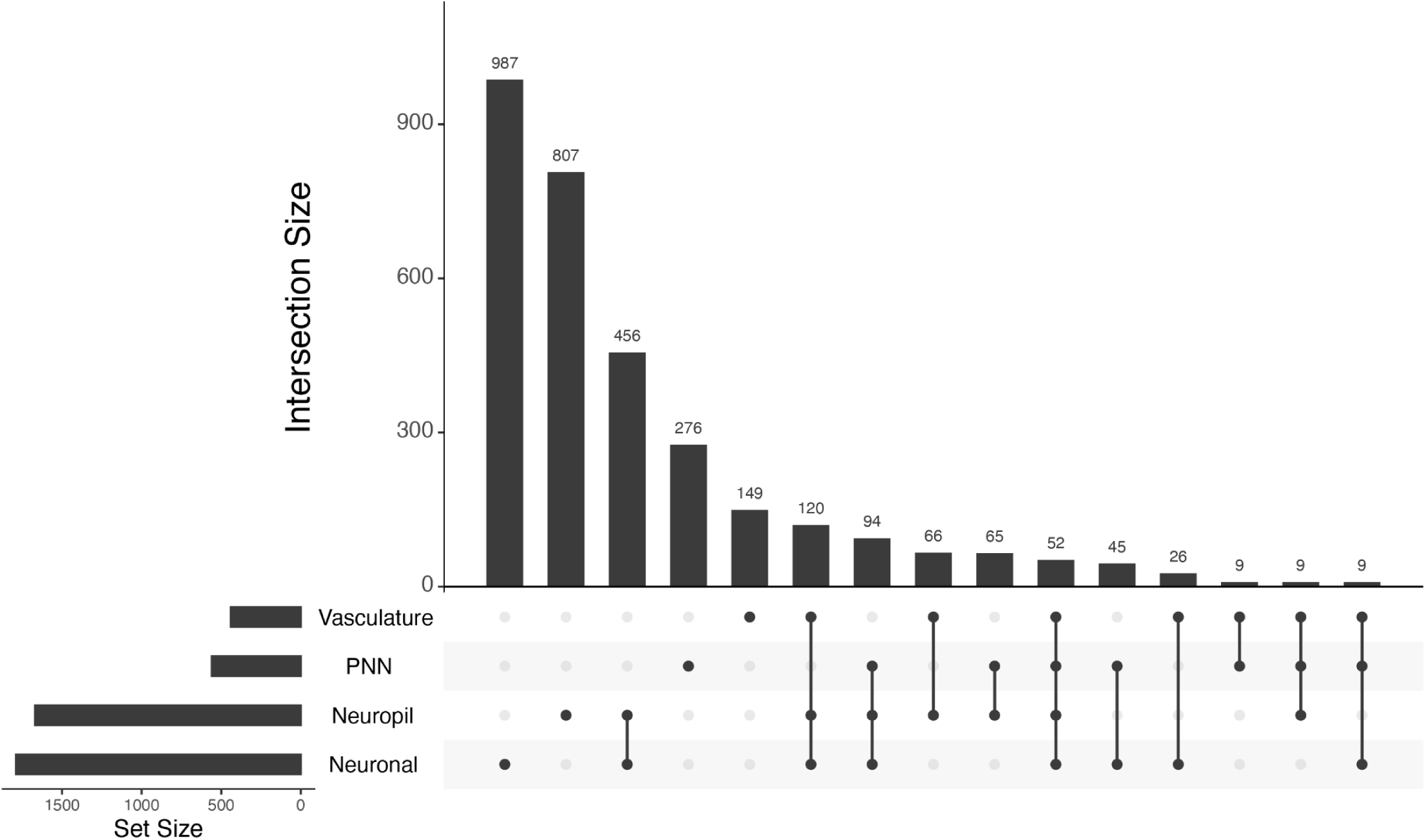
UpSet plot summarizing the intersection of microenvironment-restricted SCZ-DEGs identified within four SPG-defined microenvironments (neuropil, neuronal, PNN, vasculature). The bar plot (top) indicates the number of microenvironment-restricted SCZ-DEGs that are uniquely or jointly identified across microenvironments, ordered by intersection size. Each black dot and connecting line (bottom) specifies the particular combination of microenvironments included in the intersection. Horizontal bars on the left indicate the total number of SCZ-DEGs identified within each microenvironment at nominal *p* < 0.05. All intersections and individual gene lists are provided in **Supplementary Table 10**.

**Figure S37.**
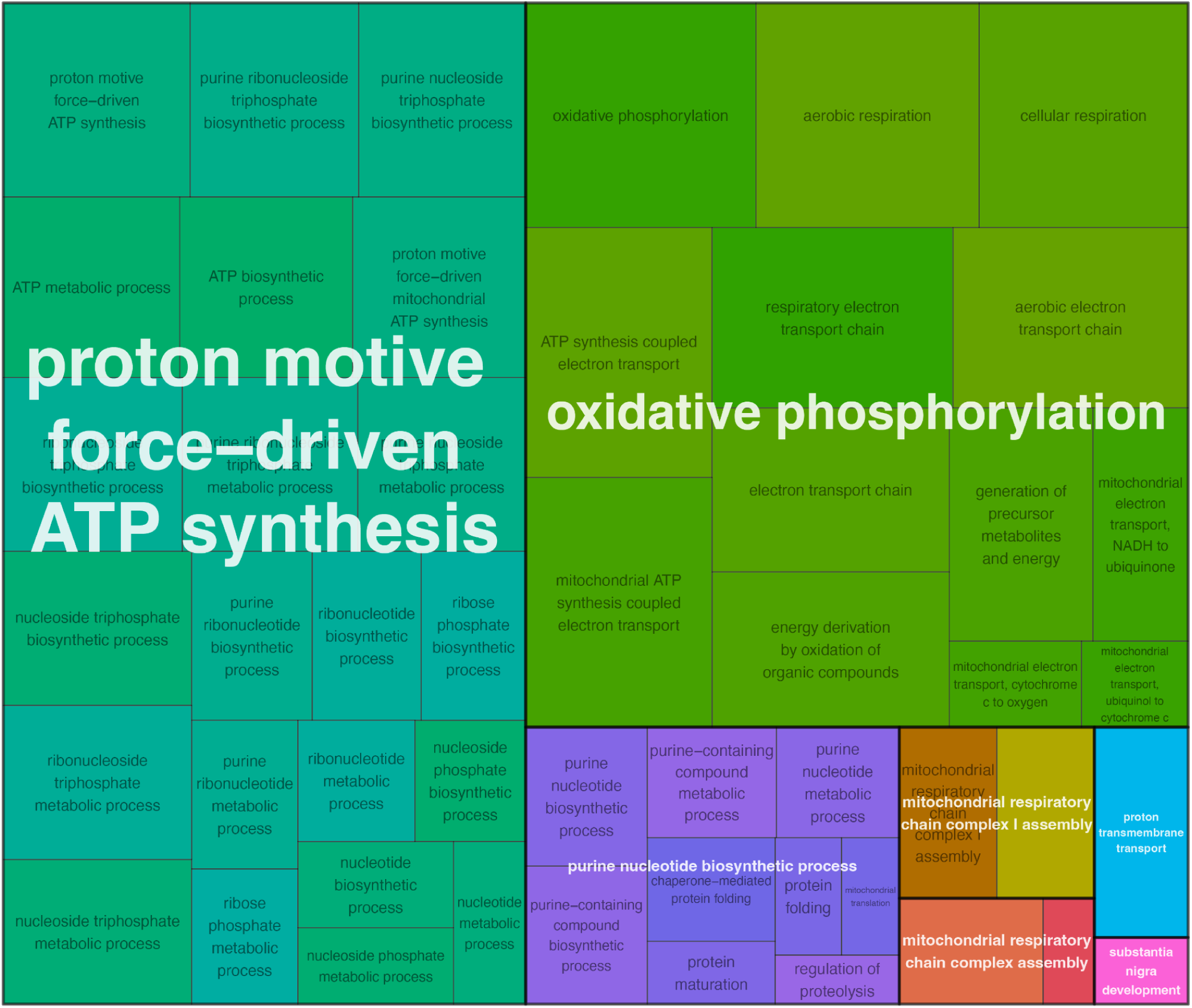
Treemap representation of GO-ORA, showing enriched BP terms among microenvironment-restricted SCZ-DEGs shared across two or more microenvironments. Non-shared microenvironment-restricted SCZ-DEGs identified in only one microenvironment at nominal *p* < 0.05 were excluded prior to this analysis. Each rectangle represents a significantly enriched GO-BP term (FDR ≤ 0.05), with related terms grouped by color and semantic similarity. Rectangle size corresponds to the statistical significance (-log_10_ (adjusted *p*-value)). The treemap displays the top 50 representative terms after redundancy reduction.

**Figure S38.**
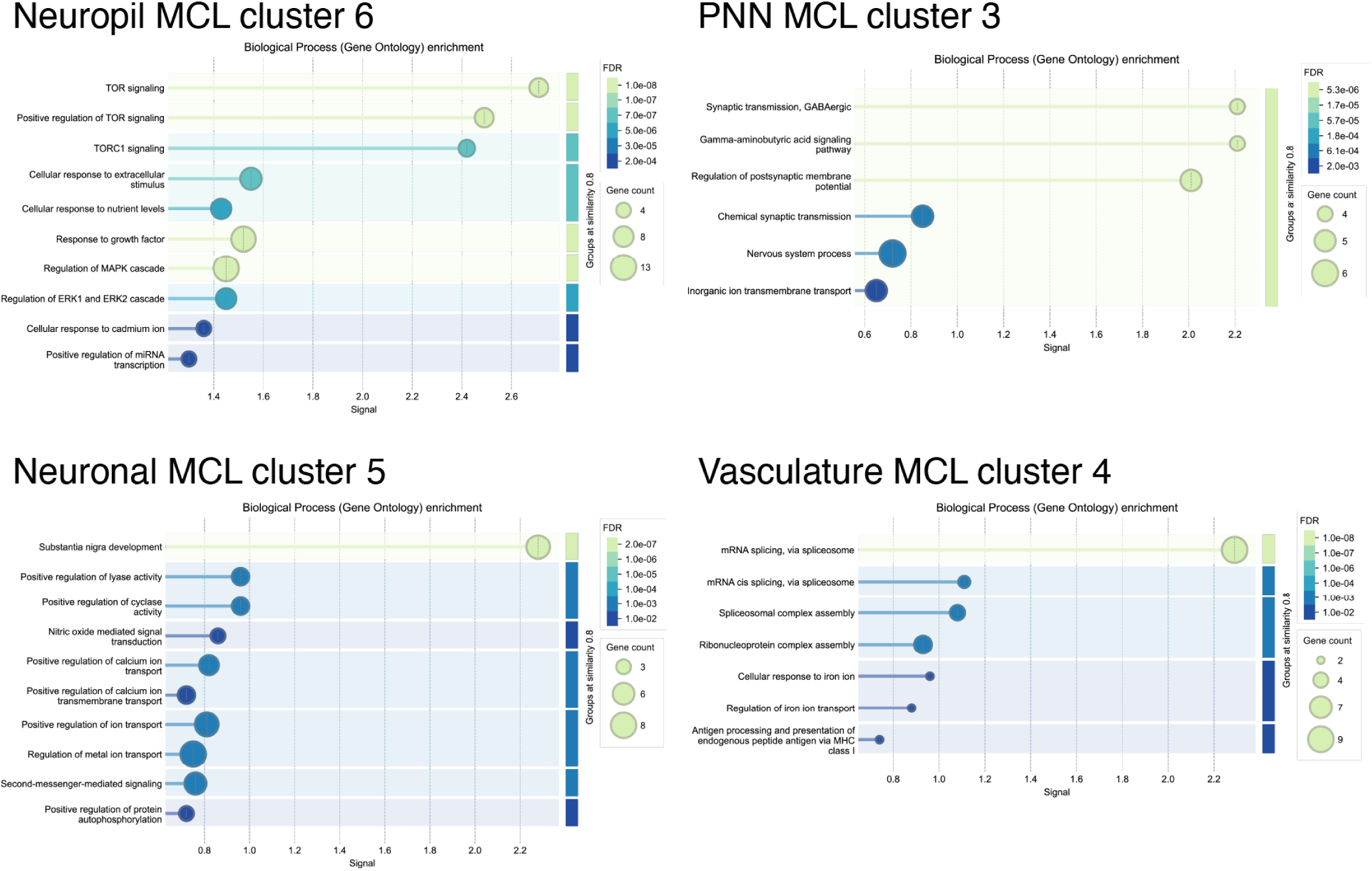
Functional characterization of selected STRING-based protein-protein interaction (PPI) network modules (MCL clusters), using STRING built-in GO-BP enrichment analyses. Each dot plot illustrates significantly enriched GO-BP terms (FDR ≤ 0.05) within a given module, with enrichment signal. Dot size corresponds to the number of module genes associated with each GO term, and dot color indicates the adjusted *p*-value. Enrichment analyses were performed separately for each microenvironment-restricted MCL module using a microenvironment-specific matching background gene set (**Methods**). The results provide additional biological context and functional interpretation of the representative signaling subnetworks highlighted in **Figure 6E**.

**Figure S39.**
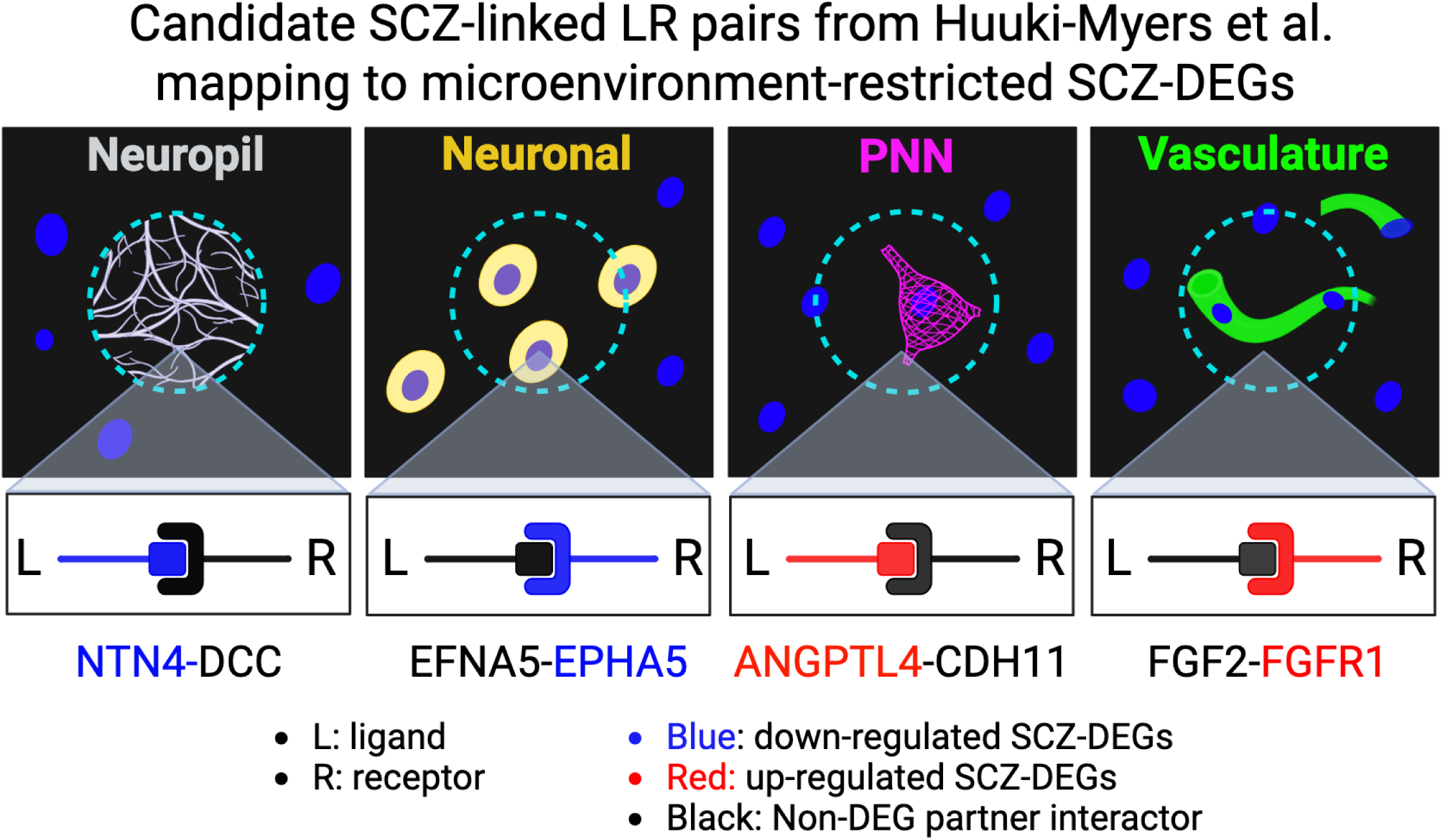
Schematic illustration of the conceptual working model exploring LR pairs altered in SCZ using LR pairs curated from a snRNA-seq study. The diagram provides a conceptual overview of how SCZ-linked candidate LR pairs from Huuki-Myers et al.^25^ were mapped onto microenvironment-restricted SCZ-DEGs identified from our Visium-SPG data. For each microenvironment (neuropil, neuronal, PNN, and vasculature), one representative LR pair is highlighted, where the gene encoding the ligand or the receptor was detected as a statistically significant microenvironment-restricted SCZ-DEG (red: up-regulation; blue: down-regulation of gene expression in SCZ). A cyan dashed circle represents a single Visium spot of the SPG-defined microenvironments. Created in BioRender. Kwon, S. H. (2026) https://BioRender.com/r7yopwt

**Figure S40.**
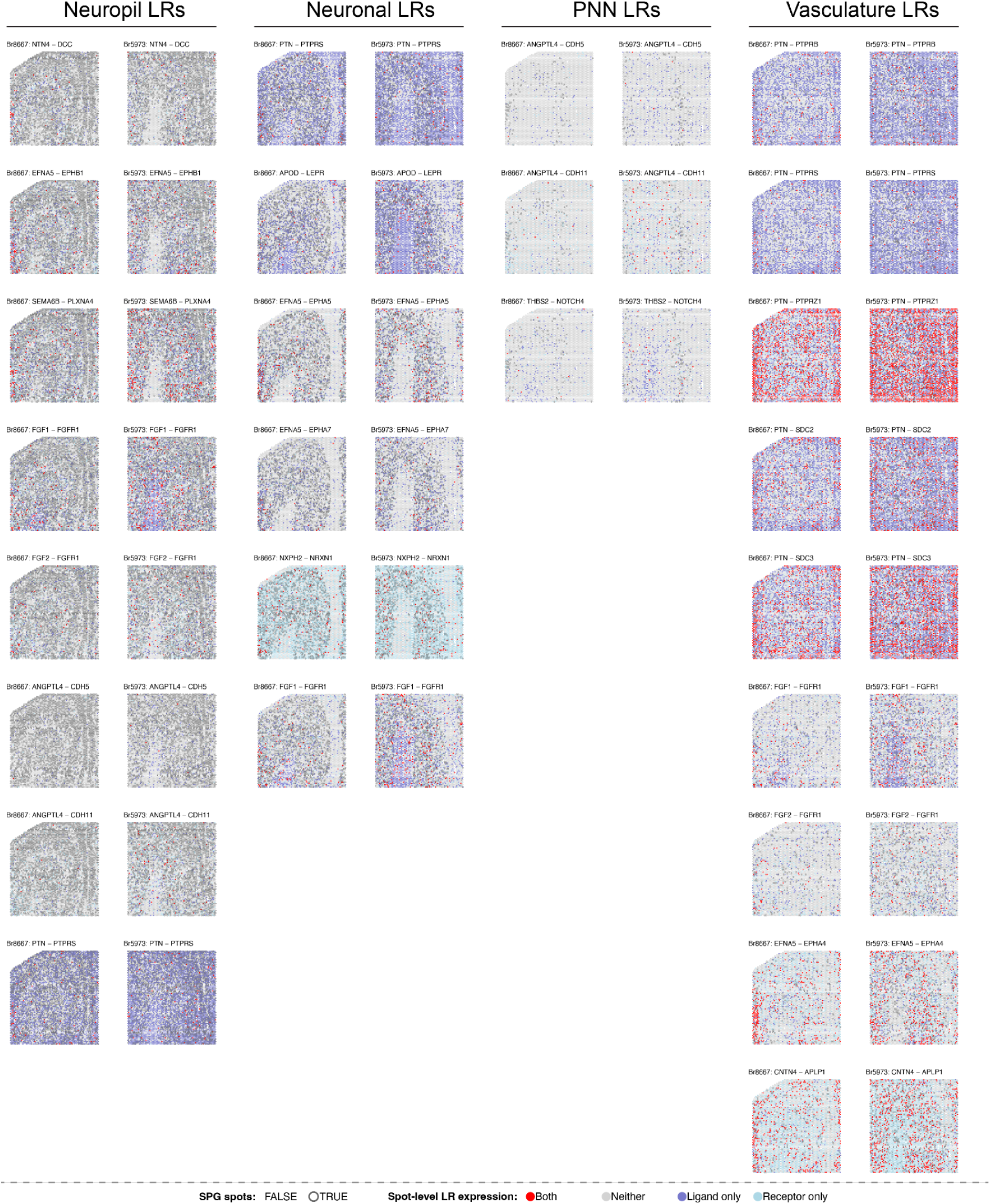
Spot plots of SCZ-linked candidate LR pairs mapping to microenvironment-restricted SCZ-DEGs across SPG-defined microenvironments. Representative LR pairs linked to microenvironment-restricted SCZ-DEGs are shown across four SPG-defined microenvironments (Br8667_NTC and Br5973_SCZ). Visium spots co-expressing both components of an LR pair are marked in red; ligand-only spots in purple; receptor-only spots in skyblue; and spots expressing neither of the components in gray. SPG microenvironment spots are outlined in black.

**Figure S41.**
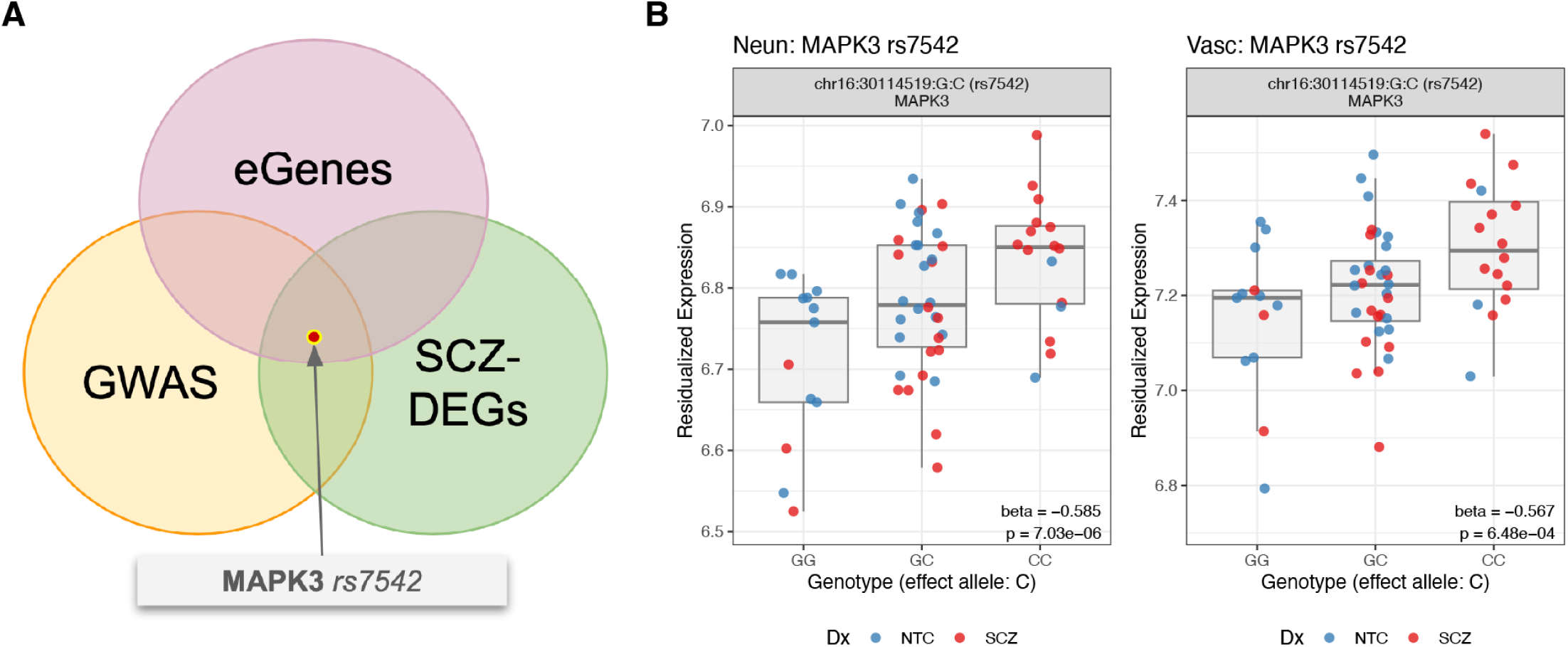
Genotype-associated expression of *MAPK3* across neuronal and vascular microenvironments. (**A**) Venn diagram illustrating the overlap between spatial eQTLs (eGenes), layer-adjusted neuropil SCZ-associated DEGs (SCZ-DEGs), and SCZ GWAS loci (GWAS). We highlight *MAPK3* as a candidate gene supported by convergent genetic association, genotype-dependent regulation, and disease-associated expression changes. (**B**) Box plots show genotype-associated expression patterns of *MAPK3* (rs7542) across neuronal (Neun, left) and vascular (Vasc, right) microenvironments. Consistent genotype-associated expression patterns were observed across neuronal and vascular microenvironments following post hoc correction of *p*-values (FDR < 0.05), suggesting that the genotype-dependent regulatory effect observed in neuropil may extend to additional microenvironments. Diagnosis (Dx) groups are colored: blue for NTC and red for SCZ.

**Figure S42.**
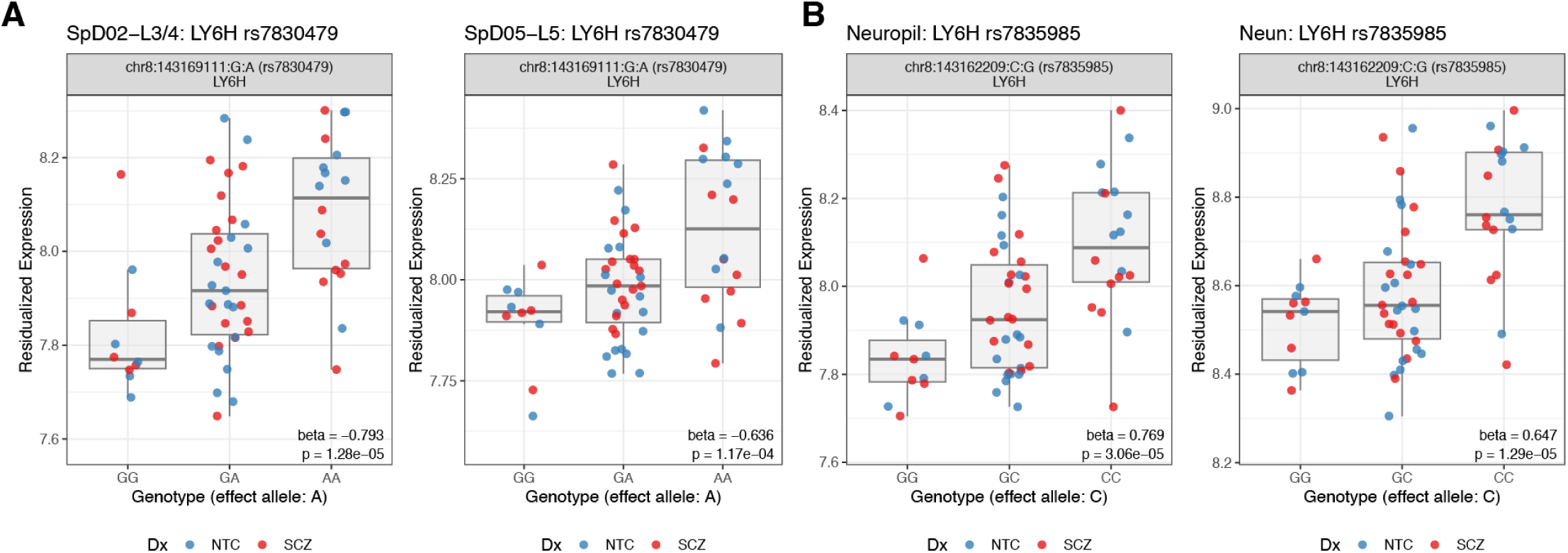
Genotype-associated expression of *LY6H* across dlPFC layers and microenvironments. Box plots show genotype-associated expression of *LY6H* highlighting two strong colocalization signals: (**A**) rs7830479 across selective dlPFC layers: SpD02-L3/4 (left) and SpD05-L5 (right) and (**B**) microenvironments: neuropil (left) and neuronal (right). Diagnosis (Dx) groups are colored: blue for NTC and red for SCZ.

**Figure S43.**
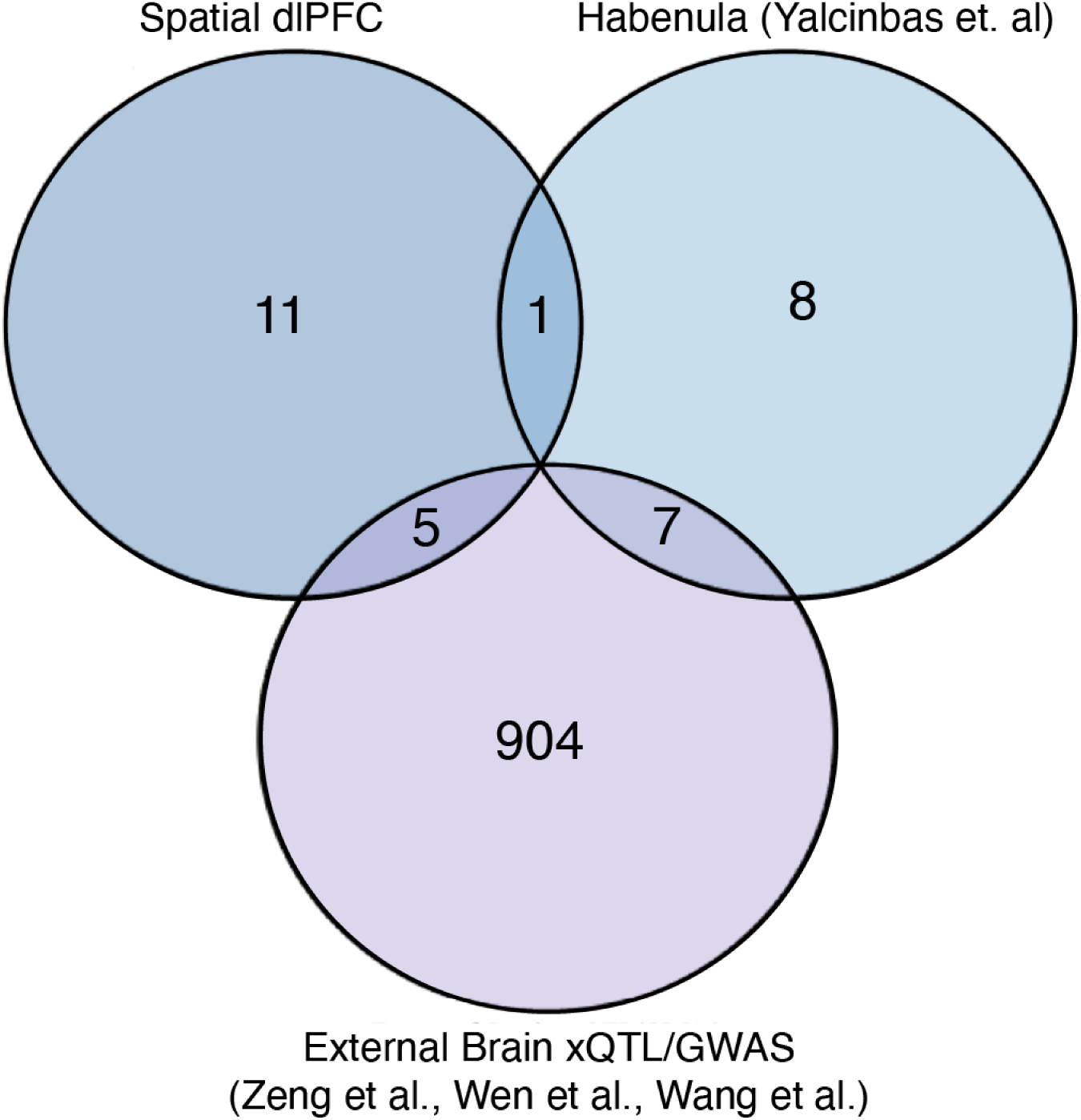
Comparison of dlPFC SCZ colocalization genes with previously published datasets. Venn diagram showing the overlap between SCZ colocalization genes identified in the current spatial dlPFC analysis, habenula eQTL data,^125^ and external brain xQTL/GWAS-based gene-prioritization resources^130–132^. 11 colocalization genes were uniquely detected in the spatial dlPFC analysis, highlighting signals not captured by previous datasets. These 11 genes are listed in **Supplementary Table 13** (coloc_genes_unique_vs_external sheet).

## Supplementary Tables

**Supplementary Table 1**. Demographic information and descriptive statistics of donors included in the Visium and Xenium SRT experiments (sex, age, RNA integrity number [RIN], and postmortem interval[PMI]).

**Supplementary Table 2**. Pairwise DE test results comparing each of the PRECAST SpDs at clustering resolution *k*=7.

**Supplementary Table 3**. DE test results between SPG spots and their matched non-SPG spots to identify highly expressed genes in neuropil, neuronal, vasculature, and PNN microenvironments within NTC samples.

**Supplementary Table 4**. Layer-adjusted DE statistics for all genes, adjusted for PRECAST SpDs at *k*=7.

**Supplementary Table 5**. Layer-restricted DE statistics for all genes, derived from comparisons of matched PRECAST SpDs between NTC and SCZ groups at *k*=7. Layer-restricted DEGs not shared across SpDs are considered layer-specific DEGs.

**Supplementary Table 6**. Predicted TFs across SpDs and directions (up- and down-regulation) of layer-restricted SCZ-DEGs, based on ChEA3 integrated mean-rank results.

**Supplementary Table 7**. PRS-based DE statistics adjusted for PRECAST SpDs at *k*=7.

**Supplementary Table 8**. List of 300 genes included in the Xenium *in situ* panel, with annotations describing their panel role and representative references where applicable.

**Supplementary Table 9**. DE statistics for the 300-gene Xenium panel, evaluated under three modeling approaches in case-control comparisons.

**Supplementary Table 10**. Microenvironment-restricted DE statistics across four SPG-defined microenvironments (neuropil, neuronal, PNN, vasculature), with an intersection summary of shared DEGs.

**Supplementary Table 11**. STRING-MCL cluster descriptions for microenvironment-restricted SCZ-DEGs, with additional GO-BP annotations for the top five MCL clusters.

**Supplementary Table 12**. Context-stratified cis-eQTL results. This Excel workbook contains worksheets **cis_independent**, **nominal_BH05**, and **eQTL summary** with eQTL results derived from `tensorQTL` and augmented with study-specific annotations. In the first 2 worksheets, rows are pooled across 11 expression contexts (column *expression_context*): spd01-spd07 and the SPG-derived microenvironment pseudobulks neun, vasc, pnn, and neuropil. Effect-size and allele-frequency fields were harmonized to the `variant_id` reference/alternate allele orientation defined in the genotype `.pvar` file. The slope direction is interpretable relative to the alleles encoded as REF:ALT in `variant_id`. Columns inherited directly from native tensorQTL outputs are defined in the official tensorQTL output documentation: https://github.com/broadinstitute/tensorqtl/blob/master/docs/outputs.md.

*The **cis_independent*** sheet contains significant cis-eQTLs from the `tensorQTL map_cis` and `map_independent` outputs. This table includes lead cis-eQTLs with FDR < 0.05 (`qval`) from tensorQTL `map_cis` output, and independent signals from `map_independent` for genes whose parent `map_cis` association satisfied `qval` < 0.05, further requiring `pval_perm` < 0.05 for the independent signal itself. One row is retained per *expression_context-ENSEMBL-variant_id* combination. If the same gene-variant pair appears in both source outputs, it is collapsed to a single row and labeled as a lead signal (`map_cis`) rather than `map_independent`-only.

- *eQTL_source* column indicates whether the row comes from `map_cis` (lead cis-eQTL) or `map_independent` (independent-only secondary signal);
- *gene_name* is the matched HGNC-style gene symbol for ENSEMBL;
- *rsid* is the dbSNP rsID from the harmonized GWAS-variant lookup and may be missing for some variants;
- *risk_SNP_GWAS* is a binary flag equal to 1 if the variant is genome-wide significant in the harmonized European-ancestry SCZ GWAS reference (P < 5e-8) and 0 otherwise;
- *SCZ_DEG* is a binary flag set to 1 if the gene is differentially expressed in SCZ versus control at FDR <= 0.1 in the matched context-level DEG resource and 0 otherwise;
- *SCZ_DEG_p05* is a binary flag set to 1 if the gene is differentially expressed at nominal P < 0.05 in the matched DEG resource and 0 otherwise;
- *strong_coloc_variant* is a binary flag set to 1 if the exact context-variant pair appears among high-confidence colocalization results and 0 otherwise;
- *strong_coloc_gene* is a binary flag equal to 1 if the exact context-gene pair appears among high-confidence colocalization results and 0 otherwise;
- *map_cis_qval* is a study-added carry-forward of the parent `map_cis` *q*-value so that independent signals can be interpreted relative to the corresponding gene-level cis significance.

In this merged sheet, the native `tensorQTL` column `qval` is populated for `map_cis` rows and is NA for `map_independent`-only rows.

“Strong” colocalization refers to the study’s high-confidence `coloc` category in downstream summaries, operationally based on high PP4 support (PP4 ≥ 0.8).

The **nominal_BH05** table contains a selection of nominal eQTL associations derived from nominal `tensorQTL` outputs after Benjamini-Hochberg correction of the nominal *p*-values. This table is restricted to significant gene-variant pairs (FDR < 0.05) within each expression context. In this worksheet, columns *expression_context*, *gene_name*, *rsid*, *risk_SNP_GWAS*, *SCZ_DEG_layer_adjusted*, *SCZ_DEG_layer_adjusted_p05*, *strong_coloc_variant*, and *strong_coloc_gene* have the same meanings as above. The column *fdr* is the study-added FDR-adjusted p-value calculated from nominal *pval_nominal* values within each context. All remaining columns in this table are retained directly from the filtered nominal `tensorQTL` results and should be interpreted using the official `tensorQTL` documentation.

**eQTL summary** worksheet provides a compact per-context summary of the significant eQTL results reported in the *cis_independent* and *nominal_BH05* worksheets. Each row corresponds to one *expression_context* and *eQTL_type combination*, where *eQTL_type* distinguishes lead cis-eQTLs (`map_cis`), conditionally independent cis-eQTLs (`map_independent`), and significant nominal eQTLs (`map_nominal`). *eQTL_count* reports the number of unique gene-variant pairs in that group, *n_genes* reports the number of unique genes, *n_SCD_DEG* and *n_SCZ_DEG_p05* report the numbers of unique genes carrying the corresponding SCZ differential-expression annotations, *n_strong_coloc_var* and *n_risk_GWAS* report the corresponding numbers of unique annotated variants, and *n_strong_coloc_gene* reports the number of unique genes annotated by strong colocalization. Thus, gene-based columns count unique genes, variant-based columns count unique variants, and *eQTL_count* counts unique gene-variant pairs.

**Intersecting_genes** worksheet provides a summary of intersecting genes across eGenes, SCZ-DEGs, and GWAS.

**Supplementary Table 13**. Colocalization between spatial dlPFC eQTLs and SCZ GWAS data. Results of colocalization analysis integrating dlPFC eQTL signals from SpD and SPG contexts with SCZ GWAS summary statistics. This workbook contains one worksheet, ***coloc_pass***, and reports context-gene-variant colocalization results retained for downstream interpretation. For each signal, the table includes the tested expression context, gene and lead variant identifiers, the number of SNPs evaluated (*nsnps*), posterior probabilities from `coloc` (*PP3*, *PP4*), study-derived posterior summaries (*PP34*, *PP4_over_PP34*), and a study-specific *hypothesis_category*. The table also includes annotations indicating whether the lead variant is a SCZ GWAS risk in European ancestry (*risk_SNP_GWAS*), whether the gene was found in differential expression results for that context (*SCZ_DEG_layer_adjusted*, FDR < 0.1; *SCZ_DEG_layer_adjusted_p05*, nominal P < 0.05), and whether the same lead variant is supported in the corresponding independent (*eQTL_independent*) or nominal (*eQTL_nominal*) eQTL results for the same context.

- *PP3* and *PP4* are native posterior probabilities from `coloc::coloc.abf`, whereas *PP34* is defined here as PP3 + PP4, and *PP4_over_PP34* as PP4 / (PP3 + PP4).
- *hypothesis_category* is a study-specific summary label derived from these posterior quantities. Native `coloc` posterior outputs can be interpreted with reference to the official `coloc` documentation and vignette: https://chr1swallace.github.io/coloc/reference/coloc-package.html and https://chr1swallace.github.io/coloc/articles/vignette.html.

This workbook additionally includes two supplementary worksheets summarizing colocalized genes in comparison with external datasets: *coloc_genes_unique_vs_Hb* (comparison with habenula eQTL data; see Figure 7I) and *coloc_genes_unique_vs_external* (genes uniquely identified relative to external xQTL/GWAS-based resources; see **Figure S43**).

**Supplementary Table 14**. Exposure times for each fluorescence channel across six different band filters used during Visium imaging with the Vectra Polaris scanner.

## Glossary

AV: Anatomical validation
BA: Brodmann area
BANKSY: Building aggregates with a neighborhood kernel and spatial yardstick
CCC: Cell-cell communication
CDF: Cumulative distributions functions
CGE: Caudal ganglionic eminence
ChEA3: ChIP-X enrichment analysis version 3
CPM: Counts per million
DAPI: 4′,6-diamidino-2-phenylindole
DE: Differential expression
DEG: Differentially expressed gene
dlPFC: Dorsolateral prefrontal cortex
Dx: Diagnosis
ECM: Extracellular matrix
eQTL: Expression quantitative trait locus
FC: Fold change
FDR: False discovery rate
GO: Gene ontology
GO-BP: Gene ontology - biological process
GO-CC: Gene ontology - cellular component
GO-ORA: Gene ontology - overrepresentation analysis
GSEA: Gene set enrichment analysis
GWAS: Genome-wide association study
H&E: Hematoxylin & Eosin
HVG: Highly variable genes
HWE: Hardy-Weinberg equilibrium
IF: Immunofluorescence
L: Cortical layer
LR: Ligand-receptor
M: Meninge
MAD: Median absolute deviation
MAF: Minor allele frequency
MAGMA: Multi-marker analysis of genomic annotation
MCL: Markov clustering
MGE: Median ganglionic eminence
NTC: Neurotypical control
OCT: Optimal cutting temperature compound
OPC: Oligodendrocyte progenitor cell
OR: Odds ratio
PNN: Perineuronal net
PC: Pericyte
PC: Principal component
PCA: Principal component analysis
PGC: Psychiatric genomics consortium
PPI: Protein-protein interaction
PRECAST: Probabilistic embedding, clustering, and alignment for integrating spatial transcriptomics
PRS: Polygenic risk score
PRS-DEG: PRS-associated differentially expressed gene
PVALB: Parvalbumin
QC: Quality control
ROI: Region of interest
scDRS: Single cell disease relevance score
SCZ: Schizophrenia
SCZ-DEG: Schizophrenia-associated differentially expressed gene
SMC: Smooth muscle cell
SMR: Summary data based mendelian randomization
snRNA-seq: SIngle-nucleus RNA-sequencing
SpD: Spatial domain
SPG: Spatial proteogenomics
SRT: Spatially resolved transcriptomics
STRING: Search Tool for the Retrieval of Interacting Genes/Proteins
TF: Transcription factor
TMM: Trimmed mean of M-values
TSS: Transcription start site
UMI: Unique molecular identifier
VLMC: Vascular leptomeningeal cell
WFA: Wisteria floribunda agglutinin
WM: White matter
WMtz: White matter transition zone
2D: 2-dimensional

## References

1. Saha, S., Chant, D., Welham, J., and McGrath, J. (2005). A systematic review of the prevalence of schizophrenia. PLoS Med. 2, e141. 10.1371/journal.pmed.0020141.

2. Moreno-Küstner, B., Martín, C., and Pastor, L. (2018). Prevalence of psychotic disorders and its association with methodological issues. A systematic review and meta-analyses. PLoS ONE 13, e0195687. 10.1371/journal.pone.0195687.

3. Velligan, D.I., and Rao, S. (2023). The epidemiology and global burden of schizophrenia. J. Clin. Psychiatry 84. 10.4088/JCP.MS21078COM5.

4. Nakamura, T., and Takata, A. (2023). The molecular pathology of schizophrenia: an overview of existing knowledge and new directions for future research. Mol. Psychiatry 28, 1868–1889. 10.1038/s41380-023-02005-2.

5. Smucny, J., Dienel, S.J., Lewis, D.A., and Carter, C.S. (2022). Mechanisms underlying dorsolateral prefrontal cortex contributions to cognitive dysfunction in schizophrenia. Neuropsychopharmacology 47, 292–308. 10.1038/s41386-021-01089-0.

6. Howes, O.D., Bukala, B.R., and Beck, K. (2024). Schizophrenia: from neurochemistry to circuits, symptoms and treatments. Nat. Rev. Neurol. 20, 22–35. 10.1038/s41582-023-00904-0.

7. McCutcheon, R.A., Keefe, R.S.E., and McGuire, P.K. (2023). Cognitive impairment in schizophrenia: aetiology, pathophysiology, and treatment. Mol. Psychiatry 28, 1902–1918. 10.1038/s41380-023-01949-9.

8. Glausier, J.R., and Lewis, D.A. (2018). Mapping pathologic circuitry in schizophrenia. Handb. Clin. Neurol. 150, 389–417. 10.1016/B978-0-444-63639-3.00025-6.

9. Semendeferi, K., Lu, A., Schenker, N., and Damasio, H. (2002). Humans and great apes share a large frontal cortex. Nat. Neurosci. 5, 272–276. 10.1038/nn814.

10. Friedman, N.P., and Robbins, T.W. (2022). The role of prefrontal cortex in cognitive control and executive function. Neuropsychopharmacology 47, 72–89. 10.1038/s41386-021-01132-0.

11. Onitsuka, T., Hirano, Y., Nakazawa, T., Ichihashi, K., Miura, K., Inada, K., Mitoma, R., Yasui-Furukori, N., and Hashimoto, R. (2022). Toward recovery in schizophrenia: Current concepts, findings, and future research directions. Psychiatry Clin. Neurosci. 76, 282–291. 10.1111/pcn.13342.

12. Owen, M.J., Bray, N.J., Walters, J.T.R., and O’Donovan, M.C. (2025). Genomics of schizophrenia, bipolar disorder and major depressive disorder. Nat. Rev. Genet. 26, 862–877. 10.1038/s41576-025-00843-0.

13. Trubetskoy, V., Pardiñas, A.F., Qi, T., Panagiotaropoulou, G., Awasthi, S., Bigdeli, T.B., Bryois, J., Chen, C.-Y., Dennison, C.A., Hall, L.S., et al. (2022). Mapping genomic loci implicates genes and synaptic biology in schizophrenia. Nature 604, 502–508. 10.1038/s41586-022-04434-5.

14. Duncan, L.E., Li, T., Salem, M., Li, W., Mortazavi, L., Senturk, H., Shahverdizadeh, N., Vesuna, S., Shen, H., Yoon, J., et al. (2025). Mapping the cellular etiology of schizophrenia and complex brain phenotypes. Nat. Neurosci. 28, 248–258. 10.1038/s41593-024-01834-w.

15. Ling, E., Nemesh, J., Goldman, M., Kamitaki, N., Reed, N., Handsaker, R.E., Genovese, G., Vogelgsang, J.S., Gerges, S., Kashin, S., et al. (2024). A concerted neuron-astrocyte program declines in ageing and schizophrenia. Nature 627, 604–611. 10.1038/s41586-024-07109-5.

16. Ruzicka, W.B., Mohammadi, S., Fullard, J.F., Davila-Velderrain, J., Subburaju, S., Tso, D.R., Hourihan, M., Jiang, S., Lee, H.-C., Bendl, J., et al. (2024). Single-cell multi-cohort dissection of the schizophrenia transcriptome. Science 384, eadg5136. 10.1126/science.adg5136.

17. Gandal, M.J., Zhang, P., Hadjimichael, E., Walker, R.L., Chen, C., Liu, S., Won, H., van Bakel, H., Varghese, M., Wang, Y., et al. (2018). Transcriptome-wide isoform-level dysregulation in ASD, schizophrenia, and bipolar disorder. Science 362. 10.1126/science.aat8127.

18. Batiuk, M.Y., Tyler, T., Dragicevic, K., Mei, S., Rydbirk, R., Petukhov, V., Deviatiiarov, R., Sedmak, D., Frank, E., Feher, V., et al. (2022). Upper cortical layer-driven network impairment in schizophrenia. Sci. Adv. 8, eabn8367. 10.1126/sciadv.abn8367.

19. Wingert, J.C., and Sorg, B.A. (2021). Impact of perineuronal nets on electrophysiology of parvalbumin interneurons, principal neurons, and brain oscillations: A review. Front. Synaptic Neurosci. 13, 673210. 10.3389/fnsyn.2021.673210.

20. Bitanihirwe, B.K.Y., and Woo, T.-U.W. (2014). Perineuronal nets and schizophrenia: the importance of neuronal coatings. Neurosci. Biobehav. Rev. 45, 85–99. 10.1016/j.neubiorev.2014.03.018.

21. Brückner, G., Szeöke, S., Pavlica, S., Grosche, J., and Kacza, J. (2006). Axon initial segment ensheathed by extracellular matrix in perineuronal nets. Neuroscience 138, 365–375. 10.1016/j.neuroscience.2005.11.068.

22. Najjar, S., Pahlajani, S., De Sanctis, V., Stern, J.N.H., Najjar, A., and Chong, D. (2017). Neurovascular Unit Dysfunction and Blood-Brain Barrier Hyperpermeability Contribute to Schizophrenia Neurobiology: A Theoretical Integration of Clinical and Experimental Evidence. Front. Psychiatry 8, 83. 10.3389/fpsyt.2017.00083.

23. Ermakov, E.A., Melamud, M.M., Buneva, V.N., and Ivanova, S.A. (2022). Immune system abnormalities in schizophrenia: an integrative view and translational perspectives. Front. Psychiatry 13, 880568. 10.3389/fpsyt.2022.880568.

24. Liu, W., Liao, X., Luo, Z., Yang, Y., Lau, M.C., Jiao, Y., Shi, X., Zhai, W., Ji, H., Yeong, J., et al. (2023). Probabilistic embedding, clustering, and alignment for integrating spatial transcriptomics data with PRECAST. Nat. Commun. 14, 296. 10.1038/s41467-023-35947-w.

25. Huuki-Myers, L.A., Spangler, A., Eagles, N.J., Montgomery, K.D., Kwon, S.H., Guo, B., Grant-Peters, M., Divecha, H.R., Tippani, M., Sriworarat, C., et al. (2024). A data-driven single-cell and spatial transcriptomic map of the human prefrontal cortex. Science 384, eadh1938. 10.1126/science.adh1938.

26. Pardo, B., Spangler, A., Weber, L.M., Page, S.C., Hicks, S.C., Jaffe, A.E., Martinowich, K., Maynard, K.R., and Collado-Torres, L. (2022). spatialLIBD: an R/Bioconductor package to visualize spatially-resolved transcriptomics data. BMC Genomics 23, 434. 10.1186/s12864-022-08601-w.

27. Maynard, K.R., Collado-Torres, L., Weber, L.M., Uytingco, C., Barry, B.K., Williams, S.R., Catallini, J.L., Tran, M.N., Besich, Z., Tippani, M., et al. (2021). Transcriptome-scale spatial gene expression in the human dorsolateral prefrontal cortex. Nat. Neurosci. 24, 425–436. 10.1038/s41593-020-00787-0.

28. Shah, K., Totty, M.S., Bach, S.V., Valentine, M.R., Chandra, A., Divecha, H.R., Miller, R.A., Kwon, S.H., Ramnauth, A.D., Tippani, M., et al. (2025). Spatio-molecular gene expression reflects dorsal anterior cingulate cortex structure and function in the human brain. BioRxiv. 10.1101/2025.07.14.664821.

29. Kwon, S.H., Parthiban, S., Tippani, M., Divecha, H.R., Eagles, N.J., Lobana, J.S., Williams, S.R., Mak, M., Bharadwaj, R.A., Kleinman, J.E., et al. (2023). Influence of Alzheimer’s disease related neuropathology on local microenvironment gene expression in the human inferior temporal cortex. GEN Biotechnology 2, 399–417. 10.1089/genbio.2023.0019.

30. Lupori, L., Totaro, V., Cornuti, S., Ciampi, L., Carrara, F., Grilli, E., Viglione, A., Tozzi, F., Putignano, E., Mazziotti, R., et al. (2023). A comprehensive atlas of perineuronal net distribution and colocalization with parvalbumin in the adult mouse brain. Cell Rep. 42, 112788. 10.1016/j.celrep.2023.112788.

31. Weller, R.O., Sharp, M.M., Christodoulides, M., Carare, R.O., and Møllgård, K. (2018). The meninges as barriers and facilitators for the movement of fluid, cells and pathogens related to the rodent and human CNS. Acta Neuropathol. 135, 363–385. 10.1007/s00401-018-1809-z.

32. DeFelipe, J., and Fariñas, I. (1992). The pyramidal neuron of the cerebral cortex: morphological and chemical characteristics of the synaptic inputs. Prog. Neurobiol. 39, 563–607. 10.1016/0301-0082(92)90015-7.

33. Elston, G.N. (2003). Cortex, cognition and the cell: new insights into the pyramidal neuron and prefrontal function. Cereb. Cortex 13, 1124–1138. 10.1093/cercor/bhg093.

34. Sherwood, C.C., Stimpson, C.D., Raghanti, M.A., Wildman, D.E., Uddin, M., Grossman, L.I., Goodman, M., Redmond, J.C., Bonar, C.J., Erwin, J.M., et al. (2006). Evolution of increased glia-neuron ratios in the human frontal cortex. Proc Natl Acad Sci USA 103, 13606–13611. 10.1073/pnas.0605843103.

35. Allen, N.J., and Barres, B.A. (2009). Neuroscience: Glia - more than just brain glue. Nature 457, 675–677. 10.1038/457675a.

36. Oohashi, T., Edamatsu, M., Bekku, Y., and Carulli, D. (2015). The hyaluronan and proteoglycan link proteins: Organizers of the brain extracellular matrix and key molecules for neuronal function and plasticity. Exp. Neurol. 274, 134–144. 10.1016/j.expneurol.2015.09.010.

37. Domínguez, S., Rey, C.C., Therreau, L., Fanton, A., Massotte, D., Verret, L., Piskorowski, R.A., and Chevaleyre, V. (2019). Maturation of PNN and ErbB4 Signaling in Area CA2 during Adolescence Underlies the Emergence of PV Interneuron Plasticity and Social Memory. Cell Rep. 29, 1099–1112.e4. 10.1016/j.celrep.2019.09.044.

38. Beroun, A., Mitra, S., Michaluk, P., Pijet, B., Stefaniuk, M., and Kaczmarek, L. (2019). MMPs in learning and memory and neuropsychiatric disorders. Cell. Mol. Life Sci. 76, 3207–3228. 10.1007/s00018-019-03180-8.

39. Niu, M., Cao, W., Wang, Y., Zhu, Q., Luo, J., Wang, B., Zheng, H., Weitz, D.A., and Zong, C. (2023). Droplet-based transcriptome profiling of individual synapses. Nat. Biotechnol. 41, 1332–1344. 10.1038/s41587-022-01635-1.

40. Garcia, F.J., Sun, N., Lee, H., Godlewski, B., Mathys, H., Galani, K., Zhou, B., Jiang, X., Ng, A.P., Mantero, J., et al. (2022). Single-cell dissection of the human brain vasculature. Nature 603, 893–899. 10.1038/s41586-022-04521-7.

41. Collado-Torres, L., Burke, E.E., Peterson, A., Shin, J., Straub, R.E., Rajpurohit, A., Semick, S.A., Ulrich, W.S., BrainSeq Consortium, Price, A.J., et al. (2019). Regional Heterogeneity in Gene Expression, Regulation, and Coherence in the Frontal Cortex and Hippocampus across Development and Schizophrenia. Neuron 103, 203–216.e8. 10.1016/j.neuron.2019.05.013.

42. Gusev, A., Mancuso, N., Won, H., Kousi, M., Finucane, H.K., Reshef, Y., Song, L., Safi, A., Schizophrenia Working Group of the Psychiatric Genomics Consortium, McCarroll, S., et al. (2018). Transcriptome-wide association study of schizophrenia and chromatin activity yields mechanistic disease insights. Nat. Genet. 50, 538–548. 10.1038/s41588-018-0092-1.

43. Liu, J., Cheng, Y., Li, M., Zhang, Z., Li, T., and Luo, X.-J. (2023). Genome-wide Mendelian randomization identifies actionable novel drug targets for psychiatric disorders. Neuropsychopharmacology 48, 270–280. 10.1038/s41386-022-01456-5.

44. Lewis, D.A., and González-Burgos, G. (2008). Neuroplasticity of neocortical circuits in schizophrenia. Neuropsychopharmacology 33, 141–165. 10.1038/sj.npp.1301563.

45. Paz, R.D., Tardito, S., Atzori, M., and Tseng, K.Y. (2008). Glutamatergic dysfunction in schizophrenia: from basic neuroscience to clinical psychopharmacology. Eur. Neuropsychopharmacol. 18, 773–786. 10.1016/j.euroneuro.2008.06.005.

46. Snijders, G.J.L.J., van Zuiden, W., Sneeboer, M.A.M., Berdenis van Berlekom, A., van der Geest, A.T., Schnieder, T., MacIntyre, D.J., Hol, E.M., Kahn, R.S., and de Witte, L.D. (2021). A loss of mature microglial markers without immune activation in schizophrenia. Glia 69, 1251–1267. 10.1002/glia.23962.

47. Gonzalez-Burgos, G., and Lewis, D.A. (2012). NMDA receptor hypofunction, parvalbumin-positive neurons, and cortical gamma oscillations in schizophrenia. Schizophr. Bull. 38, 950–957. 10.1093/schbul/sbs010.

48. Uhlhaas, P.J., and Singer, W. (2010). Abnormal neural oscillations and synchrony in schizophrenia. Nat. Rev. Neurosci. 11, 100–113. 10.1038/nrn2774.

49. Burket, J.A., Webb, J.D., and Deutsch, S.I. (2021). Perineuronal Nets and Metal Cation Concentrations in the Microenvironments of Fast-Spiking, Parvalbumin-Expressing GABAergic Interneurons: Relevance to Neurodevelopment and Neurodevelopmental Disorders. Biomolecules 11. 10.3390/biom11081235.

50. Kim, H.G., Kim, J.Y., Han, E.H., Hwang, Y.P., Choi, J.H., Park, B.H., and Jeong, H.G. (2011). Metallothionein-2A overexpression increases the expression of matrix metalloproteinase-9 and invasion of breast cancer cells. FEBS Lett. 585, 421–428. 10.1016/j.febslet.2010.12.030.

51. Cuenod, M., Steullet, P., Cabungcal, J.-H., Dwir, D., Khadimallah, I., Klauser, P., Conus, P., and Do, K.Q. (2022). Caught in vicious circles: a perspective on dynamic feed-forward loops driving oxidative stress in schizophrenia. Mol. Psychiatry 27, 1886–1897. 10.1038/s41380-021-01374-w.

52. Yılmaz, S., Kılıç, N., Kaya, Ş., and Taşcı, G. (2023). A Potential Biomarker for Predicting Schizophrenia: Metallothionein-1. Biomedicines 11. 10.3390/biomedicines11020590.

53. Skene, N.G., Bryois, J., Bakken, T.E., Breen, G., Crowley, J.J., Gaspar, H.A., Giusti-Rodriguez, P., Hodge, R.D., Miller, J.A., Muñoz-Manchado, A.B., et al. (2018). Genetic identification of brain cell types underlying schizophrenia. Nat. Genet. 50, 825–833. 10.1038/s41588-018-0129-5.

54. Mirnics, K., Middleton, F.A., Lewis, D.A., and Levitt, P. (2001). Analysis of complex brain disorders with gene expression microarrays: schizophrenia as a disease of the synapse. Trends Neurosci. 24, 479–486. 10.1016/s0166-2236(00)01862-2.

55. Howes, O.D., and Onwordi, E.C. (2023). The synaptic hypothesis of schizophrenia version III: a master mechanism. Mol. Psychiatry 28, 1843–1856. 10.1038/s41380-023-02043-w.

56. Koopmans, F., van Nierop, P., Andres-Alonso, M., Byrnes, A., Cijsouw, T., Coba, M.P., Cornelisse, L.N., Farrell, R.J., Goldschmidt, H.L., Howrigan, D.P., et al. (2019). SynGO: An Evidence-Based, Expert-Curated Knowledge Base for the Synapse. Neuron 103, 217–234.e4. 10.1016/j.neuron.2019.05.002.

57. Boulanger, L.M. (2009). Immune proteins in brain development and synaptic plasticity. Neuron 64, 93–109. 10.1016/j.neuron.2009.09.001.

58. Presumey, J., Bialas, A.R., and Carroll, M.C. (2017). Complement system in neural synapse elimination in development and disease. Adv. Immunol. 135, 53–79. 10.1016/bs.ai.2017.06.004.

59. Selden, N.R., Gitelman, D.R., Salamon-Murayama, N., Parrish, T.B., and Mesulam, M.M. (1998). Trajectories of cholinergic pathways within the cerebral hemispheres of the human brain. Brain 121 (Pt 12), 2249–2257. 10.1093/brain/121.12.2249.

60. Fields, R.D., Dutta, D.J., Belgrad, J., and Robnett, M. (2017). Cholinergic signaling in myelination. Glia 65, 687–698. 10.1002/glia.23101.

61. Foster, D.J., Bryant, Z.K., and Conn, P.J. (2021). Targeting muscarinic receptors to treat schizophrenia. Behav. Brain Res. 405, 113201. 10.1016/j.bbr.2021.113201.

62. Paramo, B., Wyatt, S., and Davies, A.M. (2018). An essential role for neuregulin-4 in the growth and elaboration of developing neocortical pyramidal dendrites. Exp. Neurol. 302, 85–92. 10.1016/j.expneurol.2018.01.002.

63. Ledonne, A., and Mercuri, N.B. (2019). On the modulatory roles of neuregulins/erbb signaling on synaptic plasticity. Int. J. Mol. Sci. 21. 10.3390/ijms21010275.

64. Mei, L., and Nave, K.-A. (2014). Neuregulin-ERBB signaling in the nervous system and neuropsychiatric diseases. Neuron 83, 27–49. 10.1016/j.neuron.2014.06.007.

65. Dietz, A.G., Goldman, S.A., and Nedergaard, M. (2020). Glial cells in schizophrenia: a unified hypothesis. Lancet Psychiatry 7, 272–281. 10.1016/S2215-0366(19)30302-5.

66. Gerstner, N., Fröhlich, A.S., Matosin, N., Gagliardi, M., Cruceanu, C., Ködel, M., Rex-Haffner, M., Tu, X., Mostafavi, S., Ziller, M.J., et al. (2025). Contrasting genetic predisposition and diagnosis in psychiatric disorders: A multi-omic single-nucleus analysis of the human OFC. Sci. Adv. 11, eadq2290. 10.1126/sciadv.adq2290.

67. Notter, T. (2021). Astrocytes in schizophrenia. Brain Neurosci. Adv. 5, 23982128211009148. 10.1177/23982128211009148.

68. Sorg, B.A., Berretta, S., Blacktop, J.M., Fawcett, J.W., Kitagawa, H., Kwok, J.C.F., and Miquel, M. (2016). Casting a wide net: role of perineuronal nets in neural plasticity. J. Neurosci. 36, 11459–11468. 10.1523/JNEUROSCI.2351-16.2016.

69. Enwright, J.F., Sanapala, S., Foglio, A., Berry, R., Fish, K.N., and Lewis, D.A. (2016). Reduced Labeling of Parvalbumin Neurons and Perineuronal Nets in the Dorsolateral Prefrontal Cortex of Subjects with Schizophrenia. Neuropsychopharmacology 41, 2206–2214. 10.1038/npp.2016.24.

70. Tewari, B.P., Woo, A.M., Prim, C.E., Chaunsali, L., Patel, D.C., Kimbrough, I.F., Engel, K., Browning, J.L., Campbell, S.L., and Sontheimer, H. (2024). Astrocytes require perineuronal nets to maintain synaptic homeostasis in mice. Nat. Neurosci. 27, 1475–1488. 10.1038/s41593-024-01714-3.

71. Suárez-Solá, M.L., González-Delgado, F.J., Pueyo-Morlans, M., Medina-Bolívar, O.C., Hernández-Acosta, N.C., González-Gómez, M., and Meyer, G. (2009). Neurons in the white matter of the adult human neocortex. Front. Neuroanat. 3, 7. 10.3389/neuro.05.007.2009.

72. Connor, C.M., Crawford, B.C., and Akbarian, S. (2011). White matter neuron alterations in schizophrenia and related disorders. Int. J. Dev. Neurosci. 29, 325–334. 10.1016/j.ijdevneu.2010.07.236.

73. Chartrand, T., Dalley, R., Close, J., Goriounova, N.A., Lee, B.R., Mann, R., Miller, J.A., Molnar, G., Mukora, A., Alfiler, L., et al. (2023). Morphoelectric and transcriptomic divergence of the layer 1 interneuron repertoire in human versus mouse neocortex. Science 382, eadf0805. 10.1126/science.adf0805.

74. Ament, S.A., and Poulopoulos, A. (2023). The brain’s dark transcriptome: Sequencing RNA in distal compartments of neurons and glia. Curr. Opin. Neurobiol. 81, 102725. 10.1016/j.conb.2023.102725.

75. Qian, X., Coleman, K., Jiang, S., Kriz, A.J., Marciano, J.H., Luo, C., Cai, C., Manam, M.D., Caglayan, E., Lai, A., et al. (2025). Spatial transcriptomics reveals human cortical layer and area specification. Nature 644, 153–163. 10.1038/s41586-025-09010-1.

76. Keenan, A.B., Torre, D., Lachmann, A., Leong, A.K., Wojciechowicz, M.L., Utti, V., Jagodnik, K.M., Kropiwnicki, E., Wang, Z., and Ma’ayan, A. (2019). ChEA3: transcription factor enrichment analysis by orthogonal omics integration. Nucleic Acids Res. 47, W212–W224. 10.1093/nar/gkz446.

77. Szklarczyk, D., Gable, A.L., Lyon, D., Junge, A., Wyder, S., Huerta-Cepas, J., Simonovic, M., Doncheva, N.T., Morris, J.H., Bork, P., et al. (2019). STRING v11: protein-protein association networks with increased coverage, supporting functional discovery in genome-wide experimental datasets. Nucleic Acids Res. 47, D607–D613. 10.1093/nar/gky1131.

78. Tao, R., Cousijn, H., Jaffe, A.E., Burnet, P.W.J., Edwards, F., Eastwood, S.L., Shin, J.H., Lane, T.A., Walker, M.A., Maher, B.J., et al. (2014). Expression of ZNF804A in human brain and alterations in schizophrenia, bipolar disorder, and major depressive disorder: a novel transcript fetally regulated by the psychosis risk variant rs1344706. JAMA Psychiatry 71, 1112–1120. 10.1001/jamapsychiatry.2014.1079.

79. Chang, H., Xiao, X., and Li, M. (2017). The schizophrenia risk gene ZNF804A: clinical associations, biological mechanisms and neuronal functions. Mol. Psychiatry 22, 944–953. 10.1038/mp.2017.19.

80. O’Donovan, M.C., Craddock, N., Norton, N., Williams, H., Peirce, T., Moskvina, V., Nikolov, I., Hamshere, M., Carroll, L., Georgieva, L., et al. (2008). Identification of loci associated with schizophrenia by genome-wide association and follow-up. Nat. Genet. 40, 1053–1055. 10.1038/ng.201.

81. Ge, T., Chen, C.-Y., Ni, Y., Feng, Y.-C.A., and Smoller, J.W. (2019). Polygenic prediction via Bayesian regression and continuous shrinkage priors. Nat. Commun. 10, 1776. 10.1038/s41467-019-09718-5.

82. Zhang, M.J., Hou, K., Dey, K.K., Sakaue, S., Jagadeesh, K.A., Weinand, K., Taychameekiatchai, A., Rao, P., Pisco, A.O., Zou, J., et al. (2022). Polygenic enrichment distinguishes disease associations of individual cells in single-cell RNA-seq data. Nat. Genet. 54, 1572–1580. 10.1038/s41588-022-01167-z.

83. de Leeuw, C.A., Mooij, J.M., Heskes, T., and Posthuma, D. (2015). MAGMA: generalized gene-set analysis of GWAS data. PLoS Comput. Biol. 11, e1004219. 10.1371/journal.pcbi.1004219.

84. Millet, A., Ledo, J.H., and Tavazoie, S.F. (2024). An exhausted-like microglial population accumulates in aged and APOE4 genotype Alzheimer’s brains. Immunity 57, 153–170.e6. 10.1016/j.immuni.2023.12.001.

85. Thrupp, N., Sala Frigerio, C., Wolfs, L., Skene, N.G., Fattorelli, N., Poovathingal, S., Fourne, Y., Matthews, P.M., Theys, T., Mancuso, R., et al. (2020). Single-Nucleus RNA-Seq Is Not Suitable for Detection of Microglial Activation Genes in Humans. Cell Rep. 32, 108189. 10.1016/j.celrep.2020.108189.

86. Koskuvi, M., Pörsti, E., Hewitt, T., Räsänen, N., Wu, Y.-C., Trontti, K., McQuade, A., Kalyanaraman, S., Ojansuu, I., Vaurio, O., et al. (2024). Genetic contribution to microglial activation in schizophrenia. Mol. Psychiatry 29, 2622–2633. 10.1038/s41380-024-02529-1.

87. Puvogel, S., Alsema, A., Kracht, L., Webster, M.J., Weickert, C.S., Sommer, I.E.C., and Eggen, B.J.L. (2022). Single-nucleus RNA sequencing of midbrain blood-brain barrier cells in schizophrenia reveals subtle transcriptional changes with overall preservation of cellular proportions and phenotypes. Mol. Psychiatry 27, 4731–4740. 10.1038/s41380-022-01796-0.

88. Cai, W., Song, W., Liu, Z., Maharjan, D.T., Liang, J., and Lin, G.N. (2022). An Integrative Analysis of Identified Schizophrenia-Associated Brain Cell Types and Gene Expression Changes. Int. J. Mol. Sci. 23. 10.3390/ijms231911581.

89. Gober, R., Ardalan, M., Shiadeh, S.M.J., Duque, L., Garamszegi, S.P., Ascona, M., Barreda, A., Sun, X., Mallard, C., and Vontell, R.T. (2022). Microglia activation in postmortem brains with schizophrenia demonstrates distinct morphological changes between brain regions. Brain Pathol. 32, e13003. 10.1111/bpa.13003.

90. Bast, L., Yao, S., Martínez-López, J.A., Memic, F., French, H., Valiukonyte, M., Karlsson, R., Wen, J., Song, J., Zhang, R., et al. (2025). Transcriptomic and genetic analysis suggests a role for mitochondrial dysregulation in schizophrenia. medRxiv. 10.1101/2025.03.14.25323827.

91. Dienel, S.J., Fish, K.N., and Lewis, D.A. (2023). The nature of prefrontal cortical GABA neuron alterations in schizophrenia: markedly lower somatostatin and parvalbumin gene expression without missing neurons. Am. J. Psychiatry 180, 495–507. 10.1176/appi.ajp.20220676.

92. Fang, C., Montgomery, K.D., Maguire, S.E., Ramnauth, A.D., Guo, B., Miller, R., Kleinmann, J.E., Hyde, T.M., Martinowich, K., Maynard, K.R., et al. (2025). spaTransfer: transfer learning for single-cell and spatial transcriptomics data using non-negative matrix factorization. BioRxiv. 10.64898/2025.12.12.694021.

93. Singhal, V., Chou, N., Lee, J., Yue, Y., Liu, J., Chock, W.K., Lin, L., Chang, Y.-C., Teo, E.M.L., Aow, J., et al. (2024). BANKSY unifies cell typing and tissue domain segmentation for scalable spatial omics data analysis. Nat. Genet. 56, 431–441. 10.1038/s41588-024-01664-3.

94. Rudy, B., Fishell, G., Lee, S., and Hjerling-Leffler, J. (2011). Three groups of interneurons account for nearly 100% of neocortical GABAergic neurons. Dev. Neurobiol. 71, 45–61. 10.1002/dneu.20853.

95. Fishell, G., and Kepecs, A. (2020). Interneuron types as attractors and controllers. Annu. Rev. Neurosci. 43, 1–30. 10.1146/annurev-neuro-070918-050421.

96. Kaar, S.J., Angelescu, I., Marques, T.R., and Howes, O.D. (2019). Pre-frontal parvalbumin interneurons in schizophrenia: a meta-analysis of post-mortem studies. J. Neural Transm. 126, 1637–1651. 10.1007/s00702-019-02080-2.

97. Höistad, M., Segal, D., Takahashi, N., Sakurai, T., Buxbaum, J.D., and Hof, P.R. (2009). Linking white and grey matter in schizophrenia: oligodendrocyte and neuron pathology in the prefrontal cortex. Front. Neuroanat. 3, 9. 10.3389/neuro.05.009.2009.

98. Petukhov, V., Igolkina, A., Rydbirk, R., Mei, S., Christoffersen, L., Khodosevich, K., and Kharchenko, P.V. (2022). Case-control analysis of single-cell RNA-seq studies. BioRxiv. 10.1101/2022.03.15.484475.

99. Zhou, J., Wang, M., and Deng, D. (2020). KLF2 protects BV2 microglial cells against oxygen and glucose deprivation injury by modulating BDNF/TrkB pathway. Gene 735, 144277. 10.1016/j.gene.2019.144277.

100. Arion, D., Corradi, J.P., Tang, S., Datta, D., Boothe, F., He, A., Cacace, A.M., Zaczek, R., Albright, C.F., Tseng, G., et al. (2015). Distinctive transcriptome alterations of prefrontal pyramidal neurons in schizophrenia and schizoaffective disorder. Mol. Psychiatry 20, 1397–1405. 10.1038/mp.2014.171.

101. Mizoguchi, T., Minakuchi, H., Ishisaka, M., Tsuruma, K., Shimazawa, M., and Hara, H. (2017). Behavioral abnormalities with disruption of brain structure in mice overexpressing VGF. Sci. Rep. 7, 4691. 10.1038/s41598-017-04132-7.

102. Anderson, C.L., Gross, A.L., Pettigrew, C., Vazquez, J.P., Moghekar, A., Oh, S., Na, C.H., Albert, M., Worley, P., Soldan, A., et al. (2025). Cognitive resilience in preclinical Alzheimer’s disease: Higher NPTX2 and VGF levels are associated with reduced cognitive decline. Neurobiol. Aging 156, 133–142. 10.1016/j.neurobiolaging.2025.09.002.

103. Irwin, N., Chao, S., Goritchenko, L., Horiuchi, A., Greengard, P., Nairn, A.C., and Benowitz, L.I. (2002). Nerve growth factor controls GAP-43 mRNA stability via the phosphoprotein ARPP-19. Proc Natl Acad Sci USA 99, 12427–12431. 10.1073/pnas.152457399.

104. Xiao, M.-F., Roh, S.-E., Zhou, J., Chien, C.-C., Lucey, B.P., Craig, M.T., Hayes, L.N., Coughlin, J.M., Leweke, F.M., Jia, M., et al. (2021). A biomarker-authenticated model of schizophrenia implicating NPTX2 loss of function. Sci. Adv. 7, eabf6935. 10.1126/sciadv.abf6935.

105. Valdés-Tovar, M., Rodríguez-Ramírez, A.M., Rodríguez-Cárdenas, L., Sotelo-Ramírez, C.E., Camarena, B., Sanabrais-Jiménez, M.A., Solís-Chagoyán, H., Argueta, J., and López-Riquelme, G.O. (2022). Insights into myelin dysfunction in schizophrenia and bipolar disorder. World J. Psychiatry 12, 264–285. 10.5498/wjp.v12.i2.264.

106. Pellegri, G., Magistretti, P.J., and Martin, J.L. (1998). VIP and PACAP potentiate the action of glutamate on BDNF expression in mouse cortical neurones. Eur. J. Neurosci. 10, 272–280. 10.1046/j.1460-9568.1998.00052.x.

107. Egan, M.F., Goldberg, T.E., Kolachana, B.S., Callicott, J.H., Mazzanti, C.M., Straub, R.E., Goldman, D., and Weinberger, D.R. (2001). Effect of COMT Val108/158 Met genotype on frontal lobe function and risk for schizophrenia. Proc Natl Acad Sci USA 98, 6917–6922. 10.1073/pnas.111134598.

108. Blackwood, D.H., Fordyce, A., Walker, M.T., St Clair, D.M., Porteous, D.J., and Muir, W.J. (2001). Schizophrenia and affective disorders--cosegregation with a translocation at chromosome 1q42 that directly disrupts brain-expressed genes: clinical and P300 findings in a family. Am. J. Hum. Genet. 69, 428–433. 10.1086/321969.

109. Onwordi, E.C., Halff, E.F., Whitehurst, T., Mansur, A., Cotel, M.-C., Wells, L., Creeney, H., Bonsall, D., Rogdaki, M., Shatalina, E., et al. (2020). Synaptic density marker SV2A is reduced in schizophrenia patients and unaffected by antipsychotics in rats. Nat. Commun. 11, 246. 10.1038/s41467-019-14122-0.

110. Sekar, A., Bialas, A.R., de Rivera, H., Davis, A., Hammond, T.R., Kamitaki, N., Tooley, K., Presumey, J., Baum, M., Van Doren, V., et al. (2016). Schizophrenia risk from complex variation of complement component 4. Nature 530, 177–183. 10.1038/nature16549.

111. Holt, C.E., and Schuman, E.M. (2013). The central dogma decentralized: new perspectives on RNA function and local translation in neurons. Neuron 80, 648–657. 10.1016/j.neuron.2013.10.036.

112. Abbott, N.J., Rönnbäck, L., and Hansson, E. (2006). Astrocyte-endothelial interactions at the blood-brain barrier. Nat. Rev. Neurosci. 7, 41–53. 10.1038/nrn1824.

113. Castillo, P.E., Jung, H., Klann, E., and Riccio, A. (2023). Presynaptic protein synthesis in brain function and disease. J. Neurosci. 43, 7483–7488. 10.1523/JNEUROSCI.1454-23.2023.

114. Gamarra, M., de la Cruz, A., Blanco-Urrejola, M., and Baleriola, J. (2021). Local translation in nervous system pathologies. Front. Integr. Neurosci. 15, 689208. 10.3389/fnint.2021.689208.

115. Liu-Yesucevitz, L., Bassell, G.J., Gitler, A.D., Hart, A.C., Klann, E., Richter, J.D., Warren, S.T., and Wolozin, B. (2011). Local RNA translation at the synapse and in disease. J. Neurosci. 31, 16086–16093. 10.1523/JNEUROSCI.4105-11.2011.

116. Carceller, H., Guirado, R., Ripolles-Campos, E., Teruel-Marti, V., and Nacher, J. (2020). Perineuronal Nets Regulate the Inhibitory Perisomatic Input onto Parvalbumin Interneurons and γ Activity in the Prefrontal Cortex. J. Neurosci. 40, 5008–5018. 10.1523/JNEUROSCI.0291-20.2020.

117. Leppä, E., Linden, A.-M., Vekovischeva, O.Y., Swinny, J.D., Rantanen, V., Toppila, E., Höger, H., Sieghart, W., Wulff, P., Wisden, W., et al. (2011). Removal of GABA(A) receptor γ2 subunits from parvalbumin neurons causes wide-ranging behavioral alterations. PLoS ONE 6, e24159. 10.1371/journal.pone.0024159.

118. Willis, A., Pratt, J.A., and Morris, B.J. (2021). BDNF and JNK Signaling Modulate Cortical Interneuron and Perineuronal Net Development: Implications for Schizophrenia-Linked 16p11.2 Duplication Syndrome. Schizophr. Bull. 47, 812–826. 10.1093/schbul/sbaa139.

119. Lesnikova, A., Casarotto, P.C., Fred, S.M., Voipio, M., Winkel, F., Steinzeig, A., Antila, H., Umemori, J., Biojone, C., and Castrén, E. (2021). Chondroitinase and Antidepressants Promote Plasticity by Releasing TRKB from Dephosphorylating Control of PTPσ in Parvalbumin Neurons. J. Neurosci. 41, 972–980. 10.1523/JNEUROSCI.2228-20.2020.

120. Ochoa, D., Hercules, A., Carmona, M., Suveges, D., Baker, J., Malangone, C., Lopez, I., Miranda, A., Cruz-Castillo, C., Fumis, L., et al. (2023). The next-generation Open Targets Platform: reimagined, redesigned, rebuilt. Nucleic Acids Res. 51, D1353–D1359. 10.1093/nar/gkac1046.

121. Türei, D., Korcsmáros, T., and Saez-Rodriguez, J. (2016). OmniPath: guidelines and gateway for literature-curated signaling pathway resources. Nat. Methods 13, 966–967. 10.1038/nmeth.4077.

122. Klimaschewski, L., and Claus, P. (2021). Fibroblast growth factor signalling in the diseased nervous system. Mol. Neurobiol. 58, 3884–3902. 10.1007/s12035-021-02367-0.

123. Papadimitriou, E., Mourkogianni, E., Ntenekou, D., Christopoulou, M., Koutsioumpa, M., and Lamprou, M. (2022). On the role of pleiotrophin and its receptors in development and angiogenesis. Int. J. Dev. Biol. 66, 115–124. 10.1387/ijdb.210122ep.

124. Taylor-Weiner, A., Aguet, F., Haradhvala, N.J., Gosai, S., Anand, S., Kim, J., Ardlie, K., Van Allen, E.M., and Getz, G. (2019). Scaling computational genomics to millions of individuals with GPUs. Genome Biol. 20, 228. 10.1186/s13059-019-1836-7.

125. Yalcinbas, E.A., Ajanaku, B., Nelson, E.D., Garcia-Flores, R., Eagles, N.J., Montgomery, K.D., Stolz, J.M., Wu, J., Divecha, H.R., Chandra, A., et al. (2025). Transcriptomic analysis of the human habenula in schizophrenia. Am. J. Psychiatry 182, 991–1006. 10.1176/appi.ajp.20240776.

126. GTEx Consortium (2015). Human genomics. The Genotype-Tissue Expression (GTEx) pilot analysis: multitissue gene regulation in humans. Science 348, 648–660. 10.1126/science.1262110.

127. Moriwaki, Y., Kubo, N., Watanabe, M., Asano, S., Shinoda, T., Sugino, T., Ichikawa, D., Tsuji, S., Kato, F., and Misawa, H. (2020). Endogenous neurotoxin-like protein Ly6H inhibits alpha7 nicotinic acetylcholine receptor currents at the plasma membrane. Sci. Rep. 10, 11996. 10.1038/s41598-020-68947-7.

128. Tregellas, J.R., and Wylie, K.P. (2019). Alpha7 nicotinic receptors as therapeutic targets in schizophrenia. Nicotine Tob. Res. 21, 349–356. 10.1093/ntr/nty034.

129. Wingo, A.P., Liu, Y., Vattathil, S.M., Gerasimov, E.S., Mei, Z., Ravindran, S.P., Liu, J., Shantaraman, A., Seifar, F., Wang, E., et al. (2025). Multiancestry brain pQTL fine-mapping and integration with genome-wide association studies of 21 neurologic and psychiatric conditions. Nat. Genet. 57, 2156–2165. 10.1038/s41588-025-02291-2.

130. Zeng, B., Bendl, J., Kosoy, R., Fullard, J.F., Hoffman, G.E., and Roussos, P. (2022). Multi-ancestry eQTL meta-analysis of human brain identifies candidate causal variants for brain-related traits. Nat. Genet. 54, 161–169. 10.1038/s41588-021-00987-9.

131. Wen, C., Margolis, M., Dai, R., Zhang, P., Przytycki, P.F., Vo, D.D., Bhattacharya, A., Matoba, N., Tang, M., Jiao, C., et al. (2024). Cross-ancestry atlas of gene, isoform, and splicing regulation in the developing human brain. Science 384, eadh0829. 10.1126/science.adh0829.

132. Wang, D., Liu, S., Warrell, J., Won, H., Shi, X., Navarro, F.C.P., Clarke, D., Gu, M., Emani, P., Yang, Y.T., et al. (2018). Comprehensive functional genomic resource and integrative model for the human brain. Science 362. 10.1126/science.aat8464.

133. Huang, S., Wu, S.J., Sansone, G., Ibrahim, L.A., and Fishell, G. (2024). Layer 1 neocortex: Gating and integrating multidimensional signals. Neuron 112, 184–200. 10.1016/j.neuron.2023.09.041.

134. Sedmak, G., and Judaš, M. (2021). White matter interstitial neurons in the adult human brain: 3% of cortical neurons in quest for recognition. Cells 10. 10.3390/cells10010190.

135. Maden, S.K., Kwon, S.H., Huuki-Myers, L.A., Collado-Torres, L., Hicks, S.C., and Maynard, K.R. (2023). Challenges and opportunities to computationally deconvolve heterogeneous tissue with varying cell sizes using single-cell RNA-sequencing datasets. Genome Biol. 24, 288. 10.1186/s13059-023-03123-4.

136. Denisenko, E., Guo, B.B., Jones, M., Hou, R., de Kock, L., Lassmann, T., Poppe, D., Clément, O., Simmons, R.K., Lister, R., et al. (2020). Systematic assessment of tissue dissociation and storage biases in single-cell and single-nucleus RNA-seq workflows. Genome Biol. 21, 130. 10.1186/s13059-020-02048-6.

137. Pitino, E., Pascual-Reguant, A., Segato-Dezem, F., Wise, K., Salvador-Martinez, I., Crowell, H.L., Marção, M., Ruiz, M., Courtois, E., Flynn, W.F., et al. (2025). STAMP: Single-cell transcriptomics analysis and multimodal profiling through imaging. Cell 188, 5100–5117.e26. 10.1016/j.cell.2025.05.027.

138. Baker, A., Kalmbach, B., Morishima, M., Kim, J., Juavinett, A., Li, N., and Dembrow, N. (2018). Specialized Subpopulations of Deep-Layer Pyramidal Neurons in the Neocortex: Bridging Cellular Properties to Functional Consequences. J. Neurosci. 38, 5441–5455. 10.1523/JNEUROSCI.0150-18.2018.

139. Duchatel, R.J., Shannon Weickert, C., and Tooney, P.A. (2019). White matter neuron biology and neuropathology in schizophrenia. NPJ Schizophr. 5, 10. 10.1038/s41537-019-0078-8.

140. Muraki, K., and Tanigaki, K. (2015). Neuronal migration abnormalities and its possible implications for schizophrenia. Front. Neurosci. 9, 74. 10.3389/fnins.2015.00074.

141. Cajigas, I.J., Tushev, G., Will, T.J., tom Dieck, S., Fuerst, N., and Schuman, E.M. (2012). The local transcriptome in the synaptic neuropil revealed by deep sequencing and high-resolution imaging. Neuron 74, 453–466. 10.1016/j.neuron.2012.02.036.

142. Ledderose, J.M.T., Zolnik, T.A., Toumazou, M., Trimbuch, T., Rosenmund, C., Eickholt, B.J., Jaeger, D., Larkum, M.E., and Sachdev, R.N.S. (2023). Layer 1 of somatosensory cortex: an important site for input to a tiny cortical compartment. Cereb. Cortex 33, 11354–11372. 10.1093/cercor/bhad371.

143. Birnbaum, R., and Weinberger, D.R. (2017). Genetic insights into the neurodevelopmental origins of schizophrenia. Nat. Rev. Neurosci. 18, 727–740. 10.1038/nrn.2017.125.

144. Jaffe, A.E., Straub, R.E., Shin, J.H., Tao, R., Gao, Y., Collado-Torres, L., Kam-Thong, T., Xi, H.S., Quan, J., Chen, Q., et al. (2018). Developmental and genetic regulation of the human cortex transcriptome illuminate schizophrenia pathogenesis. Nat. Neurosci. 21, 1117–1125. 10.1038/s41593-018-0197-y.

145. Sawa, A., Pletnikov, M.V., and Kamiya, A. (2004). Neuron-glia interactions clarify genetic-environmental links in mental illness. Trends Neurosci. 27, 294–297. 10.1016/j.tins.2004.03.012.

146. Hartmann, S.-M., Heider, J., Wüst, R., Fallgatter, A.J., and Volkmer, H. (2024). Microglia-neuron interactions in schizophrenia. Front. Cell. Neurosci. 18, 1345349. 10.3389/fncel.2024.1345349.

147. Bernstein, H.-G., Nussbaumer, M., Vasilevska, V., Dobrowolny, H., Nickl-Jockschat, T., Guest, P.C., and Steiner, J. (2025). Glial cell deficits are a key feature of schizophrenia: implications for neuronal circuit maintenance and histological differentiation from classical neurodegeneration. Mol. Psychiatry 30, 1102–1116. 10.1038/s41380-024-02861-6.

148. Huang, Z.J., Kirkwood, A., Pizzorusso, T., Porciatti, V., Morales, B., Bear, M.F., Maffei, L., and Tonegawa, S. (1999). BDNF regulates the maturation of inhibition and the critical period of plasticity in mouse visual cortex. Cell 98, 739–755. 10.1016/s0092-8674(00)81509-3.

149. Zheng, K., An, J.J., Yang, F., Xu, W., Xu, Z.-Q.D., Wu, J., Hökfelt, T.G.M., Fisahn, A., Xu, B., and Lu, B. (2011). TrkB signaling in parvalbumin-positive interneurons is critical for gamma-band network synchronization in hippocampus. Proc Natl Acad Sci USA 108, 17201–17206. 10.1073/pnas.1114241108.

150. Zannas, A.S., Wiechmann, T., Gassen, N.C., and Binder, E.B. (2016). Gene-Stress-Epigenetic Regulation of FKBP5: Clinical and Translational Implications. Neuropsychopharmacology 41, 261–274. 10.1038/npp.2015.235.

151. Matosin, N., Arloth, J., Czamara, D., Edmond, K.Z., Maitra, M., Fröhlich, A.S., Martinelli, S., Kaul, D., Bartlett, R., Curry, A.R., et al. (2023). Associations of psychiatric disease and ageing with FKBP5 expression converge on superficial layer neurons of the neocortex. Acta Neuropathol. 145, 439–459. 10.1007/s00401-023-02541-9.

152. Sinclair, D., Fillman, S.G., Webster, M.J., and Weickert, C.S. (2013). Dysregulation of glucocorticoid receptor co-factors FKBP5, BAG1 and PTGES3 in prefrontal cortex in psychotic illness. Sci. Rep. 3, 3539. 10.1038/srep03539.

153. Arion, D., Enwright, J.F., Gonzalez-Burgos, G., and Lewis, D.A. (2024). Cell Type-Specific Profiles and Developmental Trajectories of Transcriptomes in Primate Prefrontal Layer 3 Pyramidal Neurons: Implications for Schizophrenia. Am. J. Psychiatry 181, 920–934. 10.1176/appi.ajp.20230541.

154. Aryal, S., Bonanno, K., Song, B., Mani, D.R., Keshishian, H., Carr, S.A., Sheng, M., and Dejanovic, B. (2023). Deep proteomics identifies shared molecular pathway alterations in synapses of patients with schizophrenia and bipolar disorder and mouse model. Cell Rep. 42, 112497. 10.1016/j.celrep.2023.112497.

155. Gottschalk, M.G., Wesseling, H., Guest, P.C., and Bahn, S. (2014). Proteomic enrichment analysis of psychotic and affective disorders reveals common signatures in presynaptic glutamatergic signaling and energy metabolism. Int. J. Neuropsychopharmacol. 18. 10.1093/ijnp/pyu019.

156. Salter, M.W., and Stevens, B. (2017). Microglia emerge as central players in brain disease. Nat. Med. 23, 1018–1027. 10.1038/nm.4397.

157. Cheng, M., Jiang, Y., Xu, J., Mentis, A.-F.A., Wang, S., Zheng, H., Sahu, S.K., Liu, L., and Xu, X. (2023). Spatially resolved transcriptomics: a comprehensive review of their technological advances, applications, and challenges. J. Genet. Genomics 50, 625–640. 10.1016/j.jgg.2023.03.011.

158. Williams, C.G., Lee, H.J., Asatsuma, T., Vento-Tormo, R., and Haque, A. (2022). An introduction to spatial transcriptomics for biomedical research. Genome Med. 14, 68. 10.1186/s13073-022-01075-1.

159. Lipska, B.K., Deep-Soboslay, A., Weickert, C.S., Hyde, T.M., Martin, C.E., Herman, M.M., and Kleinman, J.E. (2006). Critical factors in gene expression in postmortem human brain: Focus on studies in schizophrenia. Biol. Psychiatry 60, 650–658. 10.1016/j.biopsych.2006.06.019.

160. Wang, F., Flanagan, J., Su, N., Wang, L.-C., Bui, S., Nielson, A., Wu, X., Vo, H.-T., Ma, X.-J., and Luo, Y. (2012). RNAscope: a novel in situ RNA analysis platform for formalin-fixed, paraffin-embedded tissues. J. Mol. Diagn. 14, 22–29. 10.1016/j.jmoldx.2011.08.002.

161. Benavides, S.H., Monserrat, A.J., Fariña, S., and Porta, E.A. (2002). Sequential histochemical studies of neuronal lipofuscin in human cerebral cortex from the first to the ninth decade of life. Arch. Gerontol. Geriatr. 34, 219–231. 10.1016/S0167-4943(01)00223-0.

162. Tippani, M., Divecha, H.R., Catallini, J.L., Kwon, S.H., Weber, L.M., Spangler, A., Jaffe, A.E., Hyde, T.M., Kleinman, J.E., Hicks, S.C., et al. (2023). VistoSeg: Processing utilities for high-resolution images for spatially resolved transcriptomics data. Biol. Imaging 3, e23. 10.1017/S2633903X23000235.

163. 10x Genomics spaceranger count, 10x Genomics. https://support.10xgenomics.com/spatial-gene-expression/software/pipelines/latest/using/count.

164. Righelli, D., Weber, L.M., Crowell, H.L., Pardo, B., Collado-Torres, L., Ghazanfar, S., Lun, A.T.L., Hicks, S.C., and Risso, D. (2022). SpatialExperiment: infrastructure for spatially-resolved transcriptomics data in R using Bioconductor. Bioinformatics 38, 3128–3131. 10.1093/bioinformatics/btac299.

165. Totty, M., Hicks, S.C., and Guo, B. (2025). SpotSweeper: spatially aware quality control for spatial transcriptomics. Nat. Methods 22, 1520–1530. 10.1038/s41592-025-02713-3.

166. McCarthy, D.J., Campbell, K.R., Lun, A.T.L., and Wills, Q.F. (2017). Scater: pre-processing, quality control, normalization and visualization of single-cell RNA-seq data in R. Bioinformatics 33, 1179–1186. 10.1093/bioinformatics/btw777.

167. Guo, B., Huuki-Myers, L.A., Grant-Peters, M., Collado-Torres, L., and Hicks, S.C. (2023). escheR: unified multi-dimensional visualizations with Gestalt principles. Bioinformatics Advances 3, vbad179. 10.1093/bioadv/vbad179.

168. Virtanen, P., Gommers, R., Oliphant, T.E., Haberland, M., Reddy, T., Cournapeau, D., Burovski, E., Peterson, P., Weckesser, W., Bright, J., et al. (2020). SciPy 1.0: fundamental algorithms for scientific computing in Python. Nat. Methods 17, 261–272. 10.1038/s41592-019-0686-2.

169. Harris, C.R., Millman, K.J., van der Walt, S.J., Gommers, R., Virtanen, P., Cournapeau, D., Wieser, E., Taylor, J., Berg, S., Smith, N.J., et al. (2020). Array programming with NumPy. Nature 585, 357–362. 10.1038/s41586-020-2649-2.

170. Baskar, A., Rajappa, M., Vasudevan, S.K., and Murugesh, T.S. (2023). Playing with OpenCV and Python. In Digital Image Processing (Chapman and Hall/CRC), pp. 177–187. 10.1201/9781003217428-8.

171. van der Walt, S., Schönberger, J.L., Nunez-Iglesias, J., Boulogne, F., Warner, J.D., Yager, N., Gouillart, E., Yu, T., and scikit-image contributors (2014). scikit-image: image processing in Python. PeerJ 2, e453. 10.7717/peerj.453.

172. Coudray, N., Buessler, J.-L., and Urban, J.-P. (2010). Robust threshold estimation for images with unimodal histograms. Pattern Recognit. Lett. 31, 1010–1019. 10.1016/j.patrec.2009.12.025.

173. Ghaye, J., Kamat, M.A., Corbino-Giunta, L., Silacci, P., Vergères, G., De Micheli, G., and Carrara, S. (2013). Image thresholding techniques for localization of sub-resolution fluorescent biomarkers. Cytometry A 83, 1001–1016. 10.1002/cyto.a.22345.

174. Kwon, S.H., Guo, B., Fang, C., Tippani, M., Bach, S.V., Pertea, G., Miller, R.A., Maguire, S.E., Iatrou, A., Eagles, N.J., et al. (2026). LieberInstitute/spatialDLPFC_SCZ: preprint-v1.0.0. Zenodo. 10.5281/zenodo.18663825.

175. Ritchie, M.E., Phipson, B., Wu, D., Hu, Y., Law, C.W., Shi, W., and Smyth, G.K. (2015). limma powers differential expression analyses for RNA-sequencing and microarray studies. Nucleic Acids Res. 43, e47. 10.1093/nar/gkv007.

176. Yu, G., Wang, L.-G., Han, Y., and He, Q.-Y. (2012). clusterProfiler: an R package for comparing biological themes among gene clusters. OMICS 16, 284–287. 10.1089/omi.2011.0118.

177. Carlson, M. (2017). org.Hs.eg.db: Genome wide annotation for Human. R package version 3.20.0. Bioconductor. 10.18129/b9.bioc.org.hs.eg.db.

178. Sayols, S. (2023). rrvgo: a Bioconductor package for interpreting lists of Gene Ontology terms. MicroPubl. Biol. 2023. 10.17912/micropub.biology.000811.

179. Purcell, S., Neale, B., Todd-Brown, K., Thomas, L., Ferreira, M.A.R., Bender, D., Maller, J., Sklar, P., de Bakker, P.I.W., Daly, M.J., et al. (2007). PLINK: a tool set for whole-genome association and population-based linkage analyses. Am. J. Hum. Genet. 81, 559–575. 10.1086/519795.

180. Taliun, D., Harris, D.N., Kessler, M.D., Carlson, J., Szpiech, Z.A., Torres, R., Taliun, S.A.G., Corvelo, A., Gogarten, S.M., Kang, H.M., et al. (2021). Sequencing of 53,831 diverse genomes from the NHLBI TOPMed Program. Nature 590, 290–299. 10.1038/s41586-021-03205-y.

181. Chang, C.C., Chow, C.C., Tellier, L.C., Vattikuti, S., Purcell, S.M., and Lee, J.J. (2015). Second-generation PLINK: rising to the challenge of larger and richer datasets. Gigascience 4, s13742–015–0047–8. 10.1186/s13742-015-0047-8.

182. Xue, A., Yazar, S., Neavin, D., and Powell, J.E. (2023). Pitfalls and opportunities for applying latent variables in single-cell eQTL analyses. Genome Biol. 24, 33. 10.1186/s13059-023-02873-5.

183. Robinson, M.D., McCarthy, D.J., and Smyth, G.K. (2010). edgeR: a Bioconductor package for differential expression analysis of digital gene expression data. Bioinformatics 26, 139–140. 10.1093/bioinformatics/btp616.

184. GTEx Consortium (2020). The GTEx Consortium atlas of genetic regulatory effects across human tissues. Science 369, 1318–1330. 10.1126/science.aaz1776.

185. Leek, J.T., Johnson, W.E., Parker, H.S., Jaffe, A.E., and Storey, J.D. (2012). The sva package for removing batch effects and other unwanted variation in high-throughput experiments. Bioinformatics 28, 882–883. 10.1093/bioinformatics/bts034.

186. Giambartolomei, C., Vukcevic, D., Schadt, E.E., Franke, L., Hingorani, A.D., Wallace, C., and Plagnol, V. (2014). Bayesian test for colocalisation between pairs of genetic association studies using summary statistics. PLoS Genet. 10, e1004383. 10.1371/journal.pgen.1004383.

187. Wallace, C. (2020). Eliciting priors and relaxing the single causal variant assumption in colocalisation analyses. PLoS Genet. 16, e1008720. 10.1371/journal.pgen.1008720.

188. Collado-Torres, L., Jaffe, A., and Burke, E. (2019). LieberInstitute/jaffelab: Zenodo integration. Zenodo. 10.5281/zenodo.3376220.

189. Sriworarat, C., Nguyen, A., Eagles, N.J., Collado-Torres, L., Martinowich, K., Maynard, K.R., and Hicks, S.C. (2023). Performant web-based interactive visualization tool for spatially-resolved transcriptomics experiments. Biol. Imaging 3, e15. 10.1017/S2633903X2300017X.

190. Rue-Albrecht, K., Marini, F., Soneson, C., and Lun, A.T.L. (2018). iSEE: Interactive SummarizedExperiment Explorer. [version 1; peer review: 3 approved]. F1000Res. 7, 741. 10.12688/f1000research.14966.1.

191. Kwon, S.H., Guo, B., Fang, C., Tippani, M., Bach, S.V., Pertea, G., Miller, R.A., Maguire, S.E., Iatrou, A., Eagles, N.J., et al. (2026). LieberInstitute/spatialDLPFC_SCZ_XENIUM: preprint-v1.0.0. Zenodo. 10.5281/zenodo.19135970.

