## Supplementary Figures for "Mapping spatially organized molecular and genetic signatures of schizophrenia across multiple scales in human prefrontal cortex"

### Supplemental Figures

**Figure S1**. **Representative overview of excluded spots**. Spots with library size less than 100 and fewer than 200 unique genes detected, and tissue artifacts (**Figure S3**) were flagged for removal. Representative spot plots showing spots flagged for removal from a subset of the cohort (24/63 samples).

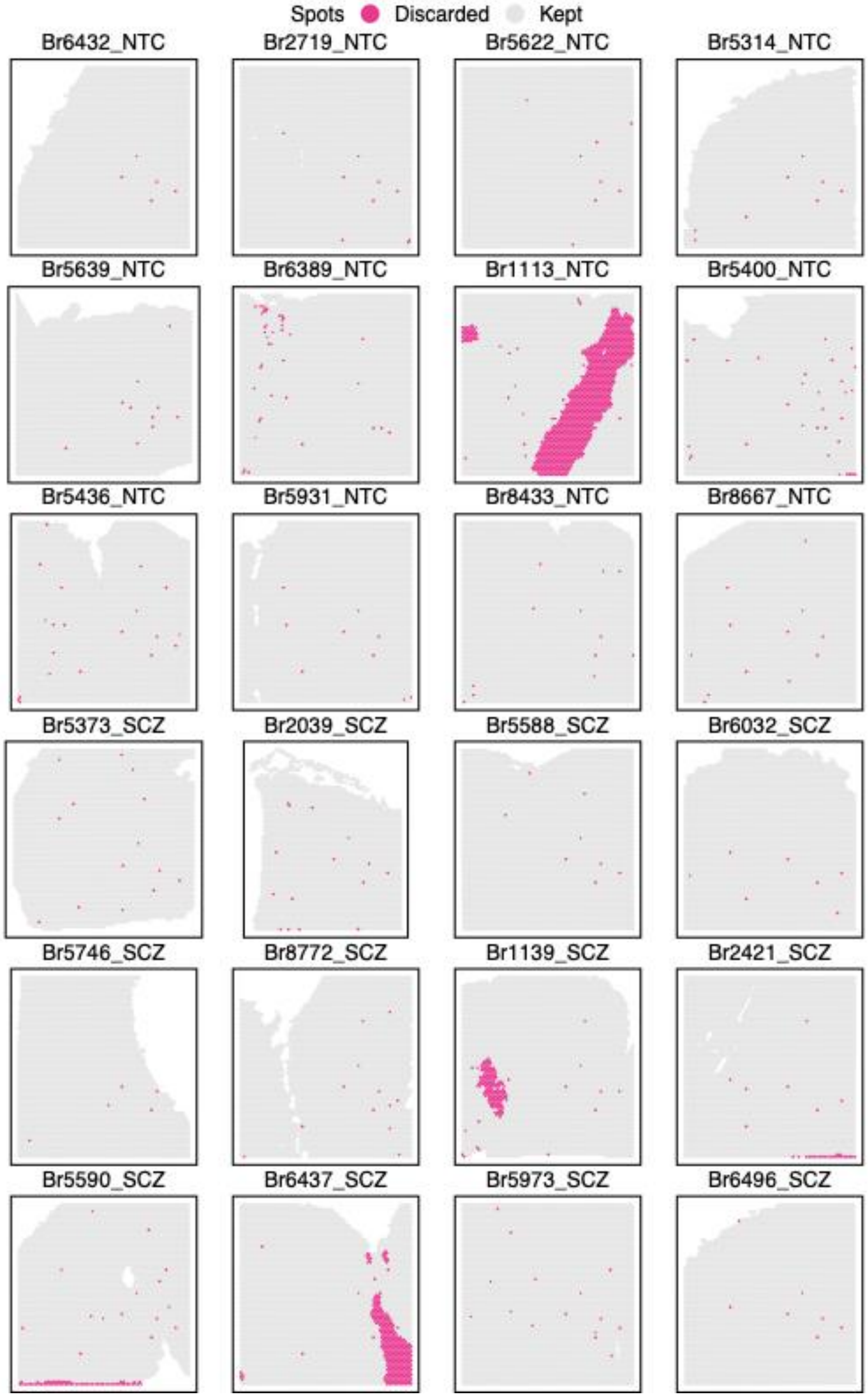

**Figure S2**. **Distributions of QC metrics stratified by diagnosis**. Violin plots showing spot-level distributions of three QC metrics, stratified by diagnosis (NTC and SCZ): total unique molecular identifiers (Total UMIs) also known as library size (**top**), number of detected genes (Unique Genes) (**middle**), and proportion of mitochondrial genes (Mito. Ratio) (**bottom**). Dotted lines represent thresholds used to identify low-quality spots (UMIs < 100 and genes < 200).

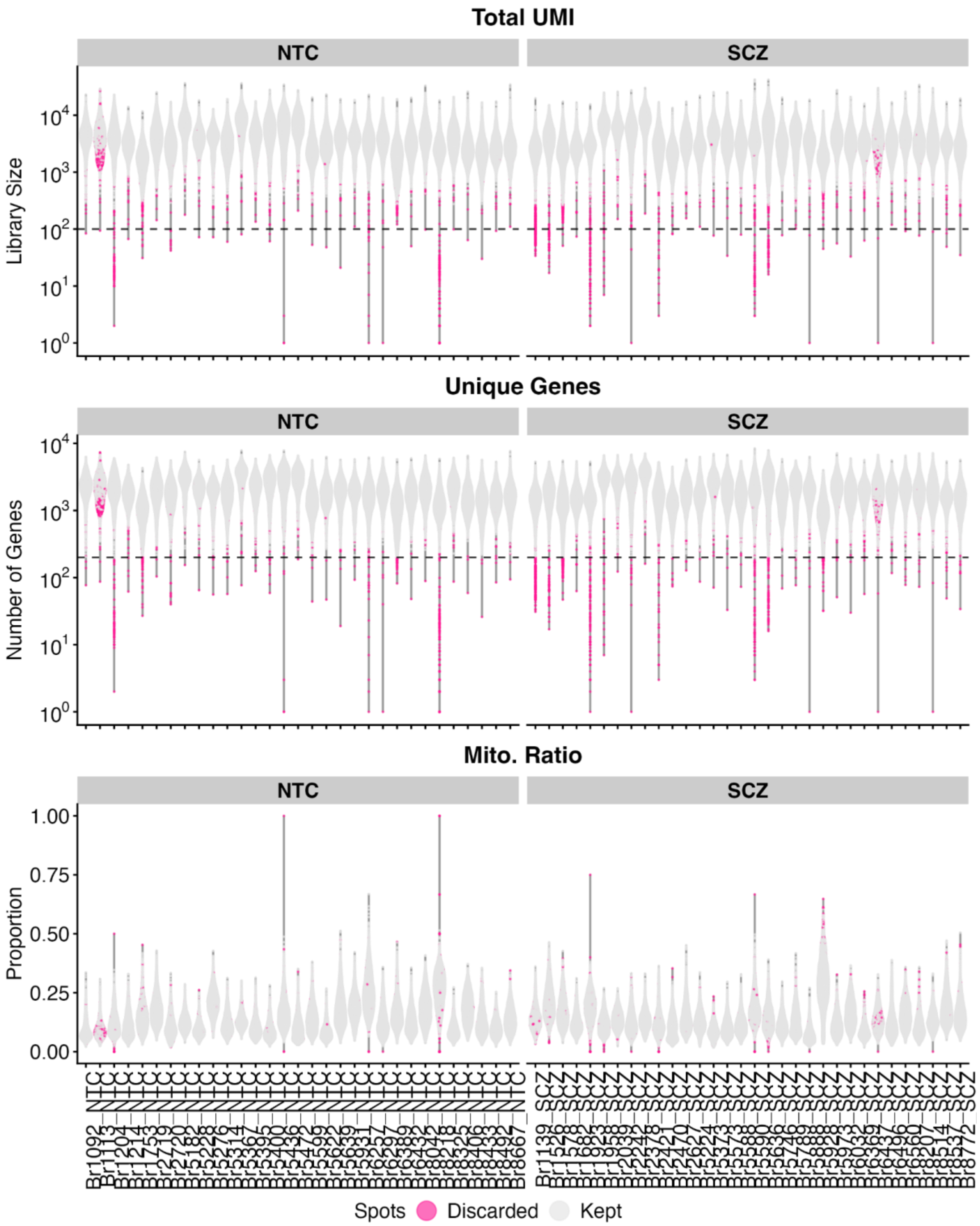

**Figure S3**. **Regional artifacts identified by SpotSweeper**. Examples from two samples (Br1113_NTC and Br6437_SCZ) illustrate technical artifacts introduced during experimental procedure, leading to diagonal scratches traversing across tissue and low-quality spots underlying the affected regions. SpotSweeper[165] identified these regional artifacts and flagged them for removal.

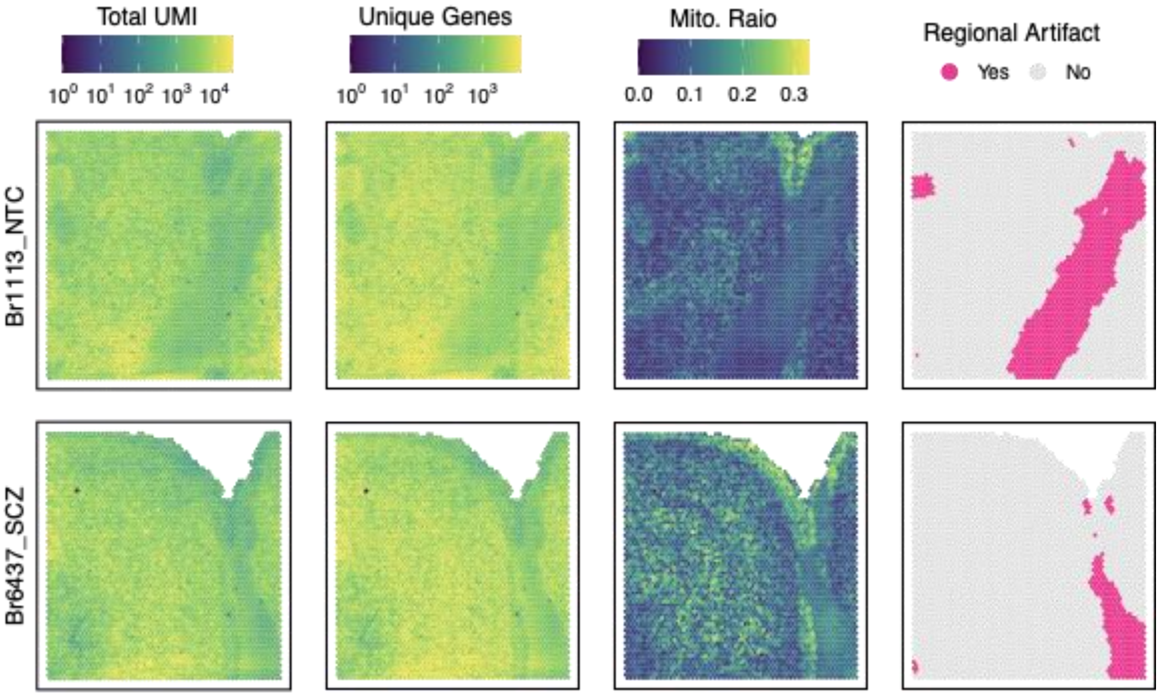

**Figure S4**. **Post-QC characterization of discarded and retained spots indicates no evidence of diagnosis-related bias**. (**Top left**) Proportion of spots discarded in NTC samples. (**Top right**) Number of spots retained after QC in NTC and SCZ samples. (**Bottom**) Empirical cumulative distribution functions (CDFs) of three QC metrics (total UMIs, number of unique genes, and proportion of mitochondrial genes) plotted for discarded (**left**) and retained (**right**) spots by diagnosis.

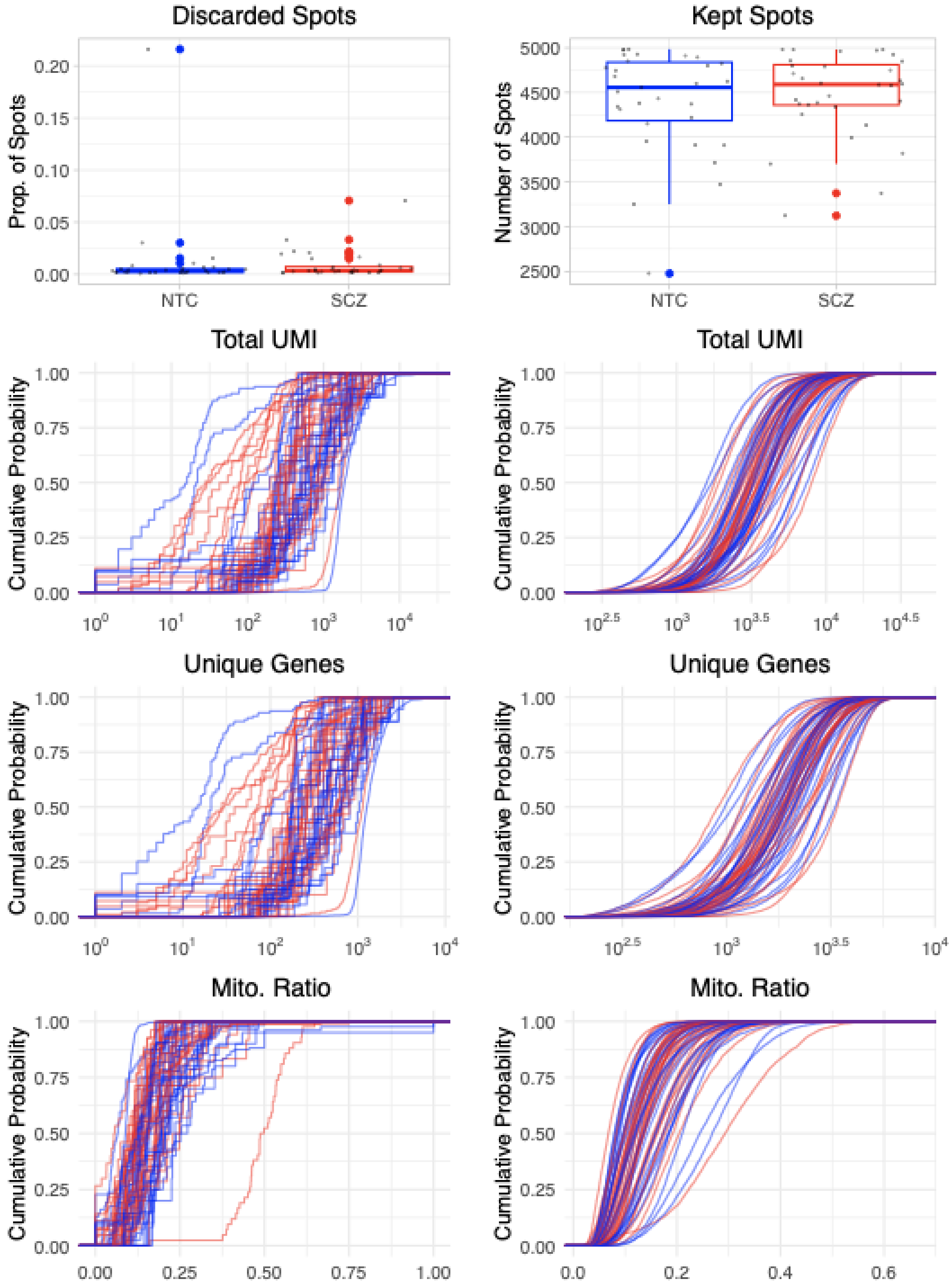

**Figure S5**. **Spot plots of data-driven SpDs at different clustering resolutions**. Unsupervised spatial clustering using PRECAST was performed at multiple resolutions (*k*=2-13), as shown in one representative sample from each diagnosis (Br8667 and Br5973, respectively). *k*=7 was chosen for downstream analysis because the patterning best correlated with established histological layers.

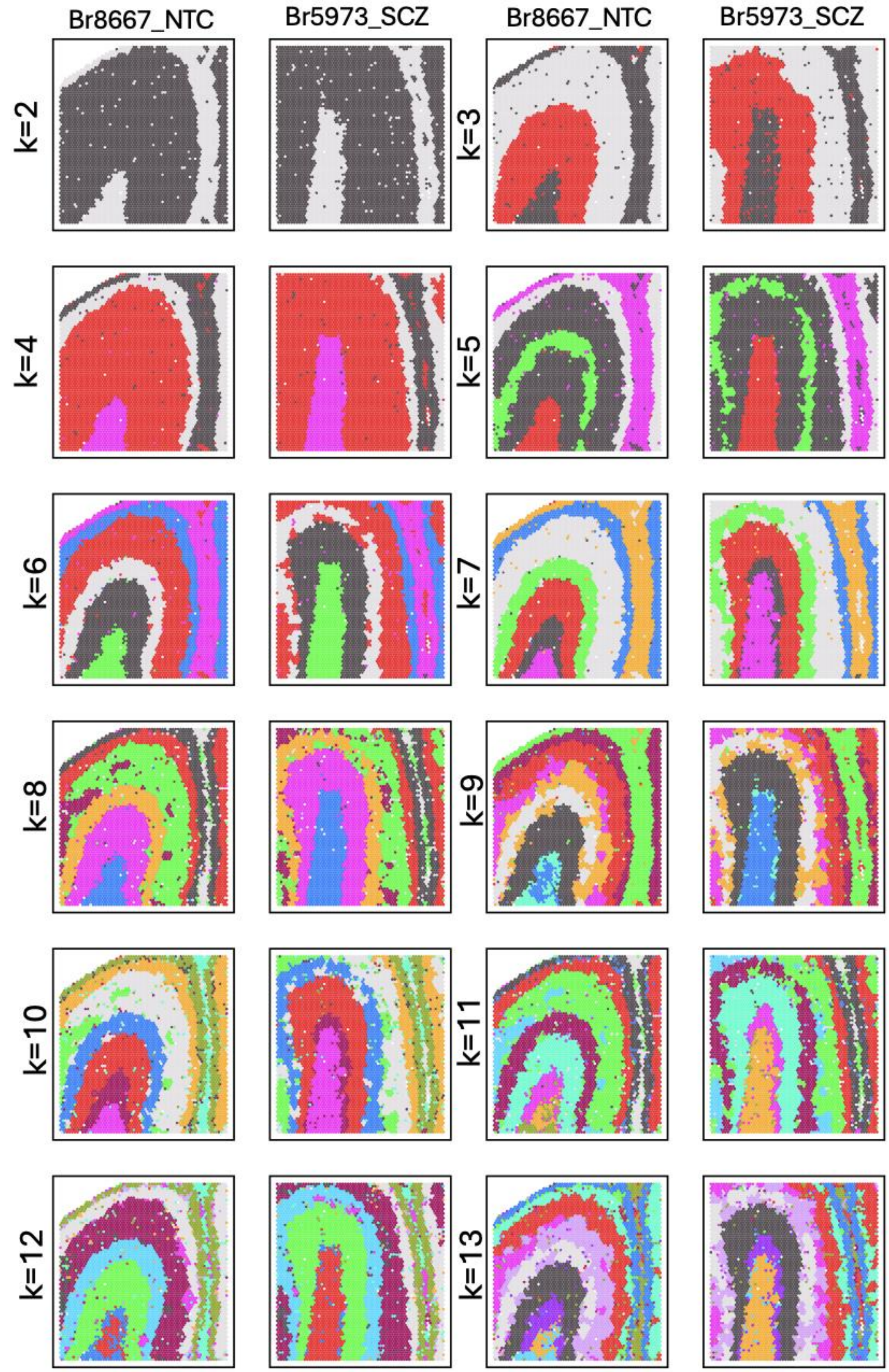

**Figure S6**. **Spot plots of the *k*=7 PRECAST SpDs across a subset of the cohort**. Spatial clustering patterns identified using PRECAST are shown for a subset of the cohort (24/63; 12 NTC and 12 SCZ). Colors represent the SpDs.

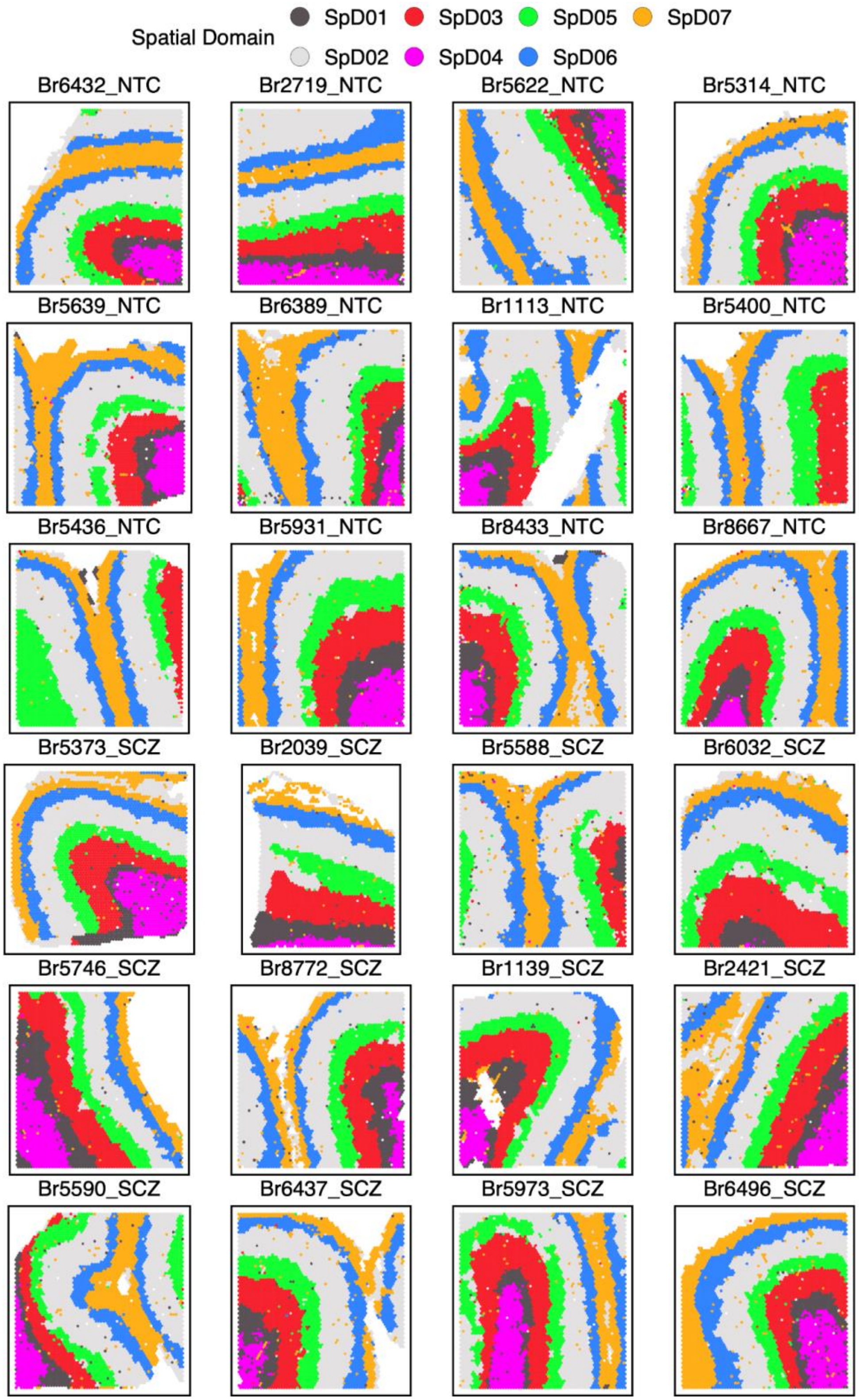

**Figure S7**. **Density curves showing the gene-level variance explained by different variables in the *k*=7 SpDs**. Spatial domain (SpD; PRECAST_07) accounts for the highest percentage of variance explained, whereas diagnosis contributes to a modest amount of variance, supporting the computational strategy to jointly clustering across the diagnostic groups.

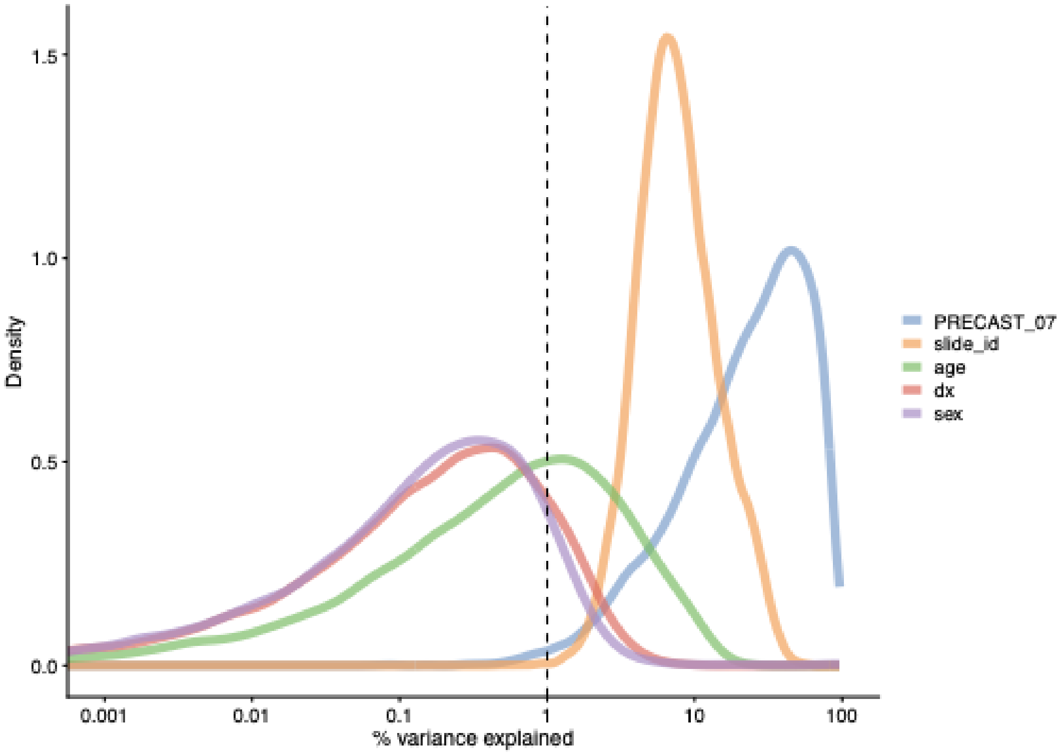

**Figure S8**. **Violin plots of representative neuronal and oligodendrocyte marker gene expression across SpDs**. Gene expression levels were pseudobulked within each individual brain donor for each SpD. Compared to SpD04-WM, SpD01-WMtz exhibited higher expression of neuronal marker genes, including *SNAP25* and *SLC17A7*, while showing relatively lower expression of oligodendrocyte genes, such as *PLP1*, *MBP*, and *MOBP*.

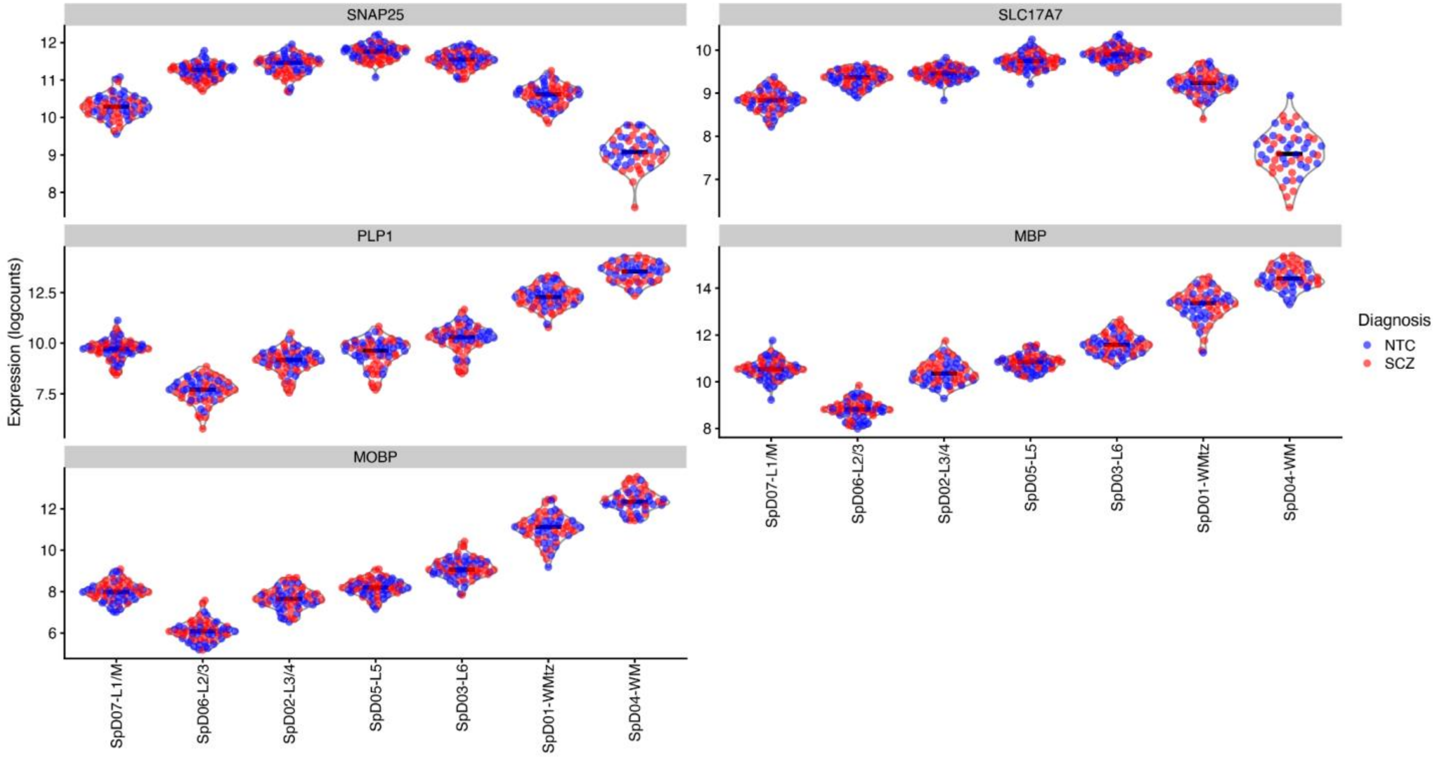

**Figure S9**. **Volcano plot of DEGs between SpD01-WMtz and SpD04-WM**. Representative cell-type-associated genes are visualized in a volcano plot (left) to illustrate relative enrichment of neuronal and synapse-related markers and depletion of oligodendrocyte markers in SpD01-WMtz compared to SpD04-WM. Genes without annotation are labeled as “NA” for “Not Annotated.” The horizontal line denotes the significance threshold of FDR=0.05. The corresponding genes highlighted in the volcano plot are listed in the table (right).

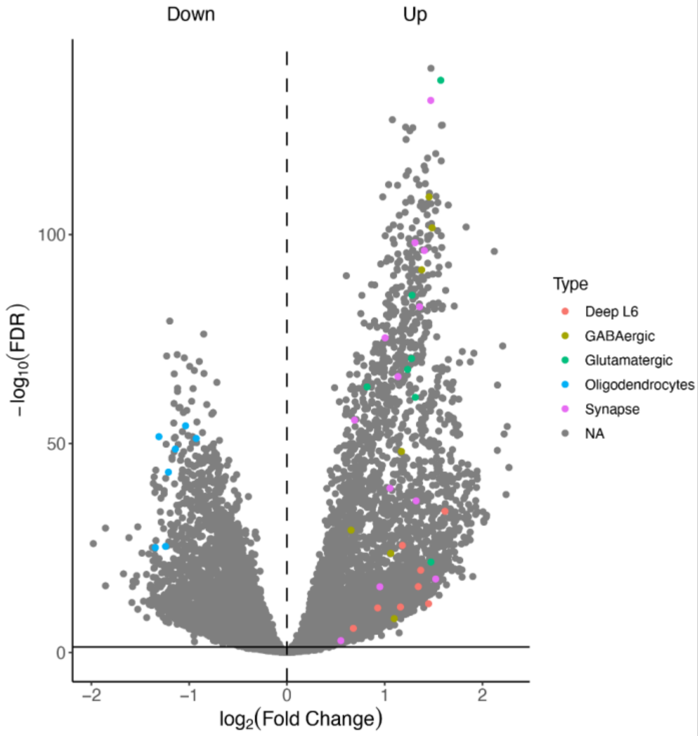


| Category | Gene | t_stat_SpD01-SpD04 | p_value_SpD01-SpD04 | fdr_SpD01-SpD04 | logFC_SpD01-SpD04 |
|---|---|---|---|---|---|
| Deep L6 neurons | CPLX3 | 5.0 | 7.1E-07 | 1.7E-06 | 0.7 |
| Deep L6 neurons | CCN2 | 10.0 | 3.3E-21 | 2.1E-20 | 1.4 |
| Deep L6 neurons | NR4A2 | 7.2 | 3.6E-12 | 1.3E-11 | 1.2 |
| Deep L6 neurons | NXPH4 | 7.4 | 5.6E-13 | 2.1E-12 | 1.4 |
| Deep L6 neurons | HS3ST2 | 13.7 | 1.3E-35 | 1.6E-34 | 1.6 |
| Deep L6 neurons | HS3ST4 | 11.6 | 2.8E-27 | 2.4E-26 | 1.2 |
| Deep L6 neurons | THEMIS | 8.8 | 3.5E-17 | 1.7E-16 | 1.3 |
| Deep L6 neurons | SEMA3E | 7.1 | 6.2E-12 | 2.2E-11 | 0.9 |
| GABAergic neurons | GABRA1 | 31.3 | 1.7E-112 | 9.3E-110 | 1.5 |
| GABAergic neurons | GABRB2 | 27.1 | 1.9E-94 | 2.7E-92 | 1.4 |
| GABAergic neurons | GABRG2 | 29.5 | 8.1E-105 | 2.4E-102 | 1.5 |
| GABAergic neurons | GABRA2 | 12.6 | 5.3E-31 | 5.3E-30 | 0.7 |
| GABAergic neurons | GAD1 | 11.1 | 2.5E-25 | 1.9E-24 | 1.1 |
| GABAergic neurons | GAD2 | 17.1 | 3.5E-50 | 7.9E-49 | 1.2 |
| GABAergic neurons | SST | 6.1 | 2.8E-09 | 8.0E-09 | 1.1 |
| Glutamatergic neurons | SLC17A6 | 10.5 | 3.2E-23 | 2.3E-22 | 1.5 |
| Glutamatergic neurons | SLC17A7 | 38.7 | 1.7E-141 | 1.2E-137 | 1.6 |
| Glutamatergic neurons | GRIN1 | 22.1 | 8.7E-73 | 4.5E-71 | 1.3 |
| Glutamatergic neurons | GRIN2A | 25.6 | 3.1E-88 | 3.2E-86 | 1.3 |
| Glutamatergic neurons | GRIN2B | 20.0 | 2.5E-63 | 8.6E-62 | 1.3 |
| Glutamatergic neurons | GRIA1 | 21.5 | 4.2E-70 | 1.9E-68 | 1.2 |
| Glutamatergic neurons | GRIA2 | 20.6 | 7.1E-66 | 2.8E-64 | 0.8 |
| Oligodendrocytes | ST18 | -11.6 | 5.0E-27 | 4.2E-26 | -1.2 |
| Oligodendrocytes | CTNNA3 | -11.5 | 9.6E-27 | 7.9E-26 | -1.3 |
| Oligodendrocytes | SLC44A1 | -18.5 | 2.1E-56 | 5.7E-55 | -1.0 |
| Oligodendrocytes | PLP1 | -15.9 | 3.7E-45 | 7.0E-44 | -1.2 |
| Oligodendrocytes | MBP | -17.2 | 9.1E-51 | 2.1E-49 | -1.1 |
| Oligodendrocytes | MOBP | -17.9 | 8.9E-54 | 2.3E-52 | -1.3 |
| Oligodendrocytes | QKI | -17.8 | 2.4E-53 | 6.0E-52 | -0.9 |
| Synapse | CAMK2A | 25.0 | 2.0E-85 | 1.8E-83 | 1.4 |
| Synapse | SNAP25 | 37.4 | 1.6E-136 | 7.6E-133 | 1.5 |
| Synapse | NRXN1 | 23.3 | 8.0E-78 | 5.1E-76 | 1.0 |
| Synapse | RELN | 3.4 | 8.7E-04 | 1.5E-03 | 0.6 |
| Synapse | BDNF | 9.4 | 4.6E-19 | 2.6E-18 | 1.5 |
| Synapse | SYT1 | 28.6 | 4.0E-101 | 8.6E-99 | 1.3 |
| Synapse | STX1A | 28.2 | 3.4E-99 | 6.4E-97 | 1.4 |
| Synapse | DLG4 | 15.0 | 3.1E-41 | 4.9E-40 | 1.1 |
| Synapse | VAMP2 | 18.8 | 8.6E-58 | 2.5E-56 | 0.7 |
| Synapse | SHANK2 | 14.3 | 3.8E-38 | 5.1E-37 | 1.3 |
| Synapse | SHANK3 | 8.8 | 3.8E-17 | 1.9E-16 | 1.0 |
| Synapse | HOMER1 | 21.2 | 2.3E-68 | 9.6E-67 | 1.1 |

**Figure S10**. **Schematic of the IF image-based strategy for classifying SPG spots into cellular microenvironments**. (**A**) Image data (Column 1) displays representative raw IF images and their corresponding thresholded binary image for each channel. SPG data (Column 2) shows these raw and thresholded IF image data overlaid with the Visium spot grid. (**B**) Spot calling (Column 3) depicts how SPG spots (highlighted in brown) were classified into SPG-defined cellular microenvironments based on enrichment of IF signals: 1. Neuropil spots lacking DAPI IF signal, 2. Neuronal spots with positive NeuN IF signal, 3. PNN spot with WFA IF signal, and 4. Vasculature spots with Claudin-5 IF signal. Created in BioRender. Kwon, S. H. (2026) https://BioRender.com/e4mjp52

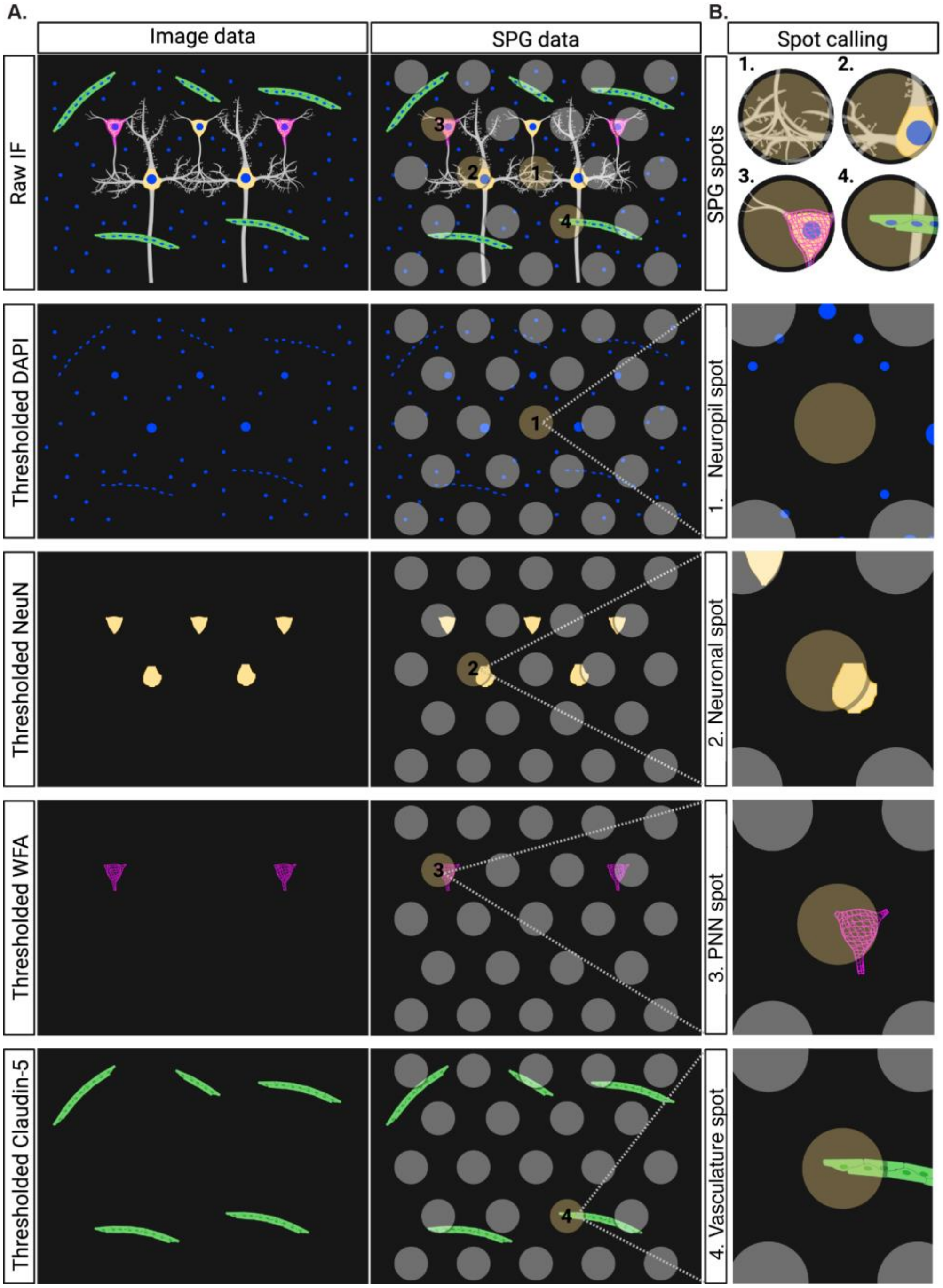

**Figure S11**. **WFA segmentation workflow to identify PNN regions of interest**. (**A**) Raw PNN channel images display the raw WFA IF images for representative NTC and SCZ samples (Br8667 and Br5973, respectively). (**B**) Intensity distribution shows the intensity histograms for the images presented in (**A**). The method for extracting the T-point threshold is illustrated,[172,173] where the red line connects the maximum and minimum intensities on the right side of the histogram. A green threshold is selected at the point where the distance between the histogram and the red line is maximized. (**C**) T-point thresholding depicts the binary images resulting from T-point thresholding applied to the raw images shown in (**A**). (**D**) Post-processing describes the additional image processing applied, including adaptive thresholding (to remove tissue background noise), autofluorescence masking, and size filtering to eliminate meninges and other noises from the T-point thresholded image. (**E**) Final segmentation shows the final segmented images containing PNN ROIs after all image processing steps, which were used for PNN spot classification.

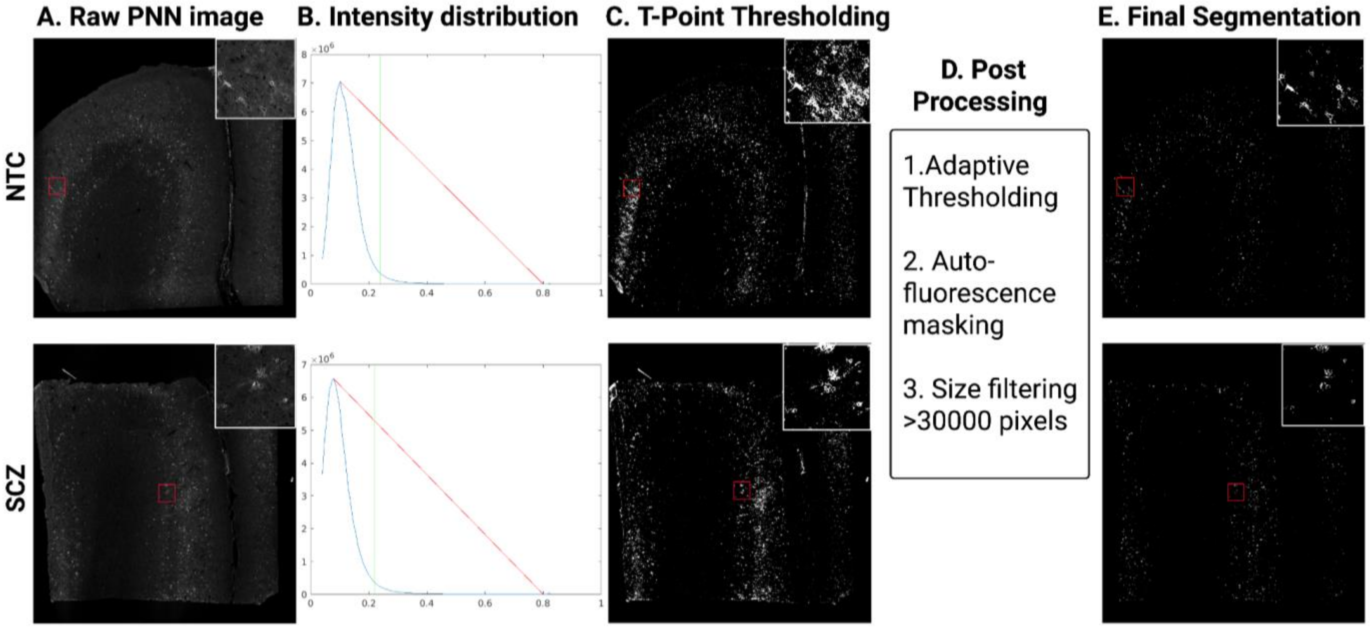

**Figure S12**. **Enrichment of SPG spots per SpD**. Heatmaps show, for each SpD, the proportion of SPG spots that are positive for each microenvironment category (neuropil, neuronal, PNN, and vasculature) calculated independently per category. Because assignments are non-mutually exclusive, spots can contribute to multiple categories, and proportions across rows do not sum to 1. Results are shown separately for NTC (left) and SCZ (right) samples. Overall, the distributions indicate highly similar SPG-defined microenvironment compositions across cortical domains between diagnostic groups.

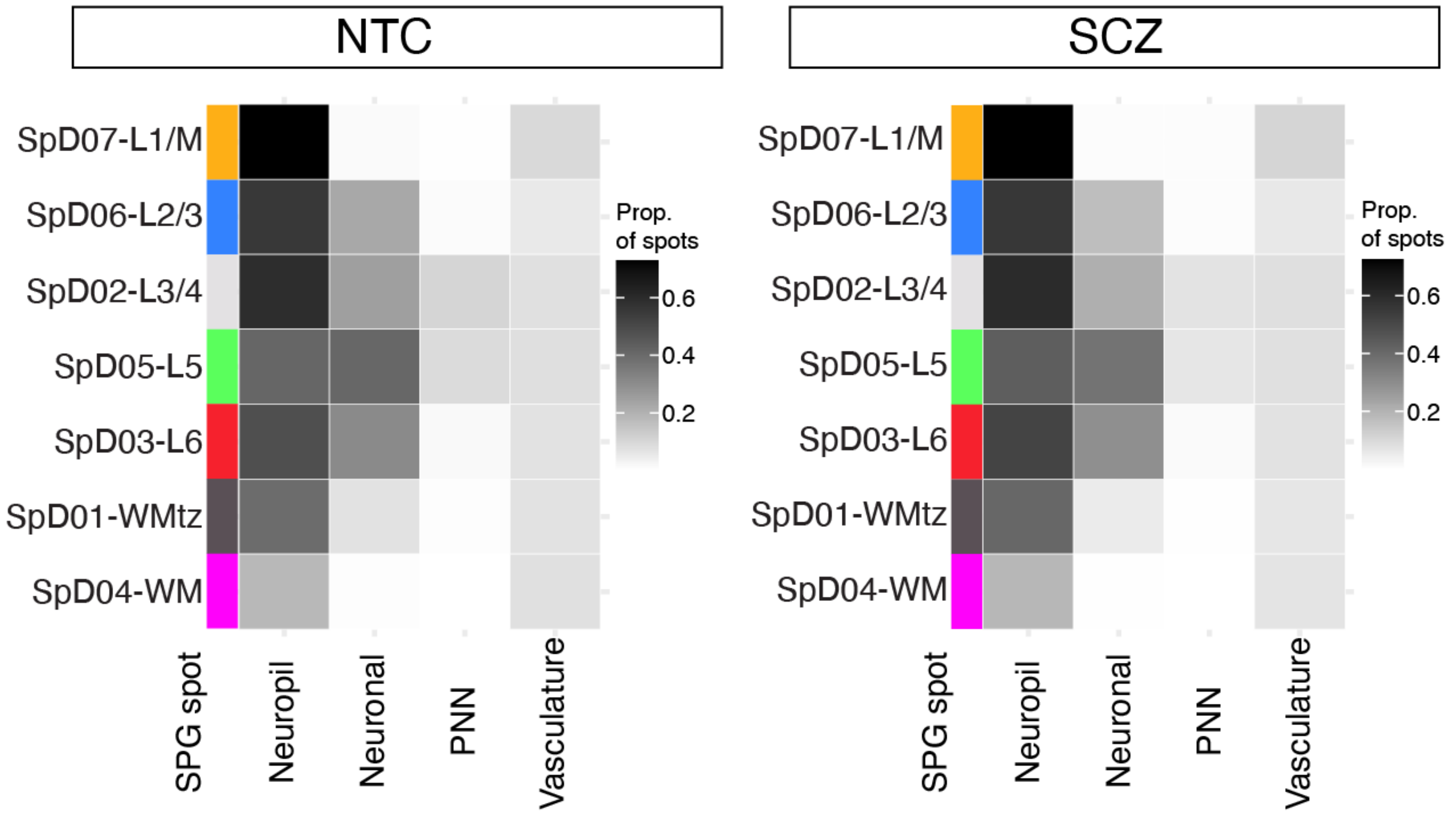

**Figure S13**. **DE analysis across SPG-defined microenvironments within NTC samples**. (**A**) Top DEGs between neuropil spots and cell body-enriched non-neuropil spots (cell body) highlight higher gene expression of synaptic genes such as *CAMK2A* in neuropil spots. Detailed DE results are provided in **Supplementary Table 3**, Sheet 1. (**B**) Top DEGs in neuronal spots demonstrate higher gene expression of neuron-associated genes such as *SNAP25* compared to non-neuronal spots. Detailed DE results are provided in **Supplementary Table 3**, Sheet 2. (**C**) Top DEGs in PNN spots highlight higher gene expression of PNN-associated genes such as *PVALB* compared to non-PNN spots. Detailed DE results are provided in **Supplementary Table 3**, Sheet 3. (**D**) Top DEGs in vasculature spots highlight higher gene expression of blood vessel-associated genes such as *CLDN5* compared to non-vasculature spots. Detailed DE results are provided in **Supplementary Table 3**, Sheet 4.

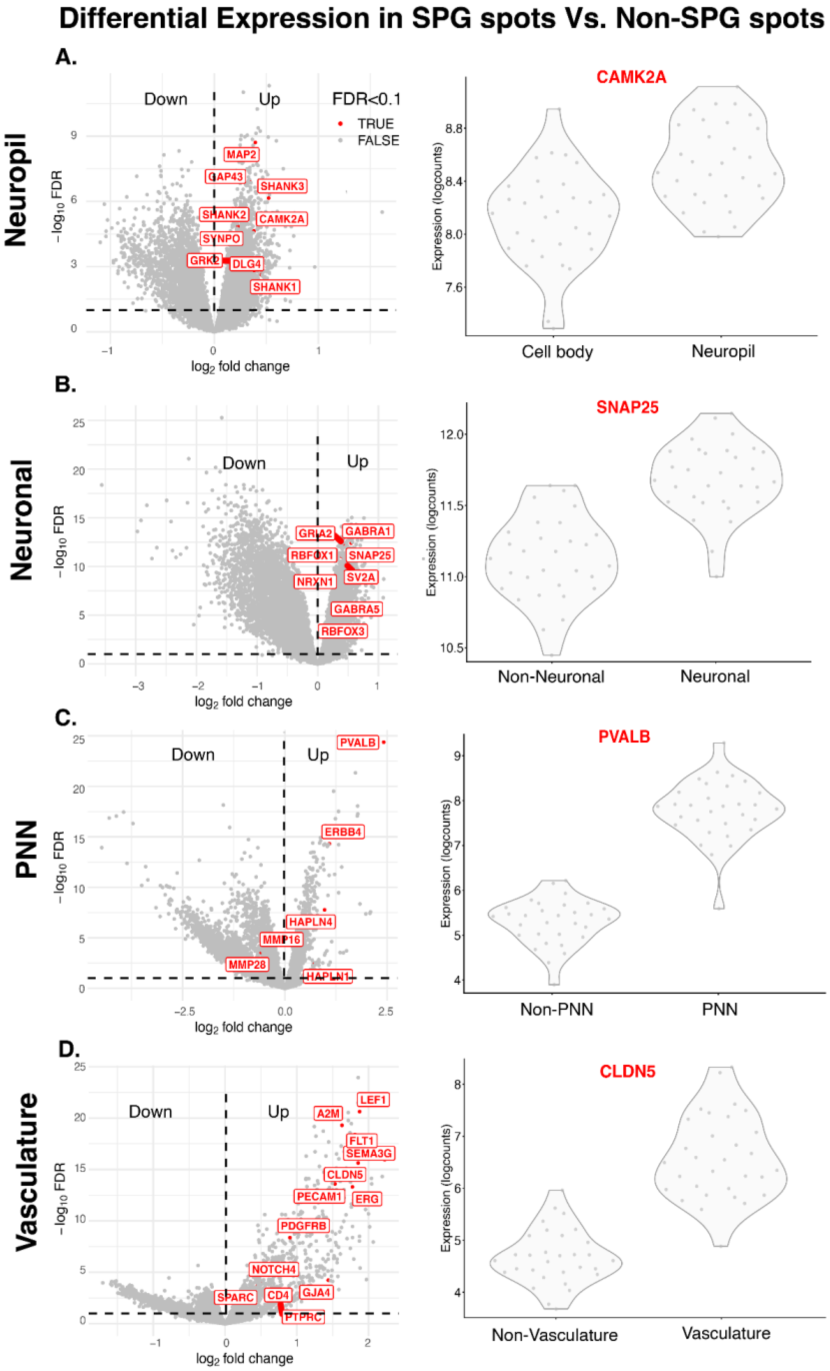

**Figure S14**. **Enrichment of SPG spots per NTC donor across SpDs**. (**A**) Proportion of neuropil spots identified: by summed marker gene expression on the left (spots with > 1,000 counts, marker genes from Niu et al.[39]), and by image-based metrics on the right (spots with < 5% DAPI signal coverage). (**B**) Left heatmap displays the proportion of neuronal spots identified by summed marker gene expression (spots with > 250 counts, marker genes from Huuki-Myers et al.[25]), compared with the right heatmap showing the proportion of neuronal spots identified using image-based metrics (spots with 5-30% NeuN signal coverage). (**C**) Left heatmap displays the proportion of PNN spots identified by summed marker gene expression (spots with > 5,000 counts, marker genes from Lupori et al.[30]), compared with the right heatmap showing the proportion of PNN spots identified using image-based metrics (spots with > 5% WFA signal coverage). (**D**) Left heatmap displays the proportion of vasculature spots identified by summed marker gene expression (spots with > 2,500 counts, marker genes from Garcia et al.[40]), compared with the right heatmap showing the proportion of vasculature spots identified using image-based metrics (spots with 5-20% Claudin-5 signal coverage).

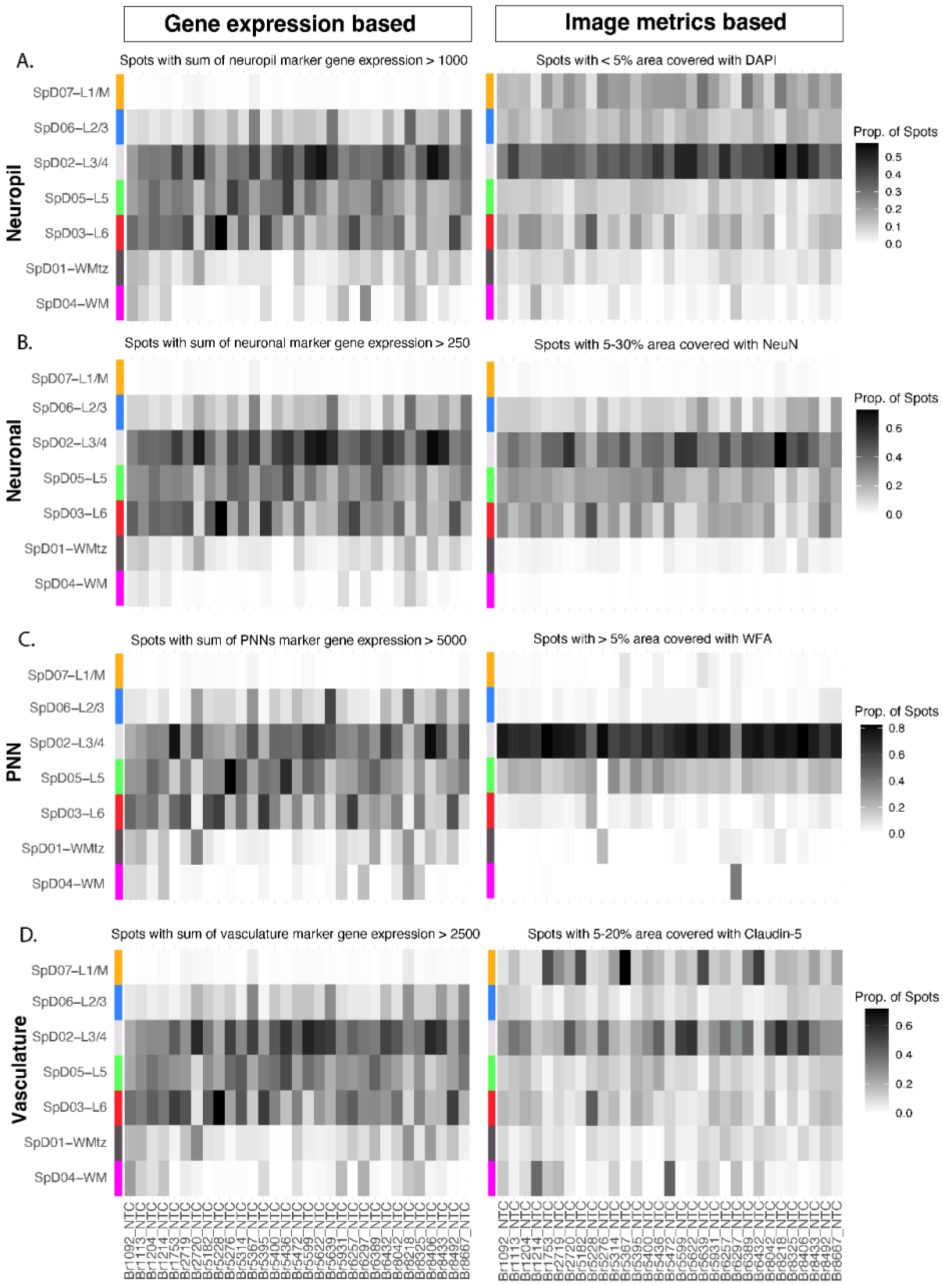

**Figure S15**. **Validation of SPG spots using reference datasets**. (**A**) Dot plots show the proportion of image-based neuropil spots (identified as spots lacking DAPI IF signal) per SpD (dot size) and the mean of summed marker gene expression from Visium spots within the SpD (dot color). Image-based SPG spot classification validated with gene expression of marker gene list derived from Niu et al.[39] (Supple_Table_10_Neuropil, sheet=10_human_synase_markers). (**B**) Neuronal spots are identified as spots containing 5-30% coverage of segmented NeuN signal. Dot plots show the proportion of image-classified neuronal spots per SpD (dot size) and the mean of summed marker gene expression from Visium spots within the same domain (dot color). Marker gene list derived from Huuki-Myers et al.[25] (TableS13_marker_stats_supp_table.xlsx, sheet=marker_stats_supp_table). (**C**) PNN spots are identified as spots containing > 5% coverage segmented WFA signal. Dot plots show the proportion of image-classified PNN spots per SpD (dot size) and the mean of summed marker gene expression from Visium spots within the same domain (dot color). Marker gene list derived from Lupori et al.[30] (Supple_Table_2_Vas.xlsx, sheet=Post Mortem Vascular Subcluster). (**D**) Vasculature spots are identified as spots containing 5-20% coverage of segmented Claudin-5 signal. Dot plots show the proportion of image-classified vasculature spots per SpD (dot size) and the mean of summed marker gene expression from Visium spots within the same domain (dot color). Marker gene list derived from Garcia et al.[40] (Supple_Table_DataS4_PNN.xlsx, sheet=PNN Energy).

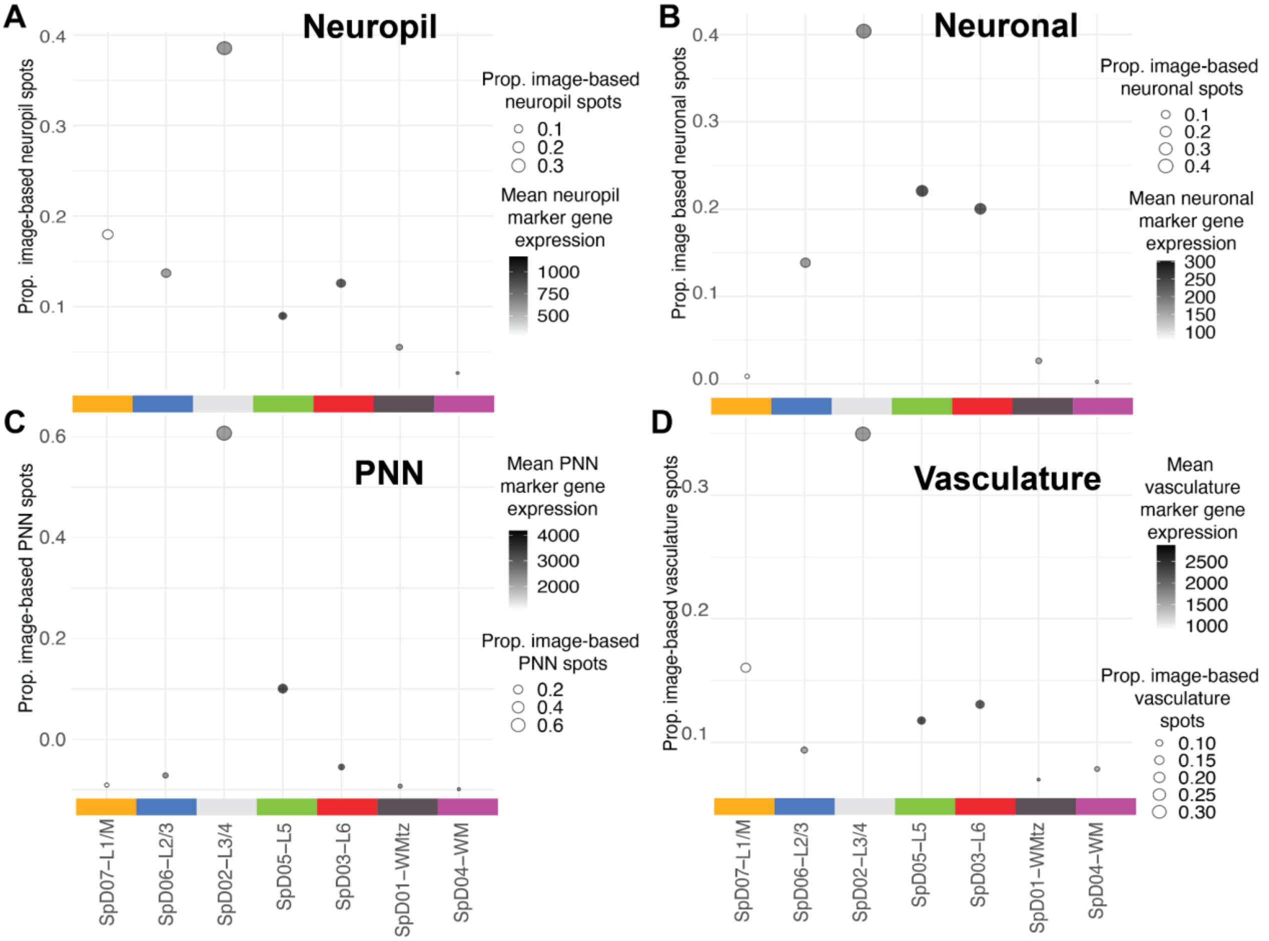

**Figure S16**. **Schematic of statistical models for identifying SCZ-associated DEGs (SCZ-DEGs)**. Three statistical models were used (I) ‘layer-adjusted’ to identify SCZ-DEGs across all cortical layers, (II) ’layer-restricted’ to identify SCZ-DEGs within each layer, and (III) ‘layer-specific’ to identify SCZ-DEGs that are unique to each layer. Created in BioRender. Kwon, S. H. (2026) https://BioRender.com/krzesaz

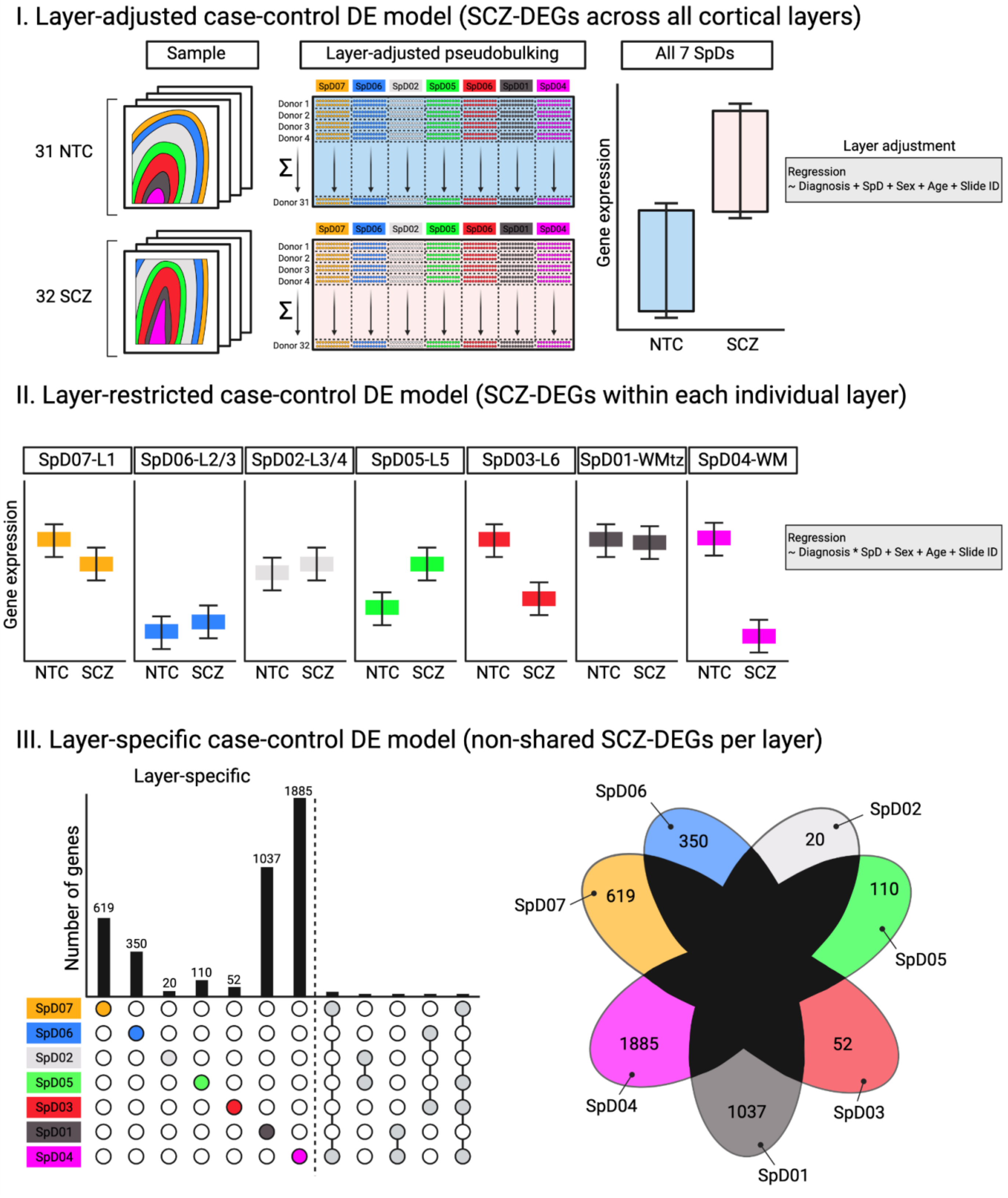

**Figure S17**. **Treemap representation of GO-BP terms enriched by GSEA based on the layer-adjusted DE model**. GSEA of all genes ranked by *t*-statistics from the layer-adjusted DE model identified significant overrepresentation of GO-BP terms (adjusted $p < 0.05$). Enrichment converging on immune-inflammatory responses and cellular responses to metal ions was observed, along with distinct enrichment of ATP metabolism and mitochondrial respiration, suggesting layer-wide metabolic alterations across cortical layers.

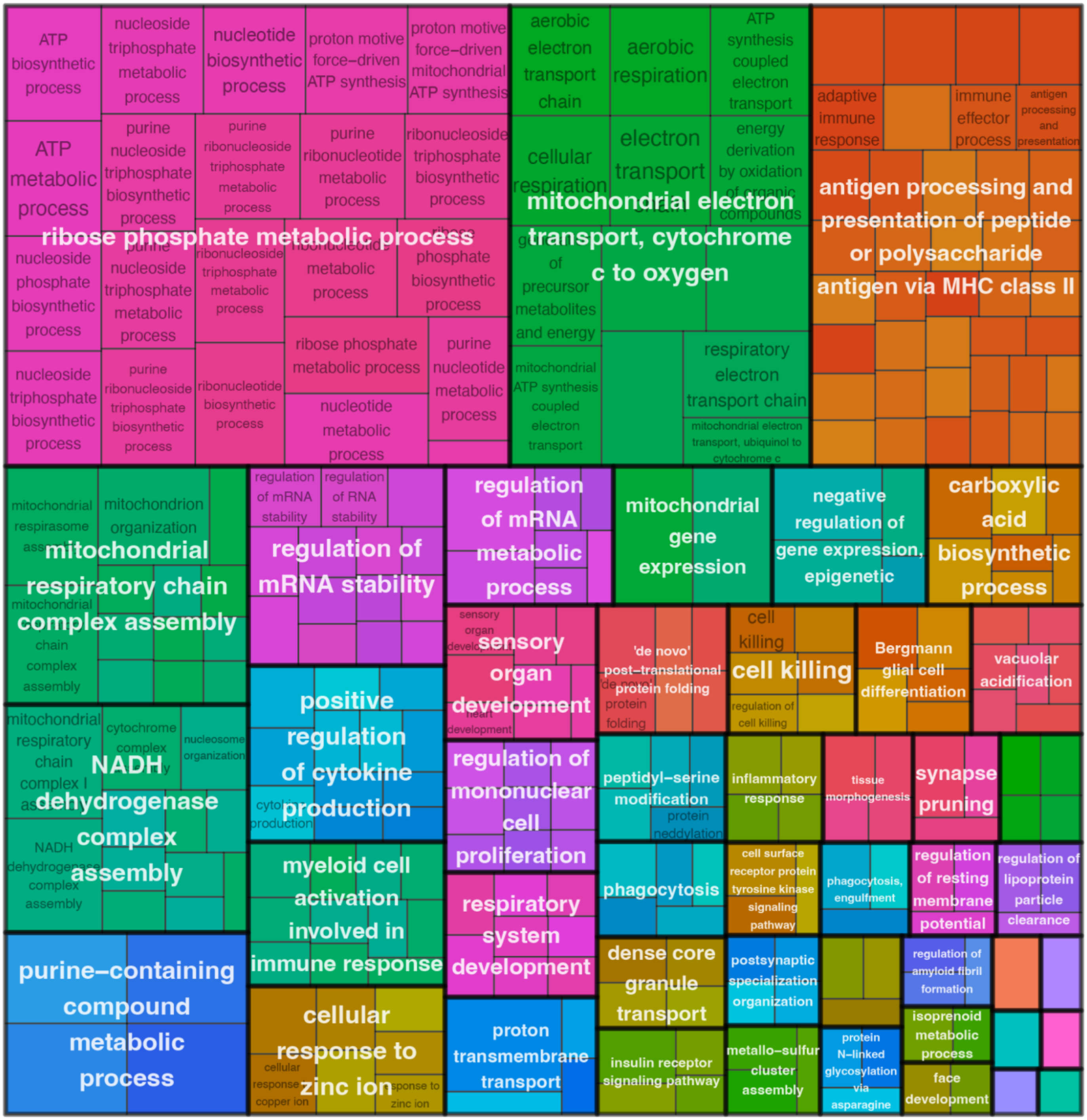

**Figure S18**. **Comparing SCZ-associated gene expression changes detected in the current Visium SRT study and in a snRNA-seq study**. (**A**) Distributions of absolute value of logFC (|log2FC|) in SCZ detected in the layer-restricted DE analysis, stratified by PRECAST SpDs. (**B**) Distributions of |log2FC| detected in cell-type-specific DE in Ruzicka et al.,[16] stratified by cell types. Due to a small amount of extreme outliers, n=148 genes (0.04%) were removed from the ridge plot due to plot truncation at 0.5. Overall, there were more glial- and WM-related signals detected than neuronal signals in the current Visium SRT dataset. This observation is consistent with cell-type-specific gene expression changes reported in Ruzicka et al..[16] Future SRT studies may require a larger sample size to detect nuanced neuronal signals in the neuronal-rich cortical domains.

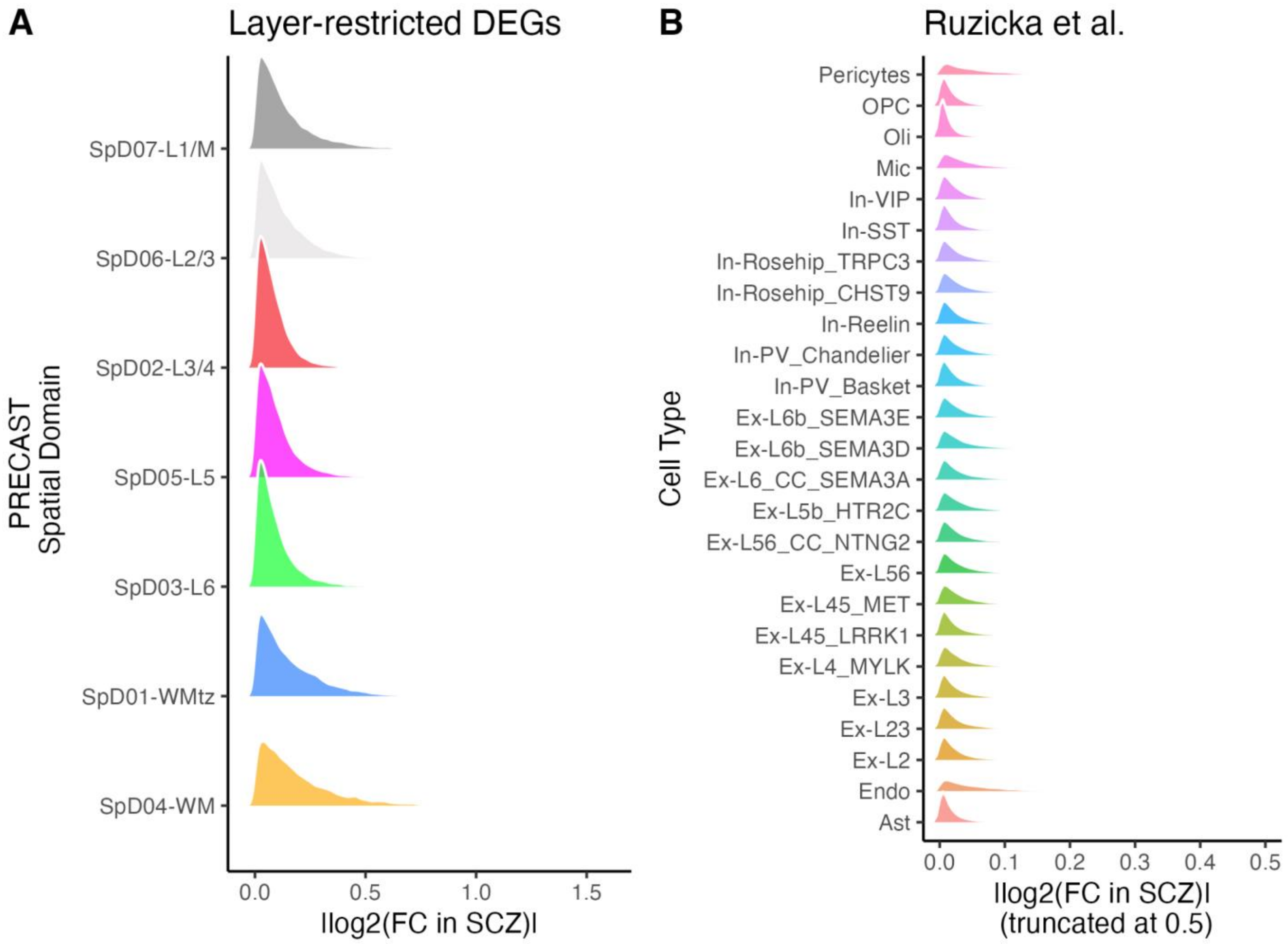

**Figure S19**. **Enrichment of BrainSEQ Phase 2 bulk RNA-seq DEGs in SPG spots across four SPG-defined microenvironments**. DEGs previously identified in the BrainSEQ Phase 2 bulk RNA-seq study[41] were tested for enrichment in SPG spots representing distinct cellular and extracellular microenvironments (All: both up- and down-regulated bulk RNA-seq DEGs; Up: up-regulated bulk RNA-seq DEGs only; Down: down-regulated bulk RNA-seq DEGs only). Robust enrichment was observed in PNN spots, particularly for down-regulated DEGs, whereas up-regulated DEGs were predominantly enriched in neuropil spots. These findings emphasize the significance of synaptic and extracellular components in SCZ and demonstrate the utility of our SRT dataset in contextualizing prior findings within anatomical tissue compartments. Color scales indicate -$\log_{10}$ ($p$-value), denoting the significance of the enrichments, and the numbers within the cells indicate the odds ratios for the significant enrichments.

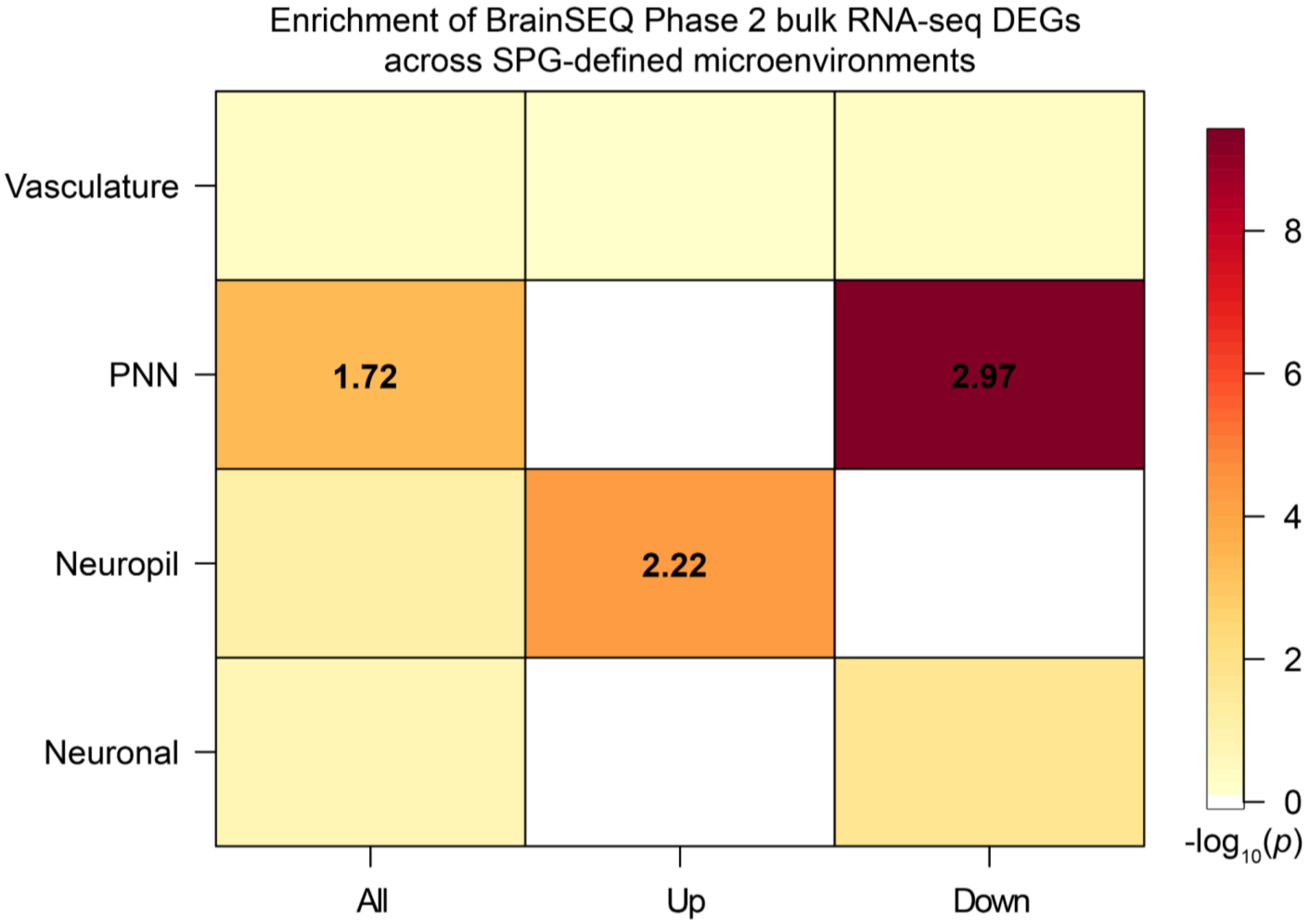

**Figure S20**. **STRING network analysis of ChEA3-derived TFs linked to layer-restricted SCZ-DEGs across non-neuronal-rich and neuronal-rich cortical domains**. STRING TF interaction networks for non-neuronal-rich (SpD07-L1/M, SpD01-WMtz, SpD04-WM) and neuronal-rich (SpD06-L2/3, SpD02-L3/4, SpD05-L5, SpD03-L6) domains highlighting functional and physical TF modules suggestive of transcriptional dysregulation hubs in SCZ. For each SpD, the top 10 TFs linked to both up- and down-regulated layer-restricted SCZ-DEGs per SpD were selected, yielding initial input sets of 60 TFs (non-neuronal) and 80 TFs (neuronal). After removing overlapping TFs and unmapped TFs (e.g., NKX62, NKX22), 48 TFs (non-neuronal) and 42 TFs (neuronal) remained for network construction. Networks are shown with all nodes including those without connections at an explorative interaction confidence (score > 0.4) compared to **Figure 4B**, revealing more extensive network connectivity.

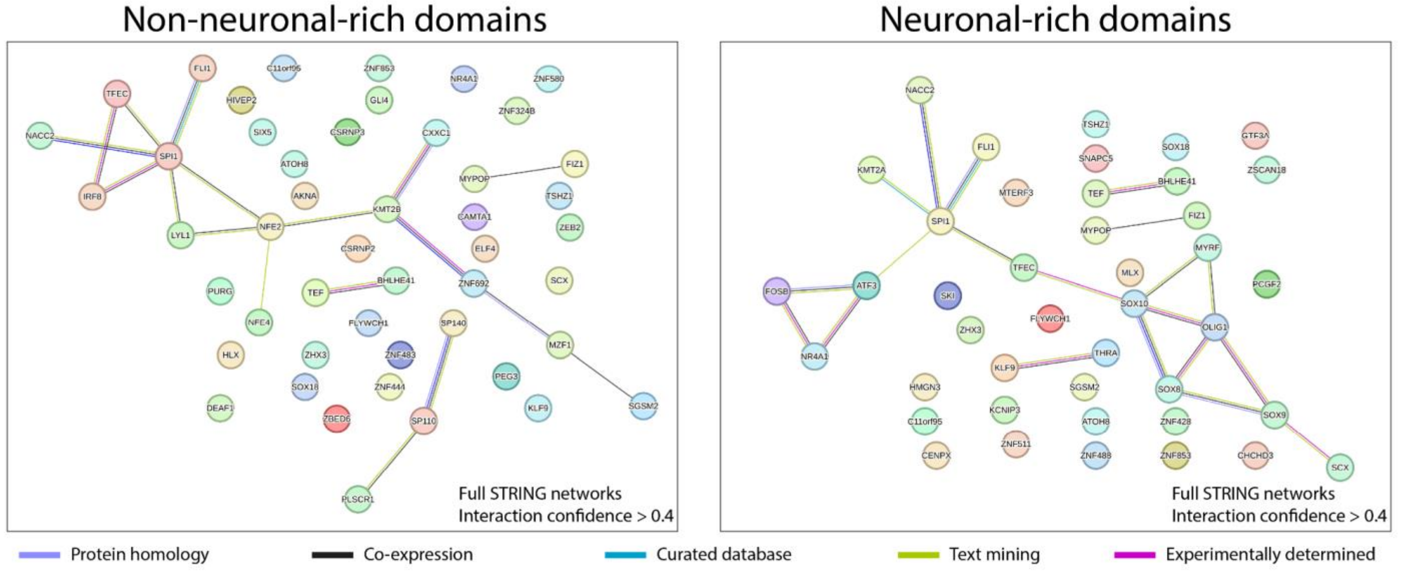

**Figure S21**. **Network diagram illustrating the ChEA3-predicted ZNF804A regulon with its associated layer-restricted SCZ-DEGs across selected SpDs (SpD07-L1/M, SpD06-L2/3, and SpD01-WMtz)**. ZNF804A-associated SCZ-DEGs were identified using ChEA3 performed on input sets of both up- and down-regulated layer-restricted SCZ-DEGs. The networks shown here represent the ZNF804A-related subset of the full ChEA3 results (provided in **Supplementary Table 6**), highlighting the predicted regulons formed by ZNF804A and its associated SCZ-DEGs in the selected SpDs referenced in **Figure 4C**. Each panel shows the predicted associations between ZNF804A (orange) and its associated input SCZ-DEGs (blue). Edges indicate predicted TF-target associations only and do not represent interaction strength or biological significance. Neuron- and synapse-associated genes among the associated SCZ-DEGs are highlighted in pink.

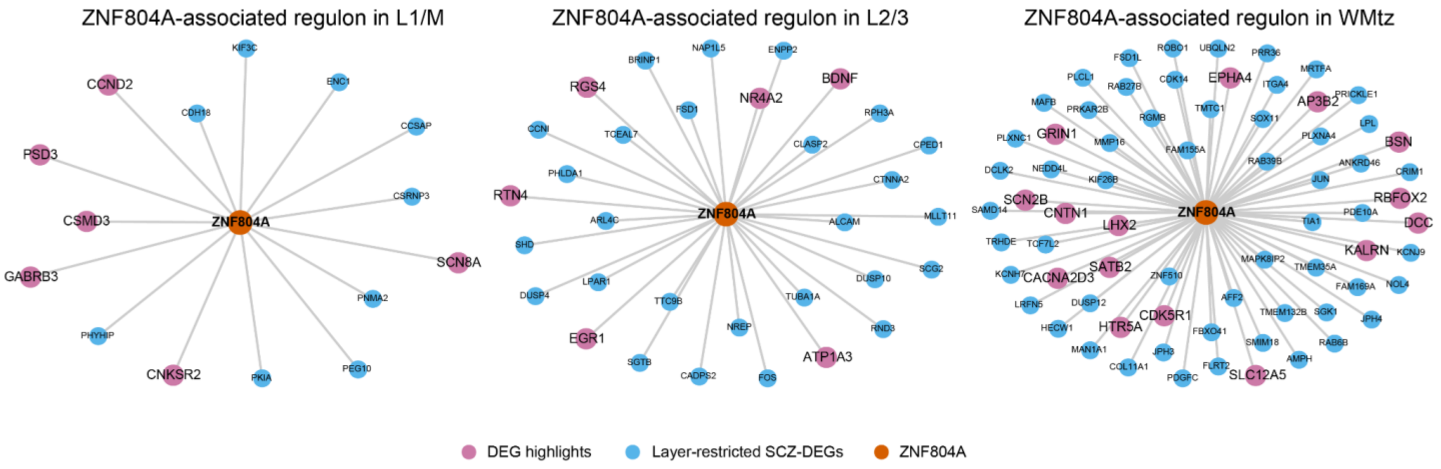

**Figure S22**. **Treemap representation of GO-BP terms enriched by GSEA from the PRS-based DE model**. GSEA of all genes ranked by *t*-statistics from the PRS-based DE model illustrates biological pathways associated with genetic risk for SCZ, identifying significant GO-BP terms (adjusted $p$ < 0.05). Enrichment converged on metabolism, gliogenesis, immune regulation, and vascular-associated processes, some of which relate to non-neuronal cellular processes.

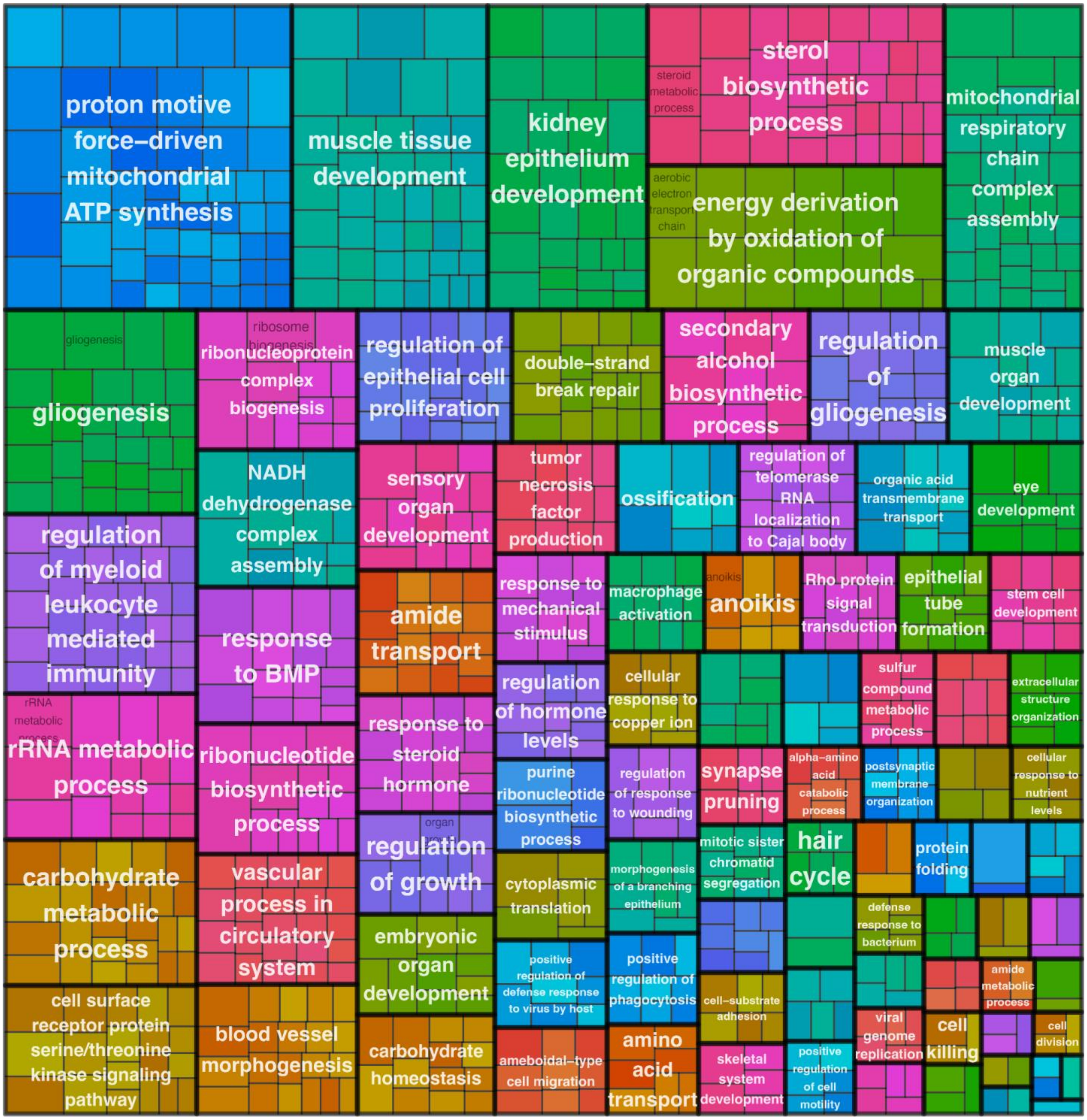

**Figure S23**. **Density plots showing spatial distribution of SCZ scDRS *z*-scores across PRECAST SpDs**. (**A**) Density plots of SCZ scDRS *z*-scores across PRECAST SpDs in all NTC and SCZ donors, colored by diagnostic group (i. normalized), and shown separately for (ii.) NTC and (iii.) SCZ donors. (**B**) Density plots of SCZ scDRS *z*-scores within each SpD across all NTC and SCZ donors, colored by diagnostic group (i. normalized), and shown separately for (ii.) NTC and (iii.) SCZ donors.

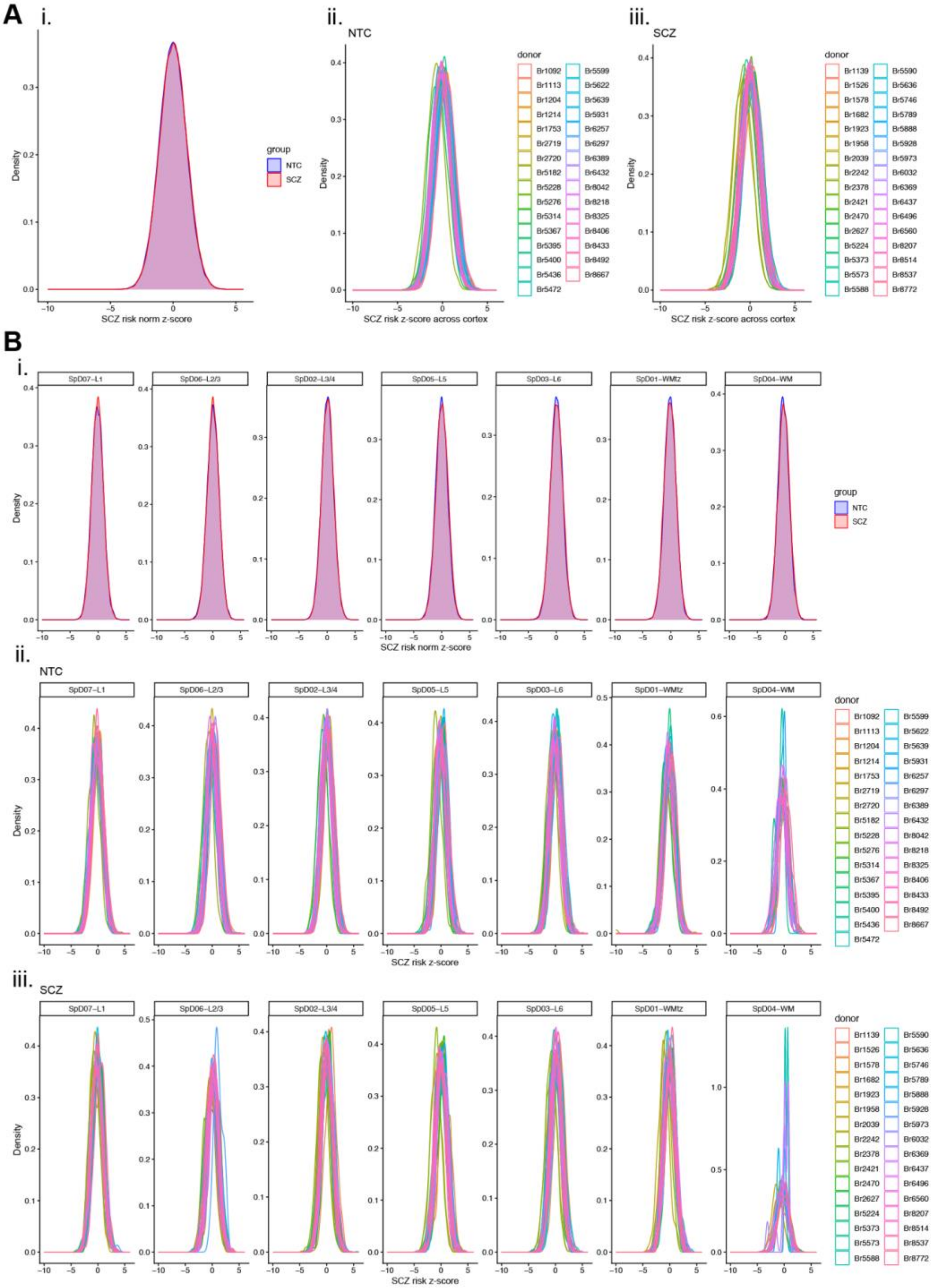

**Figure S24**. **MAGMA gene set analysis showing predominant localization of SCZ risk enrichment patterns in neuronal-rich cortical domains (L2-6)**. The MAGMA[83] heatmap shows the regression coefficient (beta) in each cortical layer with the asterisk denoting significant enrichment (FDR-adjusted $p < 0.05$) of each layer's marker genes (# in bar plot) in SCZ GWAS. Relatively high SCZ risk enrichment patterns across neuronal-rich domains spanning L2-6 were observed, consistent with the scDRS *z*-score analysis in **Figure 4H**.

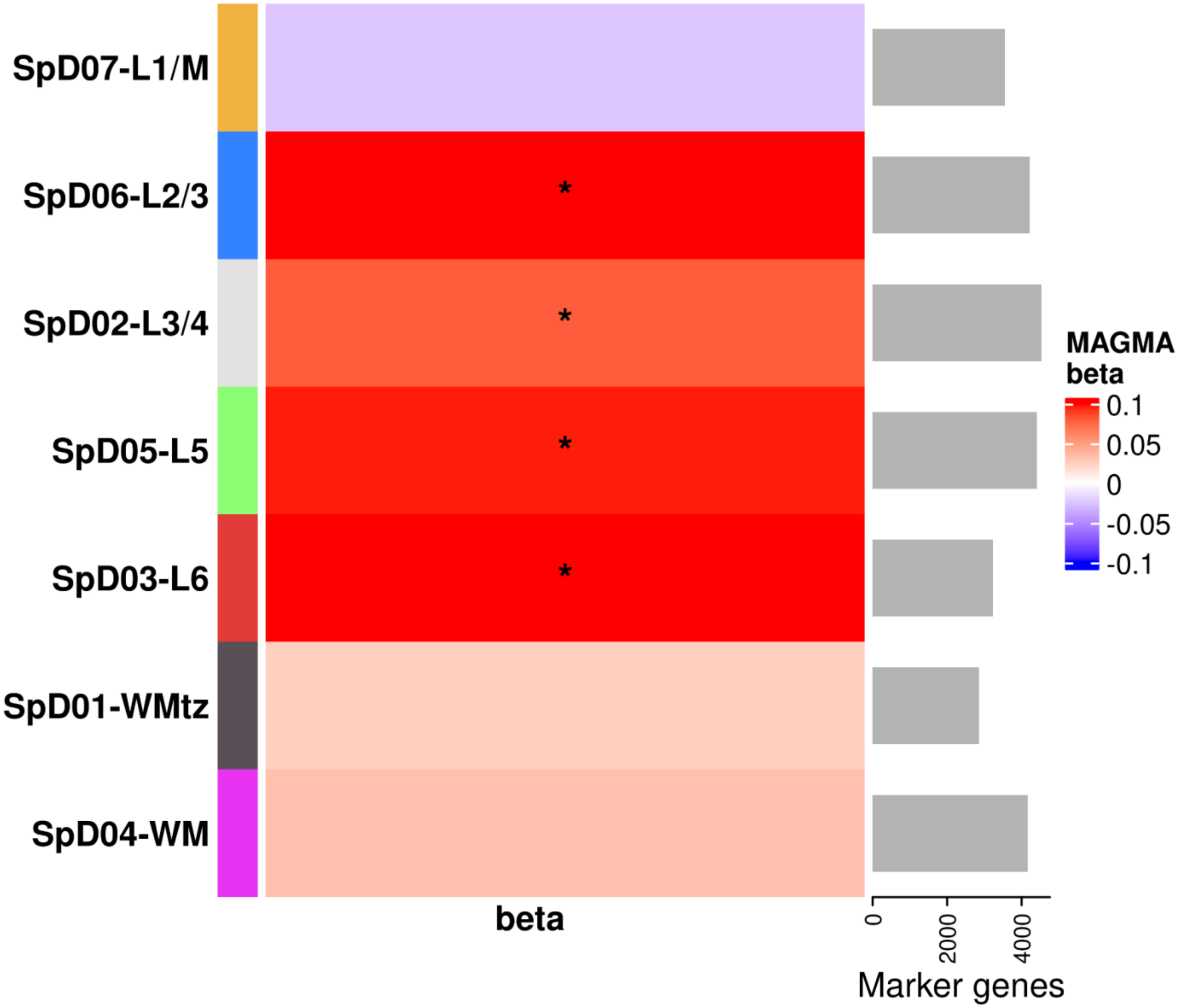

**Figure S25**. **Cell-level plots of excluded cells in the Xenium dataset**. Each point represents a single cell for the n=12 NTC and n=12 SCZ tissue samples. Discarded cells are pink while retained cells are gray. Cells with expression of negative control probes, negative control codewords, or unassigned codewords above the 99th percentile of all cells within a tissue section were discarded. Cells that were outliers in terms of number of genes detected (5 MADs below the median) or in terms of total counts (5 MADs above or below the median) were also discarded.

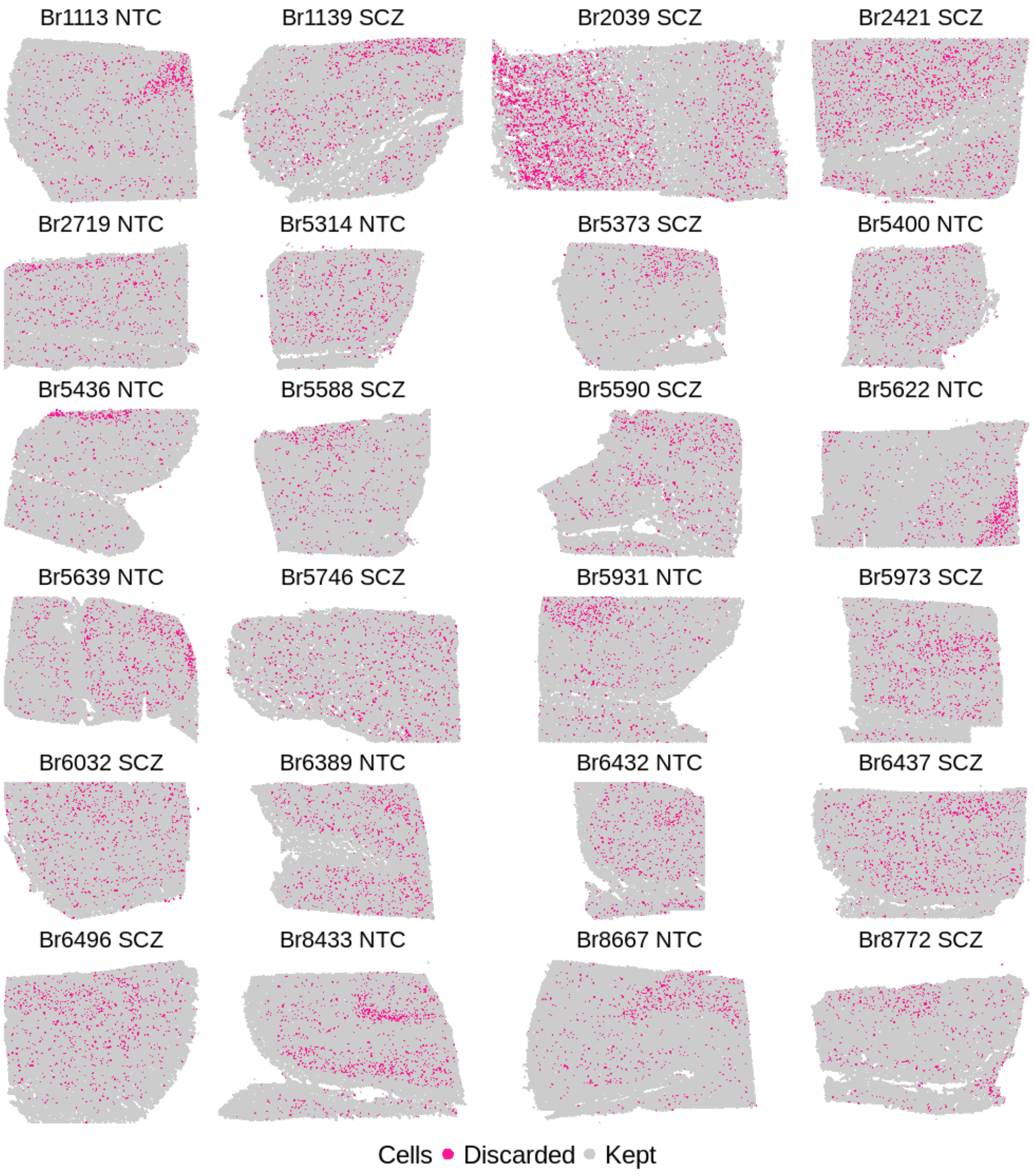

**Figure S26**. **Violin plots showing distribution of QC metrics within each sample**. The per-cell QC metrics used were the number of detected genes, the total counts across all genes, the percentage of counts from negative control probes, and the percentage of counts from negative control codewords. Each point represents a cell, and cells are colored by whether they were discarded (pink) or kept (gray). Cells that were outliers in terms of number of genes detected (5 MADs below the median) or in terms of total counts (5 MADs above or below the median) were also discarded. Cells with expression of negative control probes, negative control codewords, or unassigned codewords above the 99th percentile of all cells within a tissue section were discarded.

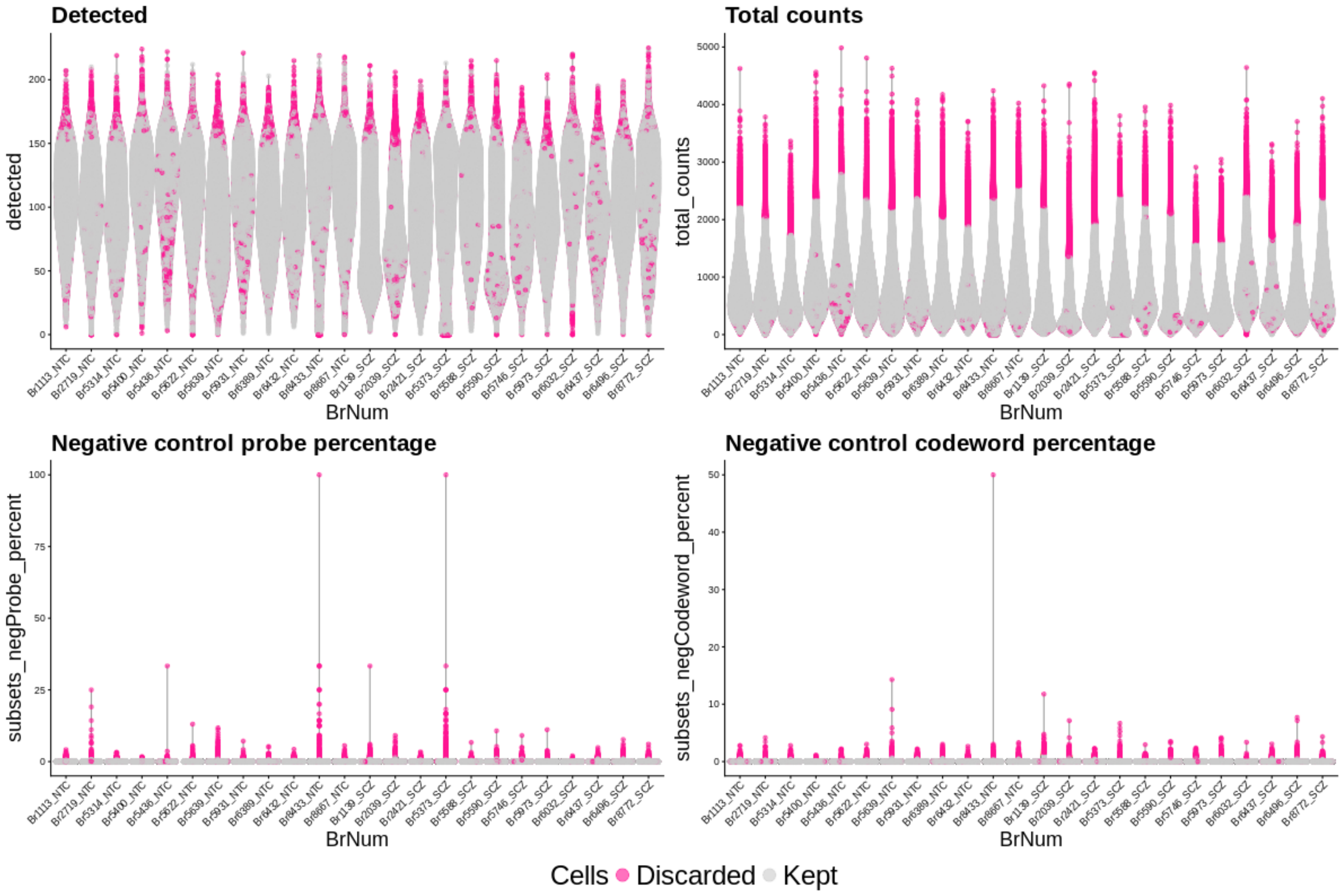

**Figure S27**. **Scatter plots of the top 4 principal components (PC) from principal component analysis (PCA) performed on the pseudobulked Xenium data**. Each point represents a unique donor-SpD combination. Points are shaped by the diagnosis of the donor and colored by the SpD. PC1 and PC2 both appear to be driven by the difference between the SpDs, with PC1 specifically being driven by SpD07-L1/M. For brevity, 'SpD0X' labels are not shown in the plot.

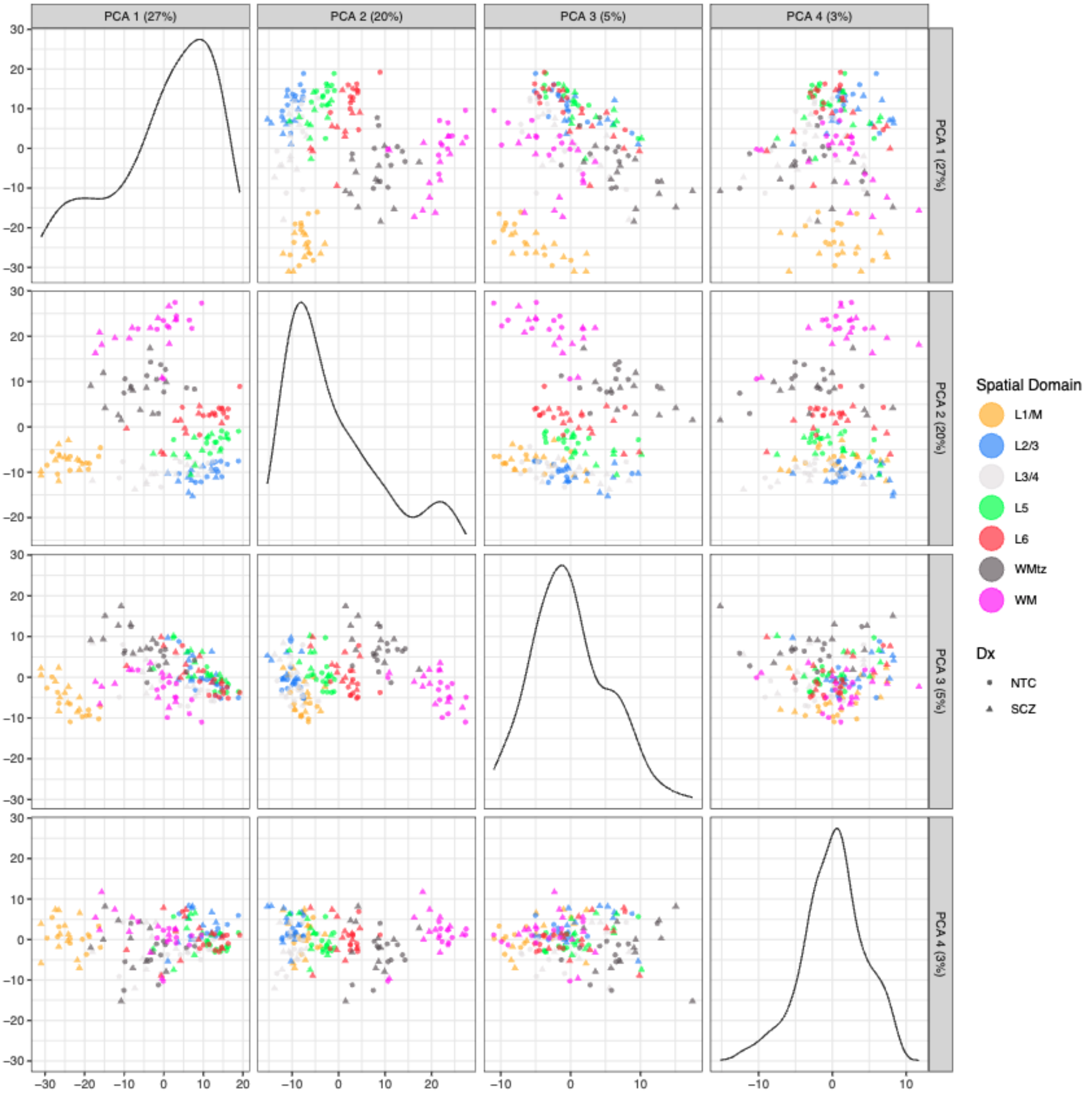

**Figure S28**. **Spatial registration correlation heatmap of Xenium predicted cell-level domains against manual annotations**. Spatial registration of spaTransfer predicted domains on the Xenium data (y-axis) against manual annotations of dlPFC layers from Maynard, Collado-Torres et al..[27] Higher values (green) indicate higher Pearson correlation in enrichment *t*-statistics between predicted cell-level domains and manual annotations. To validate our predictions, predicted cell-level domains were then annotated with their best matching manual annotation based on this correlation. Annotations with high confidence are denoted with an 'X'. For brevity, 'SpD0X' labels are not shown in the heatmap.

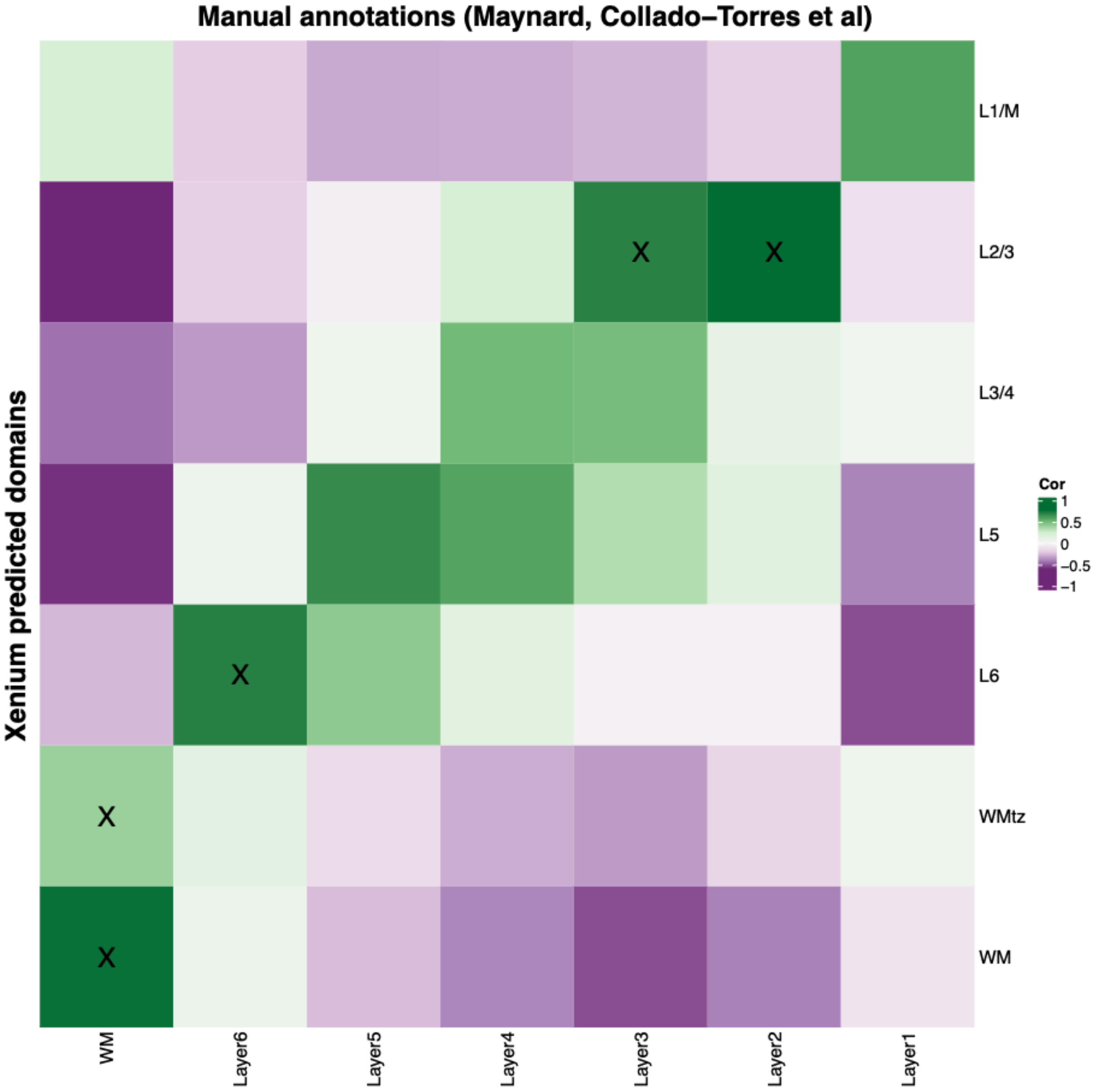

**Figure S29**. **Heatmap of the marker gene expression for each of the 18 BANKSY clusters**. Each column is a pseudobulked sample at the donor-cluster level, and each row is a cell-type marker gene included on the Xenium panel. Expression values are log-transformed library-size-normalized counts. Rows were standardized to have a mean of 0 and a variance of 1. Marker genes are grouped by their representative cell type. Cell type abbreviations: Ast, astrocytes; Endo, endothelial cells; Ex, excitatory neurons; In, inhibitory neurons; MIC, microglia; MIC_immune, microglia- or immune-associated cells; Oligo, oligodendrocytes.

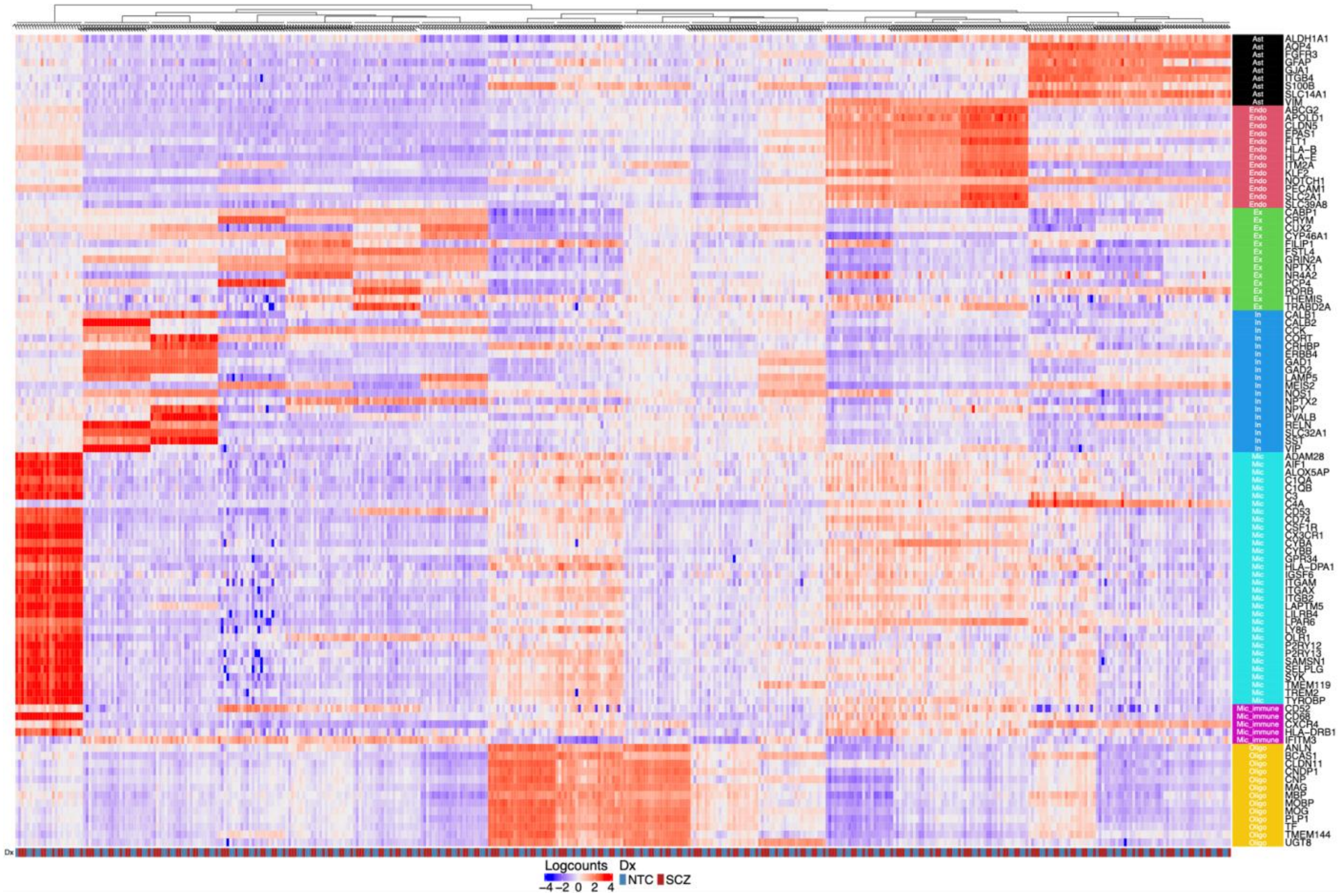

**Figure S30**. **Heatmap of cell-type proportion within each SpD**. Each column represents a unique cell type-diagnosis combination, and each row represents a SpD. The heatmap is colored by the proportion of a cell type within a SpD, computed across all donors within a diagnostic group. Across diagnosis, astrocytes and oligodendrocytes are enriched in the SpD04-WM and SpD01-WMtz, and endothelial cells are enriched in SpD07-L1/M. The neuronal cell types show enrichment in SpD06-L2/3 through SpD03-L6. For brevity, 'SpD0X' labels are not shown in the heatmap.

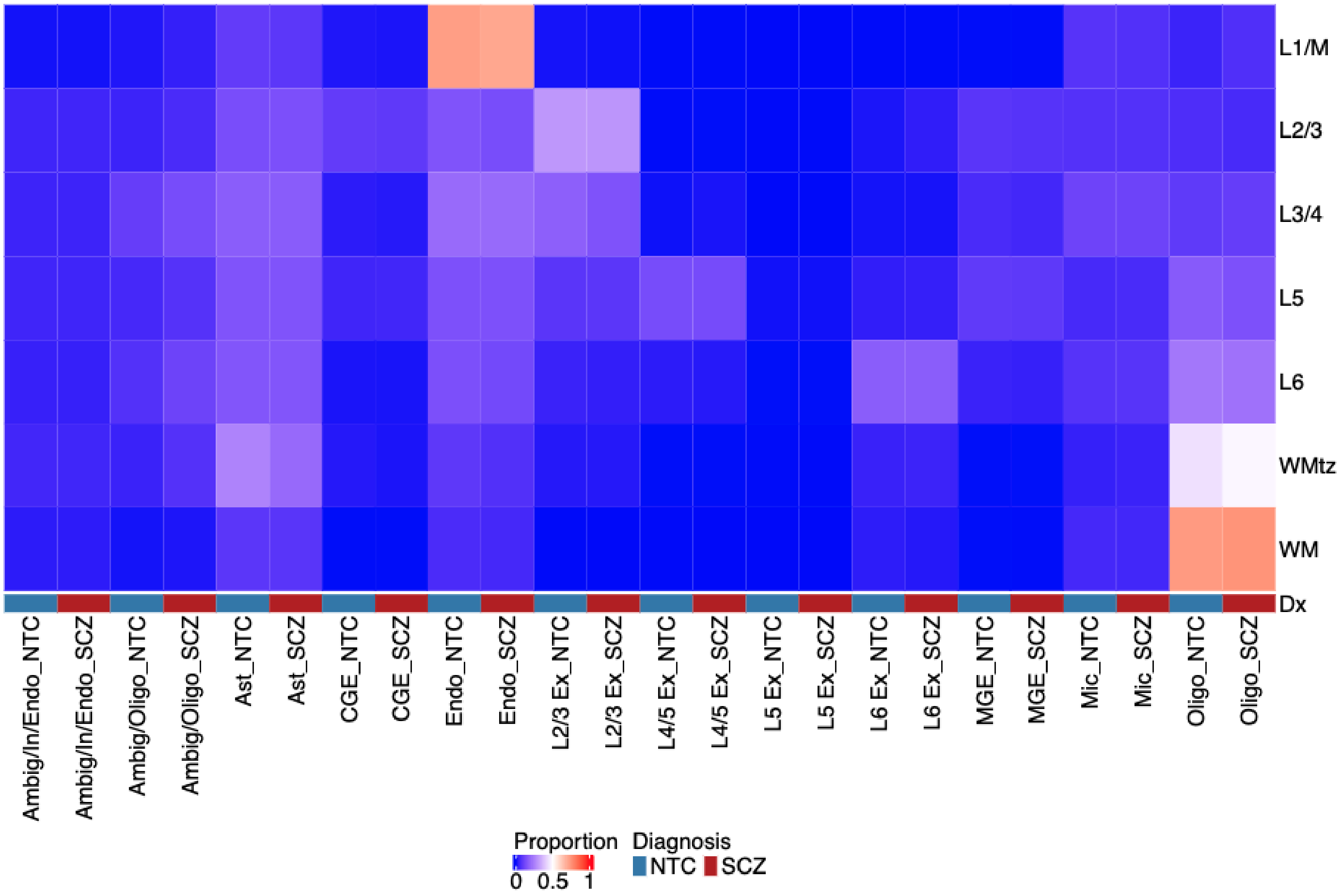

**Figure S31**. **Cell-type composition analysis reveals enrichment of L6 Ex neurons in L2/3 in SCZ**. (**A**) Compositional data analysis results for SpD06-L2/3 from the cacoa analysis.[98] The x-axis of the box plot represents the loadings from linear discriminant analysis. Loadings greater than zero indicate enrichment in SCZ, and loadings less than zero indicate enrichment in NTC. Cell types in the box plot are ordered vertically from lowest to highest *p*-value. Cell types above the red line have statistically significant enrichment. The corresponding bar chart depicts -log$_{10}$ (adjusted *p*-value). The black line indicates the *p*-value cutoff for significance. (**B**) Bar chart showing proportions of cells in SpD06-L2/3 that are L6 Ex neurons. Despite significant enrichment of L6 Ex neurons seen in (**A**), the proportion of L6 Ex neurons in L2/3 remains low for both NTC and SCZ. Notably, the percentage of L6 Ex neurons in SpD06-L2/3 ranged from 1.68% to 4.26% for SCZ donors.

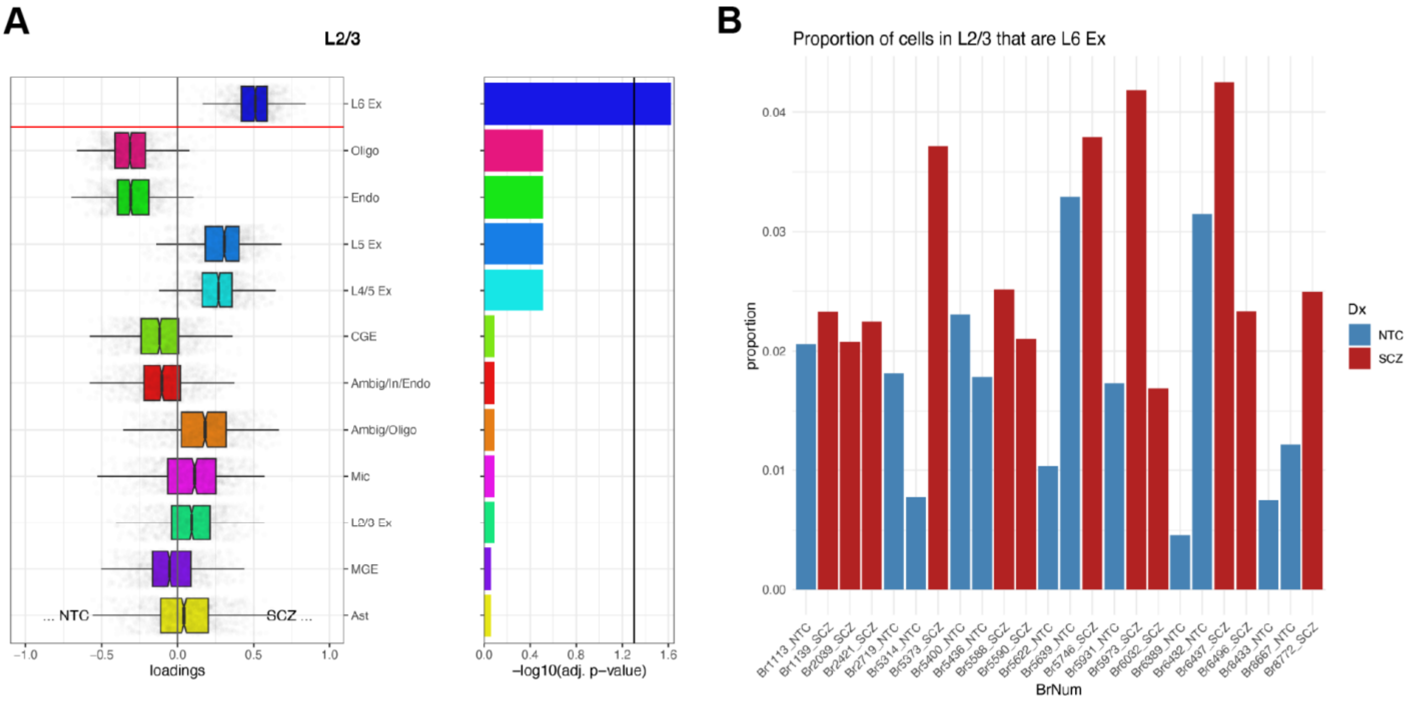

**Figure S32**. **Heatmap showing pseudobulked expression of transcription factor ZNF804A in the Xenium data**. Each column represents a unique pseudobulked donor-SpD combination and each row represents a cell type. Expression values are log-transformed library-size-normalized counts of *ZNF804A*. *ZNF804A* appears to be enriched in the MGE cell type across diagnosis.

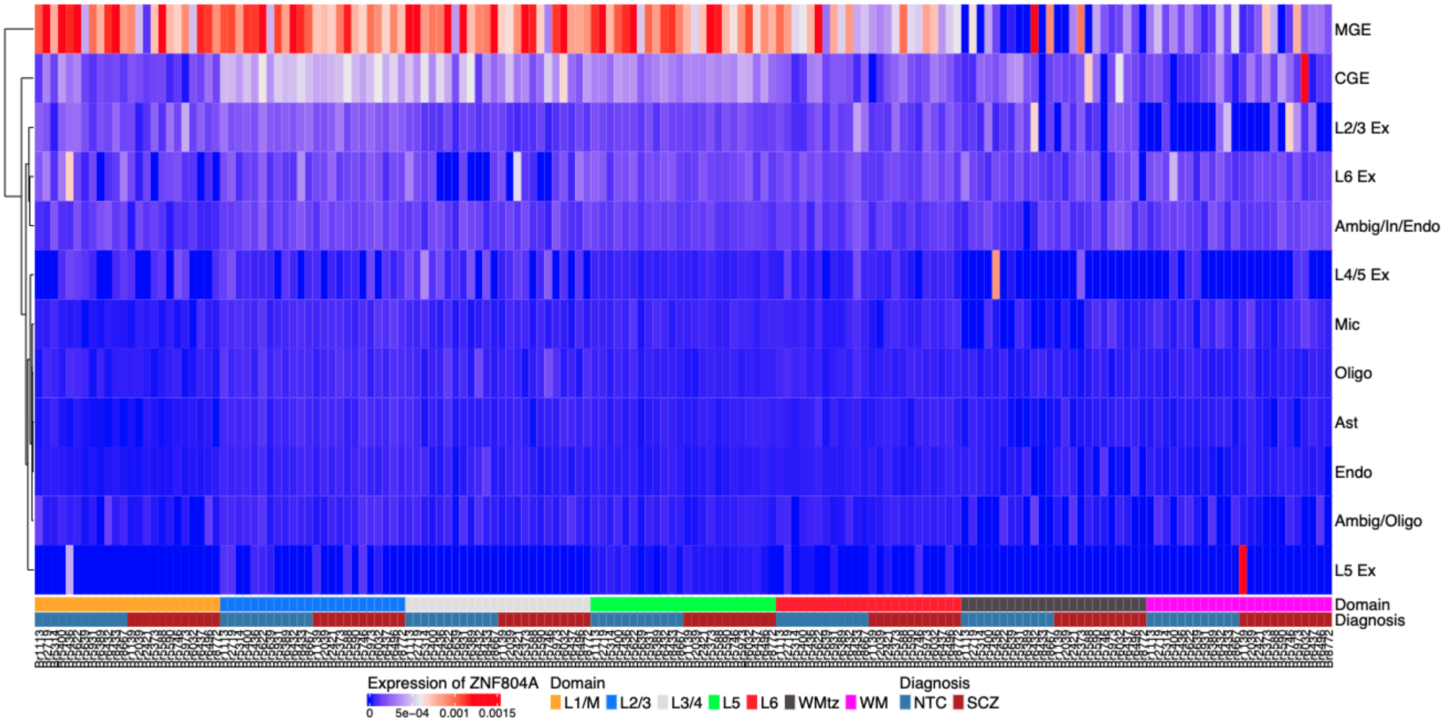

**Figure S33**.**Complementary SynGO enrichment analysis of PNN microenvironment-restricted SCZ-DEGs**.

SynGO enrichment analysis (CC) was performed for microenvironment-restricted SCZ-DEGs identified at nominal *p* < 0.05 within PNN microenvironments using the default ‘brain-expressed’ background gene set, supporting significant enrichment of genes associated with postsynaptic compartments at a stringent threshold (FDR < 0.01). Tile color indicates enrichment significance ($-\log_{10}(q\text{-value})$), with representative synaptic genes labeled.

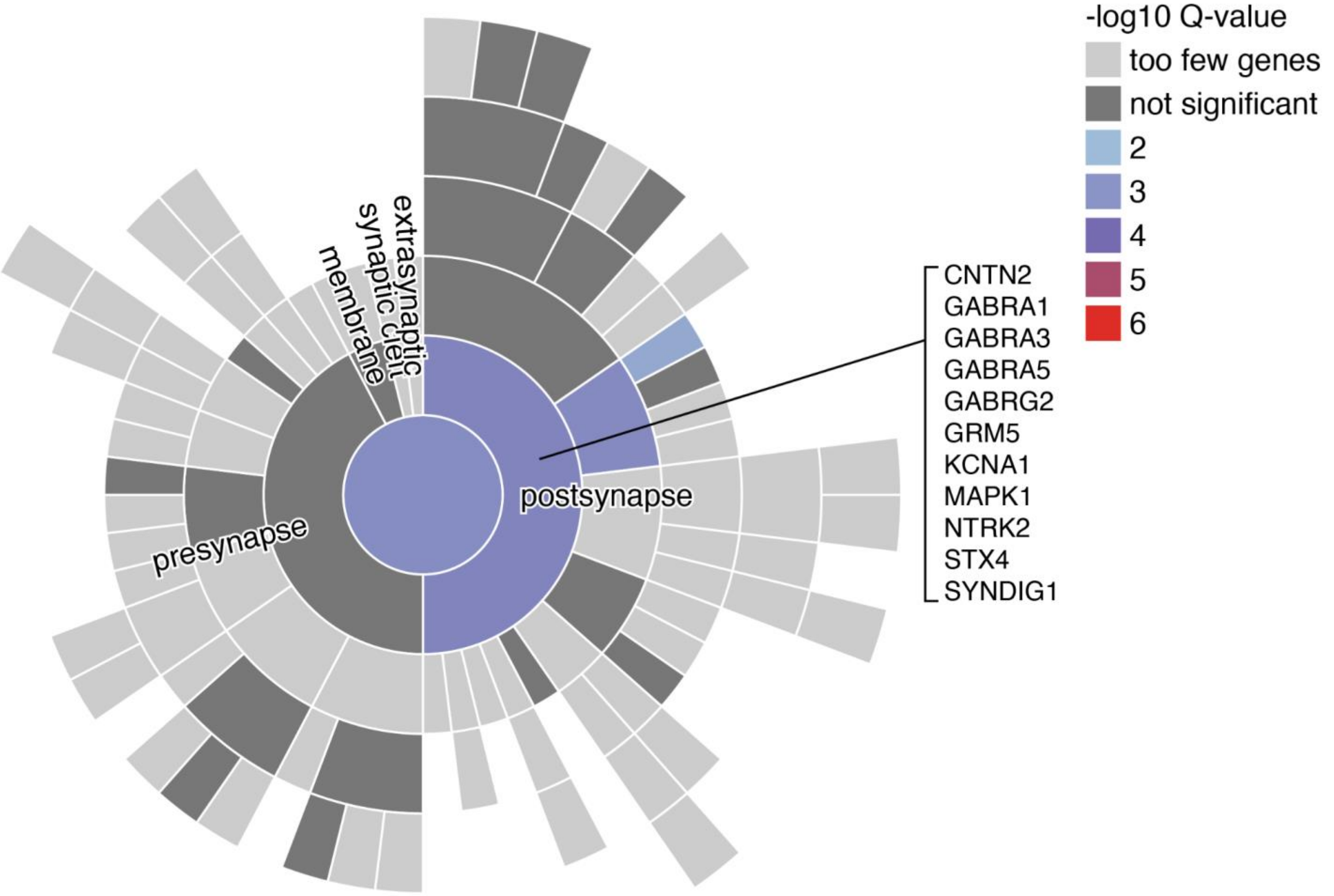

**Figure S34**. **GO-BP enrichment treemap highlighting neuronal-related biological processes in the SPG defined neuronal microenvironments**. GO-ORA analysis (GO-BP, FDR ≤ 0.05) was performed on neuronal microenvironment-restricted SCZ-DEGs identified at nominal $p < 0.05$. The top 50 neuron-associated enriched GO-BP terms (similarity threshold=0.7) were summarized and visualized as a treemap, supporting enrichment of neuronal signatures within these microenvironment-restricted SCZ-DEGs. Tile size reflects enrichment significance ($-\log_{10}$ (adjusted $p$-value)).

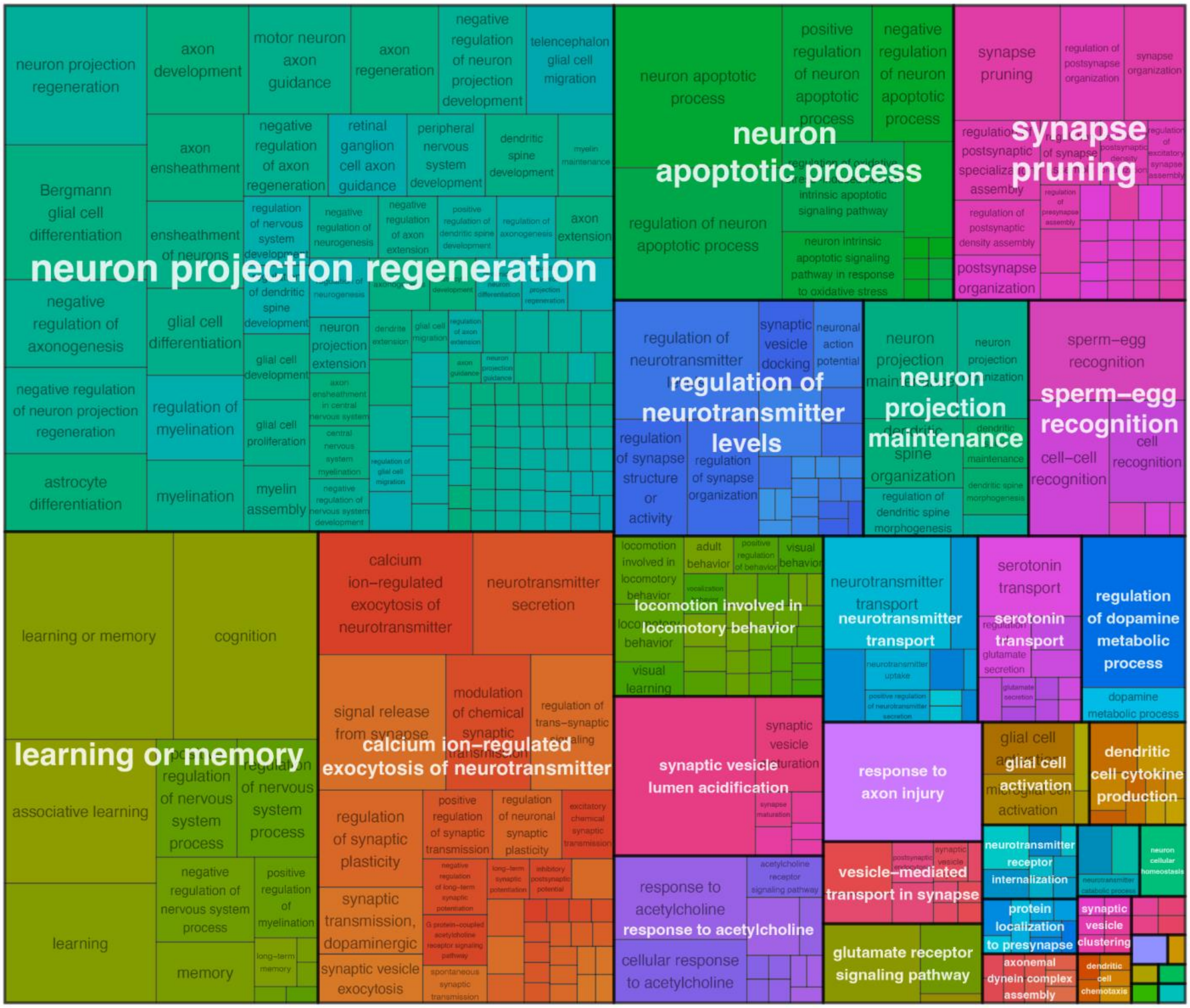

**Figure S35**. **Heatmap and bar plot summarizing the representation of key biological processes across microenvironments, focusing on the top 5 clusters (Clusters 1-5) identified from STRING-based protein-protein interaction (PPI) networks using Markov clustering (MCL)**. The bar plot (top) shows the gene share (%) of microenvironment-restricted SCZ-DEGs (nominal $p < 0.05$) contributing to three core cellular processes, 1) energy metabolism & mitochondrial function, 2) protein homeostasis, and 3) RNA processing & metabolism, calculated relative to the total number of genes aggregated across the top five clusters within each microenvironment. The heatmap (bottom) displays the total number of microenvironment-restricted SCZ-DEGs assigned to each biological category, with darker blue indicating higher gene counts. All analyses were performed separately for each microenvironment (neuropil, PNN, neuronal, vasculature).

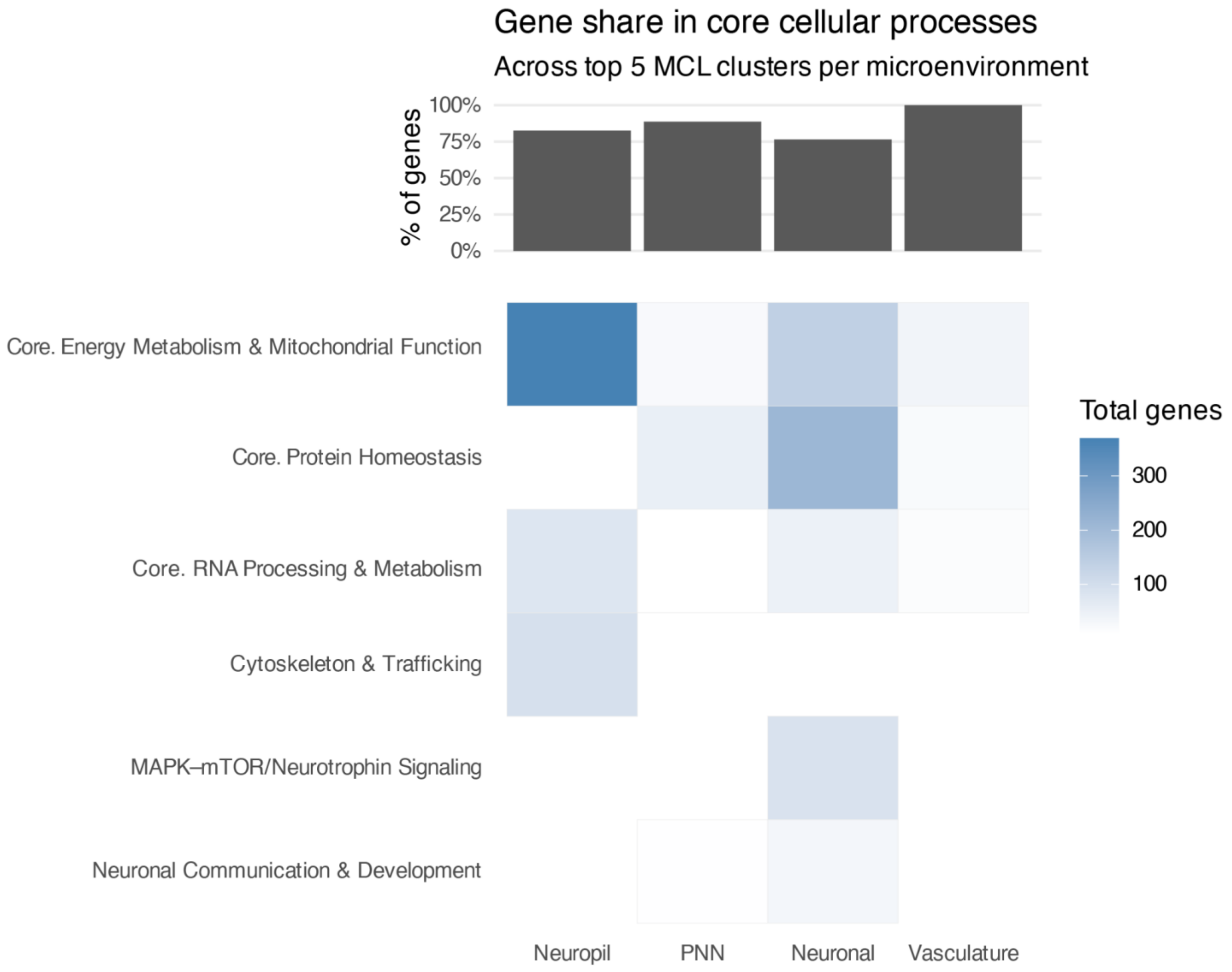

**Figure S36**. **UpSet plot summarizing the intersection of microenvironment-restricted SCZ-DEGs identified within four SPG-defined microenvironments (neuropil**, **neuronal**, **PNN**, **vasculature)**. The bar plot (top) indicates the number of microenvironment-restricted SCZ-DEGs that are uniquely or jointly identified across microenvironments, ordered by intersection size. Each black dot and connecting line (bottom) specifies the particular combination of microenvironments included in the intersection. Horizontal bars on the left indicate the total number of SCZ-DEGs identified within each microenvironment at nominal $p < 0.05$. All intersections and individual gene lists are provided in **Supplementary Table 10**.

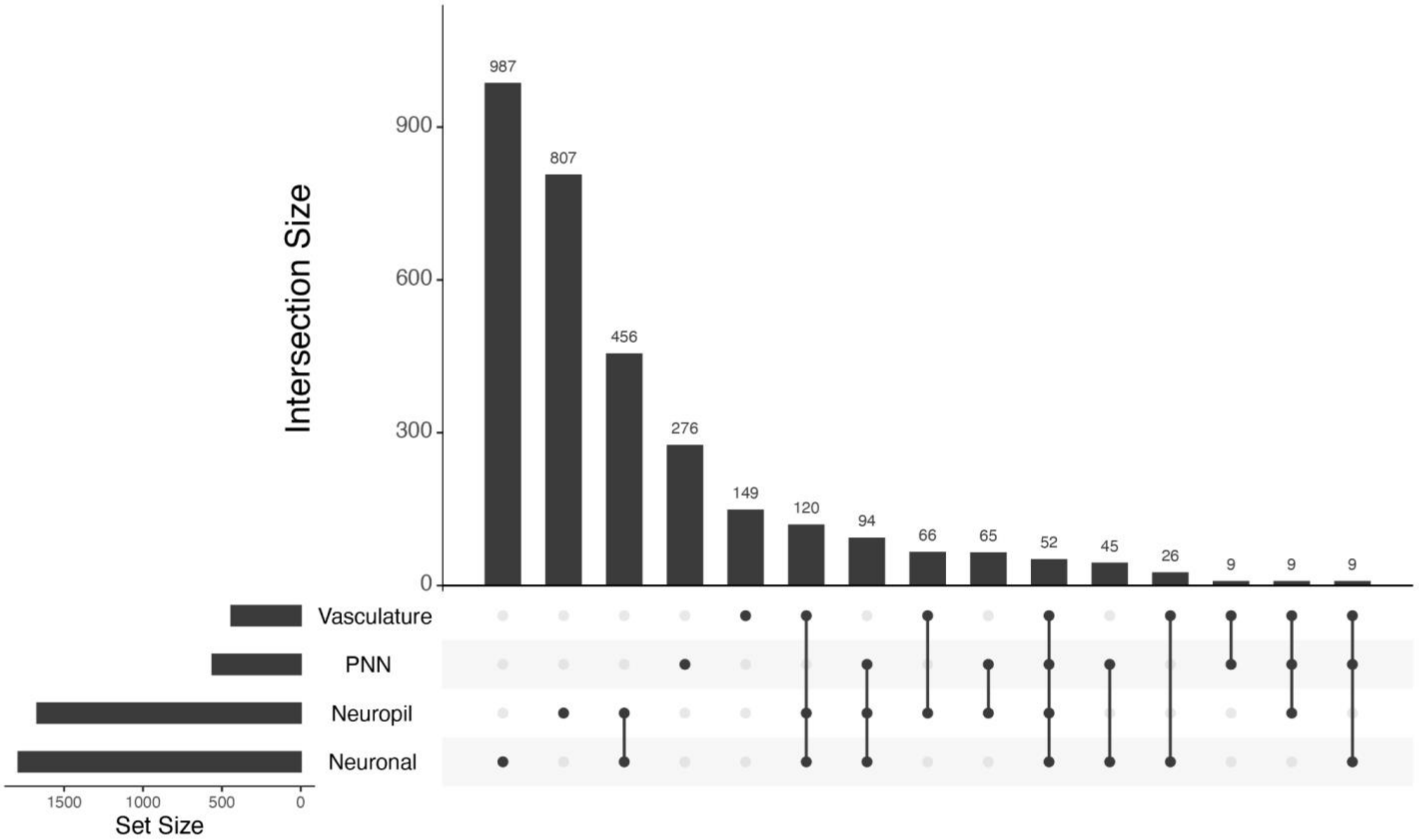

**Figure S37**. **Treemap representation of GO-ORA, showing enriched BP terms among microenvironment-restricted SCZ-DEGs shared across two or more microenvironments**. Non-shared microenvironment-restricted SCZ-DEGs identified in only one microenvironment at nominal $p < 0.05$ were excluded prior to this analysis. Each rectangle represents a significantly enriched GO-BP term (FDR ≤ 0.05), with related terms grouped by color and semantic similarity. Rectangle size corresponds to the statistical significance ($-\log_{10}$ (adjusted $p$-value)). The treemap displays the top 50 representative terms after redundancy reduction.

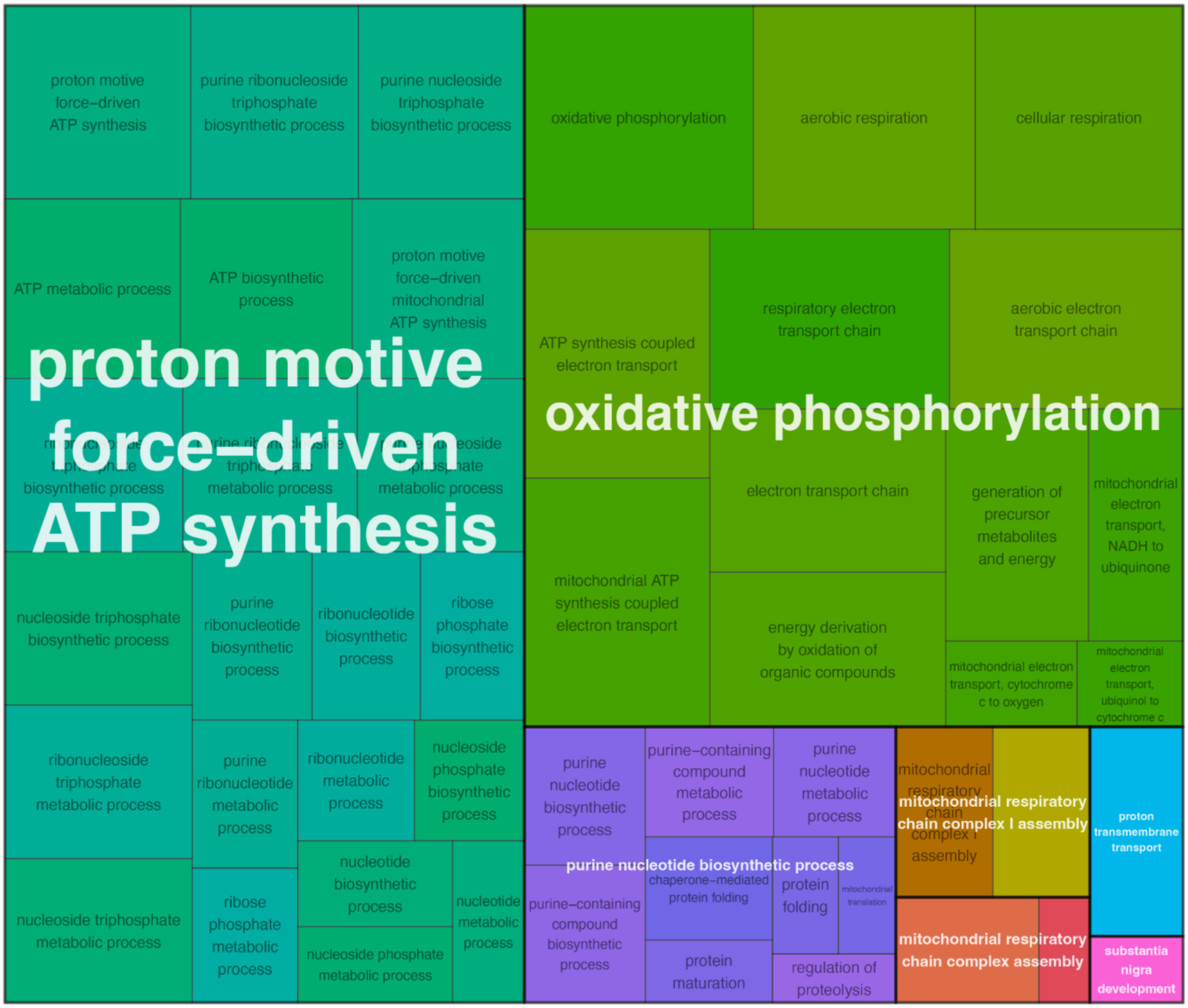

**Figure S38**. **Functional characterization of selected STRING-based protein-protein interaction (PPI) network modules (MCL clusters), using STRING built-in GO-BP enrichment analyses**. Each dot plot illustrates significantly enriched GO-BP terms (FDR ≤ 0.05) within a given module, with enrichment signal. Dot size corresponds to the number of module genes associated with each GO term, and dot color indicates the adjusted *p*-value. Enrichment analyses were performed separately for each microenvironment-restricted MCL module using a microenvironment-specific matching background gene set (**Methods**). The results provide additional biological context and functional interpretation of the representative signaling subnetworks highlighted in **Figure 6E**.

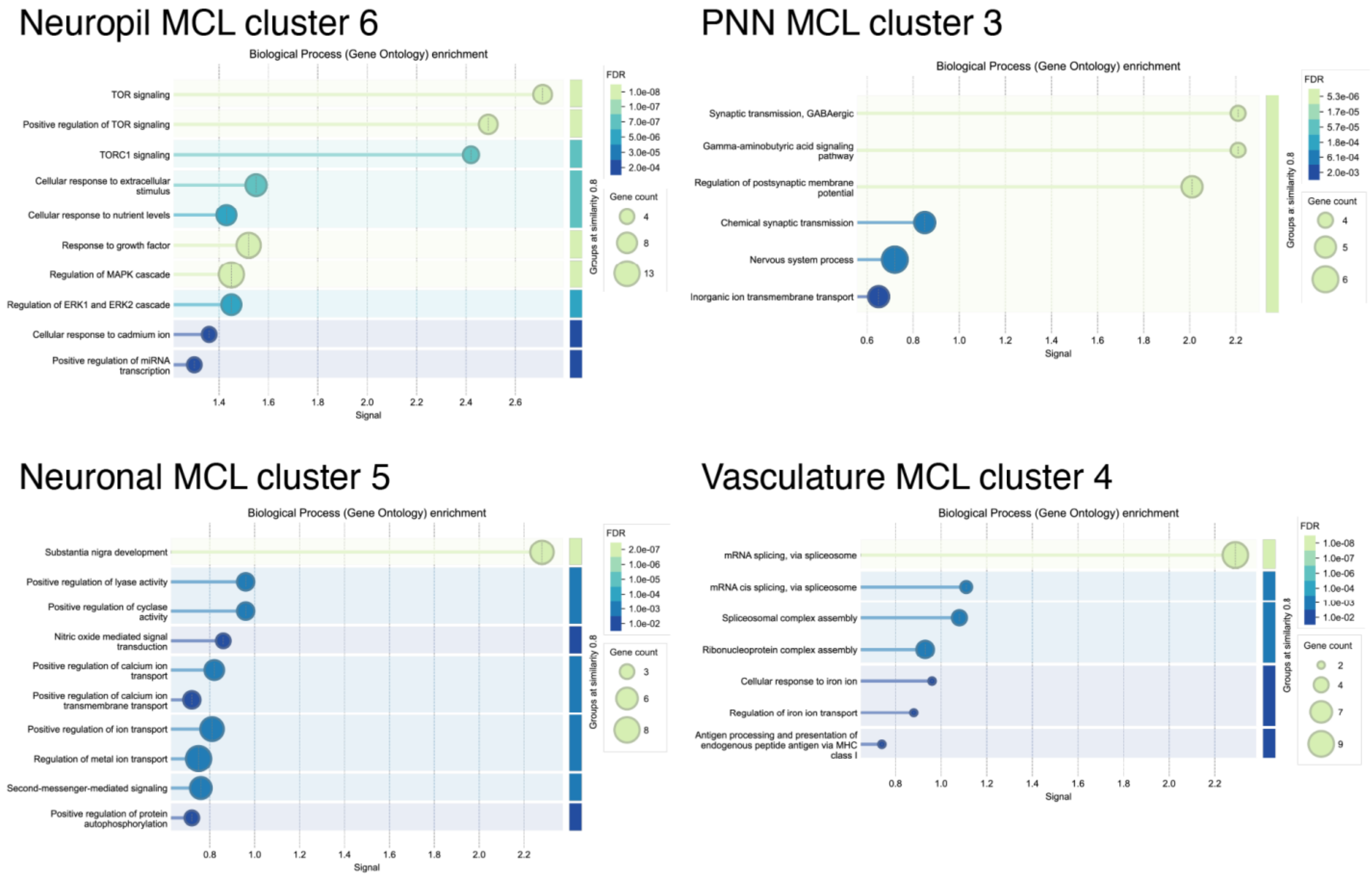

**Figure S39**. **Schematic illustration of the conceptual working model exploring LR pairs altered in SCZ using LR pairs curated from a snRNA-seq study**. The diagram provides a conceptual overview of how SCZ-linked candidate LR pairs from Huuki-Myers et al.[25] were mapped onto microenvironment-restricted SCZ-DEGs identified from our Visium-SPG data. For each microenvironment (neuropil, neuronal, PNN, and vasculature), one representative LR pair is highlighted, where the gene encoding the ligand or the receptor was detected as a statistically significant microenvironment-restricted SCZ-DEG (red: up-regulation; blue: down-regulation of gene expression in SCZ). A cyan dashed circle represents a single Visium spot of the SPG-defined microenvironments. Created in BioRender. Kwon, S. H. (2026) https://BioRender.com/r7yopwt

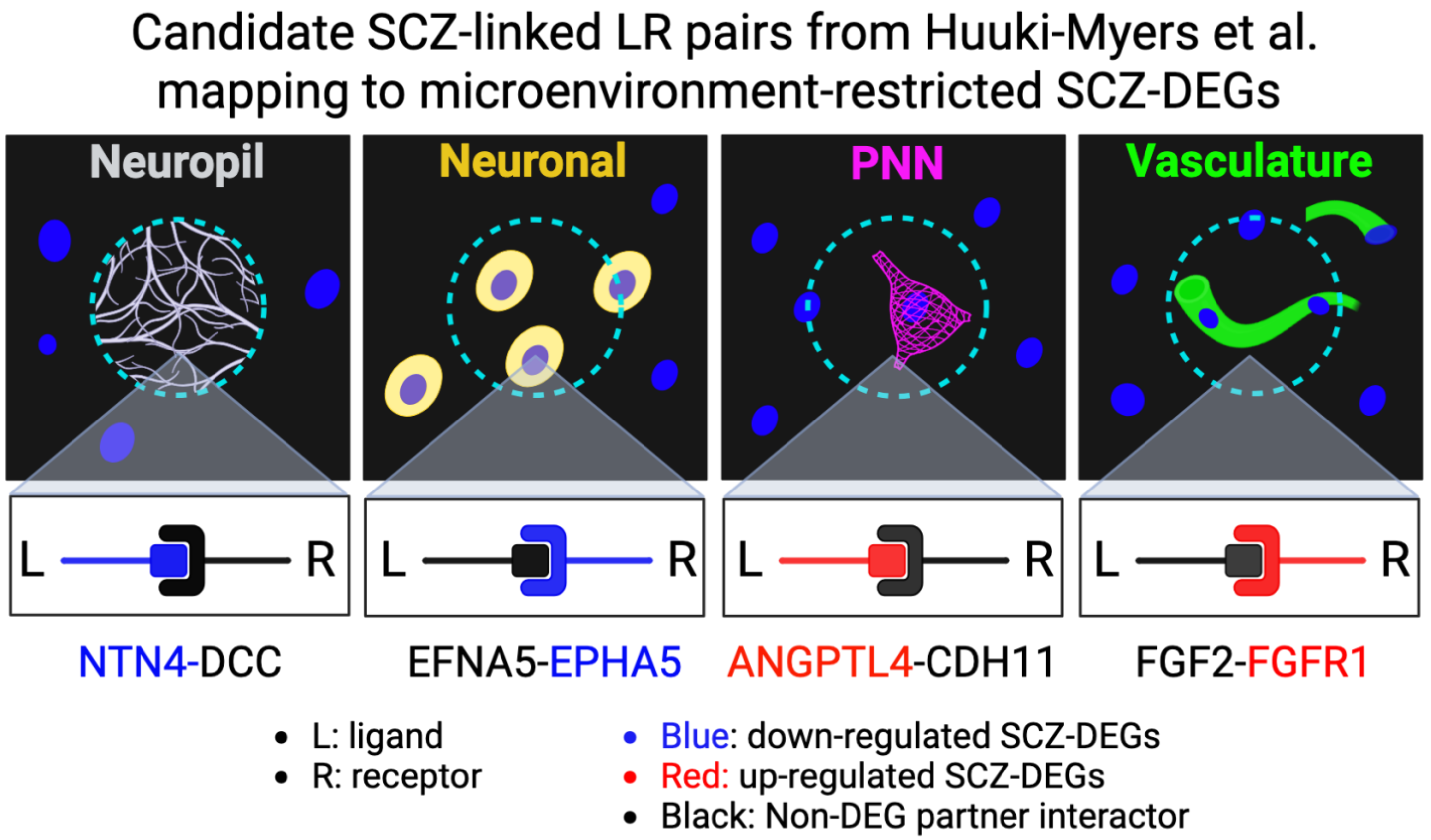

**Figure S40**. **Spot plots of SCZ-linked candidate LR pairs mapping to microenvironment-restricted SCZ-DEGs across SPG-defined microenvironments**. Representative LR pairs linked to microenvironment-restricted SCZ-DEGs are shown across four SPG-defined microenvironments (Br8667_NTC and Br5973_SCZ). Visium spots co-expressing both components of an LR pair are marked in red; ligand-only spots in purple; receptor-only spots in skyblue; and spots expressing neither of the components in gray. SPG microenvironment spots are outlined in black.

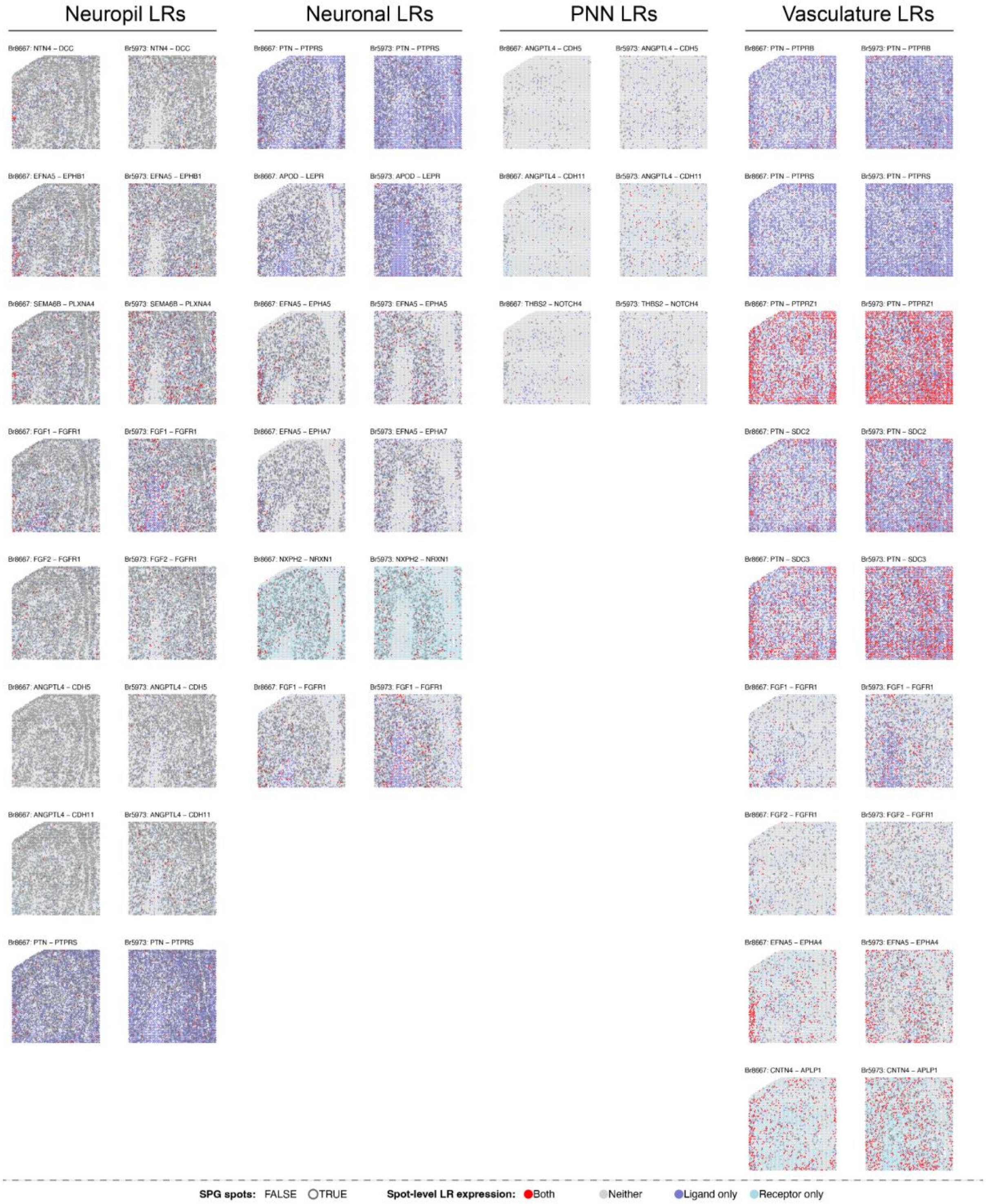

**Figure S41**. **Genotype-associated expression of *MAPK3* across neuronal and vascular microenvironments**. (**A**) Venn diagram illustrating the overlap between spatial eQTLs (eGenes), layer-adjusted neuropil SCZ-associated DEGs (SCZ-DEGs), and SCZ GWAS loci (GWAS). We highlight *MAPK3* as a candidate gene supported by convergent genetic association, genotype-dependent regulation, and disease-associated expression changes. (**B**) Box plots show genotype-associated expression patterns of *MAPK3* (rs7542) across neuronal (Neun, left) and vascular (Vasc, right) microenvironments. Consistent genotype-associated expression patterns were observed across neuronal and vascular microenvironments following post hoc correction of *p*-values (FDR < 0.05), suggesting that the genotype-dependent regulatory effect observed in neuropil may extend to additional microenvironments. Diagnosis (Dx) groups are colored: blue for NTC and red for SCZ.

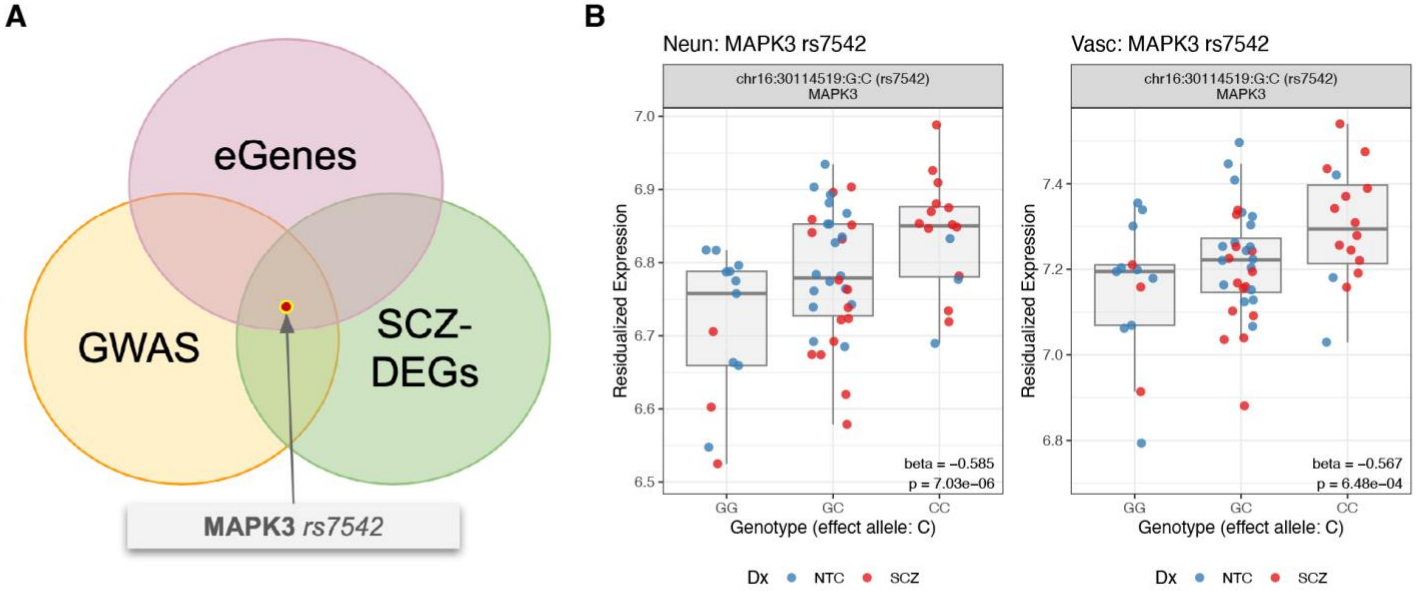

**Figure S42**. **Genotype-associated expression of *LY6H* across dlPFC layers and microenvironments**. Box plots show genotype-associated expression of *LY6H* highlighting two strong colocalization signals: (**A**) rs7830479 across selective dlPFC layers: SpD02-L3/4 (left) and SpD05-L5 (right) and (**B**) microenvironments: neuropil (left) and neuronal (right). Diagnosis (Dx) groups are colored: blue for NTC and red for SCZ.

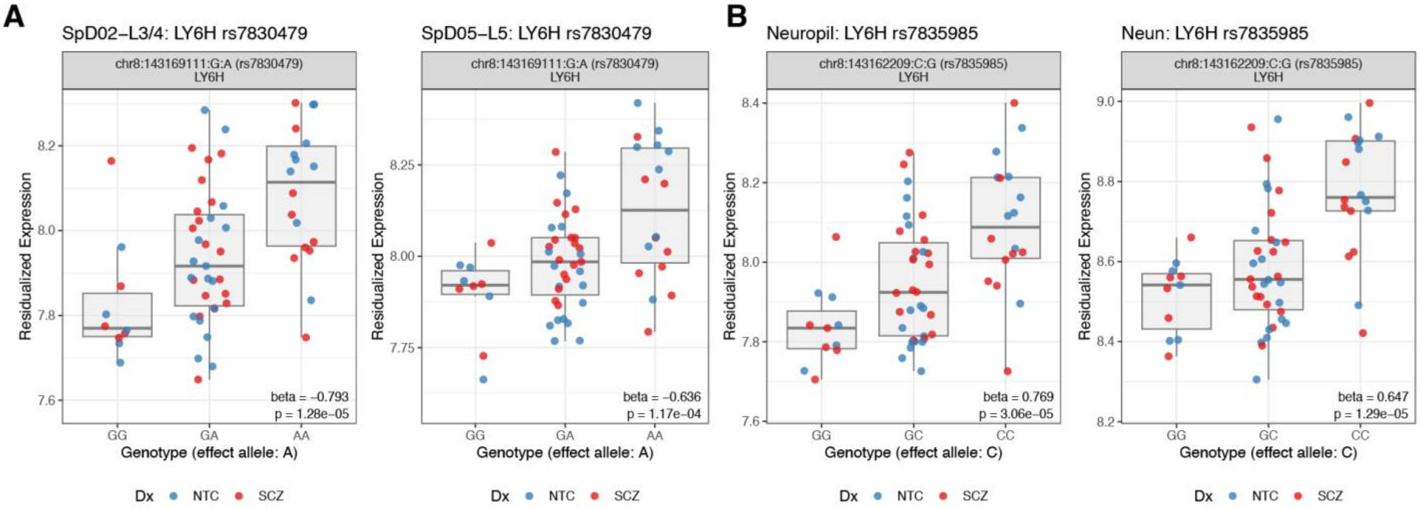

**Figure S43**. **Comparison of dlPFC SCZ colocalization genes with previously published datasets.** Venn diagram showing the overlap between SCZ colocalization genes identified in the current spatial dlPFC analysis, habenula eQTL data,[125] and external brain xQTL/GWAS-based gene-prioritization resources[130–132]. 11 colocalization genes were uniquely detected in the spatial dlPFC analysis, highlighting signals not captured by previous datasets. These 11 genes are listed in **Supplementary Table 13** (coloc_genes_unique_vs_external sheet).

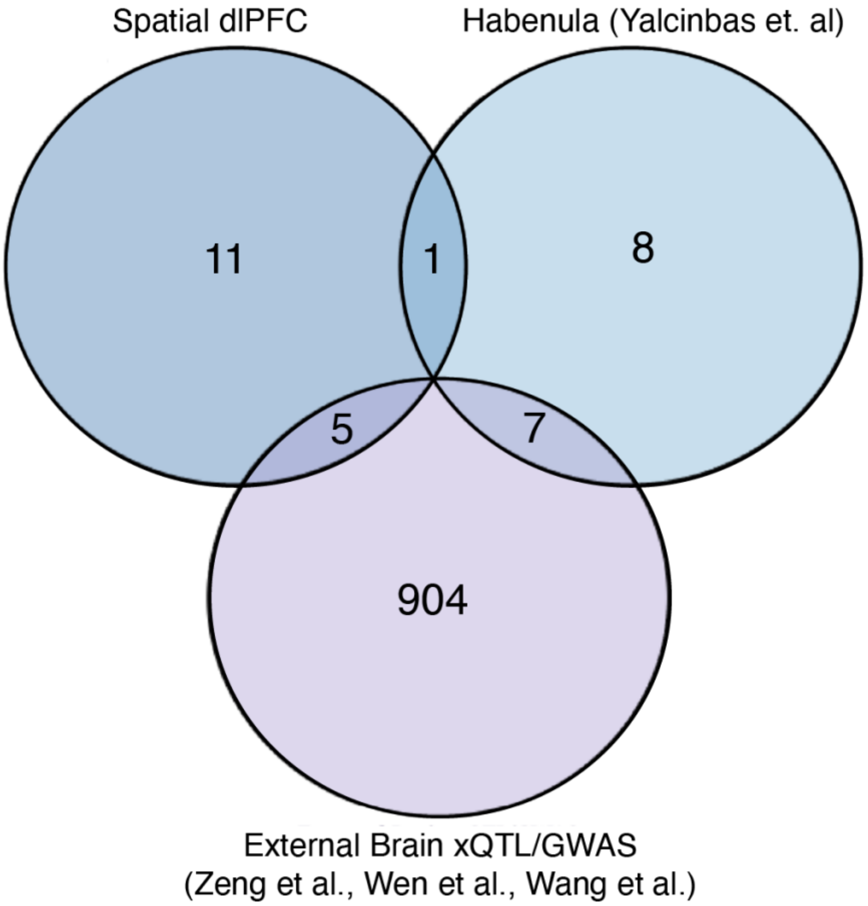

### Supplementary Tables

**Supplementary Table 1**. Demographic information and descriptive statistics of donors included in the Visium and Xenium SRT experiments (sex, age, RNA integrity number [RIN], and postmortem interval[PMI]).
